# Targeted Protein Degradation through E2 Recruitment

**DOI:** 10.1101/2022.12.19.520812

**Authors:** Nafsika Forte, Dustin Dovala, Matthew J. Hesse, Jeffrey M. McKenna, John A. Tallarico, Markus Schirle, Daniel K. Nomura

## Abstract

Targeted protein degradation (TPD) with Proteolysis Targeting Chimeras (PROTACs), heterobifunctional compounds consisting of protein targeting ligands linked to recruiters of E3 ubiquitin ligases, has arisen as a powerful therapeutic modality to induce the proximity of target proteins with E3 ligases to ubiquitinate and degrade specific proteins in cells. Thus far, PROTACs have primarily exploited the recruitment of E3 ubiquitin ligases or their substrate adapter proteins but have not exploited the recruitment of more core components of the ubiquitin-proteasome system (UPS). In this study, we used covalent chemoproteomic approaches to discover a covalent recruiter against the E2 ubiquitin conjugating enzyme UBE2D—EN67—that targets an allosteric cysteine, C111, without affecting the enzymatic activity of the protein. We demonstrated that this UBE2D recruiter could be used in heterobifunctional degraders to degrade neo-substrate targets in a UBE2D-dependent manner, including BRD4 and the androgen receptor. Overall, our data highlight the potential for the recruitment of core components of the UPS machinery, such as E2 ubiquitin conjugating enzymes, for TPD, and underscore the utility of covalent chemoproteomic strategies for identifying novel recruiters for additional components of the UPS.

## Introduction

Targeted protein degradation (TPD) using Proteolysis Targeting Chimeras (PROTACs) or molecular glues has arisen as a powerful therapeutic modality for chemically inducing the proximity of E3 ubiquitin ligases with neo-substrate target proteins to ubiquitinate and degrade specific proteins through the proteasome ^1,2^. Despite the wide-spread development of PROTACs for many target proteins, most PROTACs have utilized recruiters against only a select few E3 ligases or substrate receptors despite the existence of >600 protein factors within the ubiquitin-proteasome system (UPS) ^3^. Most PROTACs have employed recruiters against cereblon and VHL, with additional utilization of MDM2, cIAP, and DCAF15 recruiters ^3^. Covalent chemoproteomic approaches have also revealed the ligandability of proteins within the UPS machinery and have led to the discovery of several additional recruiters against DCAF16, DCAF11, DCAF1, RNF114, RNF4, and FEM1B ^3–11^. However, these recruiters thus far have all been against either E3 ligases or the substrate receptors of Cullin E3 ligase complexes and have not exploited more core and shared components of the UPS system, such as E2 ubiquitin-conjugating enzymes or commonly shared adapter proteins in Cullin E3 ligases (e.g. DDB1 or SKP1).

Recent discoveries of molecular glue degraders and their mechanisms reveal the conceptual possibility of exploiting these core UPS components in PROTACs. The CDK inhibitor CR8 was found to act as a molecular glue degrader that forms a ternary complex between the CUL4 adaptor protein DDB1 and the CDK12-cyclin K complex to induce the ubiquitination and degradation of cyclin K ^12^. HQ461 was also found to promote ternary complex formation between CDK12 and DDB1 as well to induce the ubiquitination and degradation of cyclin K^13^. Our group also recently discovered the anti-cancer covalent ligand EN450, which recognizes the allosteric C111 on UBE2D conferring recognition for the transcription factor NFKB1--leading to its ubiquitination and proteasome-dependent degradation ^14^. While these discoveries were all with molecular glue degraders and not PROTACs, they speak to the possibility that these core components of the Cullin complex (adapter proteins and E2s) can be recruited with small-molecules to form ternary complexes with neo-substrate proteins to induce their ubiquitination and degradation. Recruitment of these core components of the UPS system which are essential genes for many Cullin E3 ligase complexes and also for cell viability may enable degradation of a broader or different scope of neo-substrates and potentially avoid resistance mechanisms to PROTACs that exploit nonessential E3 ligase substrate receptors, such as cereblon^15,16^.

In this study, we sought to determine whether E2 ubiquitin conjugating enzymes can be recruited for heterobifunctional PROTAC applications. E2 ubiquitin conjugating enzymes accept ubiquitin from the E1 complex and catalyze its covalent attachment to other proteins ^17^. Using covalent chemoproteomic approaches, we discovered a covalent recruiter that targets the allosteric C111 on the E2 UBE2D and showed that this recruiter can be used in PROTACs to degrade neo-substrate proteins in a UBE2D-dependent manner.

## Results and Discussion

### Developing a Covalent Recruiter Against the E2 Ubiquitin Conjugating Enzyme UBE2D

Among the E2 ubiquitin conjugating enzymes, we prioritized efforts to discover covalent recruiters against the UBE2D family which consists of nearly identical isoforms UBE2D1, UBE2D2, UBE2D3, and UBE2D4 that all bear an active-site ubiquitin conjugating C85 as well as an allosteric cysteine C111, since we recently discovered a covalent molecular glue degrader EN450 that exploited this allosteric C111 without targeting the ubiquitin-conjugating C85 ^14,17^. This recent discovery pointed to the potential feasibility for recruitment of UBE2D for PROTACs. UBE2D1-4 are utilized as a ubiquitin-transfer enzymes for many different types of RING E3 ligases and Cullin E3 ligase complexes ^18,19^. Given that we previously showed that EN450 covalently targeted C111 on UBE2D, albeit not very potently with mid-micromolar affinity ^14^, we initially developed NF142, a PROTAC linking EN450 to the BET family inhibitor JQ1 through a C4 alkyl linker **(Figure S1A)**. NF142 dose-responsively competed against fluorophore-functionalized cysteine-reactive idoacetamide probe (IA-rhodamine) labeling of recombinant UBE2D2 C85S mutant protein at mid- to low-micromolar concentrations by gel-based activity-based protein profiling (ABPP) **(Figure S1B)**^10^. When treated in HEK293T cells, NF142 modestly degraded only the short isoform of BRD4 by ~70% without degrading the long BRD4 isoform **(Figure S1C-S1D)**.

Given that EN450 was discovered through a phenotypic screen for molecular glue degraders and not through a directed screen against UBE2D, we next screened a library of 569 cysteine-reactive acrylamide and chloroacetamide covalent ligands against recombinant human UBE2D2 C85S protein by gel-based ABPP competing the binding of covalent ligands against IA-rhodamine labeling **(Figure 1A; Table S1)**. Through this screen, we identified EN67 as the top hit that dose-responsively bound to UBE2D2 **(Figure 1B)**. Through reconstitution of TP53 ubiquitination activity by the E1 ubiquitin-activating enzyme, UBE2D2, MDM2, ubiquitin, and ATP, we confirmed that EN67 does not inhibit overall TP53 ubiquitination mediated by UBE2D2 and the whole ubiquitination machinery, suggesting the activity of UBE2D is not compromised by the EN67 covalent adduct **(Figure 1C)**. We further mapped the site of modification of the EN67 covalent adduct on pure recombinant human wild-type UBE2D2 through liquid-chromatography-tandem mass spectrometry (LC-MS/MS) analysis of resulting tryptic digests and demonstrated that EN67 selectively targeted C111 without targeting the catalytic C85 **(Figure 1D)**.

**Figure 1.**
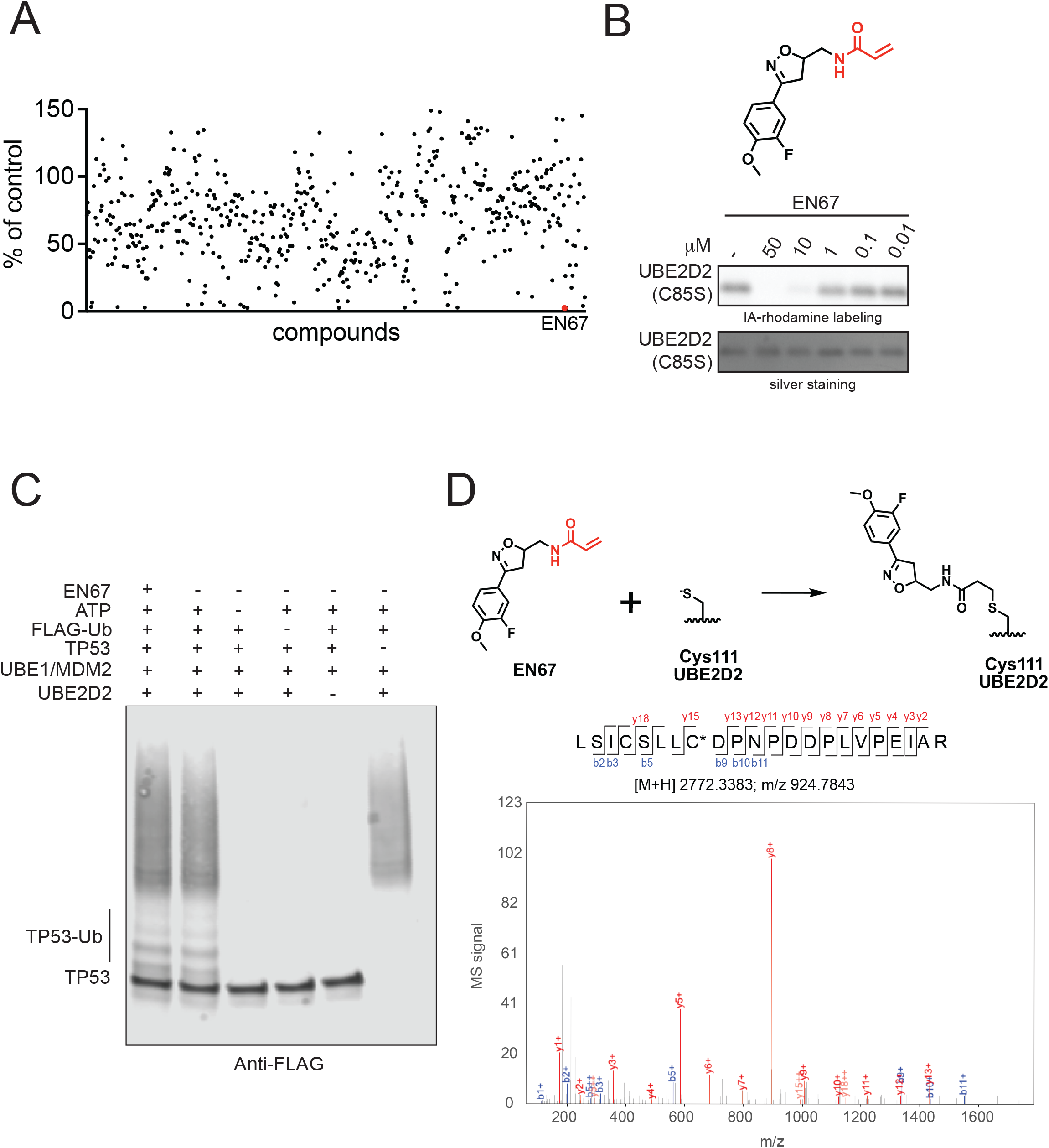
Discovering a covalent recruiter for E2 ubiquitin conjugating enzyme UBE2D. **(A)** Gel-based ABPP screen of cysteine-reactive covalent ligands against human pure UBE2D2 C85S protein showing EN67 as the top hit. **(B)** Structure of EN67 with the cysteine-reactive acrylamide warhead highlighted in red. Gel-based ABPP showing competition of EN67 against IA-rhodamine binding to UBE2D2 C85S pure protein and silver staining of protein showing equal protein loading. **(C)** TP53 ubiquitination activity by E1 UBE1, E2 UBE2D2, E3 MDM2, FLAG-ubiquitin, and ATP showing that EN67 does not inhibit TP53 ubiquitination activity. **(D)** Mapping of EN67 site of modification on human pure UBE2D2 C111 by LC-MS/MS. Experiments in **(B, C, D)** are representative of n=3 biological replicates/group.

### Assessing Target Engagement and Selectivity of UBE2D by EN67

We next sought to confirm target engagement and overall proteome-wide selectivity of EN67 for C111 of UBE2D in cells. We synthesized four alkyne-functionalized probes of EN67 with different spacer lengths (**Figure 2A**). NF363C exhibited the highest degree of binding to UBE2D2 (C85S) via gel-based ABPP **(Figure 2B)**. Using this alkyne-functionalized probe, we showed engagement and enrichment of UBE2D2 but not unrelated proteins such as GAPDH from treatment of this probe in HEK293T cells, followed by appending on a biotin-azide enrichment handle by copper-catalyzed azide-alkyne cycloaddition (CuAAC) in resulting lysates, avidin-enrichment, and blotting for specific targets **(Figure 2B-2C)**. To assess the proteome-wide selectivity, cysteine-reactivity, and degree of UBE2D2 engagement, we next performed isotopically labeled desthiobiotin tag-based ABPP (isoDTB-ABPP) competing *in situ* EN67 treatment in HEK293T cells against the alkyne-functionalized iodoacetamide (IA-alkyne) probe ^4,20,21^. Among 3798 probe-modified peptides detected and quantified across all 3 biological replicates, there were only 11 targets that showed control vs EN67-treated probe-modified peptide ratios greater than 1.3 with an adjusted p-value of less than 0.01, among which C111 of UBE2D was the only protein involved in the UPS, showing a ratio of 1.3 **(Figure 2D; Table S2)**. This indicates approximately 25 % target engagement of UBE2D in cells. The other targets—GATA6, HK2, WDSUB1, SON, AK6, GTSE1, PYGO2, ACAA2, ASXL2, and ANP32A—were not related to the UPS and not expected to interfere with validating EN67 as a UBE2D recruiter in TPD applications. We note that the tryptic peptide sequence bearing C111 detected and quantified by isoDTB-ABPP is identical across UBE2D1, UBE2D2, UBE2D3, and UBE2D4 and thus we cannot distinguish between them. Given the high degree of sequence identity between these UBE2D enzymes, we believe that EN67 likely targets all four isoforms of UBE2D. We deemed the relatively modest degree of engagement of UBE2D in cells to be acceptable given previous studies indicating that only fractional occupancy would likely be required to enable target degradation with heterobifunctional compounds that demonstrate sub-stoichiometric activity ^5,8,9,22^.

**Figure 2.**
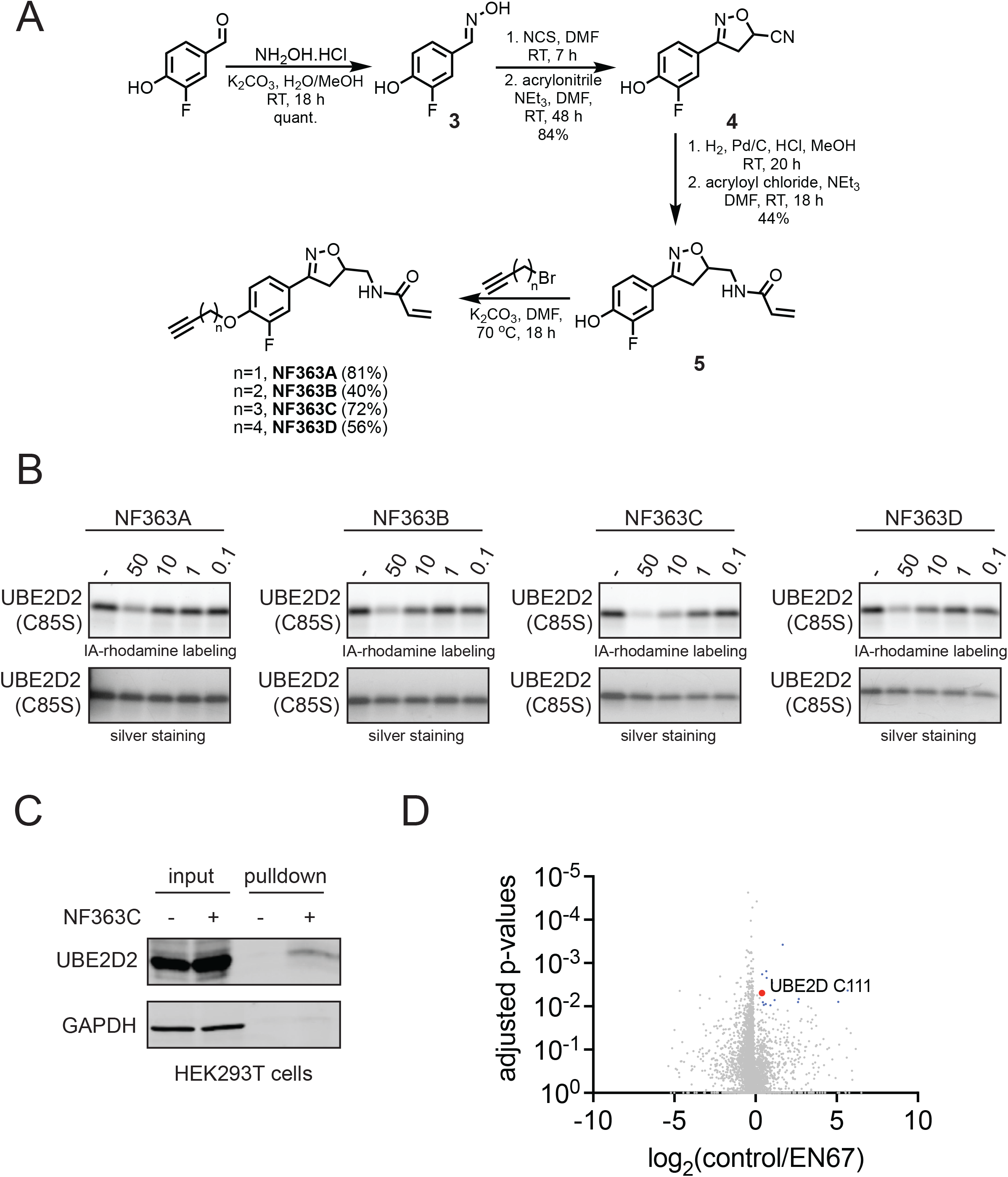
Validating target engagement and selectivity of EN67 in cells. **(A)** Synthetic route for making the alkyne-functionalized probe of EN67. **(B)** Gel-based ABPP of alkyne-functionalized EN67 probes against pure recombinant human UBE2D2 C85S protein and corresponding silver staining. **(C)** NF363C engagement and enrichment of UBE2D2 in HEK293T cells. Cells were treated with DMSO or NF36C (50 μM) for 24 h. Probe-modified proteins were subsequently appended with an azide-functionalized biotin handle by CuAAC, and proteins were avidin-enriched and eluted for detection of UBE2D2 and an unrelated negative control protein GAPDH by Western blotting. Both input and pulldown of UBE2D2 and GAPDH are shown. **(D)** isoDTB-ABPP cysteine chemoproteomic profiling of EN67 in HEK293T cells. HEK293T cells were treated with DMSO vehicle or EN67 (50 μM) for 4 h. Lysates were labeled with IA-alkyne (200 μM) for 1 h and isotopic desthiobiotin tags were appended by CuAAC and taken through the isoDTB-ABPP procedure. Shown are ratios of control/EN67-treated probe-modified peptide ratios and adjusted p-values from n=3 biological replicates/group. Data are shown in **Table S2**. Data in **(B, C)** is representative from n=3 biological replicates/group.

### Testing EN67 as a UBE2D Recruiter for PROTAC Applications

Having shown that EN67 covalently targets C111 relatively selectively in cells, at least among the UPS, we next sought to demonstrate its utility in PROTACs to degrade neo-substrate proteins. We synthesized five PROTACs linking the EN67 UBE2D recruiter to the BET family inhibitor JQ1 with either a C2, C4, C5, C7 or PEG3 linker—NF129, NF90, NF369, NF370, and NF91, respectively **(Figure 3A; Figure S2A)**. These degraders still exhibited mid-micromolar potency against UBE2D2 C85S by gel-based ABPP **(Figure 3B; Figure S2B-S2D)**. We tested these degraders in HEK293T or MDA-MB-231 breast cancer cells. Only NF90, among the five degraders, demonstrated degradation of only the short, but not long, isoform of BRD4 in both HEK293T and MDA-MB-231 cells **(Figure 3C-3D, Figure S2E-S2H; Figure S3A-S3B)**. NF90 degraded BRD4 starting around 6 h of treatment with progressively increasing degradation through 24 h **(Figure S3C)**. We further demonstrated that the BRD4 degradation by NF90 was attenuated by pre-treatment of MDA-MB-231 cells with the proteasome inhibitor bortezomib or the NEDDylation inhibitor MLN4924 **(Figure 3E)**. These latter data indicate that the observed degradation of the BRD4 short isoform is occurring through UBE2D in complex with a Cullin E3 ligase complex, rather than with another class of E3 ligases (e.g. RING), and suggests that UBE2D is not sufficient to drive degradation of targeted proteins alone.

**Figure 3.**
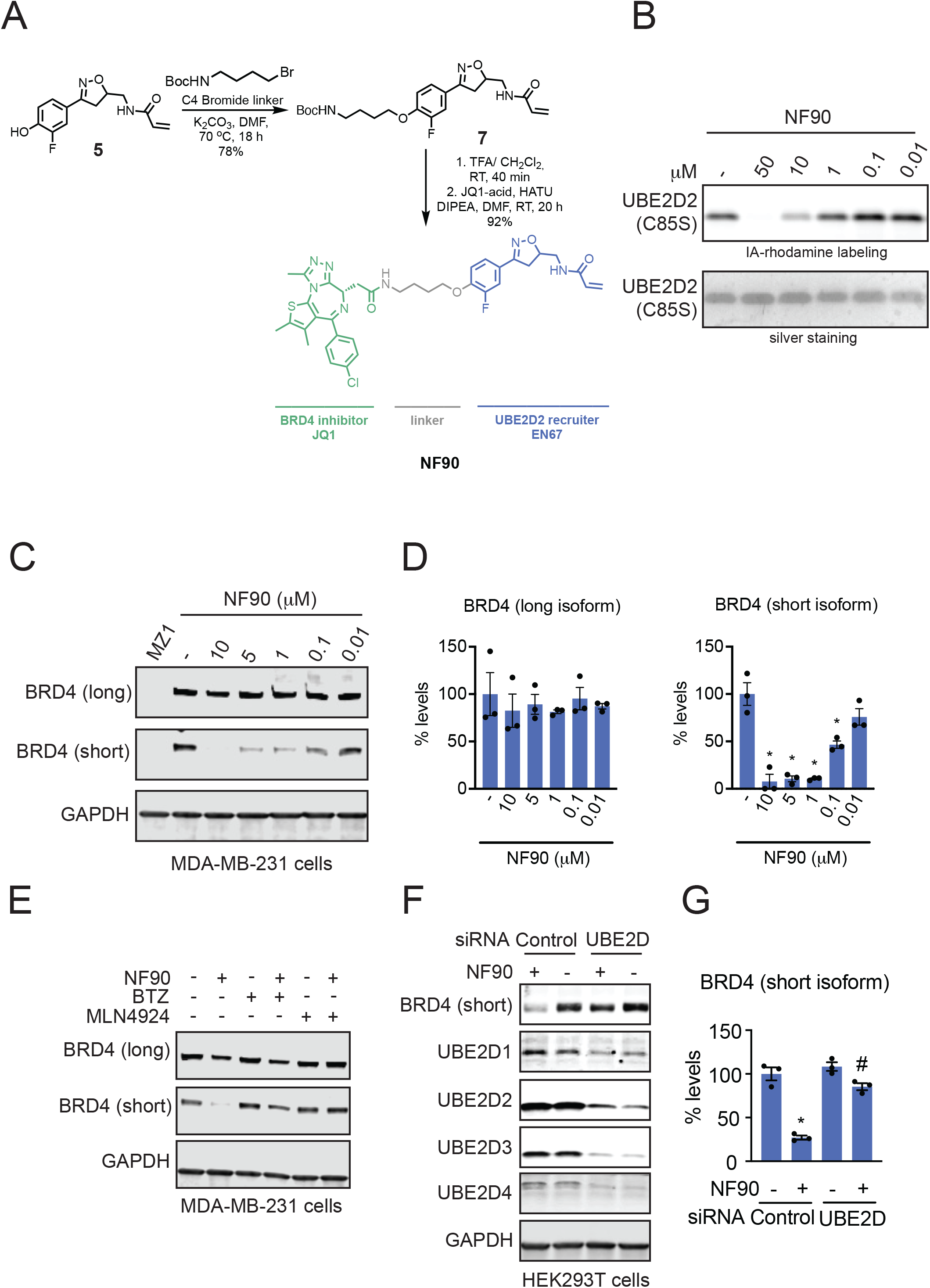
UBE2D-Based BRD4 Degrader. **(A)** Synthetic route and structure for NF90 linking UBE2D recruiter EN67 to BRD4 inhibitor JQ1. **(B)** gel-based ABPP analysis of NF90 against IA-rhodamine labeling of pure human UBE2D2 C85S protein and corresponding silver staining. **(C)** NF90 effects on BRD4 levels in MDA-MB-231 breast cancer cells. MDA-MB-231 cells were treated with DMSO vehicle or NF90 for 24 h and BRD4 and loading control GAPDH levels were assessed by Western blotting. **(D)** BRD4 levels from **(C)** quantified. **(E)** Attenuation of BRD4 degradation by proteasome inhibitor bortezomib (BTZ) or NEDDylation inhibitor (MLN4924). MDA-MB-231 cells were pre-treated with DMSO vehicle, BTZ (1 μM), or MLN4924 (1 μM) for 1 h prior to treatment of cells with DMSO vehicle or NF90 (10 μM) for 24 h. BRD4 and loading control GAPDH levels were assessed by Western blotting. **(F)** NF90 effects on BRD4 degradation upon UBE2D knockdown. HEK293T cells with siControl or siRNA knockdown of UBE2D1, UBE2D2, UBE2D3, and UBE2D4 treated with DMSO vehicle or NF90 (10 μM) for 24 h. BRD4 short isoform and UBE2D1-4 and loading control GAPDH were assessed by Western blotting. **(G)** BRD4 levels quantified from **(F)**. Gels and blots shown in **(B, C, E, F)** are representative from n=3 biological replicates/group. Bar graphs shown in **(D, G)** show average ±sem with individual replicate values. Statistical significance compared to vehicle-treated control expressed as *p<0.05 and compared to NF90-treated siControl cells as #p<0.05.

To confirm that our observed effects were on-target, we showed that the degradation of the BRD4 short isoform was completely attenuated upon knockdown of all four UBE2D1, UBE2D2, UBE2D3, and UBE2D4 enzymes **(Figure 3F-3G)**. We also synthesized a non-reactive analog of NF90 with NF457 expecting this degrader to not show UBE2D binding or target degradation. Unexpectedly, we still observed binding of this nonreactive PROTAC to UBE2D2 C85S protein by gel-based ABPP and degradation of the BRD4 short isoform in MDA-MB-231 cells, albeit to a lesser degree than NF90 **(Figure S4A-S4C)**. These data indicate that EN67 has significant degree of reversible binding to UBE2D beyond the reactivity introduced by the acrylamide warhead. Because NF90 only degraded one isoform of BRD4, we did not perform quantitative proteomic studies given that we likely not observe high degree of total BRD4 loss. The observed selective degradation of only the short isoform of BRD4 is particularly interesting given previous studies that showed tumor suppressive roles of the long isoform of BRD4 and the oncogenic roles of the short BRD4 isoform ^23^.

To demonstrate that our EN67 UBE2D recruiter was capable of degrading additional targets beyond BRD4, we next deployed this recruiter to generate androgen receptor (AR) degraders. We linked our EN67 recruiter onto an analog of the AR-targeting ligand from the Arvinas AR PROTAC ARV-110 via C4, C5, and C7 alkyl linkers to generate NF500A, NF500B, and NF500C, respectively **(Figure 4A; Figure S5A)**. We tested these PROTACs alongside the AR-targeting ligand control (NF505) as well as the ARV-110 PROTAC in AR-positive LNCaP prostate cancer cells. While all three PROTACs degraded AR, NF500C with the C7 alkyl linker showed the highest degree of AR degradation **(Figure 4B; Figure S5B-S5C)**. We further demonstrated that this loss of AR was attenuated by a NEDDylation inhibitor MLN4924 that inhibits Cullin E3 ligases, albeit MLN4924 treatment alone appeared to lower AR levels **(Figure 4C)**. To assess the selectivity of AR degradation, we performed tandem mass tagging (TMT)-based quantitative proteomic profiling of NF500C in LNCaP cells. We showed that AR was the only target degraded by >2-fold with adjusted p-value less than 0.001 **(Figure 4D; Table S3)**. We also generated a non-reactive analog of NF500C, NF534, and showed that this non-reactive PROTAC still degraded AR to an equivalent degree as NF500C **(Figure S6A-S6B)**.

**Figure 4.**
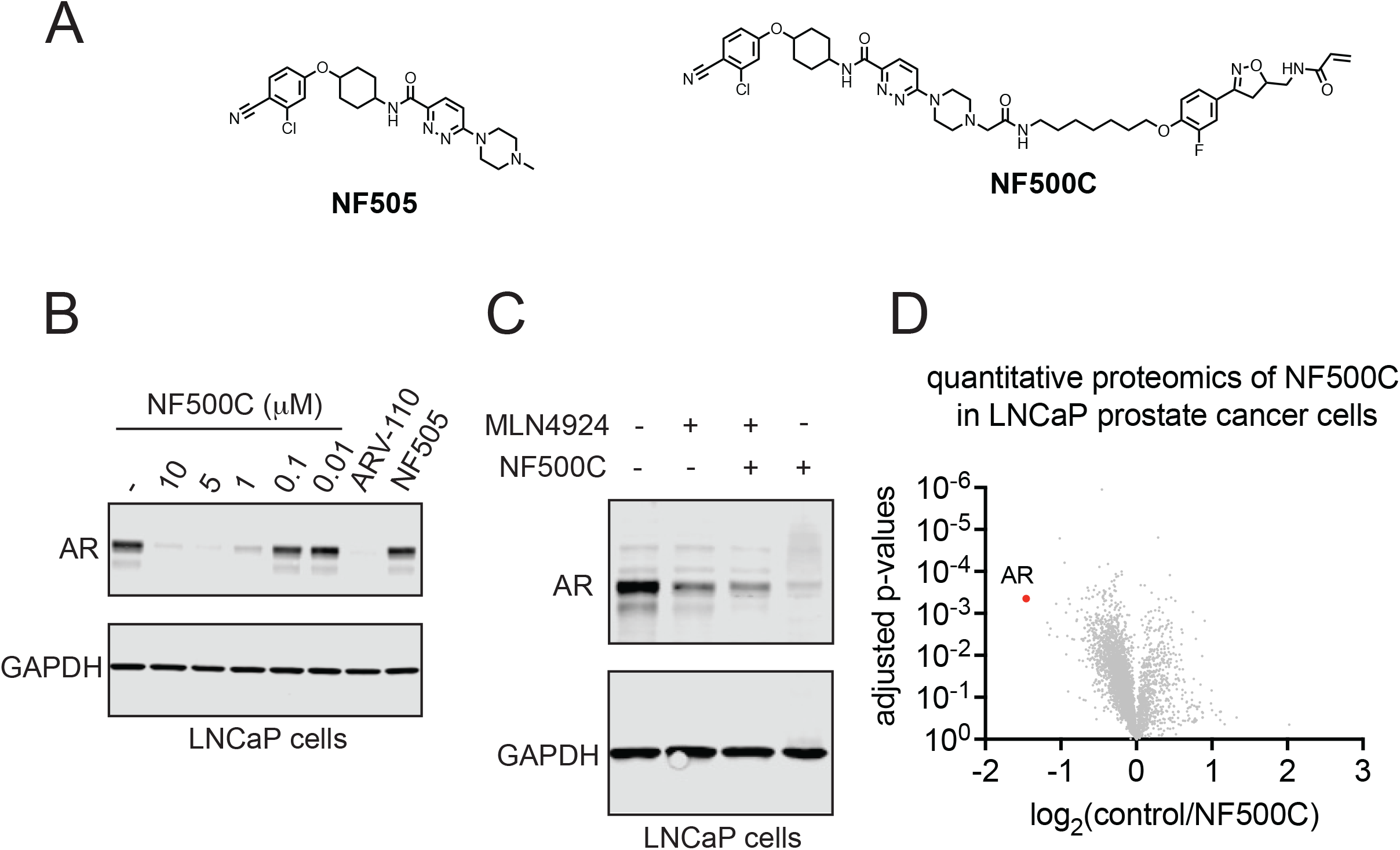
UBE2D-Based AR Degrader. **(A)** Structure of AR-targeting ligand control NF505 and UBE2D-based AR degrader NF500C linking the AR-targeting ligand to EN67. **(B)** AR degradation by NF500C. LNCaP prostate cancer cells were treated with DMSO vehicle or NF500C for 24 h and AR and loading control GAPDH levels were assessed by Western blotting. **(C)** AR degradation attenuated by NEDDylation inhibitor MLN4924. LNCaP cells were pre-treated with DMSO vehicle or NEDDylation inhibitor MLN4924 (1 μM) for 1 h prior to treating cells with DMSO vehicle or NF500C (10 μM) for 24 h. AR and loading control GAPDH levels were assessed by Western blotting. **(D)** TMT-based quantitative proteomic profiling of protein level changes conferred by NF500C treatment in LNCaP cells. LNCaP cells were treated with DMSO vehicle or NF500C (10 μM) for 24 h. Data are from n=3 biological replicates/group. Gels and blots in **(B, C)** are representative from n=3 biological replicates/group.

## Conclusions

In this study, we report the discovery of a covalent recruiter for an E2 conjugating enzyme UBE2D that can be used in heterobifunctional PROTACs to degrade neo-substrates, including BRD4 and AR. We had initially expected that the substrate scope might be broader for an E2 recruiter than an E3 ligase substrate receptor recruiter. However, we observed an even more specific substrate scope with BRD4 where we observed isoform-specific degradation of the short, but the not the longer isoform of BRD4. These data are particularly interesting given previous studies demonstrating the opposing tumor suppressive and oncogenic roles of the long versus short isoforms of BRD4, respectively ^23^. Our selective short isoform-specific BRD4 degrader may potentially eliminate the oncogenic roles of BRD4 while maintaining its tumor suppressive functions.

Attenuation of target degradation by NEDDylation inhibitors indicated that we required the NEDDylated Cullin complex to enable ubiquitination. However, we do not yet know whether recruitment of the E2 potentially bypasses the necessity for an E3 ligase substrate adapter. Formation of the ternary complex with the E2 in the Cullin complex versus the E3 ligase substrate receptor may restrict the geometry of the ternary complex for ubiquitination of the neosubstrate. Future structural elucidation of the Cullin complex with the protein targets may yield insight into the mechanism through which we are bringing together the E2 and the Cullin complex with the neosubstrates to facilitate ubiquitination.

Nonetheless, our study broadens the scope of UPS machinery that can be exploited for PROTACs to now include E2 ubiquitin conjugating enzymes and further highlights the utility of covalent chemoproteomic approaches in rapidly discovering allosteric covalent ligands that can be used as recruiters for induced proximity-based approaches.

## Acknowledgement

We thank the members of the Nomura Research Group and Novartis Institutes for BioMedical Research for critical reading of the manuscript. This work was supported by Novartis Institutes for BioMedical Research and the Novartis-Berkeley Translational Chemical Biology Institute (NB-TCBI) for all listed authors. This work was also supported by the Nomura Research Group and the Mark Foundation for Cancer Research ASPIRE Award for DKN, NF. This work was also supported by grants from the National Institutes of Health (R01CA240981 and R35CA263814 for DKN) and the National Science Foundation Molecular Foundations for Biotechnology Award (2127788). We also thank Drs. Hasan Celik, Alicia Lund, and UC Berkeley’s NMR facility in the College of Chemistry (CoC-NMR) for spectroscopic assistance. Instruments in the College of Chemistry NMR facility are supported in part by NIH S10OD024998.

## Author Contributions

NF, DKN conceived of the project idea, designed experiments, performed experiments, analyzed and interpreted the data, and wrote the paper. NF, DKN, DD performed experiments, analyzed and interpreted data, and provided intellectual contributions. DD, MJH, JAT, JMK, MS provided intellectual contributions to the project and overall design of the project.

## References

1. Burslem, G. M. & Crews, C. M. Proteolysis-Targeting Chimeras as Therapeutics and Tools for Biological Discovery. Cell 181, 102–114 (2020).

2. Schreiber, S. L. The Rise of Molecular Glues. Cell 184, 3–9 (2021).

3. Belcher, B. P., Ward, C. C. & Nomura, D. K. Ligandability of E3 Ligases for Targeted Protein Degradation Applications. Biochemistry (2021) doi:10.1021/acs.biochem.1c00464.

4. Backus, K. M. et al. Proteome-wide covalent ligand discovery in native biological systems. Nature 534, 570–574 (2016).

5. Zhang, X., Crowley, V. M., Wucherpfennig, T. G., Dix, M. M. & Cravatt, B. F. Electrophilic PROTACs that degrade nuclear proteins by engaging DCAF16. Nat. Chem. Biol. 15, 737–746 (2019).

6. Zhang, X. et al. DCAF11 Supports Targeted Protein Degradation by Electrophilic Proteolysis-Targeting Chimeras. J. Am. Chem. Soc. 143, 5141–5149 (2021).

7. Tao, Y. et al. Targeted Protein Degradation by Electrophilic PROTACs that Stereoselectively and Site-Specifically Engage DCAF1. J. Am. Chem. Soc. 144, 18688–18699 (2022).

8. Spradlin, J. N. et al. Harnessing the anti-cancer natural product nimbolide for targeted protein degradation. Nat. Chem. Biol. 15, 747–755 (2019).

9. Luo, M. et al. Chemoproteomics-enabled discovery of covalent RNF114-based degraders that mimic natural product function. Cell Chem. Biol. 28, 559–566.e15 (2021).

10. Ward, C. C. et al. Covalent Ligand Screening Uncovers a RNF4 E3 Ligase Recruiter for Targeted Protein Degradation Applications. ACS Chem. Biol. 14, 2430–2440 (2019).

11. Henning, N. J. et al. Discovery of a Covalent FEM1B Recruiter for Targeted Protein Degradation Applications. J. Am. Chem. Soc. 144, 701–708 (2022).

12. Słabicki, M. et al. The CDK inhibitor CR8 acts as a molecular glue degrader that depletes cyclin K. Nature 585, 293–297 (2020).

13. Lv, L. et al. Discovery of a molecular glue promoting CDK12-DDB1 interaction to trigger cyclin K degradation. eLife 9, e59994 (2020).

14. King, E. A. et al. Chemoproteomics-Enabled Discovery of a Covalent Molecular Glue Degrader Targeting NF-κB. 2022.05.18.492542 Preprint at https://doi.org/10.1101/2022.05.18.492542 (2022).

15. Ottis, P. et al. Cellular Resistance Mechanisms to Targeted Protein Degradation Converge Toward Impairment of the Engaged Ubiquitin Transfer Pathway. ACS Chem. Biol. 14, 2215–2223 (2019).

16. Zhang, L., Riley-Gillis, B., Vijay, P. & Shen, Y. Acquired Resistance to BET-PROTACs(Proteolysis Targeting Chimeras) Caused by Genomic Alterations in Core Components of E3 ligase Complexes. Mol. Cancer Ther. (2019) doi:10.1158/1535-7163.MCT-18-1129.

17. Stewart, M. D., Ritterhoff, T., Klevit, R. E. & Brzovic, P. S. E2 enzymes: more than just middle men. Cell Res. 26, 423–440 (2016).

18. Baek, K. et al. NEDD8 nucleates a multivalent cullin–RING–UBE2D ubiquitin ligation assembly. Nature 578, 461–466 (2020).

19. Lei, L., Bandola-Simon, J. & Roche, P. A. Ubiquitin-conjugating enzyme E2 D1 (Ube2D1) mediates lysine-independent ubiquitination of the E3 ubiquitin ligase March-I. J. Biol. Chem. 293, 3904–3912 (2018).

20. Weerapana, E. et al. Quantitative reactivity profiling predicts functional cysteines in proteomes. Nature 468, 790–795 (2010).

21. Zanon, P. R. A., Lewald, L. & Hacker, S. M. Isotopically Labeled Desthiobiotin Azide (isoDTB) Tags Enable Global Profiling of the Bacterial Cysteinome. Angew. Chem. Int. Ed. 59, 2829–2836 (2020).

22. Henning, N. J. et al. Deubiquitinase-targeting chimeras for targeted protein stabilization. Nat. Chem. Biol. 18, 412–421 (2022).

23. Wu, S.-Y. et al. Opposing Functions of BRD4 Isoforms in Breast Cancer. Mol. Cell 78, 1114–1132.e10 (2020).

