## Supporting Information for "Targeted Protein Degradation through E2 Recruitment"

**Supporting Tables**

**Table S1. Gel-based ABPP screening data.** A library of cysteine-reactive covalent ligands were screened against pure human UBE2D2 C85S by gel-based ABPP. Briefly, the protein was pre-treated with DMSO vehicle or covalent ligand (50 μM) 30 min prior to addition of IA-rhodamine (0.1 μM) for 1 h. Proteins were separated by SDS/PAGE and in-gel fluorescence was assessed and quantified. Noted in the table are percent inhibition of IA-rhodamine labeling compared to the respective controls on each gel.

**Table S2. Cysteine chemoproteomic profiling of EN67 in HEK293T cells.** isoDTB-ABPP cysteine chemoproteomic profiling of EN67 in HEK293T cells. HEK293T cells were treated with DMSO vehicle or EN67 (50 μM) for 4 h. Lysates were labeled with IA-alkyne (200 μM) for 1 h and isotopic desthiobiotin tags were appended by CuAAC and taken through the isoDTB-ABPP procedure. Tab 1 includes the raw chemoproteomic data. Tab 3 lists the quantified probe-modified peptides that were found in all 3 biological replicates that list the probe-modified peptide, uniprot ID, protein, the modified cysteine, gene name, isotopically light (EN67) over heavy (control) fold-changes, inverse fold-changes (control/EN67), p-values, false discovery rate-corrected p-values, log base 2 of control/EN67 fold-change, and the number of runs where the peptide was found and quantified.

**Table S3. TMT-based quantitative proteomic profiling of protein level changes conferred by NF500C treatment in LNCaP cells.** LNCaP cells were treated with DMSO vehicle or NF500C (10 μM) for 24 h. Data are from n=3 biological replicates/group.

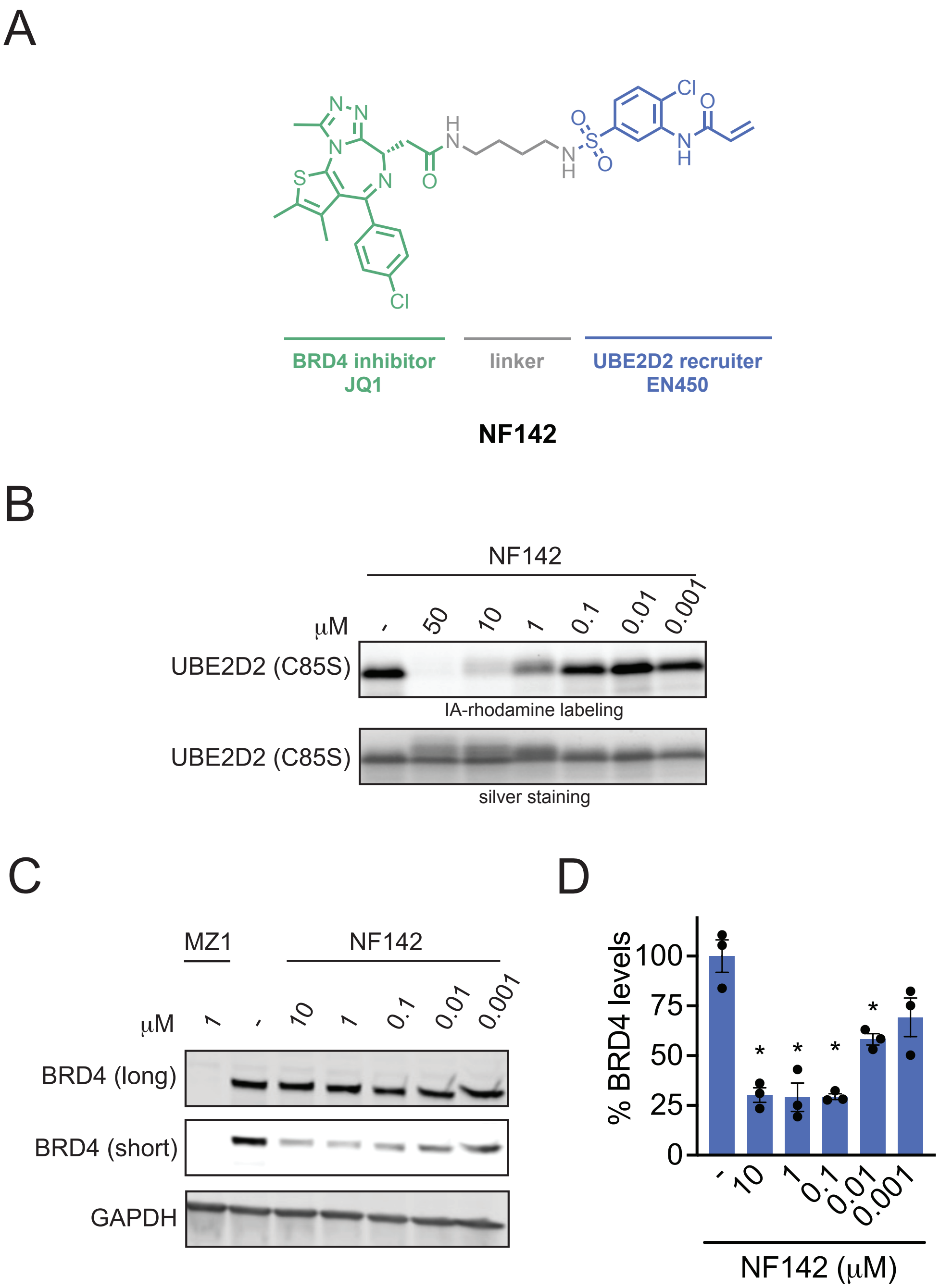


**Figure S1. EN450/UBE2D-Based BRD4 Degrader. (A)** Our previously discovered EN450 UBE2D covalent ligand linked to the BRD4 inhibitor JQ1—NF142. **(B)** gel-based ABPP of NF142 against pure human recombinant UBE2D2 C85S protein. **(C)** BRD4 degradation in HEK293T cells with NF142 treatment. HEK293T cells were treated with DMSO vehicle, MZ1, or NF142 for 24 h and BRD4 long and short isoforms and loading control GAPDH levels were assessed by Western blotting. **(D)** Quantification of BRD4 levels in **(C)**. Gels and blots shown in **(B, C)** are representative from n=3 biological replicates/group. Bar graph shown in **(D)** shows average ± sem with individual replicate values. Statistical significance compared to vehicle-treated control expressed as *$p<0.05$.

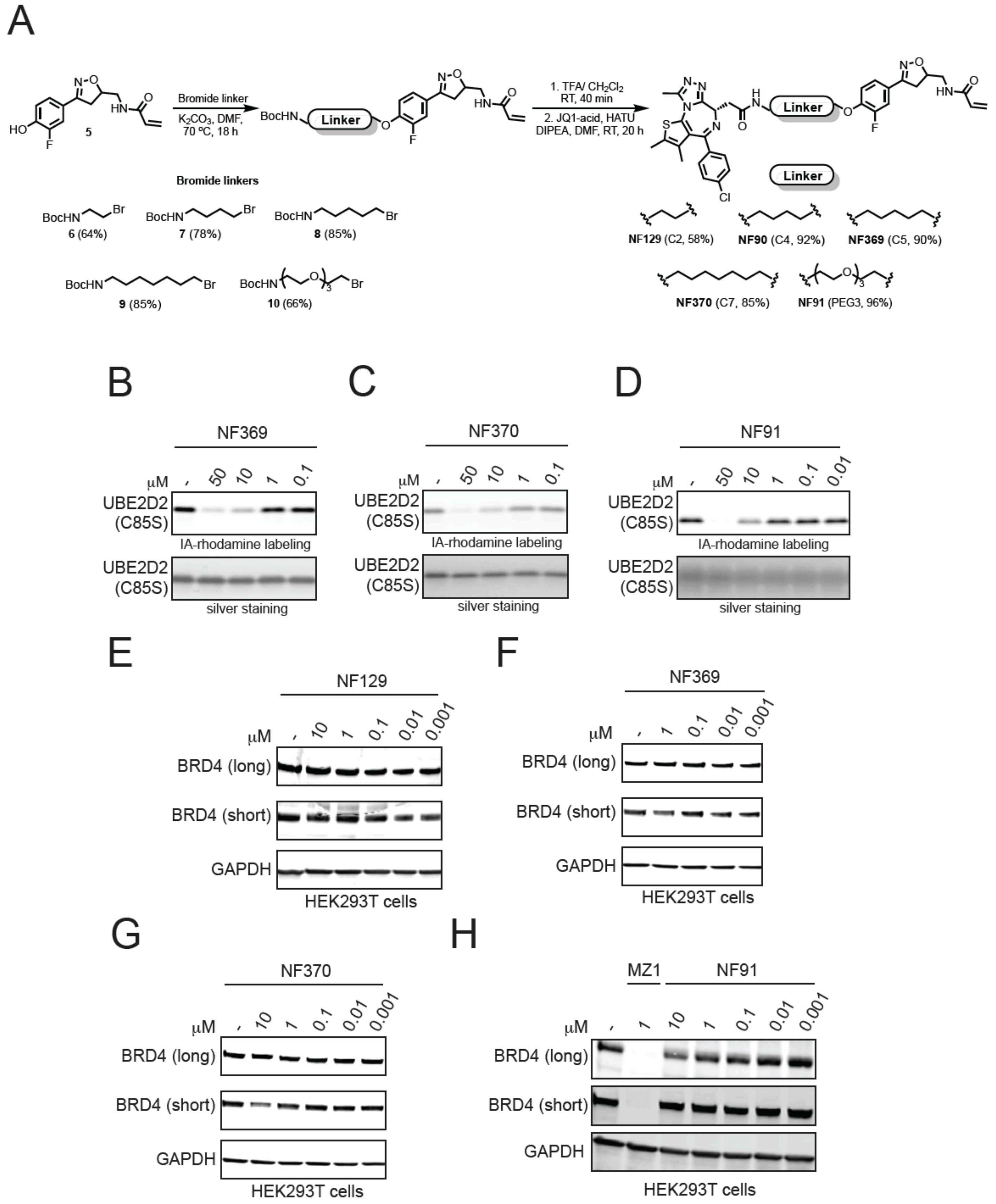
A
Bromide linker
$K_2CO_3$, DMF, 70 ºC, 18 h
1. TFA/ $CH_2Cl_2$ RT, 40 min
2. JQ1-acid, HATU DIPEA, DMF, RT, 20 h
Linker
5
Bromide linkers
6 (64%)
7 (78%)
8 (85%)
9 (85%)
10 (66%)
NF129 (C2, 58%)
NF90 (C4, 92%)
NF369 (C5, 90%)
NF370 (C7, 85%)
NF91 (PEG3, 96%)
B
NF369
μM - 50 10 1 0.1
UBE2D2 (C85S)
IA-rhodamine labeling
silver staining
C
NF370
μM - 50 10 1 0.1
UBE2D2 (C85S)
IA-rhodamine labeling
silver staining
D
NF91
μM - 50 10 1 0.1 0.01
UBE2D2 (C85S)
IA-rhodamine labeling
silver staining
E
NF129
μM - 10 1 0.1 0.01 0.001
BRD4 (long)
BRD4 (short)
GAPDH
HEK293T cells
F
NF369
μM - 1 0.1 0.01 0.001
BRD4 (long)
BRD4 (short)
GAPDH
HEK293T cells
G
NF370
μM - 10 1 0.1 0.01 0.001
BRD4 (long)
BRD4 (short)
GAPDH
HEK293T cells
H
MZ1
NF91
μM - 1 10 1 0.1 0.01 0.001
BRD4 (long)
BRD4 (short)
GAPDH
HEK293T cells

**Figure S2. Characterization of EN67-Based BRD4 degraders. (A)** Sythetic scheme for synthesis of EN67-based BRD4 degraders. **(B-D)** Gel-based ABPP of NF369, NF370, and NF91 against pure human recombinant UBE2D2 C85S protein. **(E-H)** BRD4 levels in cells treated with NF129, NF369, NF370, and NF91 in HEK293T cells. HEK293T cells were treated with DMSO vehicle, a positive control BRD4 PROTAC MZ1, or degrader compounds for 24 h and long and short isoforms of BRD4 and loading control GAPDH levels were assessed by Western blotting. Gels and blots shown in **(B-H)** are representative from n=3 biological replicates/group.

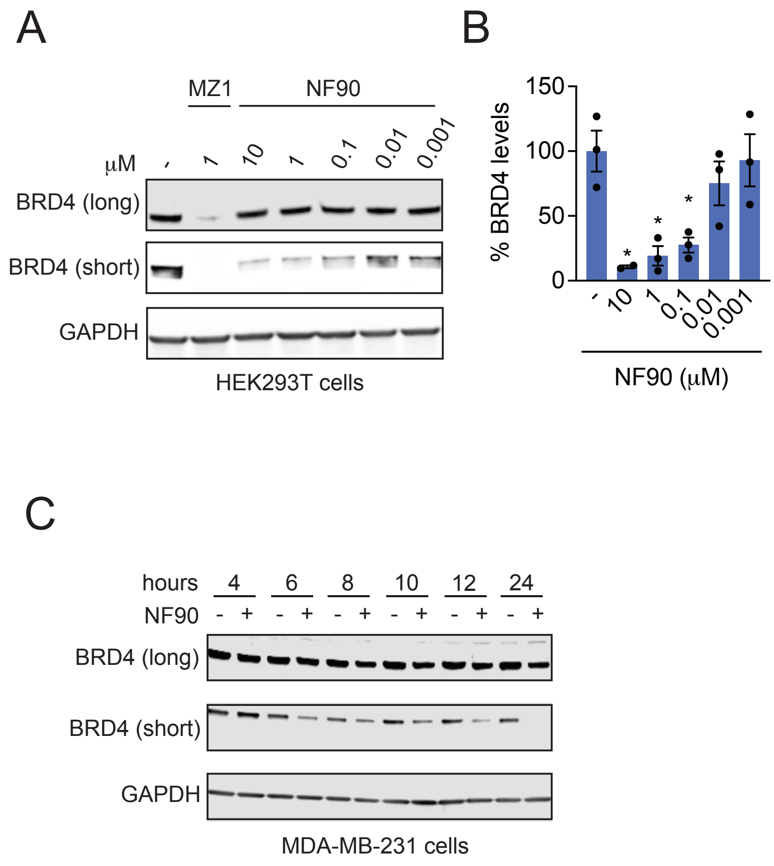


**Figure S3. Characterization of EN67-Based BRD4 degraders. (A)** BRD4 degradation by NF90 in HEK293T cells. HEK293T cells were treated with DMSO vehicle, a positive control BRD4 PROTAC MZ1, or NF90 for 24 h and long and short isoforms of BRD4 and loading control GAPDH levels were assessed by Western blotting. **(B)** Quantification of experiment described in **(A)**. **(C)** BRD4 degradation by NF90 in MDA-MB-231 cells. Cells were treated with DMSO vehicle or NF90 (10 µM) for listed times and BRD4 and loading control GAPDH levels were assessed by Western blotting. Blots shown in **(A, C)** are representative from n=3 biological replicates/group. Bar graph shown in **(B)** shows average ± sem with individual replicate values. Statistical significance compared to vehicle-treated control expressed as *p<0.05.

A

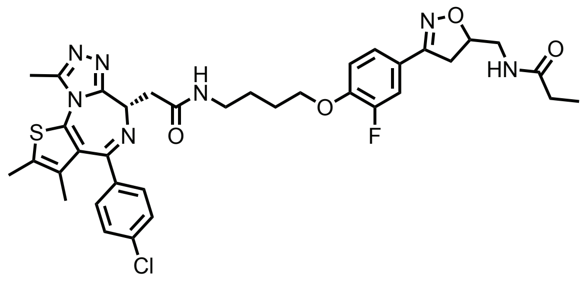


**NF457**

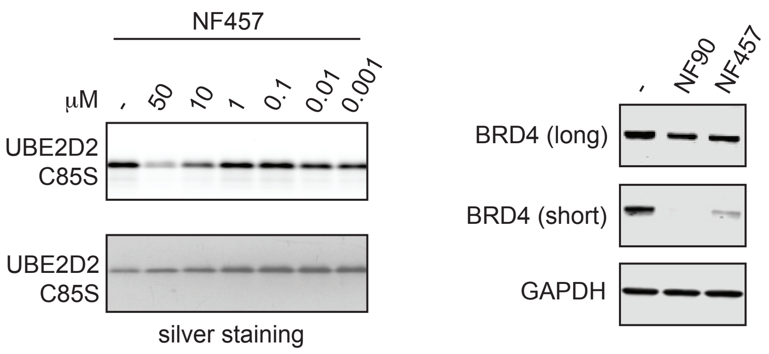


**Figure S4. Non-Reactive NF90 Analog NF457. (A)** Structure of non-reactive analog of NF90—NF457. **(B)** Gel-based ABPP of NF457 against pure human recombinant UBE2D2 C85S protein. **(C)** BRD4 degradation by NF90 in MDA-MB-231 cells. Cells were treated with DMSO vehicle, NF90 (10 μM), or NF457 (10 μM) for 24 h and BRD4 and loading control GAPDH levels were assessed by Western blotting. Gels and blots shown in **(B, C)** are representative from n=3 biological replicates/group.

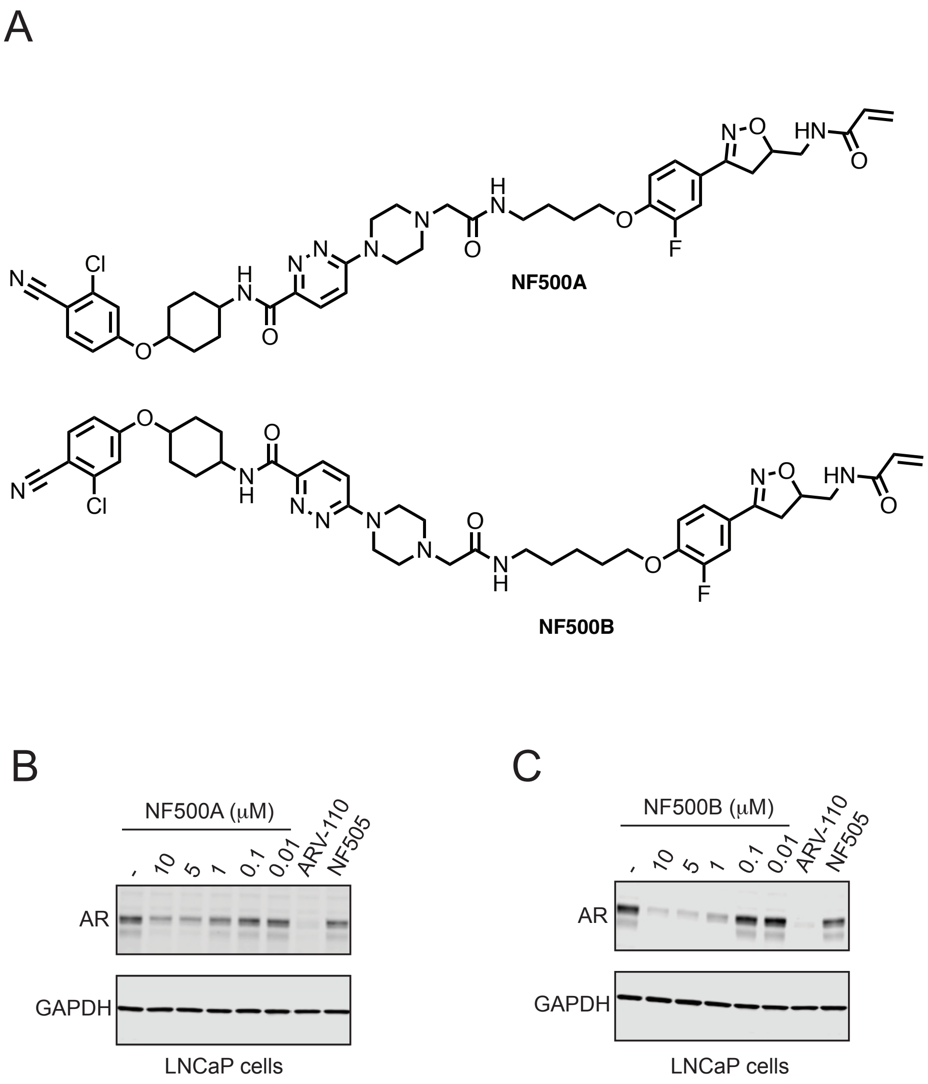


**Figure S5. UBE2D-Based AR Degrader. (A)** Structure of UBE2D-based AR degrader NF500A and NF500B linking the AR-targeting ligand to EN67 via a C4 or C5 alkyl linker, respectively. **(B, C)** AR degradation by NF500A and NF500B. LNCaP prostate cancer cells were treated with DMSO vehicle or NF500A or NF500C for 24 h and AR and loading control GAPDH levels were assessed by Western blotting. Gels and blots shown in **(B, C)** are representative from n=3 biological replicates/group.

A

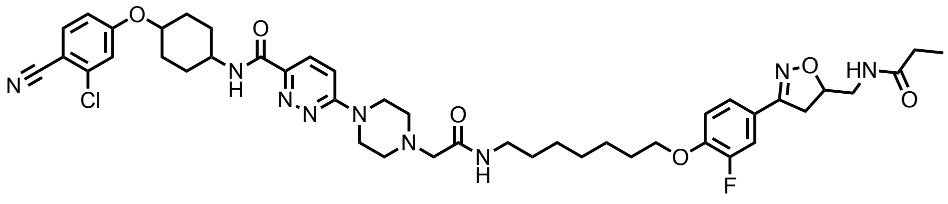

**NF534**

B

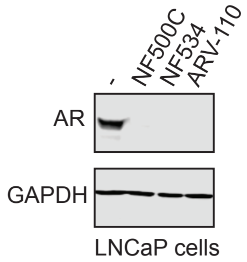


**Figure S6. Non-Reactive NF90 Analog NF500C. (A)** Structure of non-reactive analog of NF500C—NF534. **(B)** BRD4 degradation by NF500C, NF534, and ARV-110 in LNCaP cells. Cells were treated with DMSO vehicle, NF500C (10 μM), NF534 (10 μM), or ARV-110 (1 μM) for 24 h and AR and loading control GAPDH levels were assessed by Western blotting. Blot shown in **(B)** is representative from n=3 biological replicates/group.

### Methods

#### Gel-Based ABPP

UBE2D2 (C85S) (0.1µg/25 µL in PBS) was treated with either DMSO vehicle or covalent ligand at 37 °C for 30 min, and subsequently treated with 0.1 µM IA-Rhodamine (Setareh Biotech) for 1 h at RT in the dark. The reaction was stopped by addition of 4×reducing Laemmli SDS sample loading buffer (Alfa Aesar). After boiling at 95 °C for 5 min, the samples were separated on precast 4−20% Criterion TGX gels (Bio-Rad). Probe-labeled proteins were analyzed by in-gel fluorescence using a ChemiDoc MP (Bio-Rad).

#### Cell Culture

HEK293T cells were obtained from the UC Berkeley Cell Culture Facility and were cultured in Dulbecco's Modified Eagle Medium (DMEM) containing 10% (v/v) fetal bovine serum (FBS) and maintained at 37 °C with 5% $CO_2$. MDA-MB-231 cells were obtained from the UC Berkeley Cell Culture Facility and were cultured in Dulbecco's Modified Eagle Medium (DMEM) containing 10% (v/v) FBS and maintained at 37 °C with 5% $CO_2$. LNCaP cells were obtained from the American Type Culture Collection (ATCC) and were cultured in RPMI-1640 medium containing 10% (v/v) FBS and maintained at 37 °C with 5% $CO_2$. Unless otherwise specified, all cell culture materials were purchased from Gibco. It is not known whether HEK293T cells are from male or female origin.

#### Preparation of Cell Lysates

Cells were washed twice with cold PBS, scraped, and pelleted by centrifugation (700 g, 5 min, 4 °C). Pellets were resuspended in RIPA buffer (supplemented with protease inhibitor cocktail, ThermoFisher, A32963) for western blot analysis. For all other experiments, cells were resuspended in PBS (supplemented with protease inhibitor cocktail ThermoFisher, A32963) and sonicated. Cells were clarified by centrifugation (12,000 x *g*, 10 min, 4 °C), and lysate was transferred to new low-adhesion microcentrifuge tubes. Proteome concentrations were determined using BCA assay and lysate was diluted to appropriate working concentrations.

#### Western Blotting

Proteins were resolved by SDS/PAGE and transferred to nitrocellulose membranes using the Trans-Blot Turbo transfer system (Bio-Rad). Membranes were blocked with 5% BSA in Tris-buffered saline containing Tween 20 (TBST) solution for 1 h at RT and probed with primary antibody diluted in recommended diluent per manufacturer overnight at 4 °C. After 3 washes with TBS-T, the membranes were incubated in the dark with IR680- or IR800-conjugated secondary antibodies at 1:10,000 dilution in 5 % BSA in TBS-T for 1 h at RT. After 3 additional washes with TBST, blots were visualized using an Odyssey Li-Cor fluorescent scanner. The membranes were stripped using ReBlot Plus Strong Antibody Stripping Solution (EMD Millipore) when additional primary antibody incubations were performed. Antibodies used in this study were GAPDH (Cell Signaling Technology, 14C10), BRD4 (Abcam, ab128874), UBE2D1 (ThermoFisher, CF803633), UBE2D2 (Abcam, ab155088), UBE2D3

(Abcam, ab176568), UBE2D4 (ThermoFisher, TA810786), androgen receptor (Cell Signaling Technology, 5153S) and Anti-DDDDK tag (Abcam, ab205606).

**Expression and purification of UBE2D**

Hi-control BL21(DE3) cells were transformed with plasmids expressing either UBE2D1, UBE2D1(C85S), UBE2D1(C85S/C111S), UBE2D2, UBE2D2(C85S), or UBE2D2(C85S/C111S), each containing an N-terminal polyhistidine tag followed by an HRV3C protease cleavage site. Growths were performed with mild shaking at 37 °C using Terrific Broth, with induction of protein expression by treatment with 0.5 mM IPTG once cultures reached an OD600 of 1.5-2.0. Cells were then allowed to grow overnight at 19 °C. Cells were collected by centrifugation and then resuspended in lysis buffer (50 mM Tris pH 8.0, 400 mM NaCl, 1 mM TCEP) prior to lysis by three passes through a cell homogenizer at 18,000 psi. Whole cell lysate was then cleared via centrifugation at 45,000 x *g* for 30 minutes, prior to loading onto 5 mL of Ni-NTA resin pre-equilibrated with lysis buffer. After loading, the resin was washed with 5 CV each of lysis buffer with increasing concentrations of imidazole (10 mM, 20 mM, 40 mM) followed by elution with lysis buffer + 500 mM imidazole. The polyhistidine tag was then cleaved with treatment with HRV3C protease while dialyzing against 3 L of lysis buffer (0 imidazole) overnight at 4 °C. The protein was then run over 5 mL of Ni-NTA resin (pre-equilibrated with lysis buffer) again, this time collecting the flow-through, which was concentrated and then run over a superdex 75 16/60 column for size exclusion chromatography (flow rate = 1 mL/min). Included fractions containing protein were pooled and concentrated to about 12 mg/mL. Total yield from 1 L of bacteria was approximately 70 mg; this was similar between isoforms and variants. At each step, the correct molecular weight of the protein was confirmed by ESI-LC/MS.

**UBE2D2 in vitro ubiquitination assay**

UBE2D2 (5.2 µL, 25 µM, Boston Biochem. Inc., E2-622-100) was diluted in TBS (19.6 µL) and incubated with 0.5 µL of DMSO vehicle or EN67 (10 µM final concentration) for 30 min at 37 °C. Subsequently, UBE1 (1.9 µL, 1 µM, Boston Biochem. Inc., E-305-025) was added followed by MDM2 (1.4 µL, 5 µM, Boston Biochem. Inc., E3-204-050), p53 (17.4 µL, 2 µM, Boston Biochem. Inc., SP454020), FLAG-ubiquitin (1 µL, 10 mg/mL, Boston Biochem. Inc., U12001M), $MgCl_2$ (1 µL, 500 mM), DTT (1 µL, 500 mM) and ATP (1 µL, 100 mM) to achieve a final volume of 50 µL. The mixture was incubated at 37 °C for 4 h with agitation. Then, 20 µL of Laemmli SDS sample loading buffer (Alfa Aesar) was added to quench the reaction and proteins were analyzed by Western Blot. All dilutions were made using 50 mM TBS.

**Mapping of EN67 site of modification on UBE2D2 by LC-MS/MS**

UBE2D2 (100 µg, Boston Biochem. Inc., E2-622-100) was diluted in PBS (1 mL) and pre-incubated with EN67 (10 µM final concentration) for 45 min at 37 °C. The protein was precipitated by addition of 250 µL TCA (100% w/v) and incubation at -80 °C overnight. The sample was then spun at 20,000 g for 10 min at 4 °C. The supernatant was carefully removed, and the sample was washed 3 times with 200 µL ice cold 0.01 M HCl/ 90% acetone solution, with spinning at 20,000 for 5 min at 4 °C between washes. The sample was then resuspended in 30 µL 8M urea in PBS and 30 µL ProteaseMax surfactant (20 µg/mL in 100 mM ammonium bicarbonate, Promega, V2071) with vortexing. Ammonium bicarbonate (40 µL, 100 mM) was then added for a final volume of 100 µL. The sample was reduced with 10 µL TCEP (10 mM final concentration) for 30 min at 60 °C and alkylated with 10 µL iodoacetamide (12.5 mM final concentration) for 30 min at 37 °C. The sample was then diluted with 120 µL PBS before 1.2 µL ProteaseMax surfactant (0.1 mg/mL in 100 mM ammonium bicarbonate, Promega, V2071) and sequencing grade trypsin (10 µL, 0.5 mg/mL in 50 mM ammonium bicarbonate, Promega, V5111) were added for an overnight incubation at 37 °C. The next day, the sample was acidified with formic acid (5% final concentration) and fractionated using high pH reversed-phase peptide fractionation kits (ThermoFisher, 84868) according to manufacturer's protocol.

**Pulldown of UBE2D2 from HEK293T Cells with NF363C probe**

HEK293T cells were treated at 70% confluency with DMSO or NF363C (50 µM) for 24 h. Cells were harvested, lysed via sonication, and the resulting lysate normalized to 5 mg/mL per sample. 500 µL of each lysate sample was incubated for 1 h at RT with 10 µL of 10 mM biotin picolyl azide (in DMSO) (Sigma Aldrich 900912), 10 µL of 50 mM TCEP (in $H_2O$), 30 µL of TBTA ligand (0.9 mg/mL in 1:4 DMSO/tBuOH), and 10 µL of 50 mM $CuSO_4$. Proteins were precipitated, washed 3 × with cold MeOH, resolubilized in 200 µL of 1.2% SDS/PBS (w/v) and heated for 5 min at 90 °C. 10 µL of each sample was removed for Western Blot analysis of input. To the remaining 190 µL was added 1 mL PBS and 50 µL streptavidin agarose beads (ThermoFisher, 20353). Samples were incubated at 4 °C overnight on a rotator. The following day the samples were warmed to RT and washed with 0.2% SDS and further washed 3 × with 500 µL PBS and 3 × with 500 µL $H_2O$ to remove non-probe-labeled proteins. The washed beads were resuspended in 30 µL Laemmli SDS sample loading buffer (Alfa Aesar), heated to 95 °C for 5 min and analyzed by Western Blot.

**IsoDTB-ABPP Cysteine Chemoproteomic Profiling of EN67**

HEK293T cells were treated with either EN67 (50 µM) or DMSO for 4 h before cell collection and lysis. The proteome concentrations were determined using BCA assay and adjusted to 2 mg/mL. For each biological replicate, 2 aliquots of 1 mL of 2 mg/mL were used (i.e. 4 mg per condition). Each aliquot was treated with 20 µL of IA-alkyne (26.6 mg/mL in DMSO, 200 µM final concentration) for 1 h at RT. Two master mixes of the click reagents were prepared in the meanwhile, each containing 1020 µL TBTA (0.9 mg/mL in 4:1 tBuOH/DMSO), 330 µL $CuSO_4$ (12.5 mg/mL in $H_2O$), 330 µL TCEP (14.0 mg/mL in $H_2O$) and 160 µL of either heavy or light isoDTB tags (4 mg in DMSO, Click Chemistry Tools, 1565). The samples were then treated with 120 µL of the heavy (DMSO treated) or light (compound treated) master mix for 1 h at RT. After incubation, one light and one heavy-labeled samples were combined and acetone-precipitated overnight at -20 °C. The samples were then centrifuged at 3,500 rpm for 10 min, acetone was removed and they were resuspended in cold MeOH by sonication. They were centrifuged at 3,500 rpm for 10 min and MeOH was removed (repeated 3× in total). The pellets were dissolved in 300 µL urea (8M in 0.1 M TEAB) by sonication and the urea concentration was then adjusted to 2M by adding 900 µL of TEAB (0.1 M). Two tubes containing solubilized proteins were combined, further diluted with 2400 µL 0.2% NP40 in PBS and bound to high-capacity streptavidin agarose beads (200 µL/sample, ThermoFisher, 20357) for 1 h at RT with mixing. The beads were then centrifuged for 1 min at 1,000 g, the supernatant was removed and the beads were washed 3 times with 0.1% NP40 in PBS, 3 times with PBS and 3 times with $H_2O$. They were then resuspended in 8M urea (600 µL in 0.1 M TEAB) and treated with DTT (30 µL, 31 mg/mL in $H_2O$) for 45 min at 37 °C. They were then reacted with iodoacetamide (30 µL, 74 mg/mL in $H_2O$) for 30 min at RT, followed by DTT (30 µL, 31 mg/mL in $H_2O$) for 30 min at RT. The samples were diluted with 1800 µL TEAB (0.1 M), centrifuged for 1 min at 1,000 g and the supernatant was removed. The beads were resuspended in 400 µL urea (2M in 0.1 M TEAB), trypsin (8 µL, 0.5 mg/mL) was added and incubated for 20 h at 37 °C. The samples were then diluted with 800 µL 0.1% NP40 in PBS and the beads were washed 3 times with 0.1% NP40 in PBS, 3 times with PBS and 3 times with $H_2O$. Peptides were then eluted with 0.1% formic acid in 50% acetonitrile (3 × 400 µL). The samples were then dried by using a vacuum concentrator at 30 °C, resuspended in 300 µL 0.1% TFA in $H_2O$, and fractionated using high pH reversed-phase peptide fractionation kits (ThermoFisher, 84868) according to manufacturer's protocol.

**IsoDTB Mass Spectrometry Analysis**

Mass spectrometry analysis was performed on an Orbitrap Eclipse Tribrid Mass Spectrometer with a High Field Asymmetric Waveform Ion Mobility (FAIMS Pro) Interface (Thermo Scientific) with an UltiMate 3000 Nano Flow Rapid Separation LCnano System (Thermo Scientific). Off-line fractionated samples (5 µl aliquot of 15 µl sample) were injected via an autosampler (Thermo Scientific) onto a 5 µl sample loop which was subsequently eluted onto an Acclaim PepMap 100 C18 HPLC column (75 µm x 50 cm, nanoViper). Peptides were separated at a flow rate of 0.3 µl/min using the following gradient: 2 % buffer B (100 % acetonitrile with 0.1 % formic acid) in buffer A (95:5 water:acetonitrile, 0.1 % formic acid) for 5 min, followed by a gradient from 2 to 40 % buffer B from 5 to 159 min, 40 to 95 % buffer B from 159 to 160 minutes, holding at 95 % B from 160-179 min, 95 % to 2 % buffer B from 179 to 180 min, and then 2 % buffer B from 180 to 200 min. Voltage applied to the nano-LC electrospray ionization source was 2.1 kV. Data was acquired through an MS1 master scan (Orbitrap analysis, resolution 120,000, 400-1800 m/z, RF lens 30 %, heated capillary temperature 250 °C) with dynamic exclusion enabled (repeat count 1, duration 60 s). Data-dependent data acquisition comprised a full MS1 scan followed by sequential MS2 scans based on 2 s cycle times. FAIMS compensation voltages (CV) of -35, -45, and -55 were applied. MS2 analysis consisted of: quadrupole isolation window of 0.7 m/z of precursor ion followed by higher energy collision dissociation (HCD) energy of 38 % with a orbitrap resolution of 50,000.

Data were extracted in the form of MS1 and MS2 files using Raw Converter (Scripps Research Institute) and searched against the Uniprot human database using ProLuCID search methodology in IP2 v.3-v.5 (Integrated Proteomics Applications, Inc.) [1]. Cysteine residues were searched with a static modification for carboxyaminomethylation (+57.02146) and up to two differential modifications for methionine oxidation and either the light or heavy isoDTB tags (+561.33872 or +567.34621, respectively). Peptides were required to be fully tryptic peptides and to contain the TEV modification. ProLUCID data were filtered through DTASelect to achieve a peptide false-positive rate below 5%. Only those probe-modified peptides that were evident across two out of three biological replicates were interpreted for their isotopic light to heavy ratios. Light versus heavy isotopic probe-modified peptide ratios are calculated by taking the mean of the ratios of each replicate paired light versus heavy precursor abundance for all peptide-spectral matches associated with a peptide. The paired abundances were also used to calculate a paired sample *t*-test *P* value in an effort to estimate constancy in paired abundances and significance in change between treatment and control. *P* values were corrected using the Benjamini–Hochberg method.

**Knockdown Studies**

RNA interference was performed using siRNA purchased from Dharmacon. HEK293T cells were seeded at 250,000 cells per 6 cm plate and allowed to adhere overnight. Cells were transfected with either 33 nM of non-targeting (ON-TARGETplus Non-targeting Control Pool, Dharmacon, D-001810-10-20) or 16.5 nM of anti-UBE2D1 (Dharmacon, L-009387-00-0005), anti-UBE2D2 (Dharmacon, L-010383-00-0005), anti-UBE2D3 (Dharmacon, L-008478-00-0005) and anti-UBE2D4 siRNA (Dharmacon, L-009435-00-0005) using 5 µL of Lipofectamine 2000 (ThermoFisher, 11668027). Transfection reagent was added to Opti-MEM (ThermoFisher, 31985070) media and allowed to incubate for 5 min at RT. Meanwhile siRNA was added to an equal amount of Opti-MEM. Solutions of transfection reagent and siRNA in Opti-MEM were then combined and allowed to incubate for 30 min at RT. These combined solutions were diluted with complete DMEM to provide 2 mL per well, and the media exchanged. Cells were incubated with transfection reagents for 48 h, at which point the media was replaced with media containing DMSO or NF90 (10 µM) and incubated for another 24 h. Cells were then harvested, and protein abundance was analyzed by Western blotting.

**TMT-based quantitative proteomic profiling**

LNCaP cells were treated at 70% confluency with either DMSO vehicle or NF500C (10 µM) for 24 h and lysate was prepared as described above. Briefly, 25-100 µg protein from each sample was reduced, alkylated and tryptically digested overnight. Individual samples were then labeled with isobaric tags using commercially available TMTsixplex (ThermoFisher, 90061) kits, in accordance with the manufacturer's protocols. Tagged samples (20 µg per sample) were combined, dried with SpeedVac, resuspended with 300 µL 0.1% TFA in H2O, and fractionated using high pH reversed-phase peptide fractionation kits (ThermoFisher, 84868) according to manufacturer's protocol. Fractions were dried with SpeedVac, resuspended with 50 µL 0.1% FA in $H_2O$, and analyzed by LC-MS/MS as described below.

Mass spectrometry analysis of resulting TMT peptides were performed as described above. Trypsin cleavage specificity (cleavage at K, R except if followed by P) allowed for up to 2 missed cleavages. Carbamidomethylation of cysteine was set as a fixed modification, methionine oxidation, and TMT-modification of *N*-termini and lysine residues were set as variable modifications. Reporter ion ratio calculations were performed using summed abundances with most confident centroid selected from 20 ppm window. Only peptide-to-spectrum matches that are unique assignments to a given identified protein within the total dataset are considered for protein quantitation. High confidence protein identifications were reported with a <1% false discovery rate (FDR) cut-off. Differential abundance significance was estimated using ANOVA with Benjamini-Hochberg correction to determine p-values.

### Synthetic Methods and Characterization

All chemical reactions were carried out under a nitrogen atmosphere with dry solvents under anhydrous conditions, unless otherwise noted. Reagents were purchased at the highest commercial quality and used without further purification, unless otherwise stated. Room temperature is defined as between 21-25 °C. Reactions were stirred magnetically and monitored by thin layer chromatography (TLC) using TLC plates pre-coated with silica gel 60 F254 on aluminium (Merck KGaA). Detection was by UV (254 nm and 365 nm) or chemical stain ($KMnO_4$, ninhydrin, iodine). Solvents were removed *in vacuo* using a Buchi R-300 Rotavapor (equipped with an I-300 Pro Interface, B-300 Base Heating Bath, Welch 2037B-01 DryFast pump, and VWR AD15R-40-V11B Circulating Bath). Solvents for silica gel chromatography were used as supplied by Sigma-Aldrich. Automated flash chromatography was performed on a Biotage Isolera instrument, equipped with a UV detector. Chromatograms were recorded at 254 and 280 nm. High-resolution mass spectra (HRMS) were obtained at the Catalysis Center at the College of Chemistry, University of California, Berkeley. $^{1}H$ and $^{13}C$ Nuclear Magnetic Resonance (NMR) spectra were recorded on BRUKER AV spectrometer operating at 600 MHz for $^{1}H$ and at 150 MHz for $^{13}C$ NMR. Measurements were carried out at ambient temperature. Chemical shifts (δ) are reported in ppm with the residual solvent signal as internal standard (chloroform at 7.26 and 77.2 ppm for $^{1}H$ NMR and $^{13}C$ NMR, respectively, methanol at 3.31 and 49.0, respectively and DMSO at 2.50 and 39.5, respectively). The multiplicity of each signal is indicated as s = singlet, d = doublet, t = triplet, q = quartet, quin = quintet, m = multiplet (i.e. complex peak obtained due to overlap), app = apparent or a combination of these. Coupling constants (J) are reported in Hertz (Hz). $^{13}C$ NMR spectra were recorded with broadband $^{1}H$ decoupling.

#### ***tert*-butyl (4-((4-chloro-3-nitrophenyl)sulfonamido)butyl)carbamate (1)**

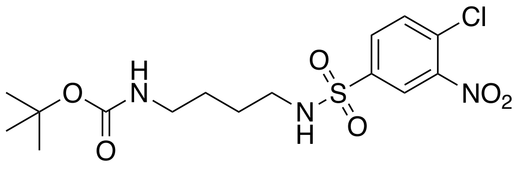


To 4-chloro-3-nitrobenzenesulfonyl chloride (700 mg, 2.73 mmol) in $CH_2Cl_2$ (20 mL) was added *N*-Boc-1,4-butanediamine (475 µL, 2.48 mmol), followed by $NEt_3$ (416 µL, 2.98 mmol). The resultant mixture was stirred at RT for 22 h. The solvent was then removed *in vacuo* and purification by column chromatography (gradient elution from $CH_2Cl_2$ to 10% MeOH in $CH_2Cl_2$) afforded the target product as a yellow crystalline solid (860 mg, 2.11 mmol, 77%).

$^1$H NMR ($CD_3OD$, 600 MHz) $\delta_H$ 8.37 (d, 1H, $J$ = 2.1 Hz), 8.06 (dd, 1H, $J$ = 8.4, 2.1 Hz), 7.89 (d, 1H, $J$ = 8.5 Hz), 6.50 (br s, 1H), 3.02 (q, 2H, $J$ = 6.2 Hz), 2.95 (t, 2H, $J$ = 6.9 Hz), 1.53-1.47 (m, 4H), 1.43 (s, 9H); $^{13}$C NMR $CD_3OD$, 150 MHz) $\delta_C$ 157.2, 148.0, 141.3, 132.7, 130.9, 130.0, 123.8, 78.6, 42.4, 39.5, 27.4, 26.7, 26.6; HRMS (ES+) calcd for $C_{15}H_{22}O_6N_3ClNaS$ $[M+Na]^+$ 430.0810, found 430.0812.

***tert*-butyl (4-((3-acrylamido-4-chlorophenyl)sulfonamido)butyl)carbamate (2)**

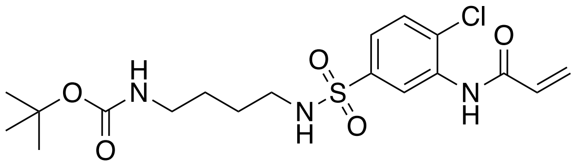


To reagent **1** (237 mg, 0.588 mmol) in a mixture of EtOH (8 mL)/ $H_2O$ (2 mL) was added $NH_4Cl$ (201 mg, 3.76 mmol) followed by iron powder (105 mg, 1.88 mmol). The resultant mixture was stirred at 80 °C for 18 h. It was then filtered through a pad of celite and concentrated *in vacuo*. It was dissolved in EtOAc (50 mL), washed with $H_2O$ (3 × 30 mL) and dried ($MgSO_4$). The crude product was re-dissolved in $CH_2Cl_2$ (5 mL) and it was added in a stirring mixture of acrylic acid (56.0 µL, 0.816 mmol), T3P (445 µL, 0.748 mmol) and $NEt_3$ (347 µL, 2.49 mmol) in $CH_2Cl_2$ (5 mL). The resultant mixture was stirred at 30 °C for 18 h. It was then washed with 1M citric acid (30 mL) and extracted in $CH_2Cl_2$ (3 × 50 mL). The combined organic extracts were washed with brine (30 mL), dried ($MgSO_4$) and concentrated. Purification by column chromatography (gradient elution from 30% diethyl ether in $CH_2Cl_2$ to diethyl ether) afforded the target product as a clear oil (65.0 mg, 0.151 mmol, 26%).

$^1$H NMR ($CD_3OD$, 600 MHz) $\delta_H$ 8.47 (d, 1H, $J$ = 2.1 Hz), 7.68-7.64 (m, 2H), 6.62 (dd, 1H, $J$ = 17.0, 10.3 Hz), 6.46 (dd, 1H, $J$ = 17.0, 1.6 Hz), 5.88 (dd, 1H, $J$ = 10.3, 1.6 Hz), 3.01 (q, 2H, $J$ = 6.3 Hz), 2.94 (t, 2H, $J$ = 6.5 Hz), 1.50-1.47 (m, 4H), 1.43 (s, 9H); $^{13}$C NMR ($CD_3OD$, 150 MHz) $\delta_C$ 165.1, 157.2, 157.1, 139.9, 135.2, 130.2, 130.1, 128.0, 124.0, 123.4, 78.5, 42.5, 39.4, 27.4, 26.6, 26.5; HRMS (ES+) calcd for $C_{18}H_{26}O_5N_3ClNaS$ $[M+Na]^+$ 454.1174, found 454.1178.

**(*S*)-*N*-(2-chloro-5-(*N*-(4-(2-(4-(4-chlorophenyl)-2,3,9-trimethyl-6*H*-thieno[3,2-*f*][1,2,4] triazolo[4,3-*a*][1,4] diazepin-6-yl)acetamido)butyl)sulfamoyl)phenyl)acrylamide (NF142)**

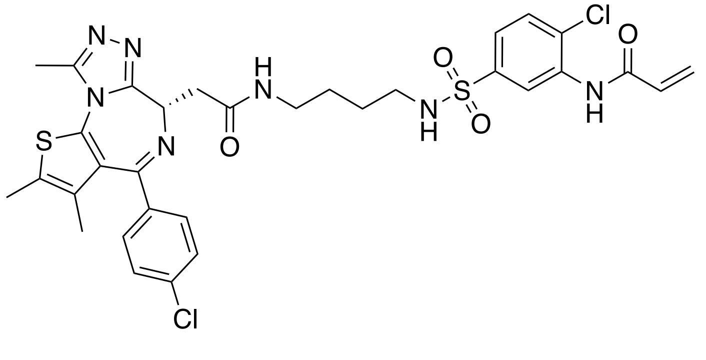

To reagent **2** (100 mg, 0.232 mmol) in $CH_2Cl_2$ (4 mL) was added TFA (2 mL). The resultant mixture was stirred at RT for 1 h. It was then concentrated *in vacuo*, dissolved in DMF (4 mL) and added to a pre-stirred mixture of JQ1-acid (60.0 mg, 0.150 mmol), HATU (64.0 mg, 0.168 mmol) and DIPEA (59.0 µL, 0.339 mmol) in DMF (4 mL). The resultant mixture was stirred at RT for 20 h. It was then concentrated *in vacuo* and purified by column chromatography (gradient elution $CH_2Cl_2$ to 10% MeOH in $CH_2Cl_2$) to afford the target product as a white solid (95.0 mg, 0.133 mmol, 89%).

$^1$H NMR (DMSO-d6 with a drop of $CDCl_3$, 600 MHz) $\delta_H$ 9.95 (s, 1H), 8.18-8.14 (m, 1H), 7.75-7.73 (m, 1H), 7.50-7.48 (m, 2H), 7.42-7.40 (m, 2H), 7.38 (d, 1H, *J* = 8.3 Hz), 7.22 (d, 1H, *J* = 2.2 Hz), 6.90 (dd, 1H, *J* = 8.3, 2.2 Hz), 6.67 (dd, 1H, *J* = 17.0, 10.2 Hz), 6.33 (dd, 1H, *J* = 17.0, 1.8 Hz), 5.85-5.83 (m, 1H), 4.51-4.48 (m, 1H), 3.24 (ddd, 1H, *J* = 15.0, 8.3, 1.9 Hz), 3.20-3.15 (m, 1H), 3.12-3.07 (m, 1H), 3.06-3.01 (m, 1H), 2.80-2.74 (m, 1H), 2.60 (s, 3H), 2.41 (s, 3H), 1.62 (s, 3H); $^{13}$C NMR (DMSO-d6 with a drop of $CDCl_3$, 150 MHz) $\delta_C$ 169.9, 164.2, 163.5, 140.3, 137.2, 135.8, 135.7, 132.7, 131.5, 131.2, 131.0, 130.6, 130.3, 130.2, 130.1, 129.0, 124.3, 123.8, 120.8, 114.5, 113.3, 54.3, 42.8, 38.5, 38.1, 31.2, 26.9, 14.5, 13.2, 11.8; HRMS (ES+) calcd for $C_{32}H_{34}O_4N_7Cl_2S_2$ $[M+H]^+$ 714.1485, found 714.1474.

**(*E*)-3-fluoro-4-hydroxybenzaldehyde oxime (3)**

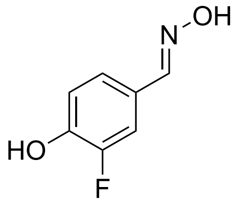


To a mixture of 3-fluoro-4-hydroxybenzaldehyde (500 mg, 3.57 mmol) and hydroxylamine hydrochloride (298 mg, 4.28 mmol) in MeOH (10 mL) and $H_2O$ (10 mL) was slowly added $K_2CO_3$ (740 mg, 5.36 mmol) over 5 min. The resultant mixture was stirred at RT for 18 h. MeOH was then removed *in vacuo*, the aqueous layer was acidified by the addition of 1M aq. HCl, extracted in EtOAc (3 × 50 mL) and dried ($MgSO_4$) to afford the target product as a pale yellow solid (552 mg, 3.56 mmol, quant.).

$^1$H NMR (DMSO-d6, 600 MHz) $\delta_H$ 11.02 (s, 1H), 10.18 (s, 1H), 8.02 (s, 1H), 7.35 (dd, 1H, *J* = 12.3, 2.0 Hz), 7.22 (dd, 1H, *J* = 8.0, 1.8 Hz), 6.96 (t, 1H, *J* = 8.8 Hz); $^{13}$C NMR (DMSO-d6, 150 MHz) $\delta_C$ 151.5 (d, *J* = 241.1 Hz), 147.6 (d, *J* = 2.8 Hz), 146.5 (d, *J* = 12.2 Hz), 125.3 (d, *J* = 6.5 Hz), 123.8 (d, *J* = 3.1 Hz), 118.3 (d, *J* = 3.2 Hz), 114.0 (d, *J* = 19.3 Hz); HRMS (ES-) calcd for $C_7H_5O_2NF$ $[M-H]^+$ 154.0310, found 154.0309.

**3-(3-fluoro-4-hydroxyphenyl)-4,5-dihydroisoxazole-5-carbonitrile (4)**

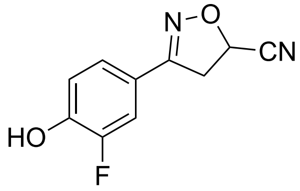

To a stirring solution of **3** (1.40 g, 9.03 mmol) in DMF (20 mL) was added *N*-chlorosuccinimide (1.45 g, 10.8 mmol) in DMF (10 mL) over 15 min. The resultant mixture was stirred at RT for 7 h, after which time acrylonitrile (710 µL, 10.8 mmol) and $NEt_3$ (1.89 mL, 13.5 mmol) were added. After 48 h at RT, the precipitate was filtered off. The filtrate was concentrated *in vacuo* and purified by column chromatography (gradient elution from 50% diethyl ether/ hexanes to 100% diethyl ether) to afford the target product as an off white solid (1.56 g, 7.57 mmol, 84%).

$^1$H NMR ($CD_3OD$, 600 MHz) $\delta_H$ 7.44 (dd, 1H, *J* = 11.9, 2.1 Hz), 7.32 (ddd, 1H, *J* = 8.5, 2.1, 1.0 Hz), 6.97 (t, 1H, *J* = 8.6 Hz), 5.56 (dd, 1H, *J* = 10.8, 5.7 Hz), 3.82-3.72 (m, 2H); $^{13}$C NMR ($CD_3OD$, 150 MHz) $\delta_C$ 156.3 (d, *J* = 2.7 Hz), 151.4 (d, *J* = 242.1 Hz), 147.8 (d, *J* = 13.2 Hz), 123.9 (d, *J* = 3.2 Hz), 119.4 (d, *J* = 6.9 Hz), 117.7 (d, *J* = 3.2 Hz), 117.6, 114.3 (d, *J* = 20.5 Hz), 66.8, 40.6; HRMS (ES-) calcd for $C_{10}H_6FN_2O_2$ $[M-H]^+$ 205.0419, found 205.0418.

***N*-((3-(3-fluoro-4-hydroxyphenyl)-4,5-dihydroisoxazol-5-yl)methyl)acrylamide (5)**

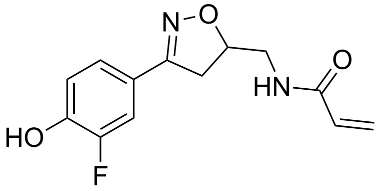

Reagent **4** (636 mg, 3.09 mmol) was dissolved in MeOH (20 mL) and degassed for 20 min. Aq. sat. HCl (0.85 mL) was then added, followed by palladium on carbon (424 mg, 10 wt.%). The resultant mixture was stirred under a $H_2$ atmosphere for 20 h, after which time it was filtered through a pad of celite and concentrated *in vacuo*. It was then dissolved in DMF (30 mL) and $NEt_3$ (1.7 mL, 12.5 mmol) was added, followed by acryloyl chloride (676 µL, 3.98 mmol). The resultant mixture was stirred at RT for 18 h. The precipitate was filtered off and the filtrate was concentrated *in vacuo* before it was redissolved in a mixture of MeOH (10 mL) and 1M aq. NaOH (10 mL). After 1 h at RT, the mixture was acidified with 1M aq. HCl and extracted into EtOAc (3 × 50 mL). The combined organic extracts were washed with brine (50 mL) and dried ($MgSO_4$). Purification by column chromatography (gradient elution from $CH_2Cl_2$ to 8% MeOH in $CH_2Cl_2$) afforded the target product as an off white foam (404 mg, 1.53 mmol, 44%).

$^1$H NMR ($CD_3OD$, 600 MHz) $\delta_H$ 7.41 (dd, 1H, *J* = 12.0, 2.1 Hz), 7.29 (ddd, 1H, 8.4, 2.1, 1.0 Hz), 6.96 (t, 1H, *J* = 8.6 Hz), 6.31-6.22 (m, 2H), 5.68 (dd, 1H, *J* = 9.7, 2.2 Hz), 4.84-4.82 (overlapped m, 1H), 3.52 (d, 2H, *J* = 5.8 Hz), 3.45 (dd, 1H, *J* = 17.0, 10.5 Hz), 3.16 (dd, 1H, *J* = 17.0, 7.1 Hz); $^{13}$C NMR ($CD_3OD$, 150 MHz) $\delta_C$ 167.3, 156.3 (d, *J* = 2.7 Hz), 151.3 (d, *J* = 241.5 Hz), 147.1 (d, *J* = 12.8 Hz), 130.4, 125.8, 123.4 (d, *J* = 3.0 Hz), 121.0

(d, $J$ = 6.6 Hz), 117.5 (d, $J$ = 3.2 Hz), 113.8 (d, $J$ = 20.4 Hz), 79.4, 42.2, 37.5; HRMS (ES+) calcd for $C_{13}H_{13}O_3N_2FNa$ $[M+Na]^+$ 287.0802, found 287.0805.

**_N_-((3-(3-fluoro-4-(prop-2-yn-1-yloxy)phenyl)-4,5-dihydroisoxazol-5-yl)methyl)acrylamide (NF363A)**

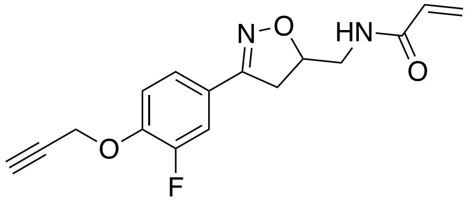

To phenol **5** (30.0 mg, 0.114 mmol) was added propargyl bromide (9.50 µL, 0.125 mmol), followed by $K_2CO_3$ (19 mg, 0.137 mmol). The resultant mixture was stirred at 70 °C for 18 h. It was then concentrated *in vacuo* and purified by column chromatography (gradient elution from $CH_2Cl_2$ to 10% MeOH in $CH_2Cl_2$) to afford the target compound as a white oil (28.0 mg, 0.0927 mmol, 81%).

$^1$H NMR ($CD_3OD$, 600 MHz) $\delta_H$ 7.49 (d, 1H, $J$ = 12.1 Hz), 7.40 (d, 1H, J = 8.5 Hz), 7.25 (t, 1H, J = 8.5 Hz), 6.30-6.21 (m, 2H), 5.68 (dd, 1H, $J$ = 9.5, 2.4 Hz), 4.91-4.86 (m, 3H), 3.53-3.52 (m, 2H), 3.47 (dd, 1H, $J$ = 17.0, 10.6 Hz), 3.17 (dd, 1H, $J$ = 17.0, 7.2 Hz), 3.04 (t, 1H, $J$ = 2.4 Hz); $^{13}$C NMR ($CD_3OD$, 150 MHz) $\delta_C$ 167.2, 156.0 (d, $J$ = 2.4 Hz), 152.4 (d, $J$ = 246.2 Hz), 147.1 (d, $J$ = 6.6 Hz), 130.4, 125.8, 123.3 (d, $J$ = 6.6 Hz), 123.0 (d, $J$ = 3.4 Hz), 115.3, 113.8 (d, $J$ = 20.4 Hz), 79.7, 77.5, 76.4, 56.4, 42.2, 37.3; HRMS (ES+) calcd for $C_{16}H_{16}O_3N_2F$ $[M+H]^+$ 303.1139, found 303.1138.

**_N_-((3-(4-(but-3-yn-1-yloxy)-3-fluorophenyl)-4,5-dihydroisoxazol-5-yl)methyl)acrylamide (NF363B)**

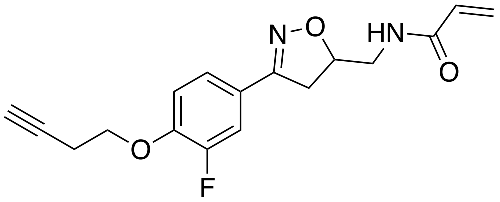

To phenol **5** (40.0 mg, 0.151 mmol) was added 4-bromo-1-butyne (15.0 µL, 0.160 mmol), followed by $K_2CO_3$ (25.0 mg, 0.181 mmol). The resultant mixture was stirred at 70 °C for 18 h. It was then concentrated *in vacuo* and purified by column chromatography (gradient elution from $CH_2Cl_2$ to 10% MeOH in $CH_2Cl_2$) to afford the target compound as a yellow foam (19.0 mg, 0.0601 mmol, 40%).

$^1$H NMR (DMSO-d6, 600 MHz) $\delta_H$ 8.39 (t, 1H, $J$ = 6.0 Hz), 7.50 (dd, 1H, $J$ = 12.2, 2.1 Hz), 7.40 (ddd, 1H, $J$ = 8.5, 2.2, 1.0 Hz), 7.27 (t, 1H, $J$ = 8.7 Hz), 6.27 (dd, 1H, $J$ = 17.1, 10.2 Hz), 6.10 (dd, 1H, $J$ = 17.1, 2.1 Hz), 5.60 (dd, 1H, $J$ = 10.2, 2.1 Hz), 4.80-4.75 (m, 1H), 4.20 (t, 2H, $J$ = 6.5 Hz), 3.45 (dd, 1H, $J$ = 17.1, 10.6 Hz), 3.36 (t, 2H, $J$ = 5.8 Hz), 3.12 (dd, 1H, $J$ = 17.1, 7.2 Hz), 2.91 (t, 1H, $J$ = 2.7 Hz), 2.69 (td, 2H, $J$ = 6.5, 2.7 Hz); $^{13}$C NMR (DMSO-d6, 150 MHz) $\delta_C$ 165.5, 156.1, 151.8 (d, $J$ = 244.8 Hz), 147.9 (d, $J$ = 10.5 Hz), 131.9, 126.0, 124.1 (d, $J$ = 3.3

Hz), 123.1 (d, *J* = 6.8 Hz), 115.5, 114.3 (d, *J* = 19.4 Hz), 81.5, 79.9, 73.1, 67.4, 42.4, 37.9, 19.3; HRMS (ES+) calcd for $C_{17}H_{18}O_3N_2F$ $[M+H]^+$ 317.1296, found 317.1298.

***N*-((3-(3-fluoro-4-(pent-4-yn-1-yloxy)phenyl)-4,5-dihydroisoxazol-5-yl)methyl)acrylamide (NF363C)**

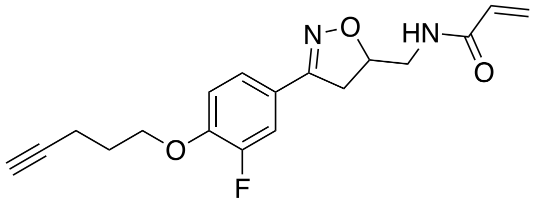

To phenol **5** (40.0 mg, 0.151 mmol) was added 5-Iodo-1-pentyne (19.0 µL, 0.167 mmol), followed by $K_2CO_3$ (25.0 mg, 0.181 mmol). The resultant mixture was stirred at 70 °C for 18 h. It was then concentrated *in vacuo* and purified by column chromatography (gradient elution from $CH_2Cl_2$ to 10% MeOH in $CH_2Cl_2$) to afford the target compound as an off white solid (36.0 mg, 0.109 mmol, 72%).

$^1$H NMR ($CD_3OD$, 600 MHz) 7.48 (dd, 1H, *J* = 12.2, 2.1 Hz) 7.39 (br d, 1H, *J* = 8.5 Hz), 7.16 (t, 1H, *J* = 8.5 Hz), 6.30-6.22 (m, 2H), 5.68 (dd, 1H, *J* = 9.5, 2.5 Hz), 4.90-4.86 (m, 1H), 4.21 (t, 2H, *J* = 6.1 Hz), 3.53-3.51 (m, 2H), 3.47 (dd, 1H, 17.0, 10.6. Hz), 3.18 (dd, 1H, *J* = 17.0, 7.1 Hz), 2.42 (td, 2H, *J* = 7.0, 2.7 Hz), 2.28 (t, 1H, *J* = 2.7 Hz), 2.02 (quin, 2H, *J* = 6.6 Hz); $^{13}$C NMR ($CD_3OD$, 150 MHz) 167.2, 156.1, 152.3 (d, *J* = 245.9 Hz), 148.7 (d, *J* = 10.9 Hz), 130.3, 125.7, 123.3 (d, *J* = 3.3 Hz), 122.4 (d, *J* = 6.7 Hz), 114.3, 113.6 (d, *J* = 20.4 Hz), 82.4, 79.6, 68.8, 67.3, 42.2, 37.3, 27.9, 14.2; HRMS (ES+) $C_{18}H_{19}O_3N_2FNa$ $[M+Na]^+$ calcd 353.1272, found 353.1272.

***N*-((3-(3-fluoro-4-(hex-5-yn-1-yloxy)phenyl)-4,5-dihydroisoxazol-5-yl)methyl)acrylamide (NF363D)**

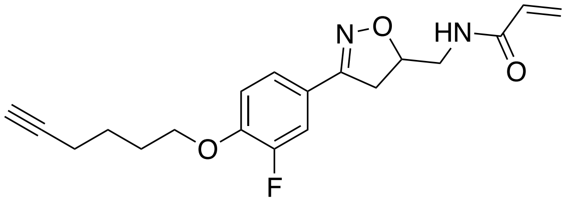

To phenol **5** (40.0 mg, 0.151 mmol) was added 6-Iodo-1-hexyne (22.0 µL, 0.167 mmol), followed by $K_2CO_3$ (25.0 mg, 0.181 mmol). The resultant mixture was stirred at 70 °C for 18 h. It was then concentrated *in vacuo* and purified by column chromatography (gradient elution from $CH_2Cl_2$ to 10% MeOH in $CH_2Cl_2$) to afford the target compound as a white solid (29.0 mg, 0.0843 mmol, 56%).

$^1$H NMR (DMSO-d6, 600 MHz) $\delta_H$ 8.38 (t, 1H, *J* = 6.0 Hz), 7.49 (dd, 1H, *J* = 12.3, 2.1 Hz), 7.40 (br d, 1H, J =8.5 Hz), 7.24 (t, 1H, *J* = 8.7 Hz), 6.27 (dd, 1H, *J* = 17.1, 10.2 Hz), 6.10 (dd, 1H, *J* = 17.1, 2.1 Hz), 5.60 (dd, 1H, *J* = 10.2, 2.1 Hz), 4.80-4.75 (m, 1H), 4.12 (t, 2H, *J* = 6.4 Hz), 3.45 (dd, 1H, *J* = 17.1, 10.6 Hz), 3.36 (t, 2H, *J* = 5.8 Hz), 3.11 (dd, 1H, *J* = 17.1, 7.1 Hz), 2.78 (t, 1H, *J* = 2.6 Hz), 2.25 (td, 2H, *J* = 7.1, 2.7 Hz), 1.84 (quin, 2H, *J* = 6.4 Hz), 1.61 (quin, 2H, *J* = 7.2 Hz); $^{13}$C NMR (DMSO-d6, 150 MHz) $\delta_C$ 165.5, 156.1 (d, *J* = 2.4 Hz), 151.9 (d, *J* = 244.3 Hz), 148.4 (d, *J* = 10.6 Hz), 131.9, 126.0, 124.1 (d, *J* = 3.2 Hz), 122.7 (d, *J* = 7.1 Hz), 115.3, 114.2 (d, *J*

= 19.5 Hz), 84.7, 79.9, 71.9, 68.7, 42.4, 37.9, 28.1, 25.0, 17.9; HRMS (ES+) calcd for $C_{19}H_{21}O_3N_2FNa$ [M+Na]$^+$ 367.1428, found 367.1430.

**tert-butyl (2-(4-(5-(acrylamidomethyl)-4,5-dihydroisoxazol-3-yl)-2-fluorophenoxy)ethyl) carbamate (6)**

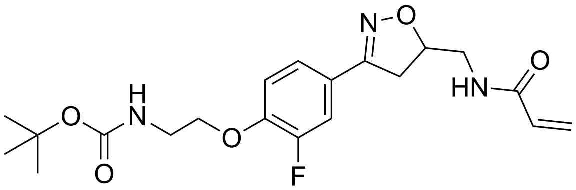

To phenol **5** (60.0 mg, 0.227 mmol) in DMF (5 mL) was added 2-(Boc-amino)ethyl bromide (56.0 mg, 0.250 mmol), followed by $K_2CO_3$ (38.0 mg, 0.275 mmol). The resultant mixture was stirred at 70 °C for 18 h. After this time, the solvent was removed *in vacuo* and purification by column chromatography (gradient elution from $CH_2Cl_2$ to 8% MeOH in $CH_2Cl_2$) afforded the target product as a clear oil (59.0 mg, 0.145 mmol, 64%).

$^1$H NMR ($CD_3OD$, 600 MHz) $\delta_H$ 7.47 (dd, 1H, *J* = 12.1, 2.1 Hz), 7.38 (br d, 1H, *J* = 8.5 Hz), 7.16 (t, 1H, *J* = 8.6 Hz), 6.30-6.21 (m, 2H), 5.68 (dd, 1H, *J* = 17.0, 7.1 Hz), 4.90-4.86 (m, 1H), 4.14 (t, 2H, *J* = 5.6 Hz), 3.53-3.50 (m, 2H), 3.50-3.43 (m, 3H), 3.17 (dd, 1H, *J* = 17.0, 7.1 Hz), 1.46 (s, 9H); $^{13}$C NMR ($CD_3OD$, 150 MHz) $\delta_C$ 167.2, 157.1, 156.0 (d, *J* = 2.6 Hz), 152.3 (d, *J* = 246.1 Hz), 148.5 (d, *J* = 10.9 Hz), 130.6, 125.7, 123.3 (d, *J* = 3.4 Hz), 122.6 (d, *J* = 7.0 Hz), 114.5, 113.7 (d, *J* = 20.0 Hz), 79.6, 78.9, 67.9, 42.2, 39.4, 37.3, 27.3; HRMS (ES+) calcd for $C_{20}H_{26}O_5N_3FNa$ [M+Na]$^+$ 430.1749, found 430.1745.

**tert-butyl (4-(4-(5-(acrylamidomethyl)-4,5-dihydroisoxazol-3-yl)-2-fluorophenoxy)butyl) carbamate (7)**

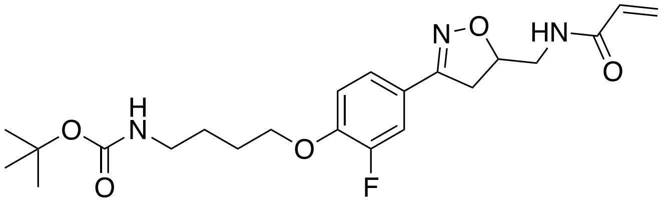

To phenol **5** (100 mg, 0.379 mmol) in DMF (10 mL) was added 4-(Boc-amino)butyl bromide (105 mg, 0.416 mmol), followed by $K_2CO_3$ (63.0 mg, 0.456 mmol). The resultant mixture was stirred at 70 °C for 18 h. After this time, the solvent was removed *in vacuo* and purification by column chromatography (gradient elution from $CH_2Cl_2$ to 8% MeOH in $CH_2Cl_2$) afforded the target product as an off white solid (128 mg, 0.294 mmol, 78%).

$^1$H NMR ($CD_3OD$, 600 MHz) $\delta_H$ 7.46 (dd, 1H, *J* = 12.1, 2.2 Hz), 7.38 (br d, 1H, *J* = 8.5 Hz), 7.14 (t, 1H, *J* = 8.6 Hz), 6.34-6.18 (m, 2H), 5.68 (dd, 1H, *J* = 9.5, 2.5 Hz), 4.89-4.86 (m, 1H), 4.12 (t, 2H, *J* = 6.3 Hz), 3.52 (d, 2H, *J* = 5.4 Hz), 3.46 (dd, 1H, *J* = 17.0, 10.5 Hz), 3.17 (dd, 1H, *J* = 16.9, 7.1 Hz), 3.13 (t, 2H, *J* = 6.9 Hz), 1.84 (quin, 2H, *J* = 6.7 Hz), 1.68 (quin, 2H, *J* = 7.1 Hz), 1.45 (s, 9H); $^{13}$C NMR ($CD_3OD$, 150 MHz) $\delta_C$ 167.2, 157.2, 156.1 (d, *J* = 2.5 Hz), 152.2 (d, *J* = 245.8 Hz), 151.4, 148.8 (d, *J* = 10.8 Hz), 130.4, 125.8, 123.3 (d, *J* = 3.4 Hz), 122.2

(d, $J$ = 6.9 Hz), 114.2, 113.6 (d, $J$ = 20.2 Hz), 79.6, 78.5, 68.6, 42.2, 39.6, 37.3, 27.4, 26.1; HRMS (ES+) calcd for $C_{22}H_{31}O_5N_3F$ $[M+H]^+$ 436.2242, found 436.2239.

***tert*-butyl (5-(4-(5-(acrylamidomethyl)-4,5-dihydroisoxazol-3-yl)-2-fluorophenoxy)pentyl) carbamate (8)**

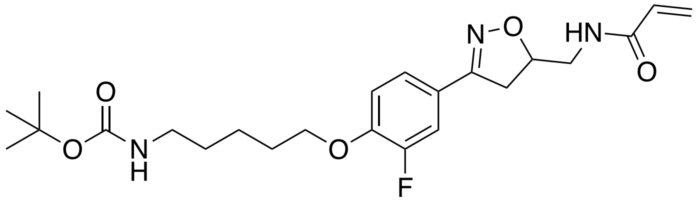

To phenol **5** (100 mg, 0.379 mmol) in DMF (10 mL) was added 5-(t-Boc-amino)-1-pentyl bromide (112 mg, 0.421 mmol), followed by $K_2CO_3$ (64.0 mg, 0.463 mmol). The resultant mixture was stirred at 70 °C for 18 h. After this time, the solvent was removed *in vacuo* and purification by column chromatography (gradient elution from $CH_2Cl_2$ to 8% MeOH in $CH_2Cl_2$) afforded the target product as an off white solid (144 mg, 0.321 mmol, 85%).

$^1$H NMR ($CDCl_3$ with a drop of $CD_3OD$, 600 MHz) $\delta_H$ 7.35 (dd, 1H, $J$ = 11.9, 2.1 Hz), 7.20 (d, 1H $J$ = 8.5 Hz), 6.88 (t, 1H, $J$ = 8.4 Hz), 6.24-6.16 (m, 1H), 6.08 (dd, 1H, $J$ = 17.0, 10.3 Hz), 5.64-5.53 (m, 1H), 4.82-4.78 (m, 1H), 3.98 (t, 2H, $J$ = 6.4 Hz), 3.55 (dd, 1H, $J$ = 14.3, 3.5 Hz), 3.47 (dd, 1H, $J$ = 14.3, 6.1 Hz), 3.31 (dd, 1H, $J$ = 16.7, 10.7 Hz), 3.07-2.99 (m, 3H), 1.77 (quin, 2H, $J$ = 6.7 Hz), 1.50-1.41 (m, 4H), 1.36 (s, 9H); $^{13}$C NMR ($CDCl_3$ with a drop of $CD_3OD$, 150 MHz) $\delta_C$ 166.8, 156.4 (overlapped), 156.4 (d, $J$ = 2.7 Hz), 152.3 (d, $J$ = 247.1 Hz), 149.0 (d, $J$ = 10.6 Hz), 130.3, 127.0, 123.4 (d, $J$ = 3.3 Hz), 121.8 (d, $J$ = 6.6 Hz), 114.3 (d, $J$ = 4.4 Hz), 114.3 (overlapped), 114.2 (d, $J$ = 15.7 Hz), 79.8, 79.3, 69.1, 53.4, 42.2, 37.7, 28.7, 28.3, 23.2; HRMS (ES+) calcd for $C_{23}H_{33}O_5N_3F$ $[M+H]^+$ 450.2399, found 450.2399.

***tert*-butyl (7-(4-(5-(acrylamidomethyl)-4,5-dihydroisoxazol-3-yl)-2-fluorophenoxy)heptyl) carbamate (9)**

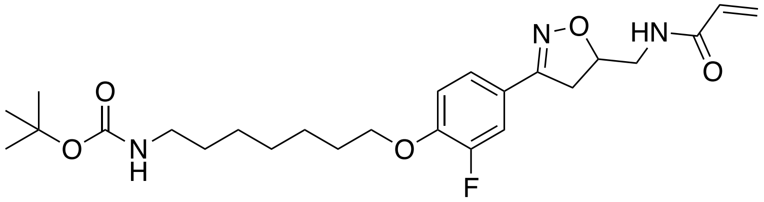

To phenol **5** (100 mg, 0.379 mmol) in DMF (10 mL) was added tert-butyl (7-bromoheptyl)carbamate (124 mg, 0.421 mmol), followed by $K_2CO_3$ (64.0 mg, 0.463 mmol). The resultant mixture was stirred at 70 °C for 18 h. After this time, the solvent was removed *in vacuo* and purification by column chromatography (gradient elution from $CH_2Cl_2$ to 8% MeOH in $CH_2Cl_2$) afforded the target product as a white solid (154 mg, 0.323 mmol, 85%).

$^1$H NMR (DMSO-d6, 600 MHz) $\delta_H$ 8.39 (t, 1H, $J$ = 6.0 Hz), 7.48 (dd, 1H, $J$ = 12.3, 2.1 Hz), 7.39 (br d, 1H, $J$ = 8.5 Hz), 7.24 (t, 1H, $J$ =8.7 Hz), 6.75 (t, 1H, $J$ =6.1 Hz), 6.27 (dd, 1H, $J$ = 17.1, 10.2 Hz), 6.10 (dd, 1H, $J$ = 17.1, 2.1 Hz), 5.60 (dd, 1H, $J$ =10.2, 2.1 Hz), 4.79-4.73 (m, 1H), 4.09 (t, 2H, $J$ = 6.5 Hz), 3.49-3.41 (m, 1H), 3.39-3.34 (m, 2H), 3.11 (dd, 1H, $J$ = 17.1, 7.2 Hz), 2.92-2.88 (m, 2H), 1.73 (quin, 2H, $J$ = 6.6 Hz), 1.46-1.38 (m, 3H), 1.37 (s,

9H), 1.34-1.29 (m, 2H), 1.29-1.20 (m, 3H); $^{13}$C NMR ($CD_3OD$, 150 MHz) $\delta_C$ 167.2, 157.2, 156.1 (d, *J* = 2.7 Hz), 152.3 (d, *J* = 245.7 Hz), 148.9 (d, *J* = 10.6 Hz), 130.4, 125.7, 123.2 (d, *J* = 3.3 Hz), 122.1 (d, *J* = 6.8 Hz), 114.2, 113.5 (d, *J* = 20.3 Hz), 79.6, 78.4, 68.9, 42.2, 39.9, 37.4, 29.5, 28.7, 28.7, 27.4, 26.4, 25.6; HRMS (ES+) calcd for $C_{25}H_{37}O_5N_3F$ $[M+H]^+$ 478.2712, found 478.2709.

***tert*-butyl (2-(2-(2-(2-(4-(5-(acrylamidomethyl)-4,5-dihydroisoxazol-3-yl)-2-fluorophenoxy) ethoxy)ethoxy) ethoxy)ethyl)carbamate (10)**

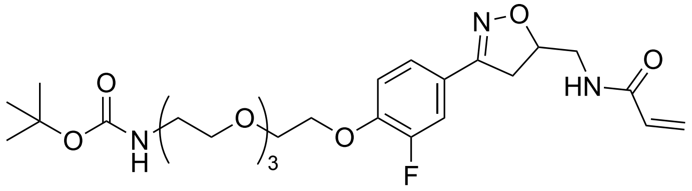

To phenol **5** (50.0 mg, 0.189 mmol) in DMF (5 mL) was added *N*-Boc-PEG3-bromide (60.0 µL, 0.208 mmol), followed by $K_2CO_3$ (31.0 mg, 0.224 mmol). The resultant mixture was stirred at 70 °C for 18 h. After this time, the solvent was removed *in vacuo* and purification by column chromatography (gradient elution from $CH_2Cl_2$ to 6% MeOH in $CH_2Cl_2$) afforded the target product as a white solid (67.0 mg, 0.124 mmol, 66%).

$^1$H NMR ($CDCl_3$, 600 MHz) $\delta_H$ 7.46 (dd, 1H, *J* = 11.9, 2.1 Hz), 7.29 (overlapped d, 1H, *J* = 8.6 Hz), 7.00 (t, 1H, *J* = 8.5 Hz), 6.30 (dd, 1H, *J* = 17.0, 1.3 Hz), 6.11 (dd, 1H, *J*=17.0, 10.3 Hz), 6.06 (br s, 1H), 5.67 (dd, 1H, *J* = 10.3, 1.3 Hz), 5.03 (br s, 1H), 4.93-4.88 (m, 1H), 4.26-4.23 (m, 2H), 3.93-3.90 (m, 2H), 3.78-3.75 (m, 2H), 3.71-3.67 (m, 3H), 3.68-3.66 (m, 4H), 3.65-3.62 (m, 3H), 3.56-3.54 (m, 2H), 3.38 (dd, 1H, *J* = 16.7, 10.6 Hz), 3.09 (dd, 1H, *J* = 16.8, 7.4 Hz), 1.46 (s, 9H); $^{13}$C NMR ($CDCl_3$, 150 MHz) $\delta_C$ 166.0, 156.1 (app d, *J* = 2.3 Hz), 156.0 (overlapped), 155.4 (d, *J* = 247.3 Hz), 148.7 (d, *J* = 10.8 Hz), 130.4, 127.2, 123.2 (d, *J* = 3.2 Hz), 122.5 (app d, *J* = 7.1 Hz), 114.7, 114.4 (d, *J* = 20.1 Hz), 79.9, 79.8, 72.7, 71.0, 70.6, 70.6, 70.5, 70.3, 70.2, 69.5, 69.0, 68.5, 42.2, 37.6, 28.4; HRMS (ES+) calcd for $C_{26}H_{38}O_8N_3FNa$ $[M+Na]^+$ 562.2533, found 562.2542.

***N*-((3-(4-(2-(2-((*S*)-4-(4-chlorophenyl)-2,3,9-trimethyl-6*H*-thieno[3,2-*f*][1,2,4]triazolo[4,3-*a*] [1,4]diazepin-6-yl)acetamido)ethoxy)-3-fluorophenyl)-4,5-dihydroisoxazol-5-yl)methyl) acrylamide (NF129)**

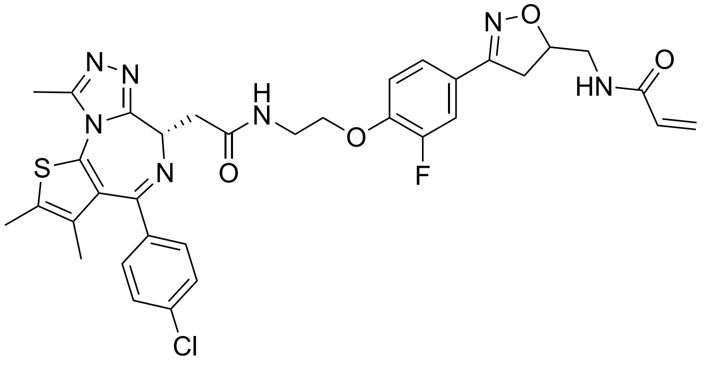

To **6** (59.0 mg, 0.145 mmol) in $CH_2Cl_2$ (2 mL) was added TFA (1 mL). The resultant mixture was stirred at RT for 30 min. It was then concentrated *in vacuo*, dissolved in DMF (4 mL) and added to a pre-stirred mixture of JQ1-acid (39.0 mg, 0.0972 mmol), HATU (41.0 mg, 0.107 mmol) and DIPEA (37.0 µL, 0.212 mmol) in DMF (4 mL). The resultant mixture was stirred at RT for 20 h. It was then concentrated *in vacuo* and purified by column chromatography (gradient elution $CH_2Cl_2$ to 10% MeOH in $CH_2Cl_2$) to afford the target product as an off white solid (39.0 mg, 0.0566 mmol, 58%).

$^1H$ NMR (DMSO-d6, 600 MHz) $\delta_H$ 8.56 (t, 1H, *J* = 5.9 Hz), 8.39 (t, 1H, *J* = 6.0 Hz), 7.52 (dt, 1H, *J* = 12.2, 1.9 Hz), 7.42 (br d, 1H, *J* = 9.6 Hz), 7.40-7.36 (m, 2H), 7.35-7.27 (m, 3H), 6.27 (dd, 1H, *J* = 17.1, 10.2 Hz), 6.11 (dd, 1H, *J* = 17.1, 2.1 Hz), 5.60 (dd, 1H, *J* = 10.2, 2,1 Hz), 4.81-4.76 (m, 1H), 4.53 (dd, 1H, *J* = 8.7, 5.5 Hz), 4.18 (t, 2H, *J* = 15.1, 5.6 Hz), 3.64-3.59 (m, 2H), 3.50-3.43 (m, 2H), 3.37 (t, 2H, *J* = 6.3 Hz), 3.20 (dd, 1H, *J* = 15.1, 5.6 Hz), 3.14-3.10 (m, 1H), 2.60 (s, 3H), 2.42 (s, 3H), 1.61 (s, 3H); $^{13}C$ NMR (DMSO-d6, 150 MHz) $\delta_C$ 170.6, 165.5, 163.5, 156.1, 155.6, 151.9 (d, *J* = 244.9 Hz), 150.3, 148.3, 137.2, 135.6, 132.8, 131.9, 131.2, 130.6, 130.3, 130.0, 128.8, 126.0, 124.2, 122.9, 115.4, 114.3, 79.9, 68.3, 54.3, 42.4, 40.5, 38.7, 38.0, 14.5, 13.2, 11.8; HRMS (ES+) calcd for $C_{34}H_{34}O_4N_7ClFS$ $[M+H]^+$ 690.2060, found 690.2057.

**N-((3-(4-(4-(2-((S)-4-(4-chlorophenyl)-2,3,9-trimethyl-6H-thieno[3,2-f][1,2,4]triazolo[4,3-a][1,4]diazepin-6-yl)acetamido)butoxy)-3-fluorophenyl)-4,5-dihydroisoxazol-5-yl)methyl) acrylamide (NF90)**

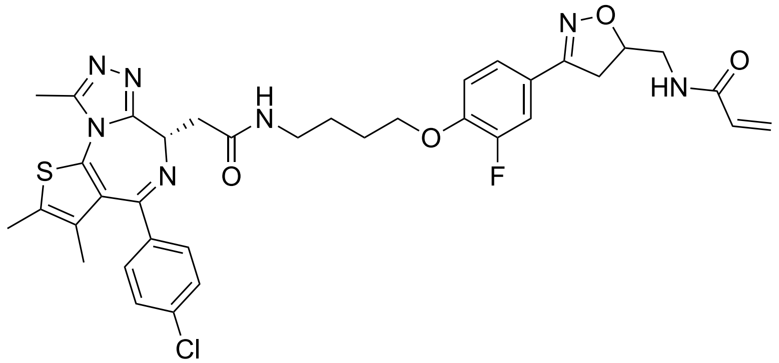


To **7** (40.0 mg, 0.0919 mmol) in $CH_2Cl_2$ (2 mL) was added TFA (1 mL). The resultant mixture was stirred at RT for 40 min. It was then concentrated *in vacuo*, dissolved in DMF (4 mL) and added to a pre-stirred mixture of JQ1-acid (28.0 mg, 0.0698 mmol), HATU (29.0 mg, 0.0763) and DIPEA (27.0 µL, 0.155 mmol) in DMF (4 mL). The resultant mixture was stirred at RT for 20 h. It was then concentrated *in vacuo* and purified by column chromatography (gradient elution $CH_2Cl_2$ to 10% MeOH in $CH_2Cl_2$) to afford the target product as a clear oil (46.0 mg, 0.0641 mmol, 92%).

$^1H$ NMR ($CD_3OD$ with a few drops of $CDCl_3$, 600 MHz) $\delta_H$ 8.42 (br s, 1H), 7.47-7.41 (m, 3H), 7.39-7.33 (m, 3H), 7.14 (t, 1H, *J* = 8.6 Hz), 6.32-6.20 (m, 2H), 5.67 (dd, 1H, *J* = 9.6, 2.4 Hz), 4.89-4.86 (m, 1H), 4.65 (ddd, 1H, *J* = 9.2, 5.1, 1.9 Hz), 4.14 (t, 2H, *J* = 6.3 Hz), 3.52 (d, 2H, *J* = 4.8 Hz), 3.48-3.38 (m, 3H), 3.32-3.22 (m, 2H), 3.16 (dd, 1H, *J* = 16.3, 7.8 Hz), 2.70 (s, 3H), 2.45 (s, 3H), 1.96-1.88 (m, 2H), 1.83-1.76 (m, 2H), 1.70 (s, 3H); $^{13}C$ NMR ($CD_3OD$, 150 MHz) $\delta_C$ 171.3, 167.2, 165.0, 156.1, 155.6, 152.2 (d, *J* = 245.9 Hz), 150.9, 148.8 (d, *J* = 11.2 Hz),

136.7, 136.5, 132.1, 132.0, 130.7, 130.6, 130.6, 129.9, 128.4, 125.8, 123.3 (d, *J* = 3.4 Hz), 122.2 (d, *J* = 7.0 Hz), 114.2, 113.6 (d, *J* = 20.5 Hz), 79.6, 68.6, 53.8, 42.2, 38.7, 37.4, 37.3, 26.2, 25.7, 13.0, 11.5, 10.2; HRMS (ES+) calcd for $C_{36}H_{38}O_4N_7ClFS$ $[M+H]^+$ 718.2373, found 718.2379.

***N*-((3-(4-((5-(2-((*S*)-4-(4-chlorophenyl)-2,3,9-trimethyl-6*H*-thieno[3,2-*f*][1,2,4]triazolo[4,3-*a*][1,4]diazepin-6-yl)acetamido)pentyl)oxy)-3-fluorophenyl)-4,5-dihydroisoxazol-5-yl) methyl)acrylamide (NF369)**

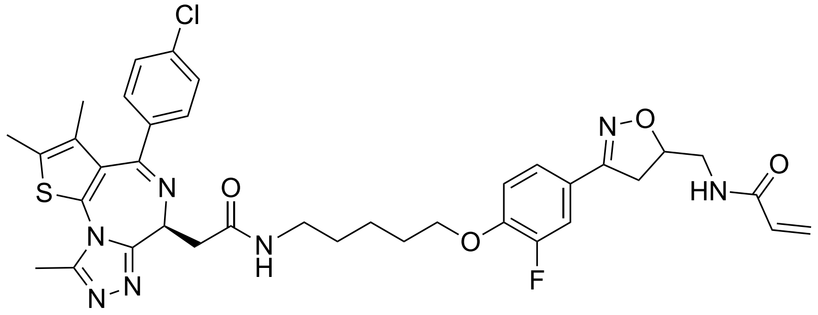

To **8** (20.0 mg, 0.0445 mmol) in $CH_2Cl_2$ (2 mL) was added TFA (1 mL). The resultant mixture was stirred at RT for 30 min. It was then concentrated *in vacuo*, dissolved in DMF (4 mL) and added to a pre-stirred mixture of JQ1-acid (14.0 mg, 0.0349 mmol), HATU (14.0 mg, 0.0368 mmol) and DIPEA (13.0 µL, 0.0746 mmol) in DMF (4 mL). The resultant mixture was stirred at RT for 20 h. It was then concentrated *in vacuo* and purified by column chromatography (gradient elution $CH_2Cl_2$ to 10% MeOH in $CH_2Cl_2$) to afford the target product as a clear oil (23.0 mg, 0.0315 mmol, 90%).

$^1$H NMR ($CD_3OD$ with a drop of $CDCl_3$, 600 MHz) $\delta_H$ 8.37 (t, 1H, *J* = 5.9 Hz), 7.47-7.42 (m, 3H), 7.39 (d, 2H, *J* = 8.7 Hz), 7.36 (br d, 1H, *J* = 8.5 Hz), 7.11 (t, 1H, *J* = 8.6 Hz), 6.30-6.22 (m, 2H), 5.67 (dd, 1H, *J* = 9.6, 2.4 Hz), 4.88-4.86 (m, 1H), 4.65 (dd, 1H, *J* = 9.0, 5.2 Hz), 4.11 (t, 2H, *J* = 6.4 Hz), 3.52 (d, 2H, *J* = 5.4 Hz), 3.49-3.41 (m, 2H), 3.30-3.24 (m, 1H), 3.16 (dd, 1H, *J* = 17.0, 7.2 Hz), 2.70 (s, 3H), 2.45 (s, 3H), 1.88 (quin, 2H, *J* = 6.8 Hz), 1.73-1.65 (m, 4H), 1.64-1.57 (m, 2H), 1.51 (dd, 1H, *J* = 6.6, 4.6 Hz), 1.40-1.137 (m, 2H); $^{13}$C NMR ($CD_3OD$ with a drop of $CDCl_3$, 150 MHz) $\delta_C$ 171.4, 167.2, 164.8, 156.1, 155.6, 152.2 (d, *J* = 245.8 Hz), 150.8, 148.8 (d, *J* = 10.7 Hz), 136.6, 136.6, 132.1, 131.8, 130.6, 130.6, 130.4, 129.9, 128.4, 125.7, 123.3, 122.2, 114.3, 113.6 (d, *J* = 20.5 Hz), 78.1, 68.9, 54.5, 53.9, 42.2, 39.0, 37.3, 28.4, 22.9, 17.3, 13.0, 11.5, 10.2; HRMS (ES+) calcd for $C_{37}H_{40}O_4N_7ClFS$ $[M+H]^+$ 732.2530, found 732.2522.

***N*-((3-(4-((7-(2-((*S*)-4-(4-chlorophenyl)-2,3,9-trimethyl-6*H*-thieno[3,2-*f*][1,2,4]triazolo[4,3-*a*][1,4]diazepin-6-yl)acetamido)heptyl)oxy)-3-fluorophenyl)-4,5-dihydroisoxazol-5-yl)methyl)acrylamide (NF370)**

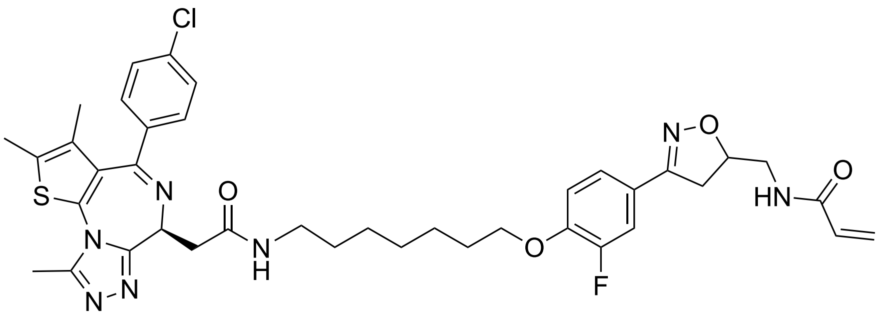

To **9** (20.0 mg, 0.0419 mmol) in $CH_2Cl_2$ (2 mL) was added TFA (1 mL). The resultant mixture was stirred at RT for 30 min. It was then concentrated *in vacuo*, dissolved in DMF (4 mL) and added to a pre-stirred mixture of JQ1-acid (13.0 mg, 0.0324 mmol), HATU (13.0 mg, 0.0342 mmol) and DIPEA (12.0 µL, 0.0689 mmol) in DMF (4 mL). The resultant mixture was stirred at RT for 20 h. It was then concentrated *in vacuo* and purified by column chromatography (gradient elution $CH_2Cl_2$ to 10% MeOH in $CH_2Cl_2$) to afford the target product as a clear oil (21.0 mg, 0.0277 mmol, 85%).

$^1$H NMR ($CD_3OD$, 600 MHz) $\delta_H$ 7.48-7.46 (m, 2H), 7.45 (dd, 1H, *J* = 12.2, 2.2 Hz), 7.43-7.40 (m, 2H), 7.36 (br d, 1H, *J* = 8.4 Hz), 7.12 (t, 1H, *J* = 8.6 Hz), 6.30-6.22 (m, 2H), 5.68 (dd, 1H, *J* = 9.5, 2.4 Hz), 4.89-4.86 (m, 1H), 4.66 (dd, 1H, *J* = 9.0, 5.2 Hz), 4.09 (t, 2H, *J* = 6.4 Hz), 3.52 (d, 2H, *J* = 5.4 Hz), 3.49-3.40 (m, 2H), 3.28-3.24 (m, 1H), 3.16 (dd, 1H, *J* = 17.0, 7.2 Hz), 2.72 (s, 3H), 2.46 (s, 3H), 1.82 (quin, 2H, *J* = 7.4 Hz), 1.71 (s, 3H), 1.61 (quin, 2H, *J* = 7.2 Hz), 1.56-1.36 (m, 2H), 1.46-1.43 (m, 2H), 1.41-1.36 (m, 4H); $^{13}$C NMR ($CD_3OD$, 150 MHz) $\delta_C$ 171.2, 167.2, 165.0, 156.1, 155.5, 152.1 (app d, *J* = 243.5 Hz), 150.9, 148.8, 136.7, 136.5, 132.1, 130.7, 130.6, 130.4, 130.0, 128.4, 125.7, 123.3, 122.1, 114.3, 113.5 (d, *J* = 20.4 Hz), 79.6, 69.0, 54.5, 53.8, 42.2, 39.0, 37.3, 29.0, 26.5, 25.6, 17.3, 15.9, 13.0, 11.5, 10.2; HRMS (ES+) calcd for $C_{39}H_{44}O_4N_7ClFS$ $[M+H]^+$ 760.2843, found 760.2834.

***N*-((3-(4-((1-((*S*)-4-(4-chlorophenyl)-2,3,9-trimethyl-6*H*-thieno[3,2-*f*][1,2,4]triazolo[4,3-*a*][1,4]diazepin-6-yl)-2-oxo-6,9,12-trioxa-3-azatetradecan-14-yl)oxy)-3-fluorophenyl)-4,5-dihydroisoxazol-5-yl)methyl)acrylamide (NF91)**

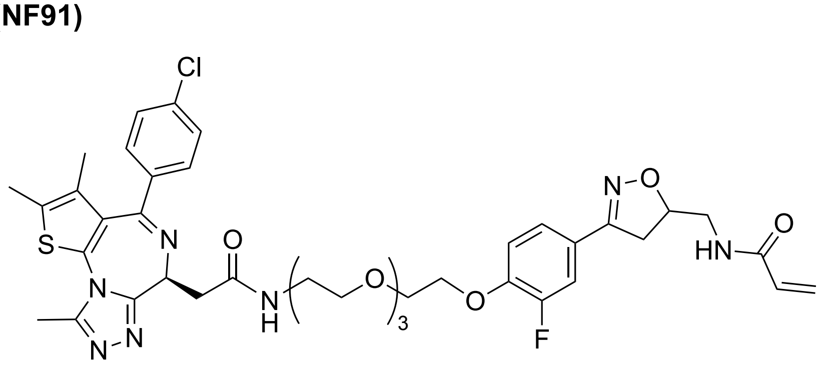

To **10** (36.0 mg, 0.0667 mmol) in $CH_2Cl_2$ (2 mL) was added TFA (1 mL). The resultant mixture was stirred at RT for 40 min. It was then concentrated *in vacuo*, dissolved in DMF (4 mL) and added to a pre-stirred mixture of

JQ1-acid (21.3 mg, 0.0531), HATU (22.0 mg, 0.0579 mmol) and DIPEA (21.0 µL, 0.121 mmol) in DMF (4 mL). The resultant mixture was stirred at RT for 20 h. It was then concentrated *in vacuo* and purified by column chromatography (gradient elution $CH_2Cl_2$ to 10% MeOH in $CH_2Cl_2$) to afford the target product as a clear oil (42.0 mg, 0.0511 mmol, 96%).

$^1$H NMR ($CD_3OD$, 600 MHz) $\delta_H$ 7.49-7.45 (m, 2H), 7.44-7.39 (m, 3H), 7.32 (br d, 1H, *J* = 8.4 Hz), 7.12 (t, 1H, *J* = 8.3 Hz), 6.30-6.21 (m, 2H), 5.67 (dd, 1H, *J* = 9.5, 2.5 Hz), 4.84-4.82 (m, 1H), 4.63 (dd, 1H, *J* = 9.1, 5.1 Hz), 4.25-4.22 (m, 2H), 3.89-3.87 (m, 2H), 3.75-3.72 (m, 2H), 3.71-3.64 (m, 6H), 3.64-3.61 (m, 2H), 3.52-3.38 (m, 6H), 3.32-3.28 (m, 1H), 3.14 (ddd, 1H, *J* = 17.0, 7.2, 2.4 Hz), 2.71 (s, 3H), 2.46 (s, 3H), 1.71 (s, 3H); $^{13}$C NMR ($CD_3OD$, 150 MHz) $\delta_C$ 171.5, 167.2, 164.8, 156.0, 155.6, 152.2 (d, *J* = 245.9 Hz), 150.8, 148.6, 136.7, 136.6, 132.1, 131.8, 130.7, 130.6, 130.4, 130.0, 128.4, 125.7, 123.2, 114.5, 113.6 (d, *J* = 20.1 Hz), 79.6, 70.5, 70.2, 70.2, 70.0, 69.2, 69.2, 68.9, 57.4, 53.8, 42.1, 39.2, 37.3, 13.0, 11.5, 10.2; HRMS (ES+) calcd for $C_{40}H_{46}O_7N_7ClFS$ $[M+H]^+$ 822.2846, found 822.2845.

***N*-((3-(3-fluoro-4-hydroxyphenyl)-4,5-dihydroisoxazol-5-yl)methyl)propionamide (11)**

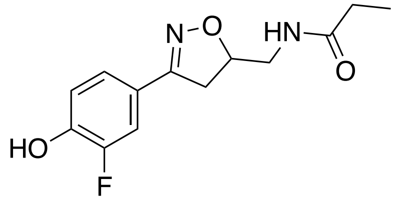


Reagent **4** (50.0 mg, 0.243 mmol) was dissolved in MeOH (5 mL) and degassed for 20 min. Aq. sat. HCl (71 µL) was then added, followed by palladium on carbon (35.0 mg, 10 wt.%). The resultant mixture was stirred under a $H_2$ atmosphere for 20 h, after which time it was filtered through a pad of celite and concentrated *in vacuo*. It was then dissolved in DMF (5 mL) and $NEt_3$ (120 µL, 0.864 mmol) was added, followed by propionyl chloride (50.0 µL, 0.576 mmol). The resultant mixture was stirred at RT for 18 h. The precipitate was filtered off and the filtrate was concentrated *in vacuo* before it was redissolved in a mixture of MeOH (10 mL) and 1M aq. NaOH (10 mL). After 1 h at RT, the mixture was acidified with 1M aq. HCl and extracted into EtOAc (3 × 30 mL). The combined organic extracts were washed with brine (30 mL) and dried ($MgSO_4$). Purification by column chromatography (gradient elution from $CH_2Cl_2$ to 5% MeOH in $CH_2Cl_2$) afforded the target product as a clear oil (28.0 mg, 0.105 mmol, 43%).

$^1$H NMR ($CD_3OD$ with a drop of $CDCl_3$, 600 MHz) $\delta_H$ 7.42 (dd, 1H, *J* = 12.0, 2.1 Hz), 7.30 (ddd, 1H, *J* = 8.4, 2.1, 1.0 Hz), 6.96 (t, 1H, *J* = 8.6 Hz), 4.83-4.79 (m, 1H), 3.46-3.40 (m, 3H), 3.15 (dd, 1H, *J* = 17.0, 7.0 Hz), 2.22 (q, 2H, *J* = 7.7 Hz), 1.10 (t, 3H, *J* = 7.7 Hz); $^{13}$C NMR ($CD_3OD$ with a drop of $CDCl_3$, 150 MHz) $\delta_C$ 176.3, 156.2 (d, *J* = 2.7 Hz), 151.4 (d, *J* = 241.5 Hz), 147.1 (d, *J* = 13.0 Hz), 123.3 (d, *J* = 3.1 Hz), 121.1 (d, *J* = 6.6 Hz), 117.5 (d, *J* = 3.3 Hz), 113.8 (d, *J* = 20.3 Hz), 79.5, 42.1, 37.3, 28.7, 9.1; HRMS (ES+) calcd $C_{13}H_{16}O_3N_2F$ $[M+H]^+$ 267.1139, found 267.1139.

***tert*-butyl (4-(2-fluoro-4-(5-(propionamidomethyl)-4,5-dihydroisoxazol-3-yl)phenoxy)butyl) carbamate (12)**

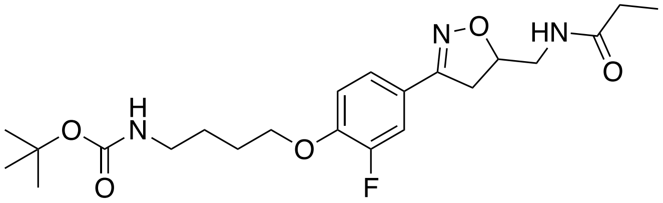

To phenol **11** (25 mg, 0.0939 mmol) in DMF (5 mL) was added 4-(Boc-amino)butyl bromide (26.0 mg, 0.103 mmol), followed by $K_2CO_3$ (16.0 mg, 0.116 mmol). The resultant mixture was stirred at 70 °C for 18 h. After this time, the solvent was removed *in vacuo* and purification by column chromatography (gradient elution from $CH_2Cl_2$ to 8% MeOH in $CH_2Cl_2$) afforded the target product as a white solid (35.0 mg, 0.0800 mmol, 85%).

$^1$H NMR ($CD_3OD$, 600 MHz) $\delta_H$ 7.46 (dd, 1H, *J* = 12.2, 2.1 Hz), 7.38 (br d, 1H, *J* = 8.5 Hz), 7.14 (t, 1H, *J* = 8.6 Hz), 6.63 (br s, 1H), 4.84-4.81 (m, 1H), 4.13 (t, 2H, *J* = 6.3 Hz), 3.47-3.40 (m, 3H), 3.18-3.12 (m, 3H), 2.22 (q, 2H, *J* = 7.6 Hz), 1.85 (quin, 2H, *J* = 6.4 Hz), 1.68 (quin, 2H, *J* = 7.1 Hz), 1.45 (s, 9H), 1.10 (t, 3H, *J* = 7.6 Hz); $^{13}$C NMR ($CD_3OD$, 150 MHz) $\delta_C$ 176.2, 157.3, 156.1 (d, *J* = 2.6 Hz), 152.2 (d, *J* = 245.6 Hz), 148.8 (d, *J* = 10.9 Hz), 123.6 (d, *J* = 3.3 Hz), 122.2 (d, *J* = 7.0 Hz), 114.3, 113.5 (d, *J* = 20.2 Hz), 79.7, 78.5, 68.8, 42.1, 39.7, 39.6, 37.2, 28.7, 27.4, 26.1, 9.1; HRMS (ES+) calcd for $C_{22}H_{33}O_5N_3F$ $[M+H]^+$ 438.2399, found 438.2400.

***N*-((3-(4-(4-(2-((*S*)-4-(4-chlorophenyl)-2,3,9-trimethyl-6*H*-thieno[3,2-*f*][1,2,4]triazolo[4,3-*a*][1,4]diazepin-6-yl)acetamido)butoxy)-3-fluorophenyl)-4,5-dihydroisoxazol-5-yl)methyl) propionamide (NF457)**

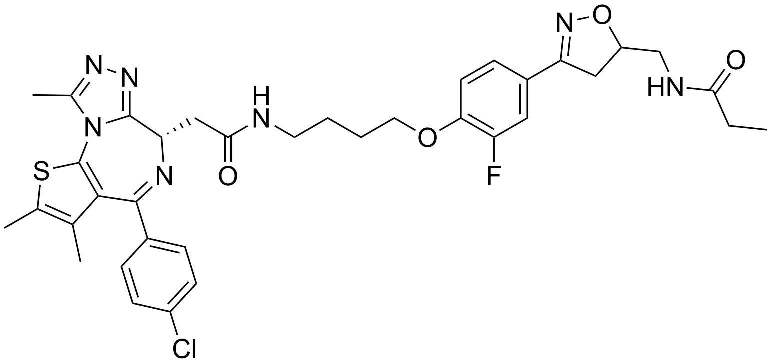

To reagent **12** (35.0 mg, 0.0827 mmol) in $CH_2Cl_2$ (4 mL) was added TFA (2 mL). The resultant mixture was stirred at RT for 40 min. It was then concentrated *in vacuo*, dissolved in DMF (4 mL) and added to pre-stirred mixture of JQ1-acid (26.0 mg, 0.0649 mml), HATU (37.0 mg, 0.0973 mmol) and DIPEA (37.0 µL, 0.211 mmol) in DMF (4 mL). The resultant mixture was stirred at RT for 20 h. It was then concentrated *in vacuo* and purified by column chromatography (gradient elution $CH_2Cl_2$ to 10% MeOH in $CH_2Cl_2$) to afford the target product as a clear oil (45.0 mg, 0.0626 mmol, 96%).

$^1$H NMR ($CD_3OD$ with a drop of $CDCl_3$, 600 MHz) $\delta_H$ 8.44-8.41 (m, 1H), 7.48-7.41 (m, 3H), 7.39-7.31 (m, 3H), 7.14 (t, 1H, *J* = 8.5 Hz), 4.85-4.79 (m, 1H), 4.67-4.63 (m, 1H), 4.14 (t, 2H, *J* = 6.3 Hz), 3.48-3.39 (m, 5H), 3.39-3.32 (m, 1H), 3.31-3.26 (m, 1H), 3.14 (dd, 1H, *J* = 17.0, 7.0 Hz), 2.70 (s, 3H), 2.45 (s, 3H), 2.22 (q, 2H, *J* = 7.7

Hz), 1.94-1.89 (m, 2H), 1.81-1.76 (m, 2H), 1.70 (s, 3H), 1.10 (t, 3H, *J* = 7.6 Hz); $^{13}$C NMR ($CD_3OD$ with a drop of $CDCl_3$, 150 MHz) $\delta_C$ 176.2, 171.5, 164.8, 156.0 (d, *J* = 2.3 Hz), 155.6, 152.2 (d, *J* = 245.9 Hz), 150.8, 148.8 (d, *J* = 10.9 Hz), 136.7, 136.6, 132.1, 131.8, 130.7, 130.6, 129.9, 128.3, 123.3 (d, *J* = 3.3 Hz), 122.3 (d, *J* = 6.9 Hz), 120.6, 114.2, 113.5 (d, *J* = 20.1 Hz), 79.7, 78.1, 68.6, 53.9, 42.1, 38.8, 37.6, 37.2, 28.8, 26.2, 25.7, 13.0, 11.5, 10.2, 9.1; HRMS (ES$^+$) calcd For $C_{36}H_{40}O_4N_7ClFS$ [M+H]$^+$ 720.2530, found 720.2526.

**tert-butyl ((1r,4r)-4-(3-chloro-4-cyanophenoxy)cyclohexyl)carbamate (13)**

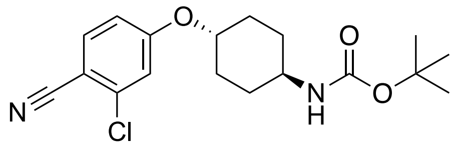

To a solution of trans-4-(Boc-amino)cyclohexanol (1.00 g, 4.65 mmol) in DMF (15 mL) at -10 °C was added NaH (223 mg, 5.58 mmol, 60% dispersion in mineral oil) and the resultant mixture was stirred for 1 h. 2-Chloro-4-fluorobenzonitrile (723 mg, 4.65 mmol) was then added and the mixture stirred at RT for further 18 h. It was then concentrated *in vacuo*, dissolved in EtOAc (50 mL), washed with $H_2O$ (30 mL), brine (30 mL) and dried ($MgSO_4$). Purification by column chromatography (gradient elution from hexanes to 20% EtOAc in hexanes) afforded the target product as a colorless oil (1.19 g, 3.40 mmol, 73%).

$^1$H NMR ($CD_3OD$ with a drop of $CDCl_3$, 600 MHz) $\delta_H$ 7.67 (d, 1H, *J* = 8.7 Hz), 7.15 (d, 1H, *J* = 2.4 Hz), 7.01 (dd, 1H, *J* = 8.8, 2.4 Hz), 4.42 (tt, 1H, *J* = 10.2, 4.1 Hz), 3.42 (td, 1H, *J* = 8.8, 5.4 Hz), 2.16-2.09 (m, 2H), 2.03-1.96 (m, 2H), 1.45 (s, 9H), 1.43-1.36 (m, 2H); $^{13}$C NMR ($CD_3OD$ with a drop of $CDCl_3$, 150 MHz) $\delta_C$ 162.2, 156.4, 137.6, 135.2, 116.7, 115.9, 114.7, 103.9, 78.6, 78.1, 75.7, 43.4, 29.6, 29.6, 27.4; HRMS (ES+) calcd for $C_{18}H_{23}O_3N_2ClNa$ [M+Na]$^+$ 373.1289, found 373.1286.

**6-chloro-*N*-((1r,4r)-4-(3-chloro-4-cyanophenoxy)cyclohexyl)pyridazine-3-carboxamide (14)**

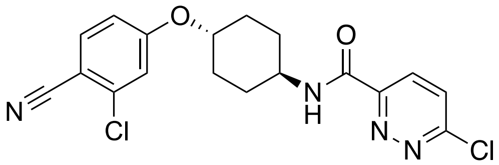

To **13** (2.00 g, 5.71 mmol) in $CH_2Cl_2$ (40 mL) was added TFA (20 mL) and the resultant mixture was stirred at RT for 4 h. It was then concentrated *in vacuo*, re-dissolved in $CH_2Cl_2$ (10 mL) and added to a pre-stirred mixture of 6-chloropyridazine-3-carboxylic acid (1.18 g, 7.43 mmol), T3P (4.1 mL, 6.85 mmol, 50% in EtOAc) and $NEt_3$ (3.18 mL, 22.8 mmol) in $CH_2Cl_2$ (15 mL). The resultant mixture was stirred at 30 °C for 16 h. It was then concentrated *in vacuo* and purified by column chromatography (gradient elution from hexanes to 80% EtOAc in hexanes) to afford the target product as a white crystalline solid (1.64 g, 4.20 mmol, 74%).

$^1$H NMR (DMSO-d6, 600 MHz) $\delta_H$ 9.19 (d, 1H, *J* = 8.2 Hz), 8.28 (d, 1H, *J* = 8.9 Hz), 8.15 (d, 1H, *J* = 8.9 Hz), 7.91 (d, 1H, *J* = 8.8 Hz), 7.44 (d, 1H, *J* = 2.4 Hz), 7.19 (dd, 1H, *J* = 8.8, 2.5 Hz), 4.62-4.56 (m, 1H), 4.00-3.93 (m, 1H),

2.21-2.15 (m, 2H), 1.99-1.93 (m, 2H), 1.81-1.71 (m, 2H), 1.62-1.53 (m, 2H); $^{13}C$ NMR (DMSO-d6, 150 MHz) $\delta_C$ 162.3, 161.7, 158.6, 153.5, 137.5, 136.2, 130.7, 129.5, 117.3, 116.9, 115.9, 103.7, 76.0, 48.2, 30.4, 29.6; HRMS (ES-) calcd for $C_{18}H_{15}O_2N_4Cl_2$ $[M-H]^+$ 389.0578, found 389.0581.

***tert*-butyl 4-(6-(((1r,4r)-4-(3-chloro-4-cyanophenoxy)cyclohexyl)carbamoyl)pyridazin-3-yl) piperazine-1-carboxylate (15)**

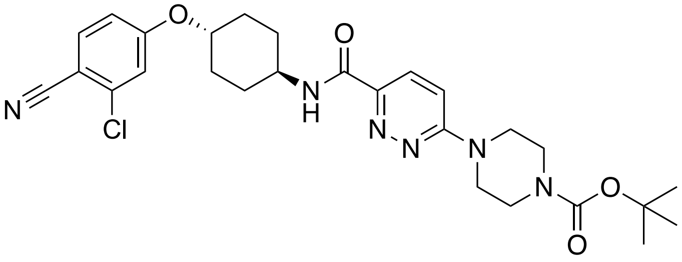

To **14** (500 mg, 1.32 mmol) in DMF (10 mL) was added 1-Boc-piperazine (367 mg, 1.97 mmol), followed by $NEt_3$ (552 µL, 3.96 mmol). The resultant mixture was stirred at 80 °C for 20 h. It was then cooled to RT and the precipitate was filtered off and dried to afford the target product as a white solid (411 mg, 0.761 mmol, 58%).
$^1H$ NMR (DMSO-d6 with a drop of $CDCl_3$, 600 MHz) $\delta_H$ 8.55 (d, 1H, $J$ = 8.2 Hz), 7.87 (d, 1H, $J$ = 9.5 Hz), 7.79 (d, 1H, $J$ = 8.8 Hz), 7.32 (d, 1H, $J$ = 9.6 Hz), 7.30 (d, 1H, $J$ = 2.4 Hz), 7.09 (dd, 1H, $J$ = 8.8, 2.4 Hz), 4.54-4.48 (m, 1H), 3.91-3.84 (m, 1H), 3.74-3.70 (m, 4H), 3.50-3.46 (m, 4H), 2.14-2.09 (m, 2H), 1.96-1.89 (m, 2H), 1.65 (qd, 2H, $J$ = 13.1, 3.1 Hz), 1.53 (qd, 2H, $J$ = 12.9, 3.4 Hz), 1.43 (s, 9H); $^{13}C$ NMR (DMSO-d6 with a drop of $CDCl_3$, 150 MHz) $\delta_C$ 162.8, 162.2, 160.5, 154.3, 145.4, 137.5, 136.0, 126.9, 117.2, 116.7, 115.7, 113.2, 103.8, 76.0, 47.5, 44.5, 40.6, 30.3, 29.9, 28.5; HRMS (ES+) calcd for $C_{27}H_{34}O_4N_6Cl$ $[M+H]^+$ 541.2325, found 541.2330.

***tert*-butyl 2-(4-(6-(((1r,4r)-4-(3-chloro-4-cyanophenoxy)cyclohexyl)carbamoyl)pyridazin-3-yl)piperazin-1-yl)acetate (16)**

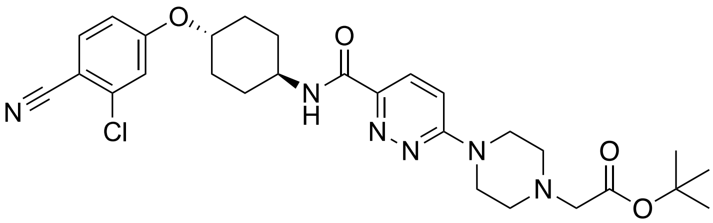

Reagent **15** (500 mg, 0.926 mmol) was dissolved in a mixture of $CH_2Cl_2$ (20 mL) and TFA (10 mL) and stirred at RT for 1 h. It was then concentrated *in vacuo* and re-dissolved in DMF (10 mL). To this was added tert-butyl bromoacetate (205 µL, 2.79 mmol), followed by $NEt_3$ (389 µL, 2.89 mmol) and the resultant mixture was stirred at 70 °C for 20 h. It was then concentrated *in vacuo* and purification by column chromatography (gradient elution from 50% EtOAc in hexanes to EtOAc) afforded the target compound as a white solid (394 mg, 0.711 mmol, 77%).

$^1H$ NMR ($CDCl_3$, 600 MHz) $\delta_H$ 7.99 (d, 1H, *J* = 9.5 Hz), 7.86 (d, 1H, *J* = 8.2 Hz), 7.55 (d, 1H, *J* = 8.7 Hz), 7.00 (d, 1H, *J* = 2.4 Hz), 6.97 (d, 1H, *J* = 9.5 Hz), 6.85 (dd, 1H, *J* = 8.7, 2.4 Hz), 4.36-4.29 (m, 1H), 4.08-4.01 (m, 1H), 3.83 (t, 4H, *J* = 5.1 Hz), 3.21 (s, 2H), 2.77 (t, 4H, *J* = 5.1 Hz), 2.20-2.15 (m, 4H), 1.71-1.65 (m, 2H), 1.51-1.43 (m, 11H); $^{13}C$ NMR ($CDCl_3$, 150 MHz) $\delta_C$ 169.1, 162.7, 161.6, 160.2, 144.5, 138.3, 135.1, 126.8, 116.9, 116.4, 114.6, 112.2, 104.7, 81.6, 75.7, 59.6, 52.2, 47.1, 44.6, 30.0, 29.6, 28.1; HRMS (ES+) calcd for $C_{28}H_{36}O_4N_6Cl$ $[M+H]^+$ 555.2481, found 555.2486.

**6-(4-(2-((4-(4-(5-(acrylamidomethyl)-4,5-dihydroisoxazol-3-yl)-2-fluorophenoxy)butyl) amino)-2-oxoethyl) piperazin-1-yl)-*N*-(4-(3-chloro-4-cyanophenoxy)cyclohexyl)pyridazine-3-carboxamide (NF500A)**

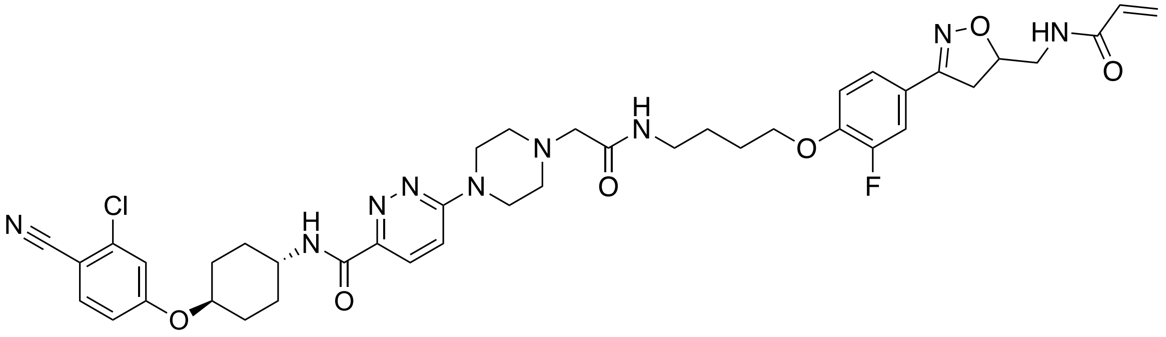

To **16** (30.0 mg, 0.054 mmol) in $CH_2Cl_2$ (4 mL) was added TFA (2 mL) and the resultant mixture was stirred at RT for 18 h. To **7** (30.0 mg, 0.069 mmol) in $CH_2Cl_2$ (4 mL) was also added TFA (2 mL) and the resultant mixture was stirred at RT for 1 h. Both mixtures were concentrated *in vacuo*. **7-amine** (0.069 mmol) was dissolved in DMF (4 mL) and added to pre-stirred mixture of **16-acid** (0.054 mmol), HATU (33.0 mg, 0.0823 mmol) and DIPEA (31.0 µL, 0.178 mmol) in DMF (4 mL). The resultant mixture was stirred at RT for 20 h. It was then concentrated *in vacuo* and purified by column chromatography (gradient elution $CH_2Cl_2$ to 10% MeOH in $CH_2Cl_2$) to afford the target product as a clear oil (35.0 mg, 0.0429 mmol, 79%).

$^1H$ NMR ($CDCl_3$, 600 MHz) $\delta_H$ 8.04 (d, 1H, *J* = 9.5 Hz), 7.88 (d, 1H, *J* = 8.2 Hz), 7.58 (d, 1H, *J* = 8.7 Hz), 7.45 (dd, 1H, *J* = 11.9, 2.1 Hz), 7.30-7.28 (m, 1H), 7.24 (br s, 1H), 7.02 (d, 1H, *J* = 2.4 Hz), 7.00-6.95 (m, 2H), 6.88 (dd, 1H, *J* = 8.8, 2.4 Hz), 6.30 (dd, 1H, *J* = 17.0, 1.3 Hz), 6.16-6.09 (m, 2H), 5.67 (dd, 1H, *J* = 10.3, 1.3 Hz), 4.93-4.88 (m, 1H), 4.37-4.31 (m, 1H), 4.13 (t, 2H, *J* = 6.0 Hz), 4.10-4.04 (m, 1H), 3.79 (br s, 4H), 3.73-3.67 (m, 1H), 3.67-3.60 (m, 1H), 3.43 (q, 2H, *J* = 6.7 Hz), 3.37 (dd, 1H, *J* = 16.8, 10.6 Hz), 3.13-3.07 (m, 2H), 2.71 (br s, 4H), 2.26-2.15 (m, 4H), 1.96-1.88 (m, 2H), 1.85-1.76 (m, 2H), 1.75-1.67 (m, 2H), 1.55-1.44 (m, 2H); $^{13}C$ NMR ($CDCl_3$, 150 MHz) $\delta_C$ 166.1, 162.6, 161.6, 160.2, 156.0, 152.3 (d, *J* = 247.1 Hz), 148.7 (d, *J* = 11.0 Hz), 138.3, 135.1, 130.4, 127.2, 127.0, 123.3 (d, *J* = 3.5 Hz), 122.4, 116.9, 116.4, 114.7, 114.3 (d, *J* = 20.0 Hz), 114.2, 112.3, 104.8, 79.9, 75.7, 68.8, 61.8, 53.0, 47.1, 44.8, 42.2, 38.5, 37.6, 30.0, 29.7, 26.5 (overlapped), 26.5; HRMS (ES+) calcd for $C_{41}H_{47}ClFN_9O_6$ $[M+H]^+$ 816.3363, found 816.3388.

**6-(4-(2-((5-(4-(5-(acrylamidomethyl)-4,5-dihydroisoxazol-3-yl)-2-fluorophenoxy)pentyl) amino)-2-oxoethyl)piperazin-1-yl)-*N*-(4-(3-chloro-4-cyanophenoxy)cyclohexyl)pyridazine-3-carboxamide (NF500B)**

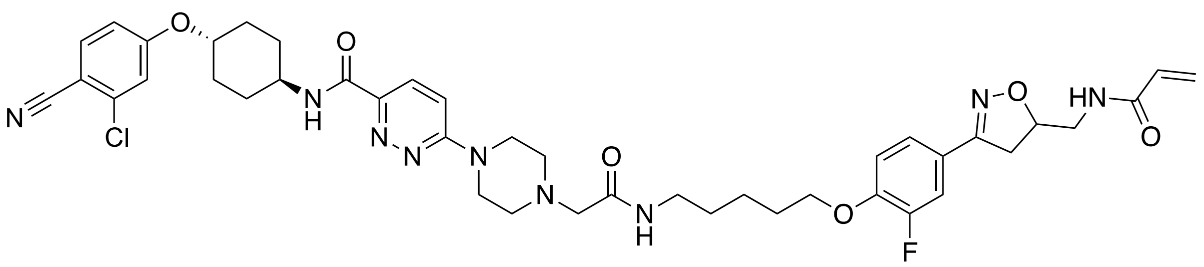

To **16** (30.0 mg, 0.0542 mmol) in $CH_2Cl_2$ (4 mL) was added TFA (2 mL) and the resultant mixture was stirred at RT for 18 h. To **8** (31 mg, 0.0690 mmol) in $CH_2Cl_2$ (4 mL) was also added TFA (2 mL) and the resultant mixture was stirred at RT for 1 h. Both mixtures were concentrated *in vacuo*. **8-amine** (0.0690 mmol) was dissolved in DMF (4 mL) and added to pre-stirred mixture of **16-acid** (0.0542 mmol), HATU (33.0 mg, 0.0823 mmol) and DIPEA (31.0 µL, 0.178 mmol) in DMF (4 mL). The resultant mixture was stirred at RT for 20 h. It was then concentrated *in vacuo* and purified by column chromatography (gradient elution $CH_2Cl_2$ to 10% MeOH in $CH_2Cl_2$) to afford the target product as a clear oil (37.0 mg, 0.0446 mmol, 82%).

$^1$H NMR ($CDCl_3$, 600 MHz) $\delta_H$ 8.01 (d, 1H, *J* = 9.5 Hz), 7.90 (d, 1H, *J* = 8.1 Hz), 7.58 (d, 1H, *J* = 8.7 Hz), 7.41 (dd, 1H, *J* =12.0, 2.1 Hz), 7.26 (br d, 1H, *J* = 8.4 Hz), 7.18 (br s, 1H), 7.03 (d, 1H, *J* = 2.4 Hz), 6.98-6.91 (m, 2H), 6.88 (dd, 1H, *J* = 8.7, 2.4 Hz), 6.34-6.29 (m, 2H), 6.15 (dd, 1H, *J* = 17.0, 10.3 Hz), 5.67 (dd, 1H, *J* = 10.3, 1.4 Hz), 4.94-4.89 (m, 1H), 4.37-4.33 (m, 1H), 4.11-4.05 (m, 3H), 3.78 (br s, 4H), 3.71-3.63 (m, 2H), 3.40-3.34 (m, 3H), 3.11-3.06 (m, 3H), 2.71 (br s, 4H), 2.25-2.18 (m, 4H), 1.92-1.87 (m, 2H), 1.76-1.66 (m, 4H), 1.60-1.55 (m, 2H), 1.54-1.47 (m, 2H); $^{13}$C NMR ($CDCl_3$, 150 MHz) $\delta_C$ 166.1, 162.6, 161.6, 156.0, 152.3 (d, *J* = 247.0 Hz), 148.8, 138.3, 135.1, 130.5, 127.1, 126.9, 123.3, 122.2, 116.9, 116.4, 114.7, 114.3, 114.2, 112.3, 104.8, 79.9, 75.6, 69.0, 61.6, 53.0, 47.2, 44.8, 42.1, 38.8, 37.5, 30.0, 29.7, 29.3, 28.7, 23.5; HRMS (ES+) $C_{42}H_{50}O_6N_9ClF$ $[M+H]^+$ 830.3551, found 830.3564.

**6-(4-(2-((7-(4-(5-(acrylamidomethyl)-4,5-dihydroisoxazol-3-yl)-2-fluorophenoxy) heptyl) amino)-2-oxoethyl)piperazin-1-yl)-*N*-((1*r*,4*r*)-4-(3-chloro-4-cyanophenoxy) cyclohexyl) pyridazine-3-carboxamide (NF500C)**

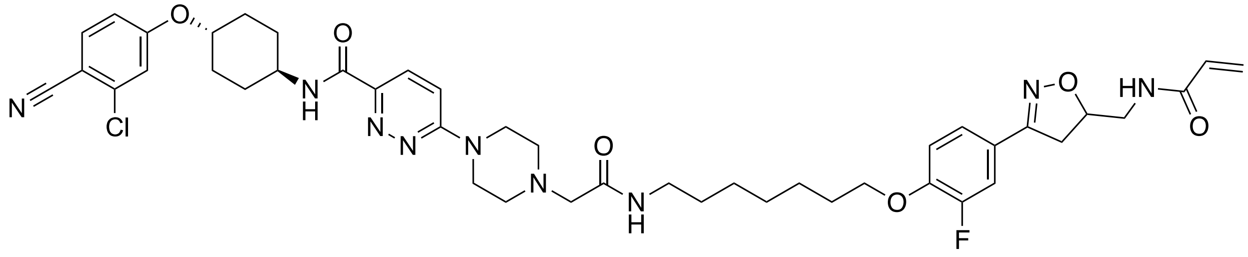

To **16** (30.0 mg, 0.0542 mmol) in $CH_2Cl_2$ (4 mL) was added TFA (2 mL) and the resultant mixture was stirred at RT for 18 h. To **9** (33 mg, 0.0690 mmol) in $CH_2Cl_2$ (4 mL) was also added TFA (2 mL) and the resultant mixture was stirred at RT for 1 h. Both mixtures were concentrated *in vacuo*. **9-amine** (0.0690 mmol) was dissolved in DMF (4 mL) and added to pre-stirred mixture of **16-acid** (0.0542 mmol), HATU (33.0 mg, 0.0823 mmol) and DIPEA (31.0 µL, 0.178 mmol) in DMF (4 mL). The resultant mixture was stirred at RT for 20 h. It was then concentrated *in vacuo* and purified by column chromatography (gradient elution $CH_2Cl_2$ to 10% MeOH in $CH_2Cl_2$) to afford the target product as a clear oil (39.0 mg, 0.0455 mmol, 84%).

$^1$H NMR ($CDCl_3$, 600 MHz) $\delta_H$ 8.02 (d, 1H, *J* = 9.5 Hz), 7.88 (d, 1H, *J* = 7.9 Hz), 7.57 (d, 1H, *J* = 8.8 Hz), 7.42 (dd, 1H, *J* = 12.0, 2.2 Hz), 7.30-7.25 (m, 1H), 7.18 (br s, 1H), 7.04-6.98 (m, 2H), 6.95 (t, 1H, *J* = 8.5 Hz), 6.87 (dd, 1H, *J* = 8.7, 2.5 Hz), 6.32-6.24 (m, 2H), 6.14 (dd, 1H, *J* = 17.0, 10.4 Hz), 5.66 (dd, 1H, *J* = 10.3, 1.4 Hz), 4.93-4.88 (m, 1H), 4.36-4.32 (m, 1H), 4.08-4.04 (m, 3H), 3.85 (br s, 4H), 3.71-3.66 (m, 1H), 3.65-3.60 (m, 1H), 3.39-3.34 (m, 1H), 3.34-3.28 (m, 2H), 3.20 (br s, 2H), 3.09 (dd, 1H, *J* = 16.8, 7.4 Hz), 2.81 (br s, 4H), 2.23-2.16 (m, 4H), 1.87-1.81 (m, 2H), 1.74-1.66 (m, 2H), 1.61-1.55 (m, 2H), 1.55-1.44 (m, 4H), 1.44-1.37 (m, 4H); $^{13}$C NMR ($CDCl_3$, 150 MHz) $\delta_C$ 166.1, 162.2, 161.6, 160.1, 156.1, 152.2 (d, *J* = 246.6 Hz), 148.9 (d, *J* = 10.8 Hz), 145.0, 138.3, 135.1, 130.5, 127.1, 127.0, 123.3 (d, *J* = 3.3 Hz), 122.1, 116.9, 116.4, 114.7, 114.3, 114.2, 112.4, 104.8, 79.8, 75.6, 69.3, 68.5, 61.2, 58.8, 54.5, 52.9, 47.2, 44.5, 42.2, 39.1, 37.6, 30.0, 29.7, 29.5, 28.9, 28.8, 26.8, 25.8, 18.6, 17.3; HRMS (ES+) calcd for $C_{44}H_{54}O_6N_9ClF$ $[M+H]^+$ 858.3864, found 858.3856.

***N*-((1*r*,4*r*)-4-(3-chloro-4-cyanophenoxy)cyclohexyl)-6-(4-methylpiperazin-1-yl)pyridazine-3-carboxamide (NF505)**

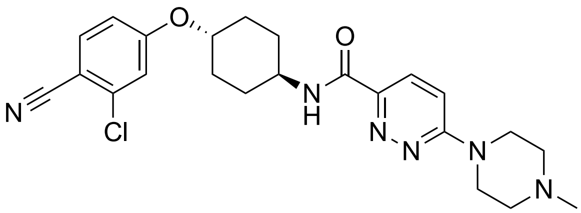


To **14** (200 mg, 0.513 mmol) in DMF (5 mL) was added 1-methylpiperazine (86.0 µL, 0.769 mmol) followed by $NEt_3$ (213 µL, 1.53 mmol). The resultant mixture was stirred at 80 °C for 18 h. It was then concentrated *in vacuo* and purified by column chromatography (gradient elution from $CH_2Cl_2$ to 10% MeOH in $CH_2Cl_2$) to afford the target product as an off white solid (202 mg, 0.445 mmol, 87%).

$^1$H NMR ($CDCl_3$, 600 MHz) $\delta_H$ 8.01 (d, 1H, *J* = 9.5 Hz), 7.87 (d, 1H, *J* = 8.1 Hz), 7.56 (d, 1H, *J* = 8.7 Hz), 7.04-6.96 (m, 2H), 6.86 (dd, 1H, *J* = 8.7, 2.4 Hz), 4.33 (tt, 1H, *J* = 9.9, 3.8 Hz), 4.06 (dtd, 1H, J = 10.8, 7.5, 4.0 Hz), 3.83 (t, 4H, *J* = 5.1 Hz), 2.63 (t, 4H, *J* = 5.1 Hz), 2.41 (s, 3H), 2.22-2.15 (m, 4H), 1.73-1.65 (m, 2H), 1.51-1.44 (m, 2H); $^{13}$C NMR ($CDCl_3$, 150 MHz) $\delta_C$ 162.7, 161.6, 160.2, 144.6, 138.3, 135.1, 126.9, 116.9, 116.4, 114.6, 112.2, 104.8, 75.7, 54.4, 47.1, 45.9, 44.5, 30.0, 29.7; HRMS (ES+) calcd for $C_{23}H_{28}O_2N_6Cl$ $[M+H]^+$ 455.1957, found 455.1962.

***tert*-butyl (7-(2-fluoro-4-(5-(propionamidomethyl)-4,5-dihydroisoxazol-3-yl)phenoxy)heptyl) carbamate (17)**

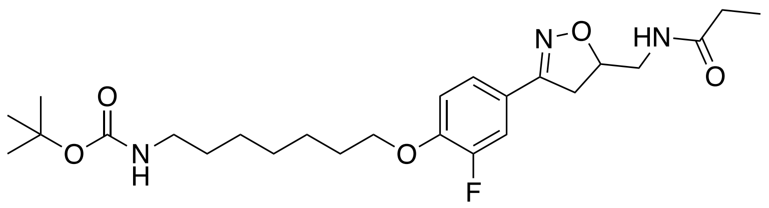

To phenol **11** (23.0 mg, 0.0864 mmol) in DMF (5 mL) was added tert-Butyl (7-bromoheptyl)carbamate (28.0 mg, 0.0951 mmol), followed by $K_2CO_3$ (14.0 mg, 0.101 mmol). The resultant mixture was stirred at 70 °C for 18 h. After this time, the solvent was removed *in vacuo* and purification by column chromatography (gradient elution from $CH_2Cl_2$ to 10% MeOH in $CH_2Cl_2$) afforded the target product as a white solid (33.0 mg, 0.0689 mmol, 80%). $^1$H NMR ($CDCl_3$, 600 MHz) $\delta_H$ 7.45 (dd, 1H, *J* = 11.9, 2.1 Hz), 7.28 (br d, 1H, *J* = 8.5 Hz), 6.95 (t, 1H, *J* = 8.5 Hz), 5.96 (t, 1H, *J* = 6.1 Hz), 4.88-4.84 (m, 1H), 4.53 (br s, 1H), 4.06 (t, 2H, *J* = 6.5 Hz), 3.62-3.53 (m, 2H), 3.09 (dd, 1H, *J* = 15.1, 7.4 Hz), 3.13-3.11 (m, 2H), 3.08 (dd, 1H, *J* = 16.8, 7.2 Hz), 2.23-2.19 (m, 2H), 1.84 (quin, 2H, *J* = 6.6 Hz), 1.52-1.48 (m, 4H), 1.45 (s, 9H), 1.41-1.34 (m, 4H), 1.11 (t, 3H, *J* = 7.6 Hz); $^{13}$C NMR ($CDCl_3$, 150 MHz) $\delta_C$ 174.5, 156.1 (d, *J* = 2.6 Hz), 156.0, 152.4 (d, *J* = 247.1 Hz), 149.0 (d, *J* = 10.8 Hz), 123.2 (d, *J* = 3.4 Hz), 122.0 (d, *J* = 7.0 Hz), 114.3, 114.2 (d, *J* = 2.2 Hz), 79.9, 79.1, 69.3, 42.2, 40.6, 37.6, 30.0, 29.7, 29.0, 29.0, 28.4, 26.7, 25.8, 9.8; HRMS (ES+) calcd for $C_{25}H_{39}O_5N_3F$ $[M+H]^+$ 480.2868, found 480.2873.

***N*-((1*r*,4*r*)-4-(3-chloro-4-cyanophenoxy)cyclohexyl)-6-(4-(2-((7-(2-fluoro-4-(5-(propiona midomethyl)-4,5-dihydroisoxazol-3-yl)phenoxy)heptyl)amino)-2-oxoethyl)piperazin-1-yl) pyridazine-3-carboxamide (NF534)**

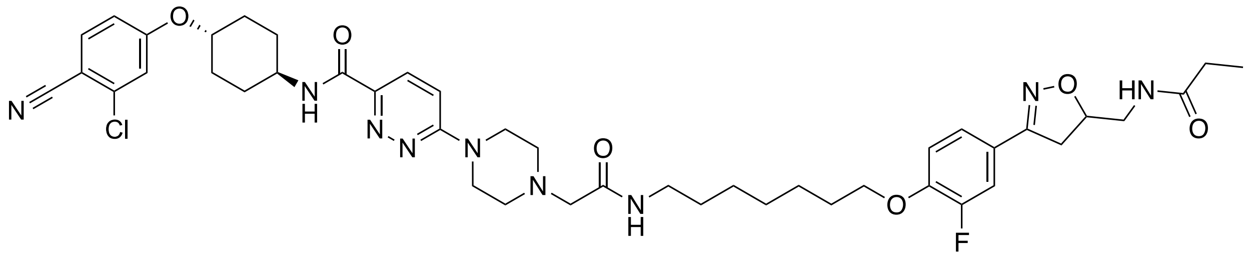

To **16** (62.0 mg, 0.112 mmol) in $CH_2Cl_2$ (10 mL) was added TFA (5 mL) and the resultant mixture was stirred at RT for 18 h. To **17** (70.0 mg, 0.146 mmol) in $CH_2Cl_2$ (10 mL) was also added TFA (5 mL) and the resultant mixture was stirred at RT for 1 h. Both mixtures were concentrated *in vacuo*. **17-amine** (0.146 mmol) was dissolved in DMF (4 mL) and added to pre-stirred mixture of **16-acid** (0.112 mmol), HATU (66.0 mg, 0.165 mmol) and DIPEA (63.0 µL, 0.362 mmol) in DMF (4 mL). The resultant mixture was stirred at RT for 20 h. It was then concentrated *in vacuo* and purified by column chromatography (gradient elution $CH_2Cl_2$ to 10% MeOH in $CH_2Cl_2$) to afford the target product as a white solid (56.0 mg, 0.0652 mmol, 58%).

$^{1}H$ NMR ($CDCl_3$ with a few drops of $CD_3OD$, 600 MHz) $\delta_H$ 7.94 (d, 1H, $J$ = 8.1 Hz), 7.89 (d, 1H, $J$ = 9.5 Hz), 7.49 (d, 1H, $J$ = 8.7 Hz), 7.33 (d, 1H, $J$ = 12.0 Hz), 7.27 (overlapped br s, 1H), 7.19 (d, 1H, $J$ = 8.4 Hz), 6.98 (d, 1H, $J$ = 9.6 Hz), 6.96-6.91 (m, 2H), 6.88 (t, 1H, $J$ = 8.5 Hz), 6.80 (dd, 1H, $J$ = 8.8, 2.1 Hz), 4.78-4.72 (m, 1H), 4.31-4.25 (m, 1H), 3.98 (t, 2H, $J$ = 6.41 Hz), 3.95-3.93 (m, 1H), 3.72 (dr s, 4H), 3.60-3.55 (m, 1H), 3.45-3.36 (m, 2H), 3.32-3.24 (m, 2H), 3.10-3.02 (m, 2H), 3.01-2.95 (m, 1H), 2.64 (br s, 4H), 2.16-2.06 (m, 6H), 1.77-1.70 (m, 2H), 1.65-1.55 (m, 2H), 1.51-1.45 (m, 2H), 1.45-1.39 (m, 4H), 1.36-1.30 (m, 4H), 1.01 (t, 3H, $J$ = 7.6 Hz); $^{13}C$ NMR ($CDCl_3$ with a few drops of $CD_3OD$, 150 MHz) $\delta_C$ 175.7, 163.0, 161.7, 160.1, 156.2, 152.2 (d, $J$ = 246.7 Hz), 148.9 (d, $J$ = 10.9 Hz), 144.5, 138.2, 135.1, 126.9, 123.3 (d, $J$ = 3.2 Hz), 121.8 (d, $J$ = 7.0 Hz), 116.9, 116.4, 116.4, 114.2, 114.1 (d, $J$ = 20.3 Hz), 112.6, 104.3, 79.8, 75.5, 69.2, 54.7, 52.3, 47.2, 44.6, 42.7, 42.3, 42.2, 38.9, 37.6, 29.7, 29.5, 29.3, 29.3, 28.8, 28.8, 26.7, 25.7, 18.2, 16.8, 12.3, 9.7; HRMS (ES+) calcd for $C_{44}H_{56}O_6N_9ClF$ $[M+H]^+$ 860.4021, found 860.4029.

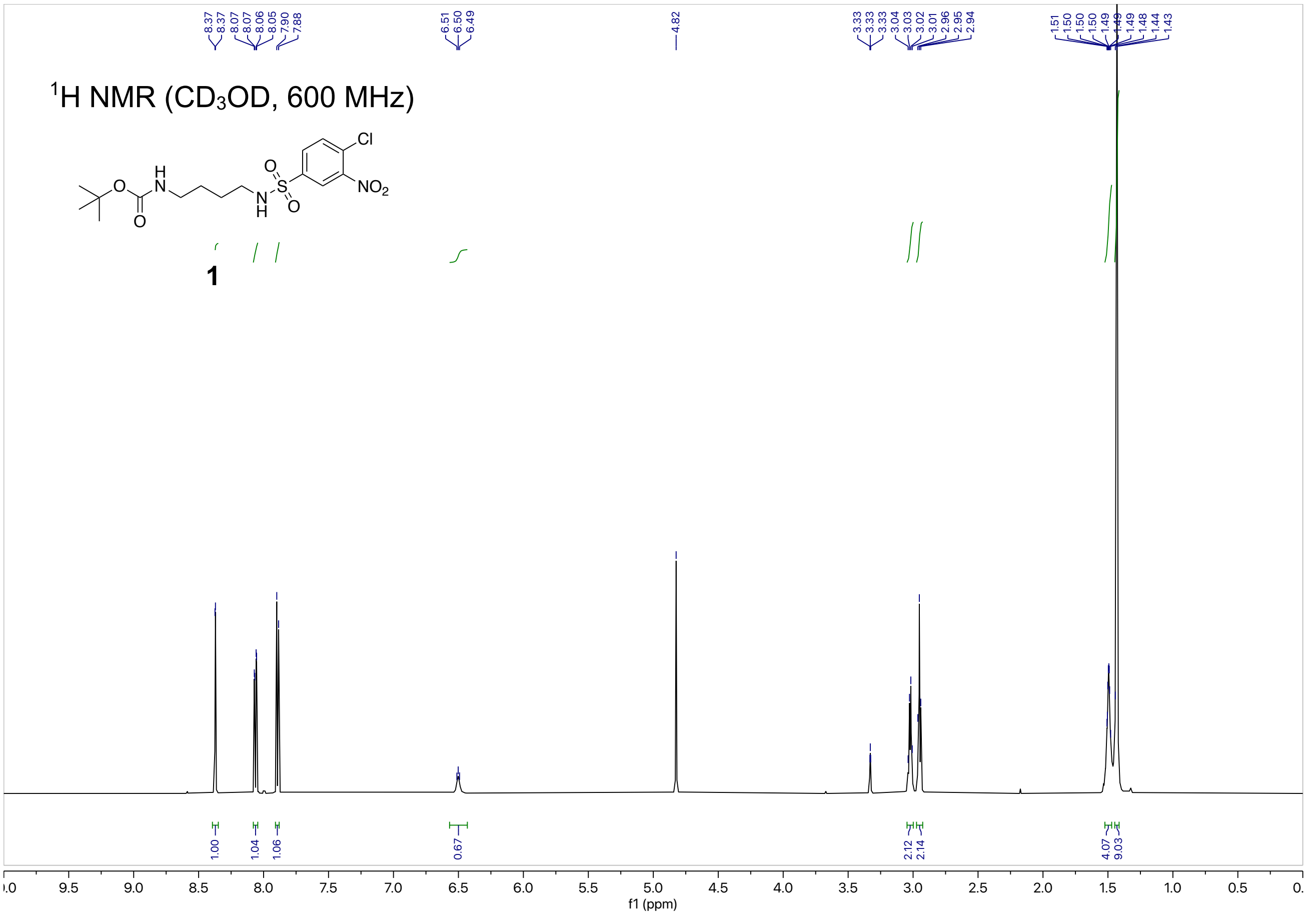

1H NMR (CD3OD, 600 MHz)
1
f1 (ppm)


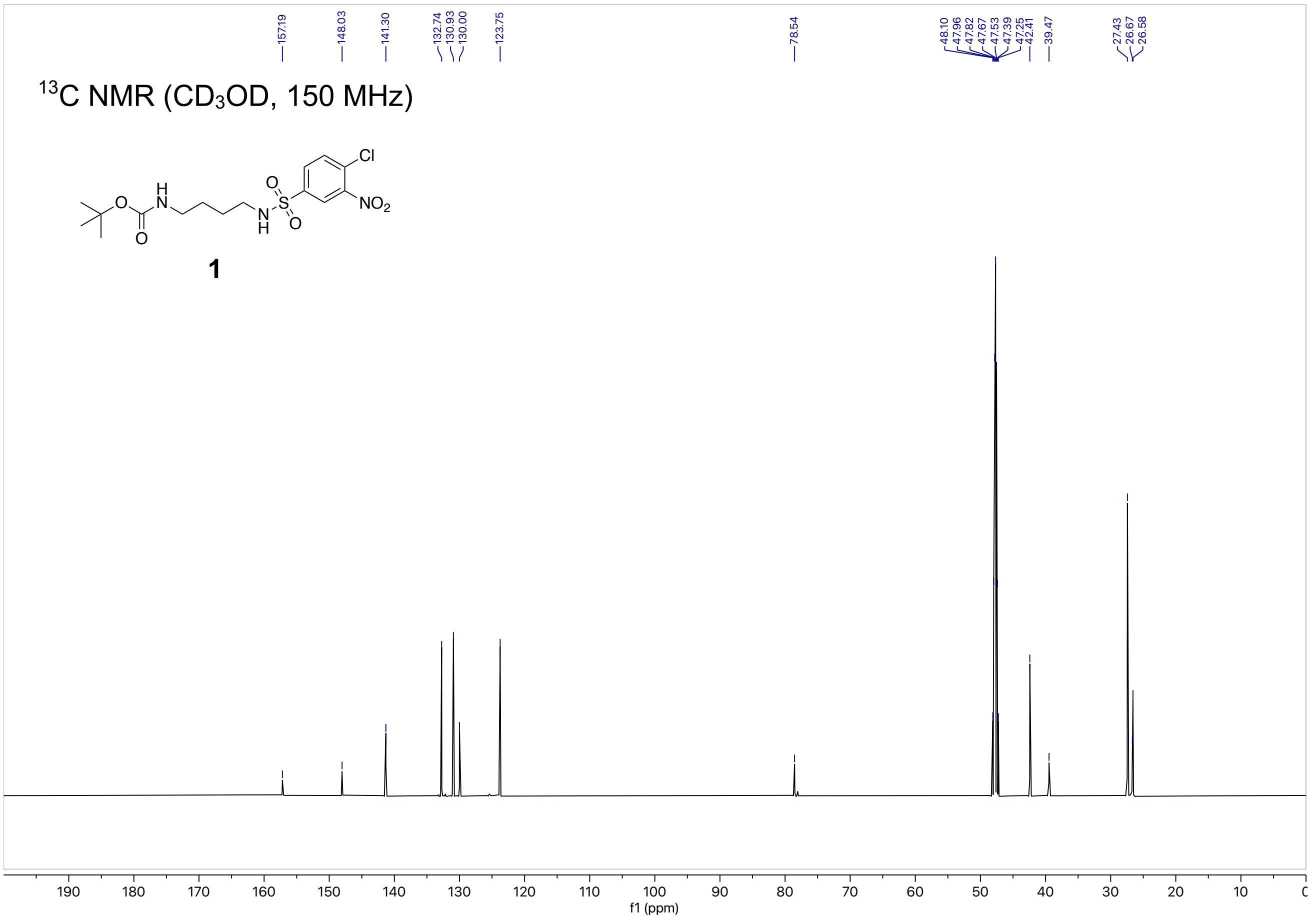

13C NMR (CD3OD, 150 MHz)
1
f1 (ppm)

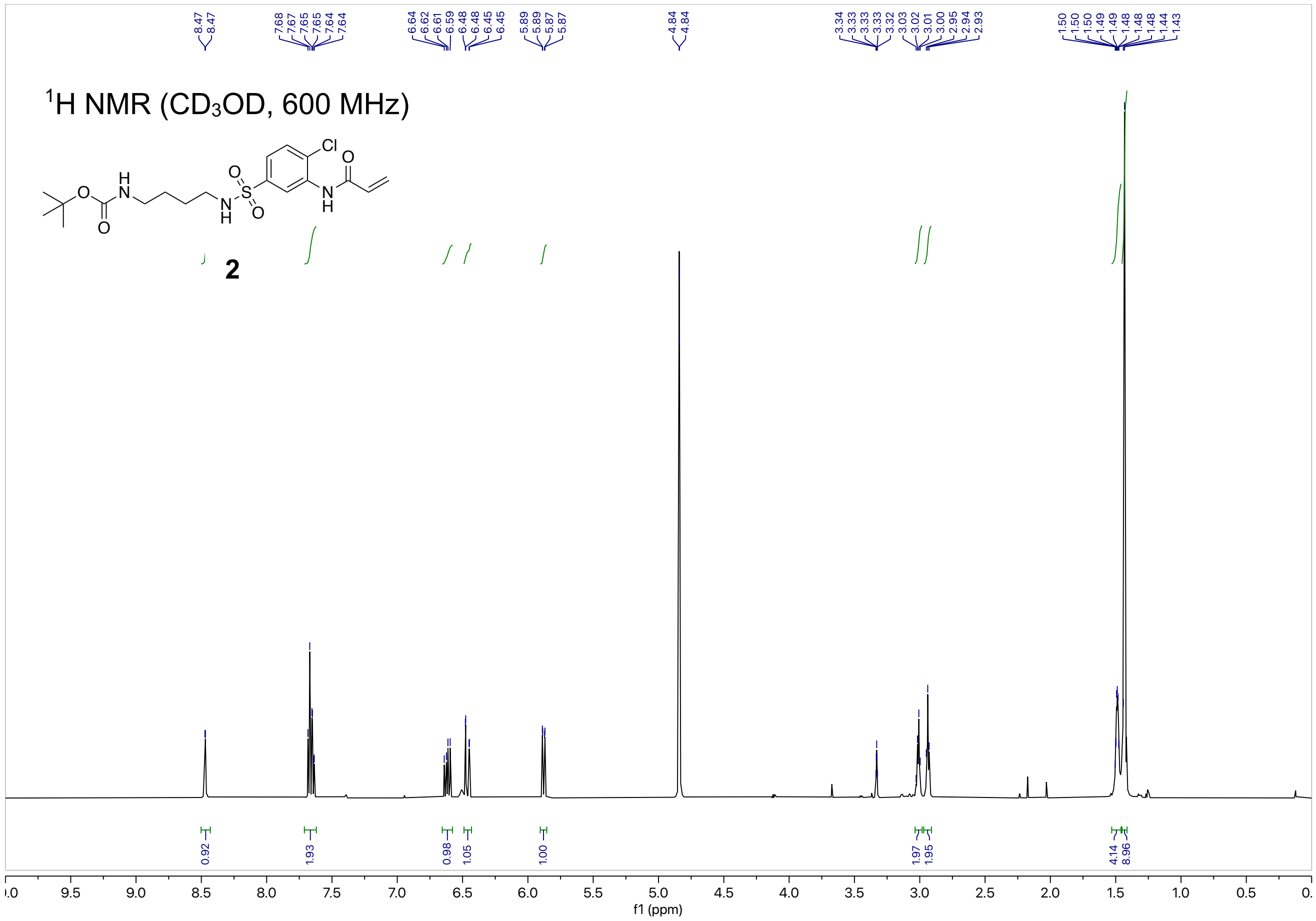
1H NMR (CD3OD, 600 MHz)
2
f1 (ppm)

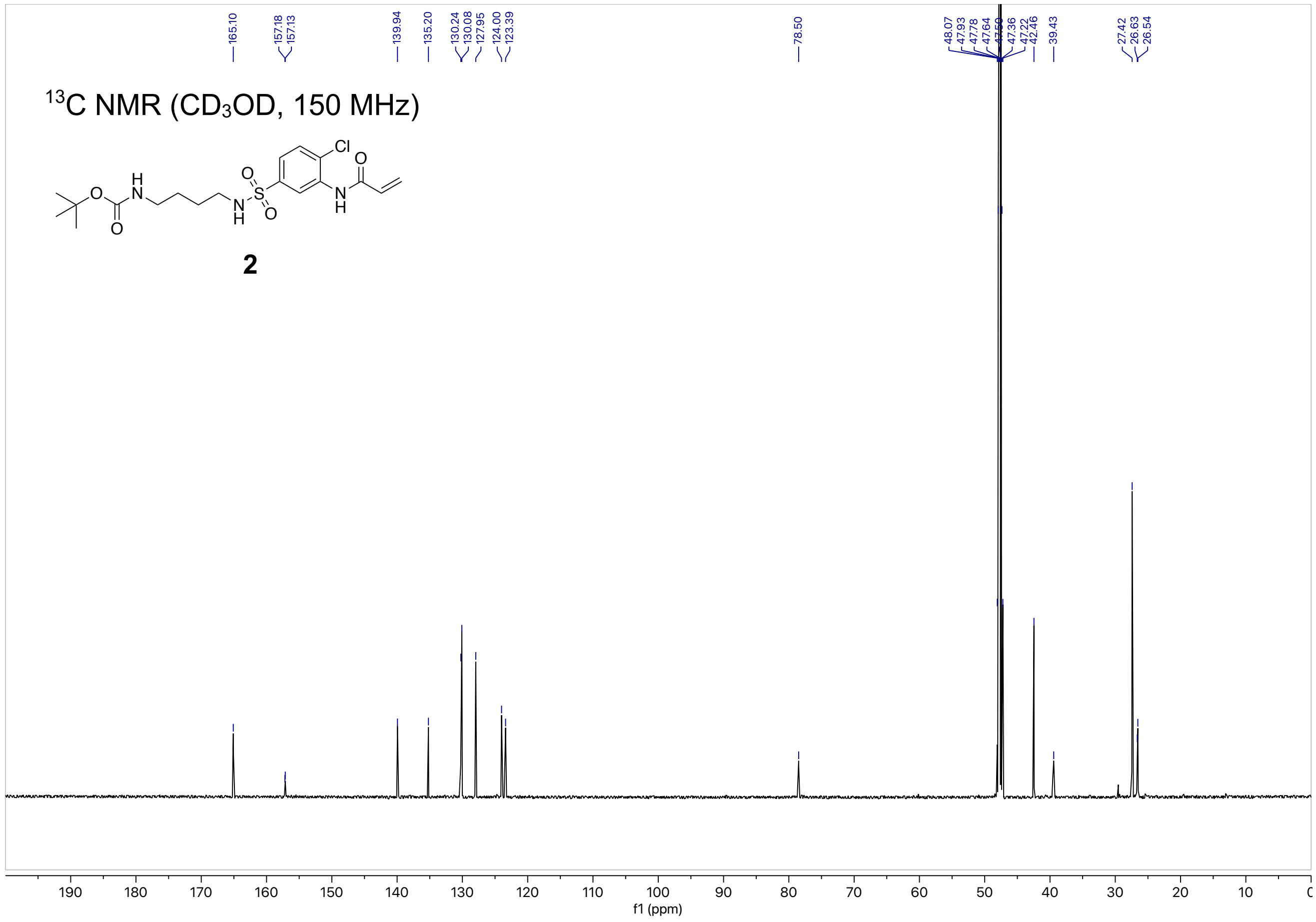
13C NMR (CD3OD, 150 MHz)
2
f1 (ppm)

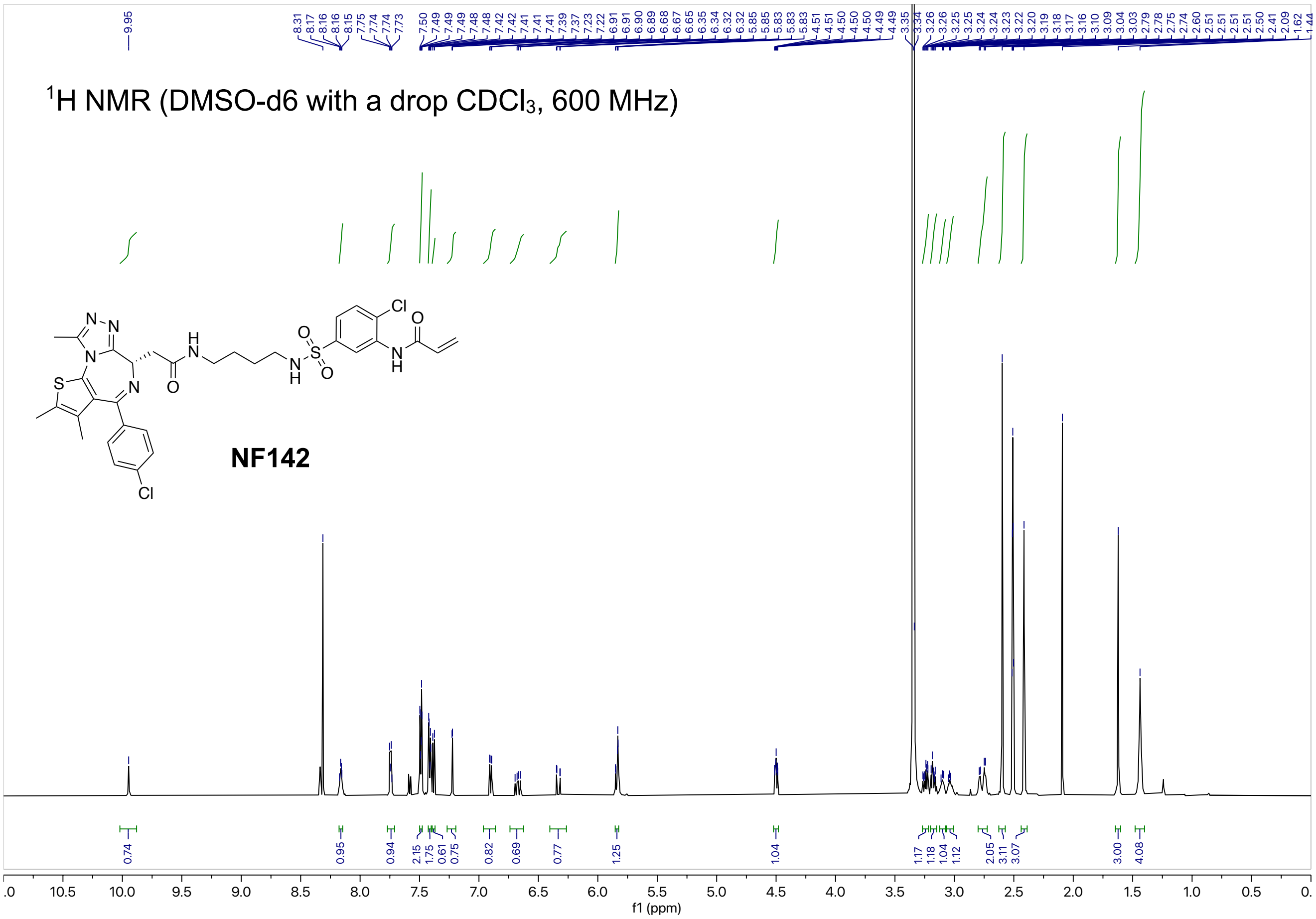

$^1$H NMR (DMSO-d6 with a drop $CDCl_3$, 600 MHz)
NF142


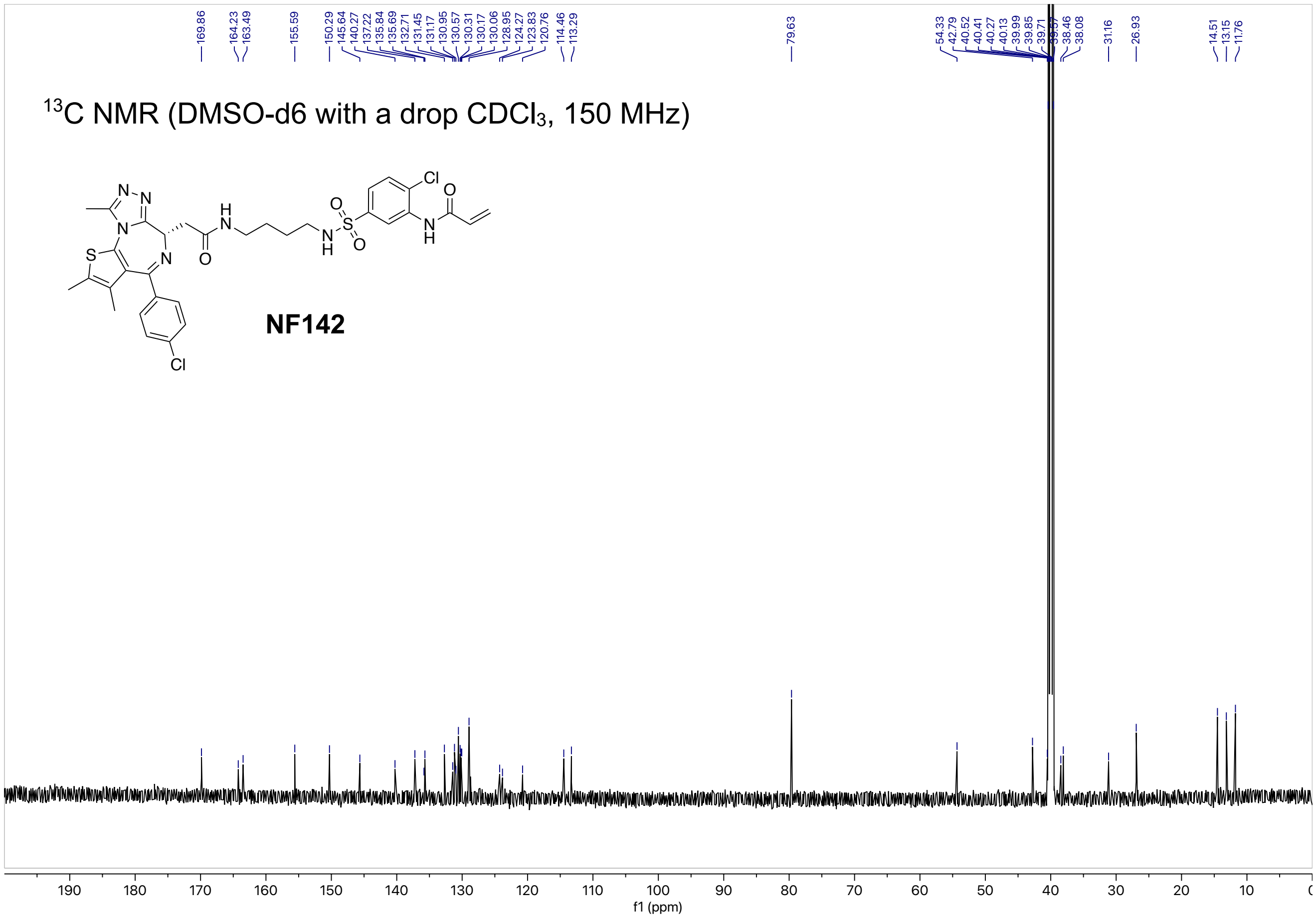

$^{13}$C NMR (DMSO-d6 with a drop $CDCl_3$, 150 MHz)
NF142

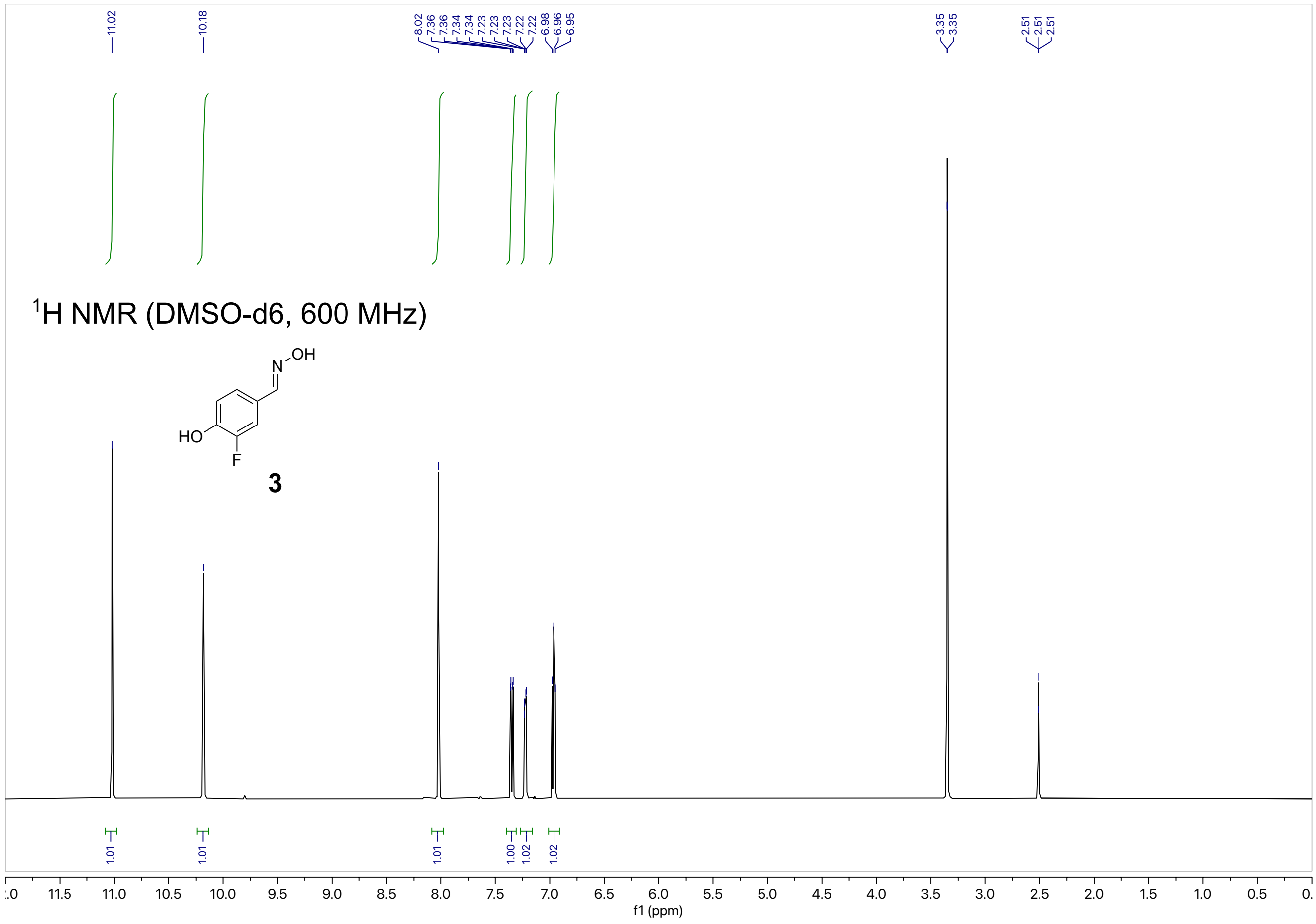

$^{1}$H NMR (DMSO-d6, 600 MHz)
3
f1 (ppm)


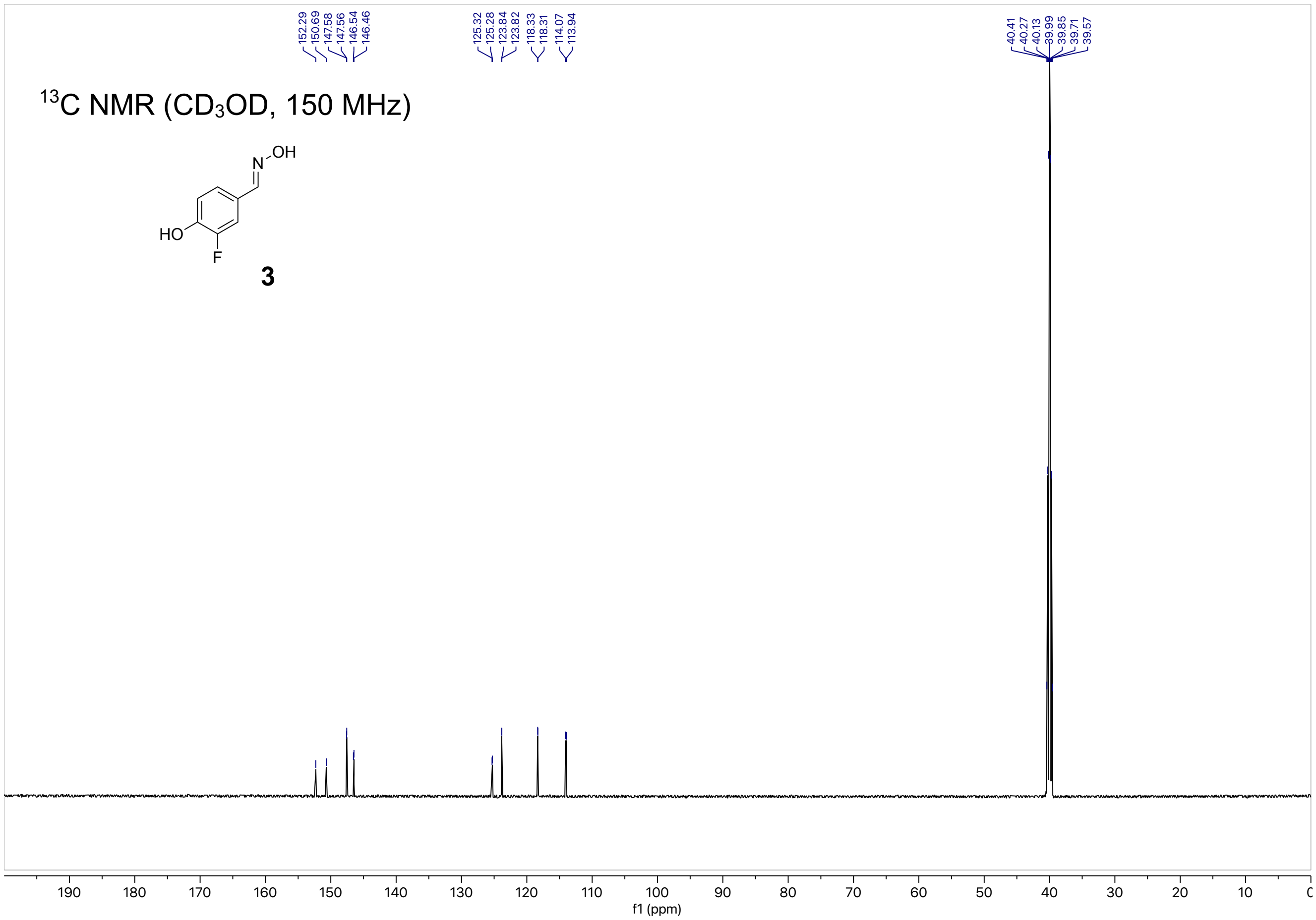

$^{13}$C NMR ($CD_3OD$, 150 MHz)
3
f1 (ppm)

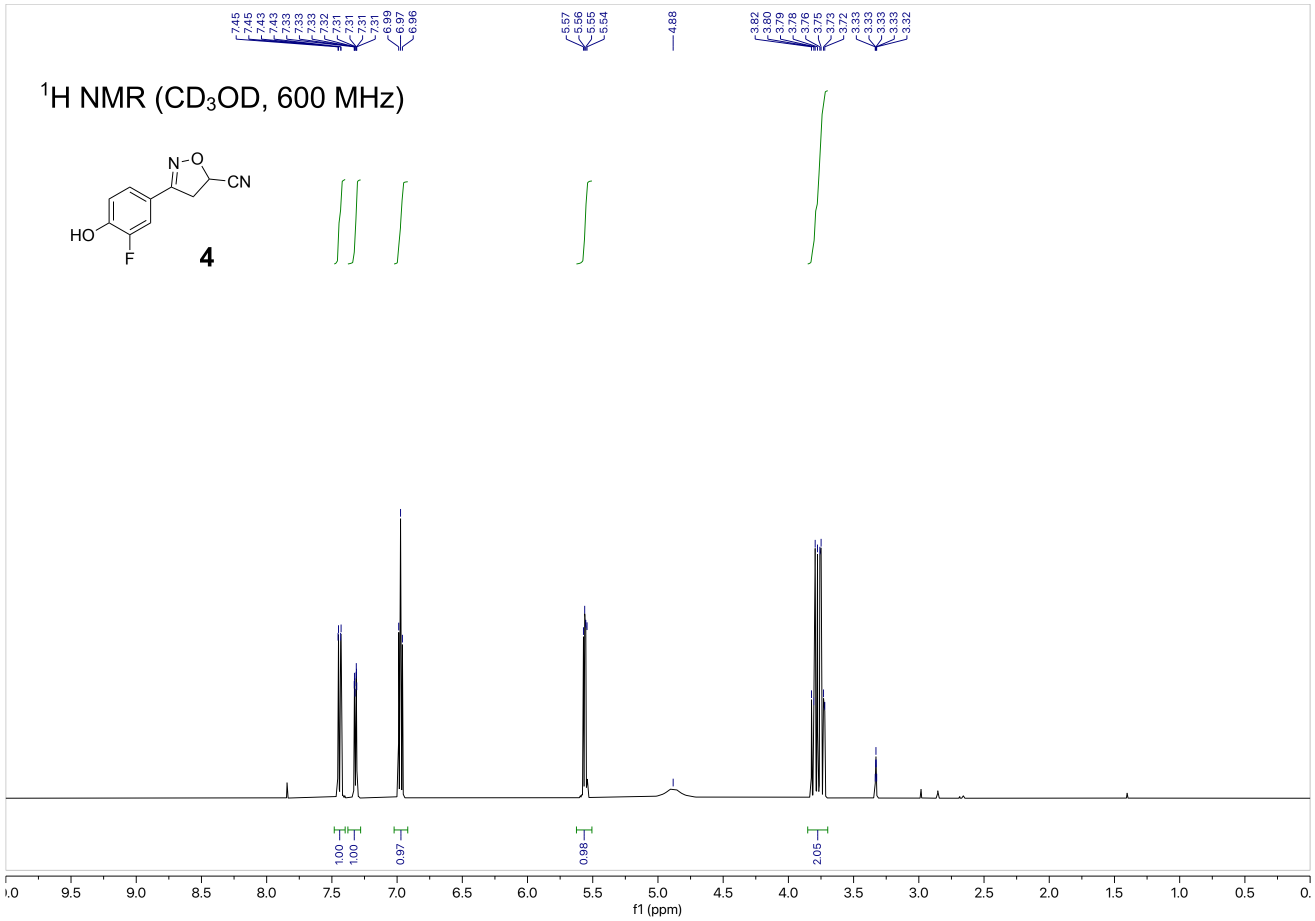
$^1$H NMR (CD$_3$OD, 600 MHz)
4
f1 (ppm)

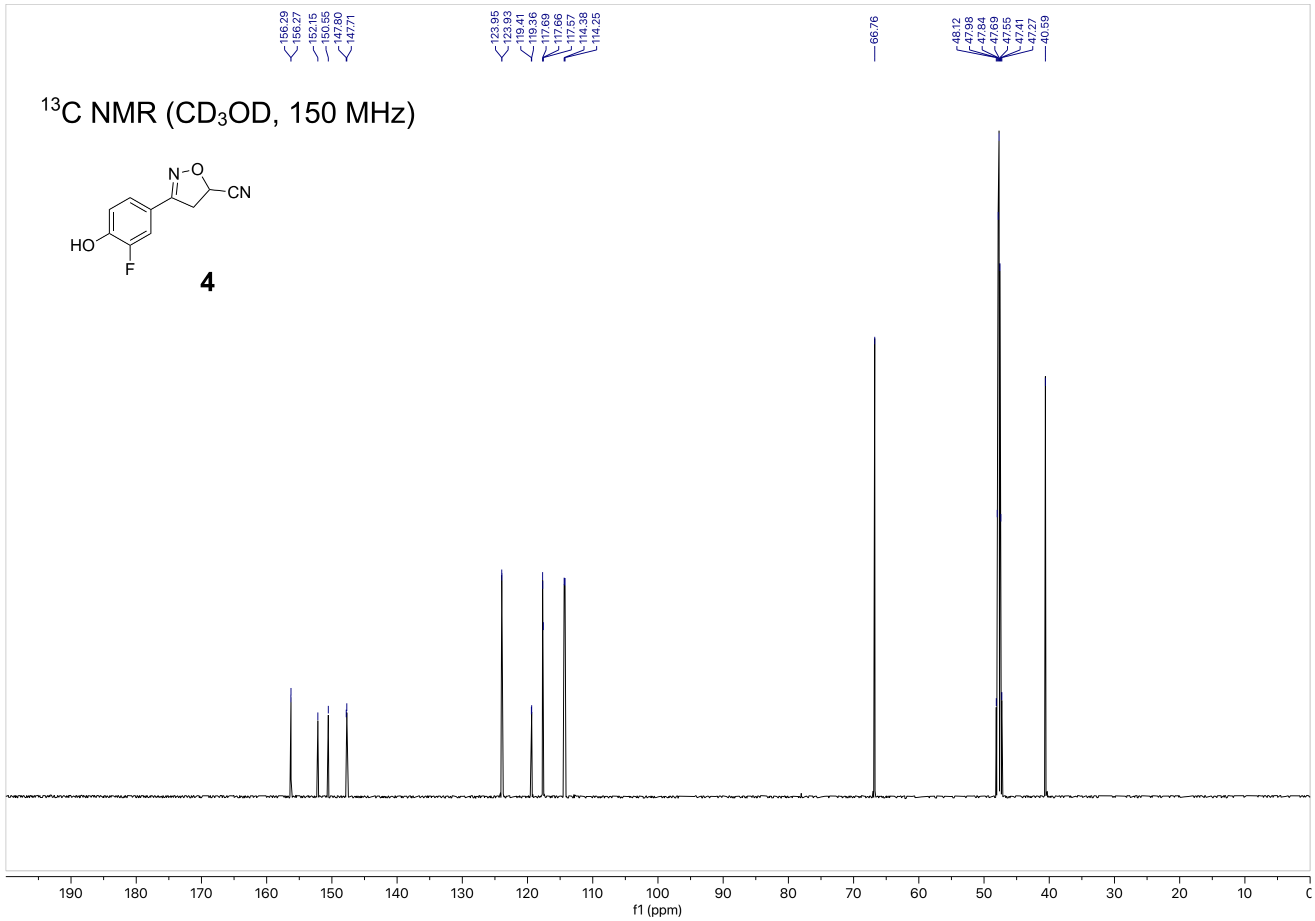
$^{13}$C NMR (CD$_3$OD, 150 MHz)
4
f1 (ppm)

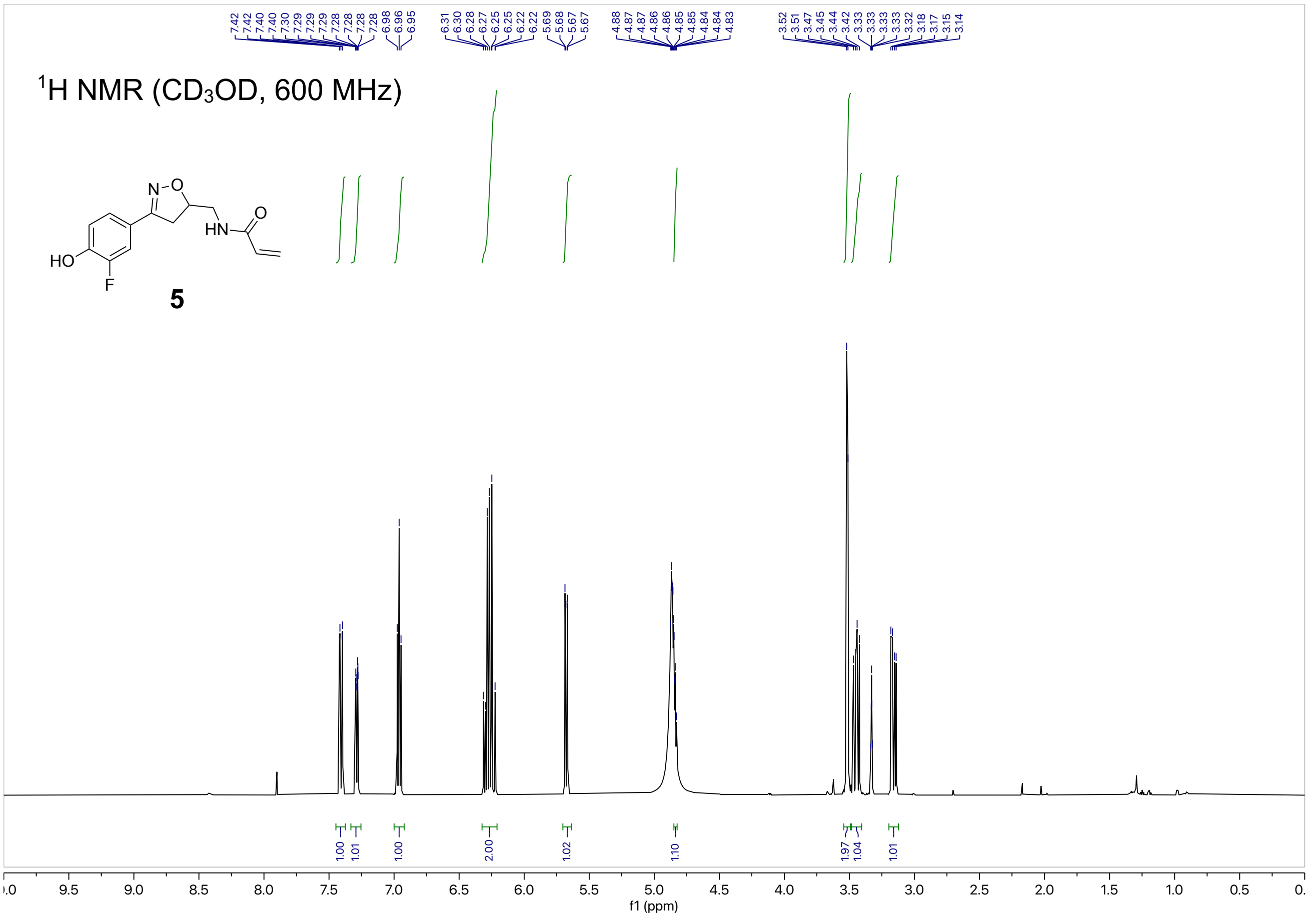
[1]H NMR (CD3OD, 600 MHz)
7.42 7.42 7.40 7.40 7.30 7.29 7.29 7.29 7.28 7.28 7.28 7.28 6.98 6.96 6.95
6.31 6.30 6.28 6.27 6.25 6.25 6.22 6.22 5.69 5.68 5.67 5.67
4.88 4.87 4.87 4.86 4.86 4.85 4.85 4.84 4.84 4.83
3.52 3.51 3.47 3.45 3.44 3.42 3.33 3.33 3.33 3.33 3.32 3.18 3.17 3.15 3.14
N-O
HN
O
HO
F
5
1.00 1.01 1.00 2.00 1.02 1.10 1.97 1.04 1.01
f1 (ppm)

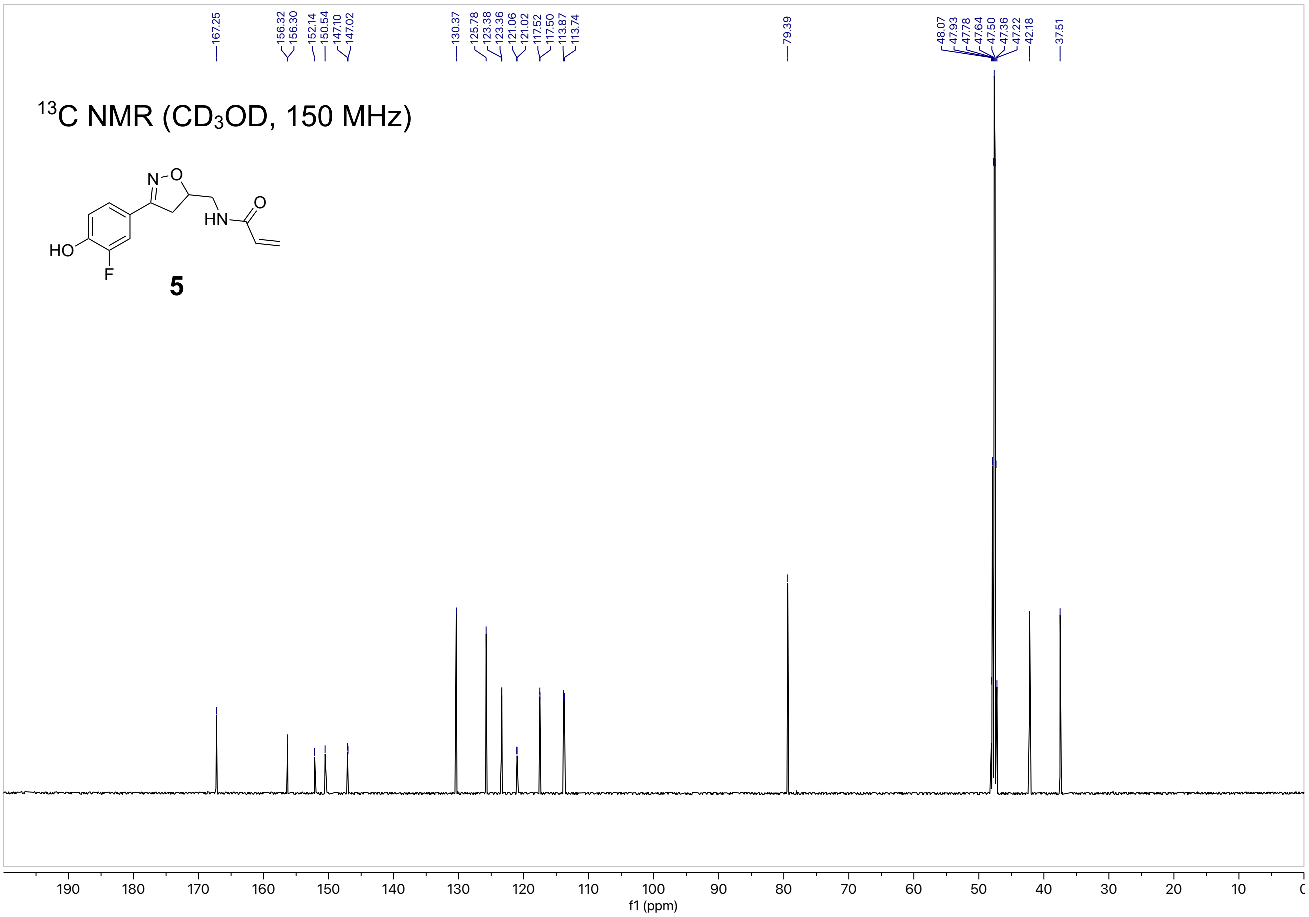
[13]C NMR (CD3OD, 150 MHz)
167.25
156.32 156.30 152.14 150.54 147.10 147.02
130.37 125.78 123.38 123.36 121.06 121.02 117.52 117.50 113.87 113.74
79.39
48.07 47.93 47.78 47.64 47.50 47.36 47.22 42.18
37.51
N-O
HN
O
HO
F
5
f1 (ppm)

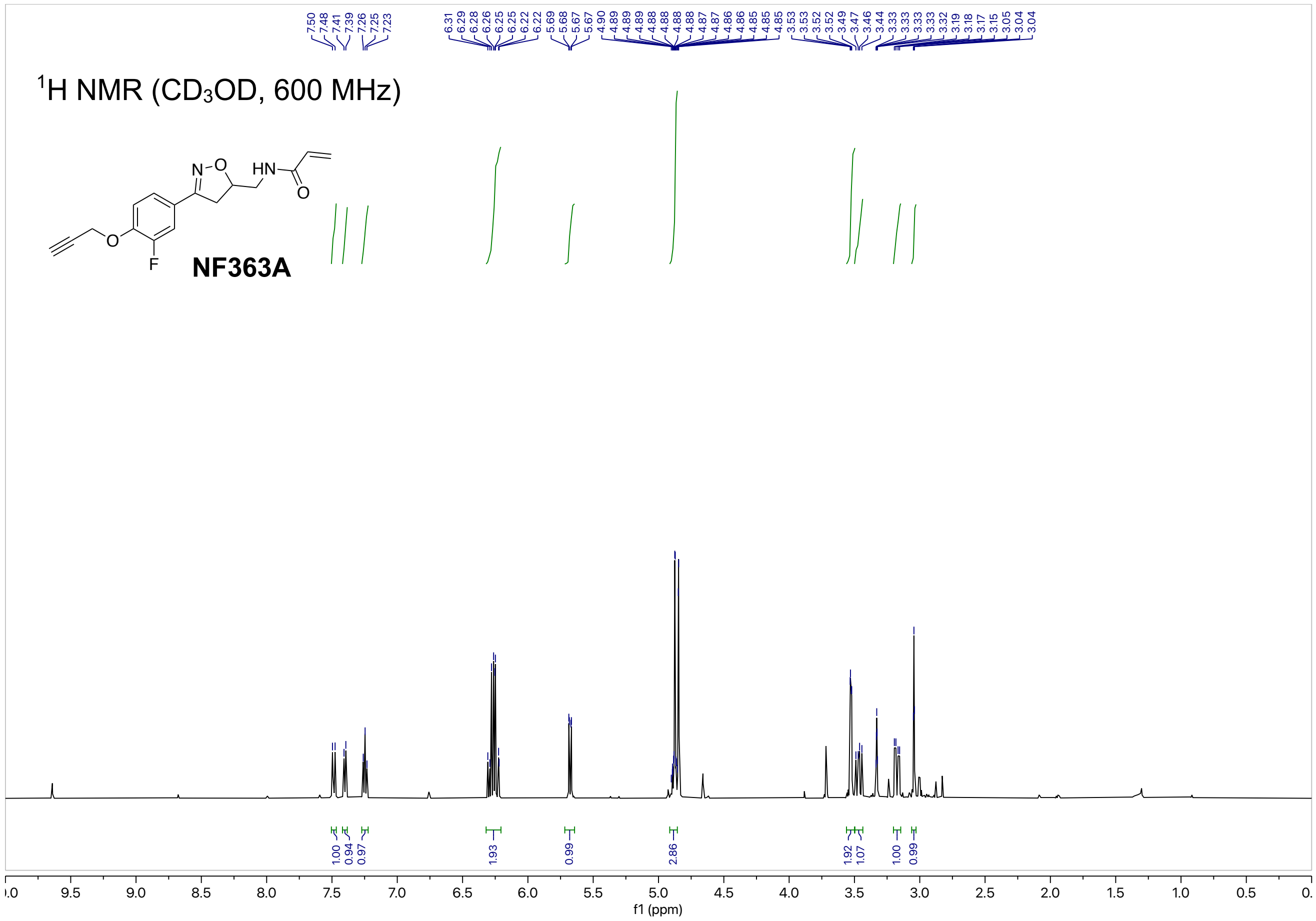
1H NMR (CD3OD, 600 MHz)
NF363A
f1 (ppm)

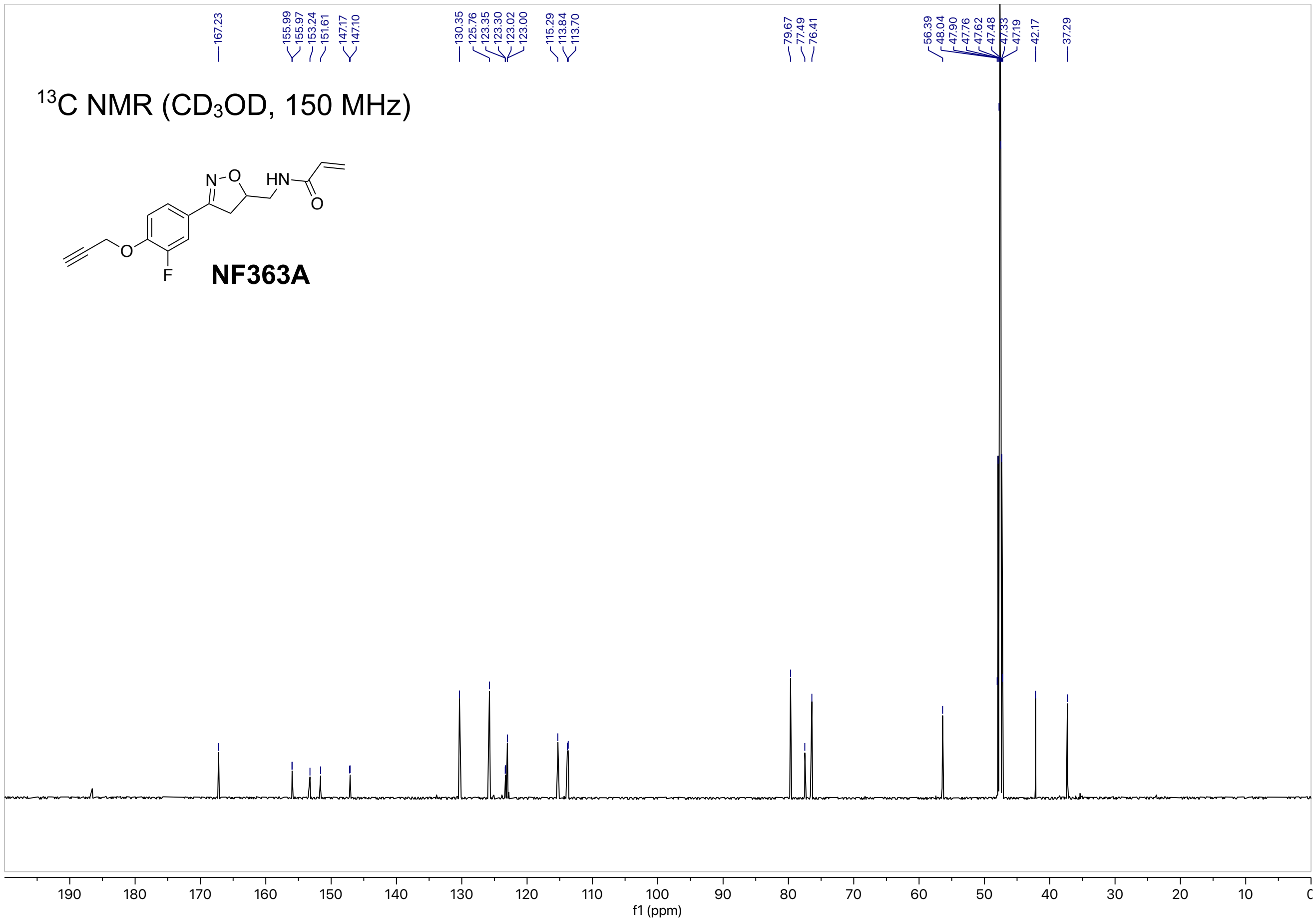
13C NMR (CD3OD, 150 MHz)
NF363A
f1 (ppm)

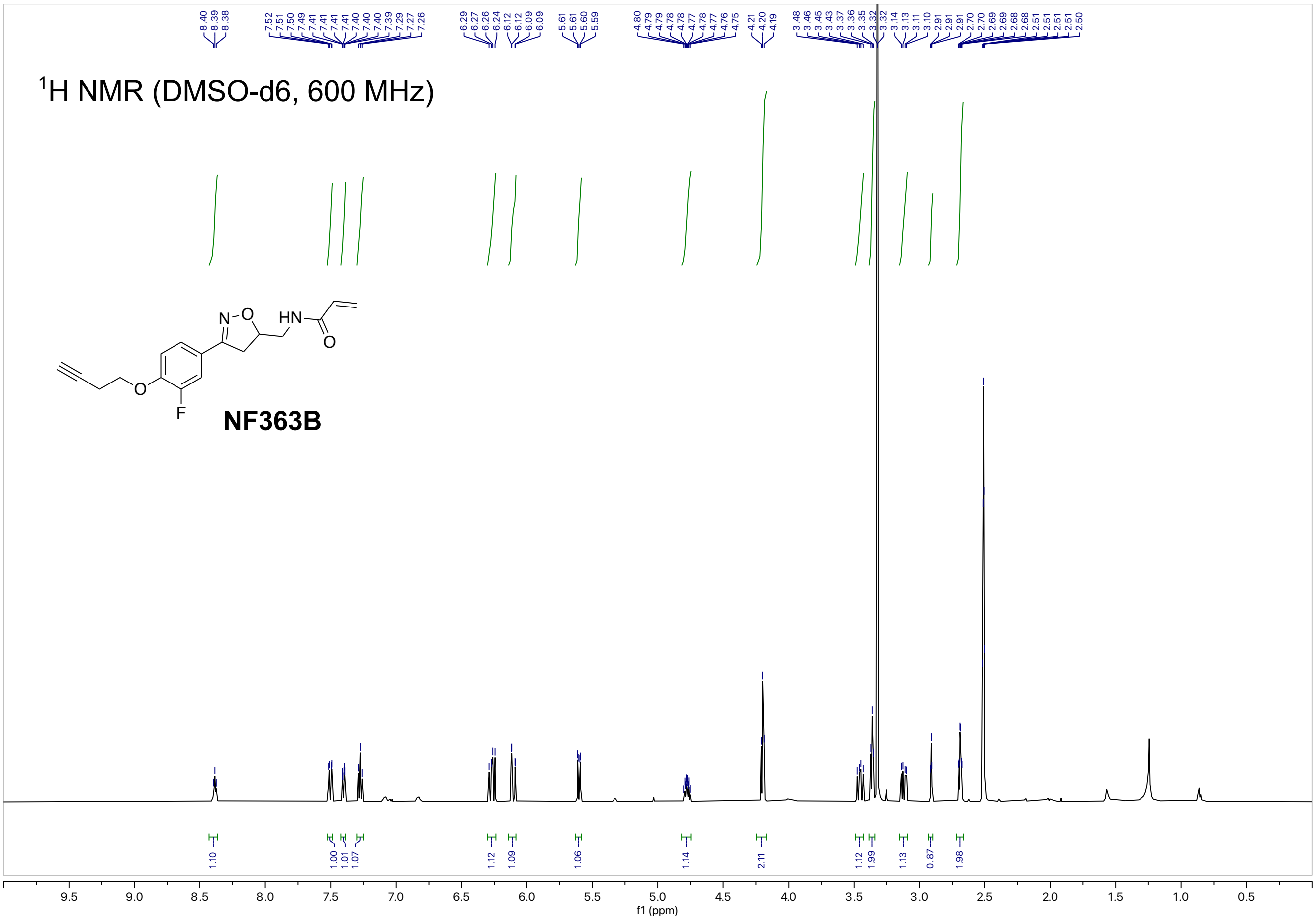

1H NMR (DMSO-d6, 600 MHz)
NF363B
f1 (ppm)


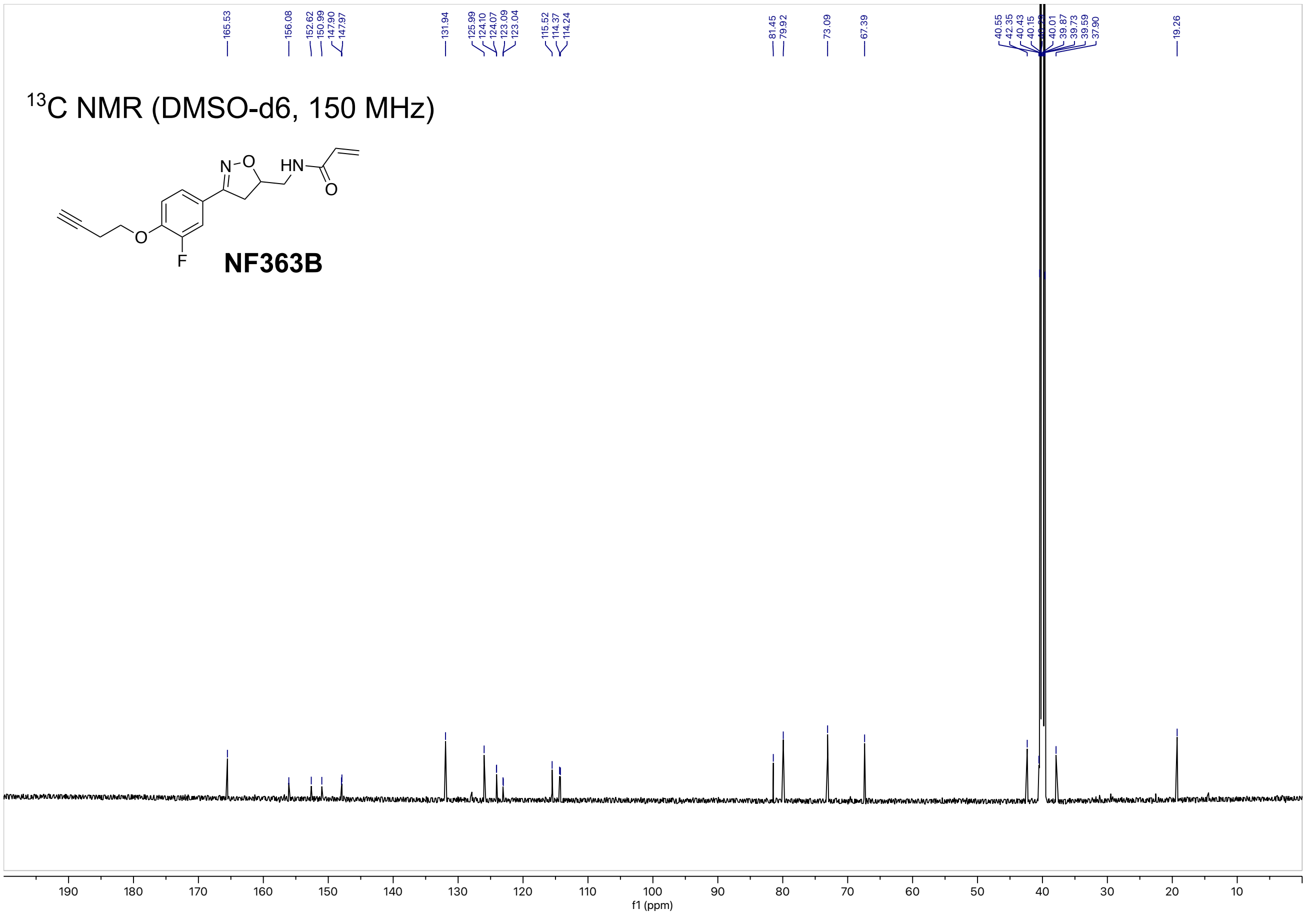

13C NMR (DMSO-d6, 150 MHz)
NF363B
f1 (ppm)

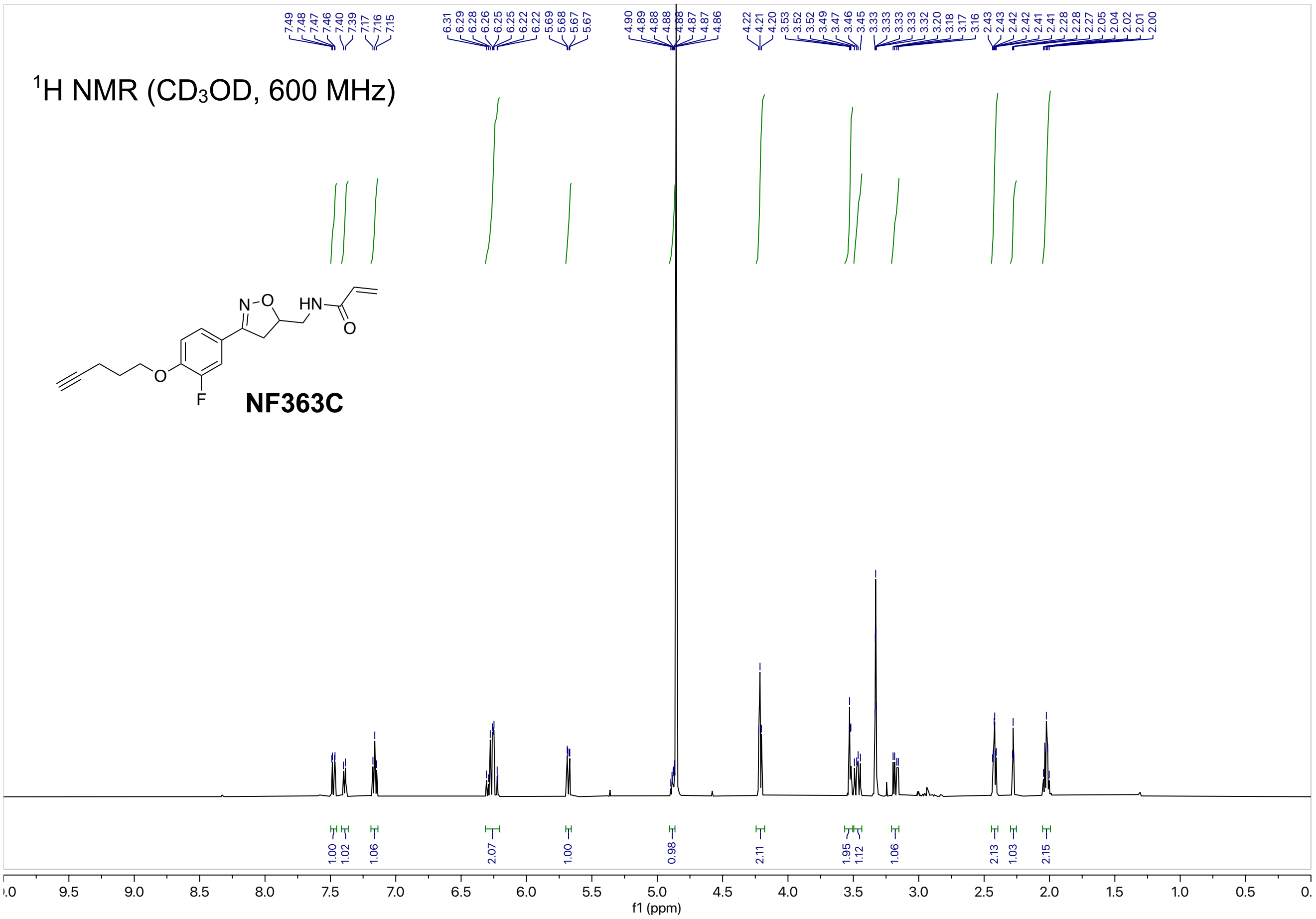

1H NMR (CD3OD, 600 MHz)
NF363C
7.49 7.48 7.47 7.46 7.40 7.39 7.17 7.16 7.15
6.31 6.29 6.28 6.26 6.25 6.25 6.22 6.22 5.69 5.68 5.67 5.67
4.90 4.89 4.88 4.88 4.88 4.87 4.87 4.86
4.22 4.21 4.20 3.53 3.52 3.52 3.49 3.47 3.46 3.45 3.33 3.33 3.33 3.33 3.32 3.20 3.18 3.17 3.16
2.43 2.43 2.42 2.42 2.41 2.41 2.28 2.28 2.27 2.05 2.04 2.02 2.01 2.00
1.00 1.02 1.06 2.07 1.00 0.98 2.11 1.95 1.12 1.06 2.13 1.03 2.15
f1 (ppm)


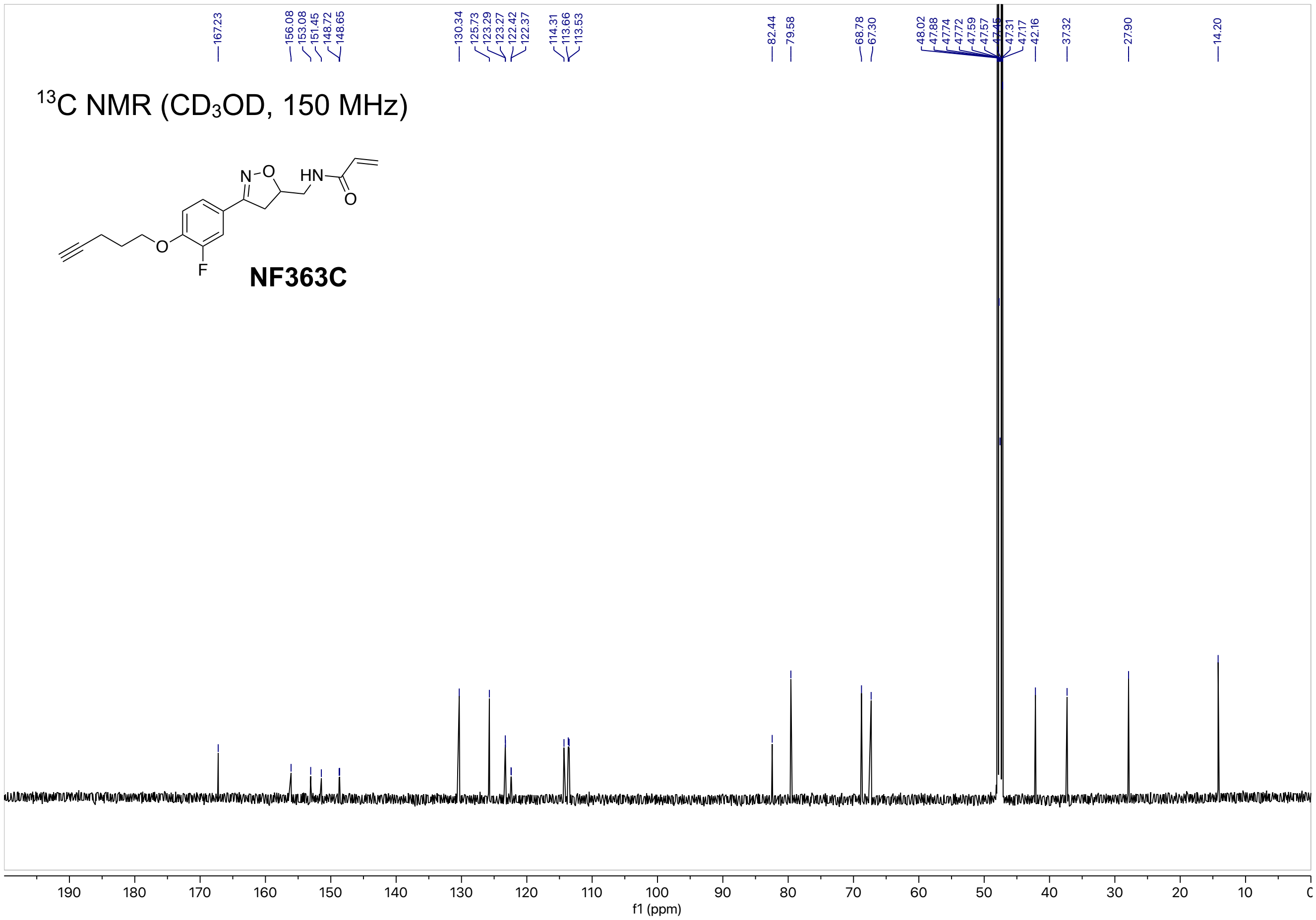

13C NMR (CD3OD, 150 MHz)
NF363C
167.23
156.08 153.08 151.45 148.72 148.65
130.34 125.73 123.29 123.27 122.42 122.37
114.31 113.66 113.53
82.44 79.58
68.78 67.30
48.02 47.88 47.74 47.72 47.59 47.57 47.45 47.31 47.17 42.16
37.32
27.90
14.20
f1 (ppm)

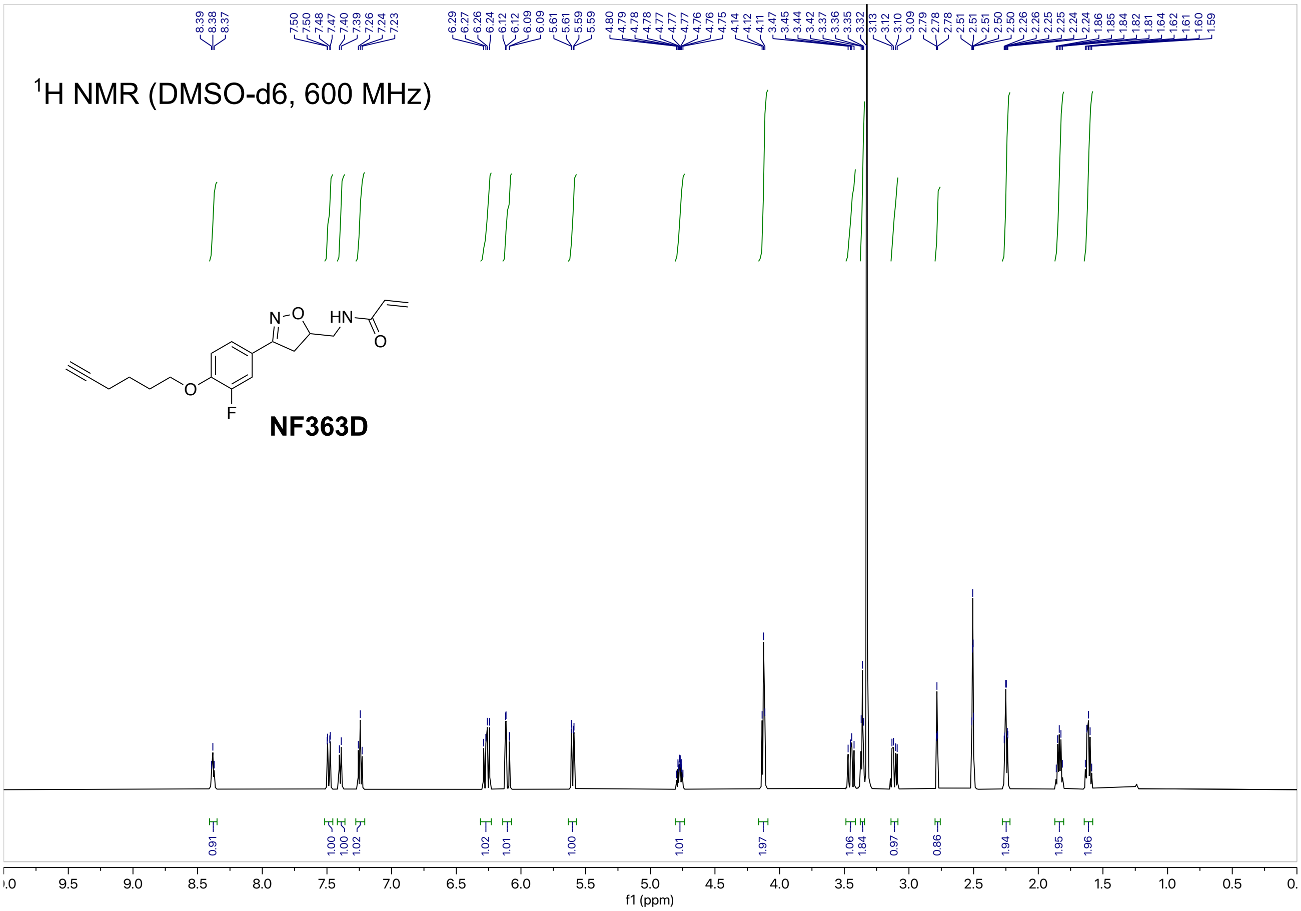
$^1$H NMR (DMSO-d6, 600 MHz)
NF363D
f1 (ppm)

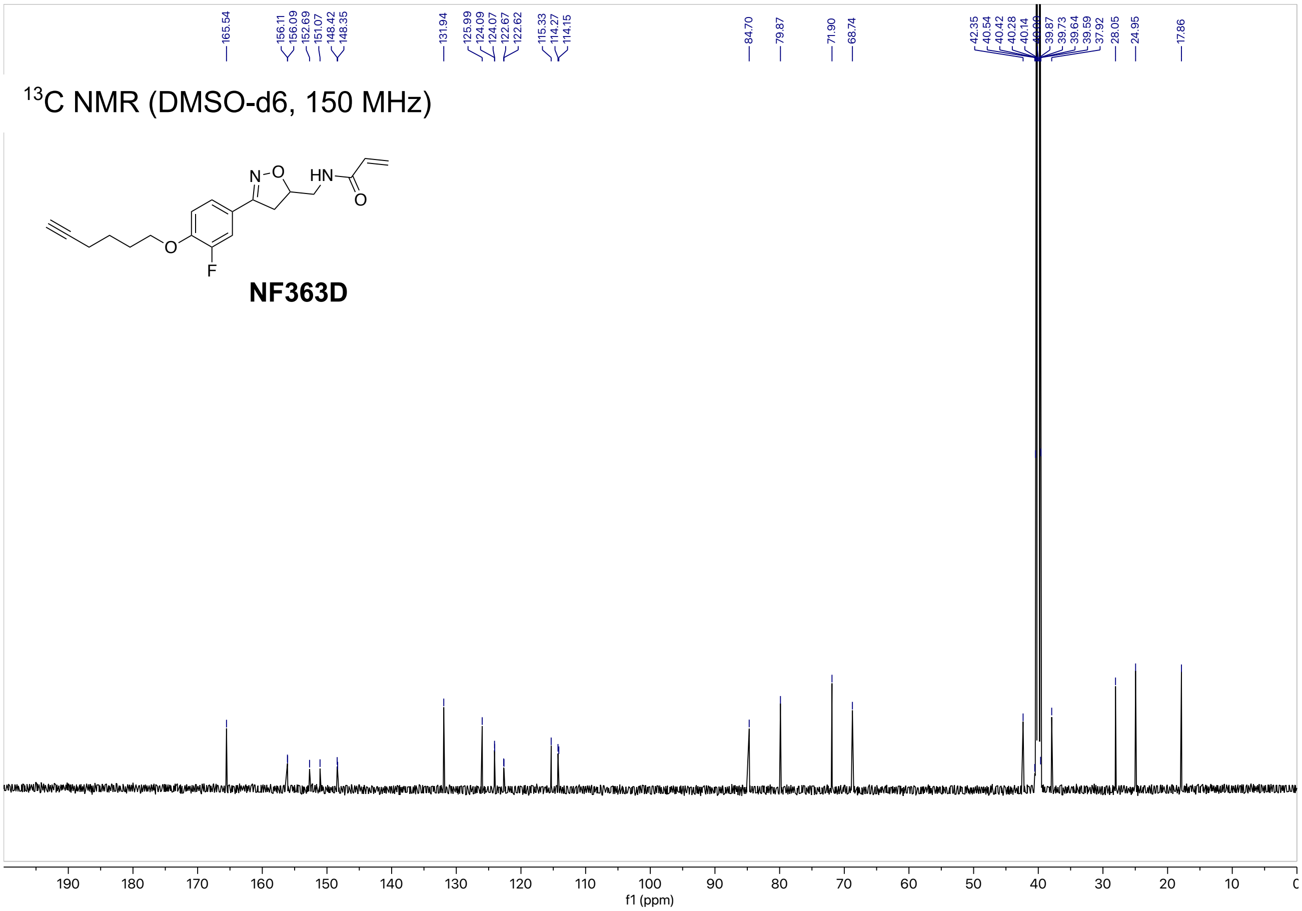
$^{13}$C NMR (DMSO-d6, 150 MHz)
NF363D
f1 (ppm)

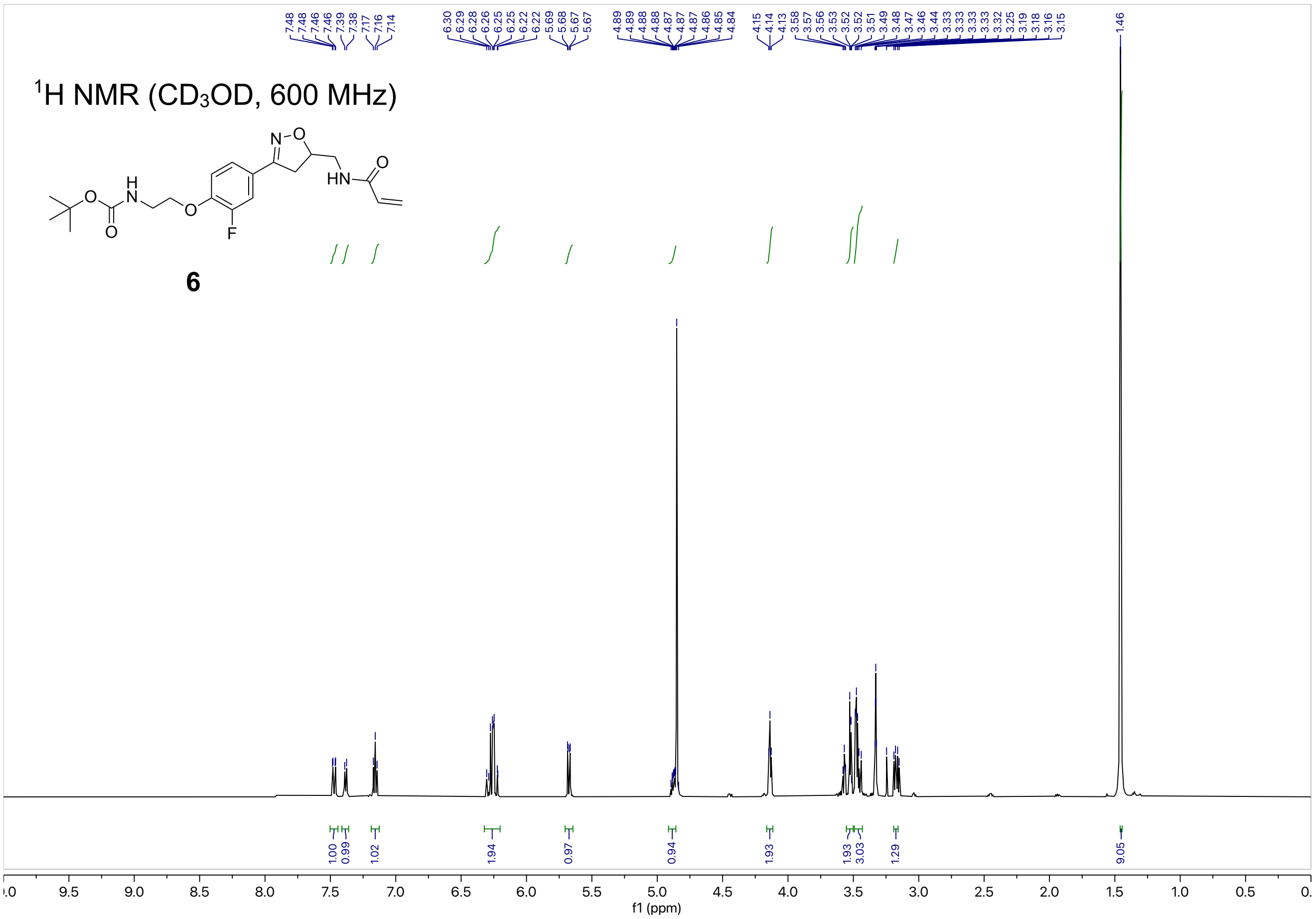
$^{1}$H NMR ($CD_3OD$, 600 MHz)
6
f1 (ppm)

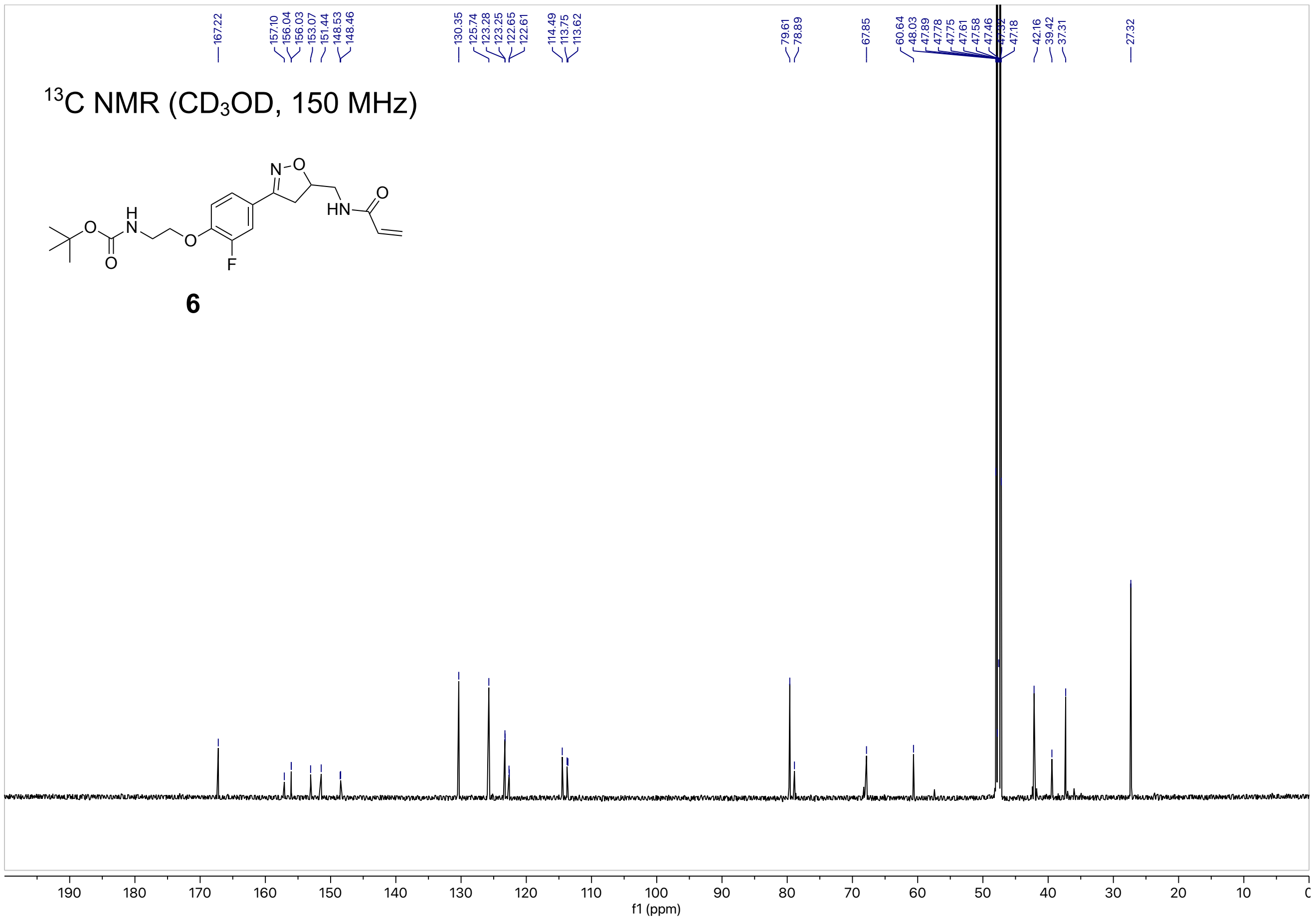
$^{13}$C NMR ($CD_3OD$, 150 MHz)
6
f1 (ppm)

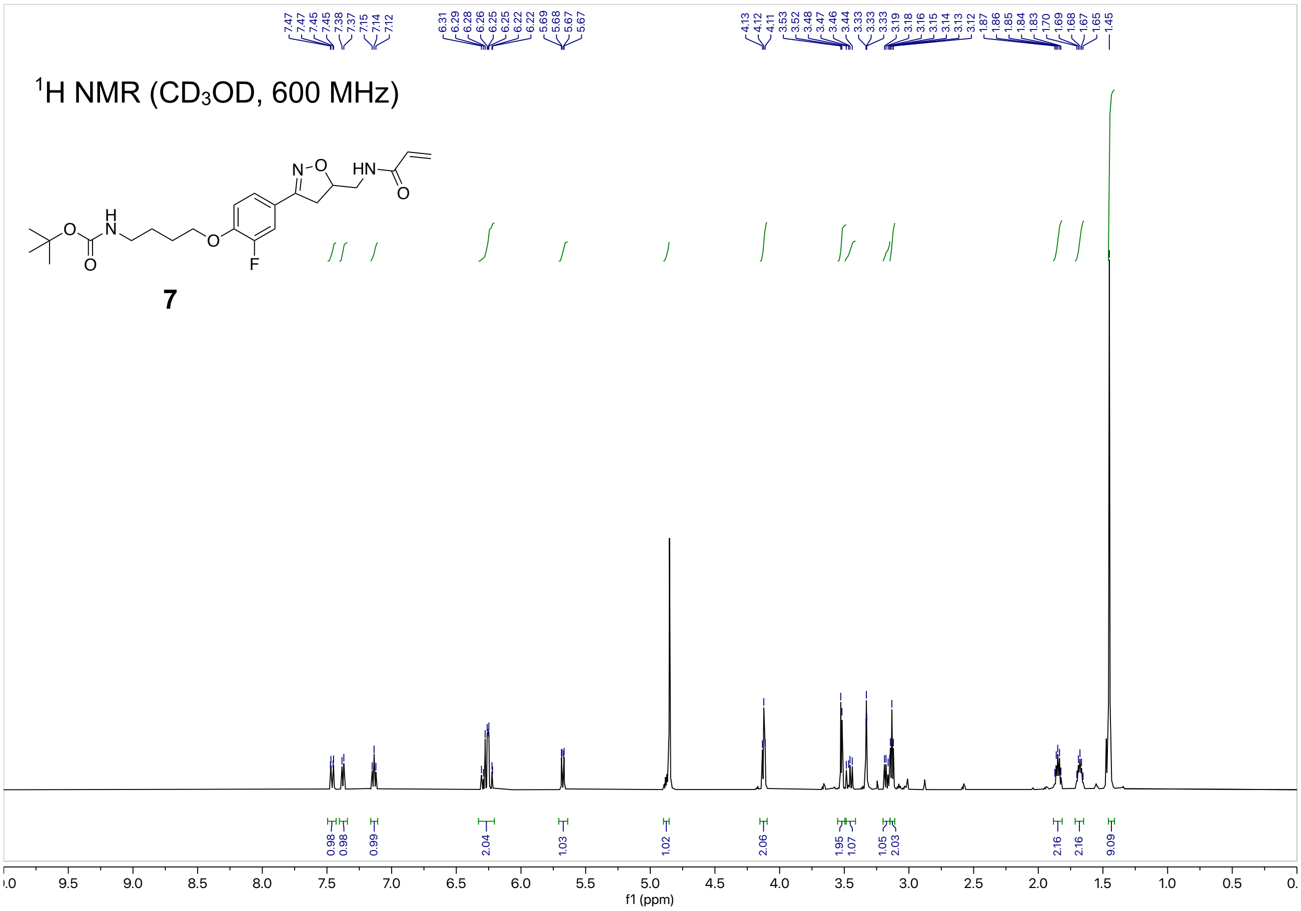

7.47
7.47
7.45
7.45
7.38
7.37
7.15
7.14
7.12
6.31
6.29
6.28
6.26
6.25
6.25
6.22
6.22
5.69
5.68
5.67
5.67
4.13
4.12
4.11
3.53
3.52
3.48
3.47
3.46
3.44
3.33
3.33
3.33
3.19
3.18
3.16
3.15
3.14
3.13
3.12
1.87
1.86
1.85
1.84
1.83
1.70
1.69
1.68
1.67
1.65
1.45
1H NMR (CD3OD, 600 MHz)
N-O
HN
O
H
N
O
O
O
F
7
0.98
0.98
0.99
2.04
1.03
1.02
2.06
1.95
1.07
1.05
2.03
2.16
2.16
9.09
f1 (ppm)


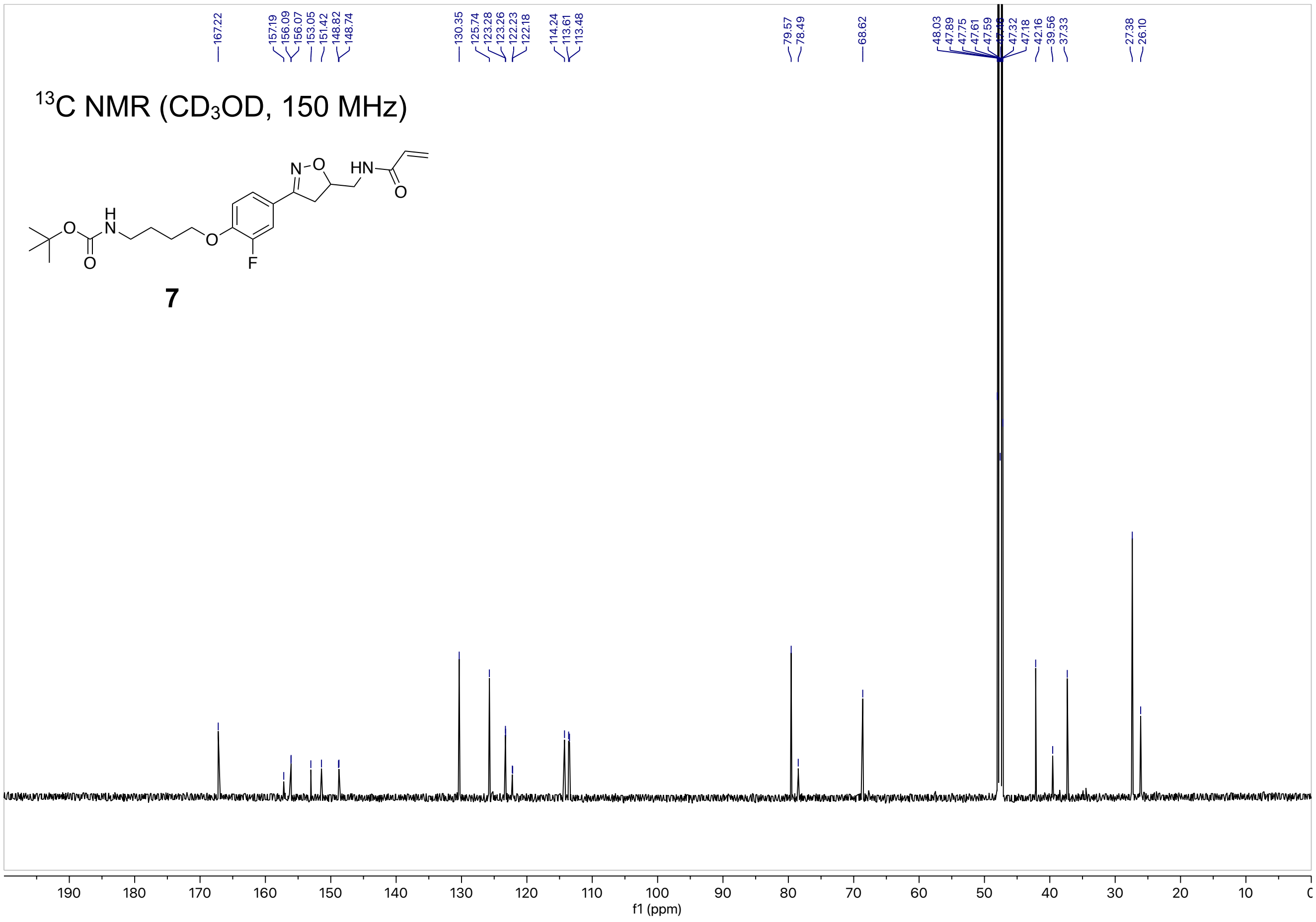

167.22
157.19
156.09
156.07
153.05
151.42
148.82
148.74
130.35
125.74
123.28
123.26
122.23
122.18
114.24
113.61
113.48
79.57
78.49
68.62
48.03
47.89
47.75
47.61
47.59
47.32
47.18
42.16
39.56
37.33
27.38
26.10
13C NMR (CD3OD, 150 MHz)
N-O
HN
O
H
N
O
O
O
F
7
f1 (ppm)

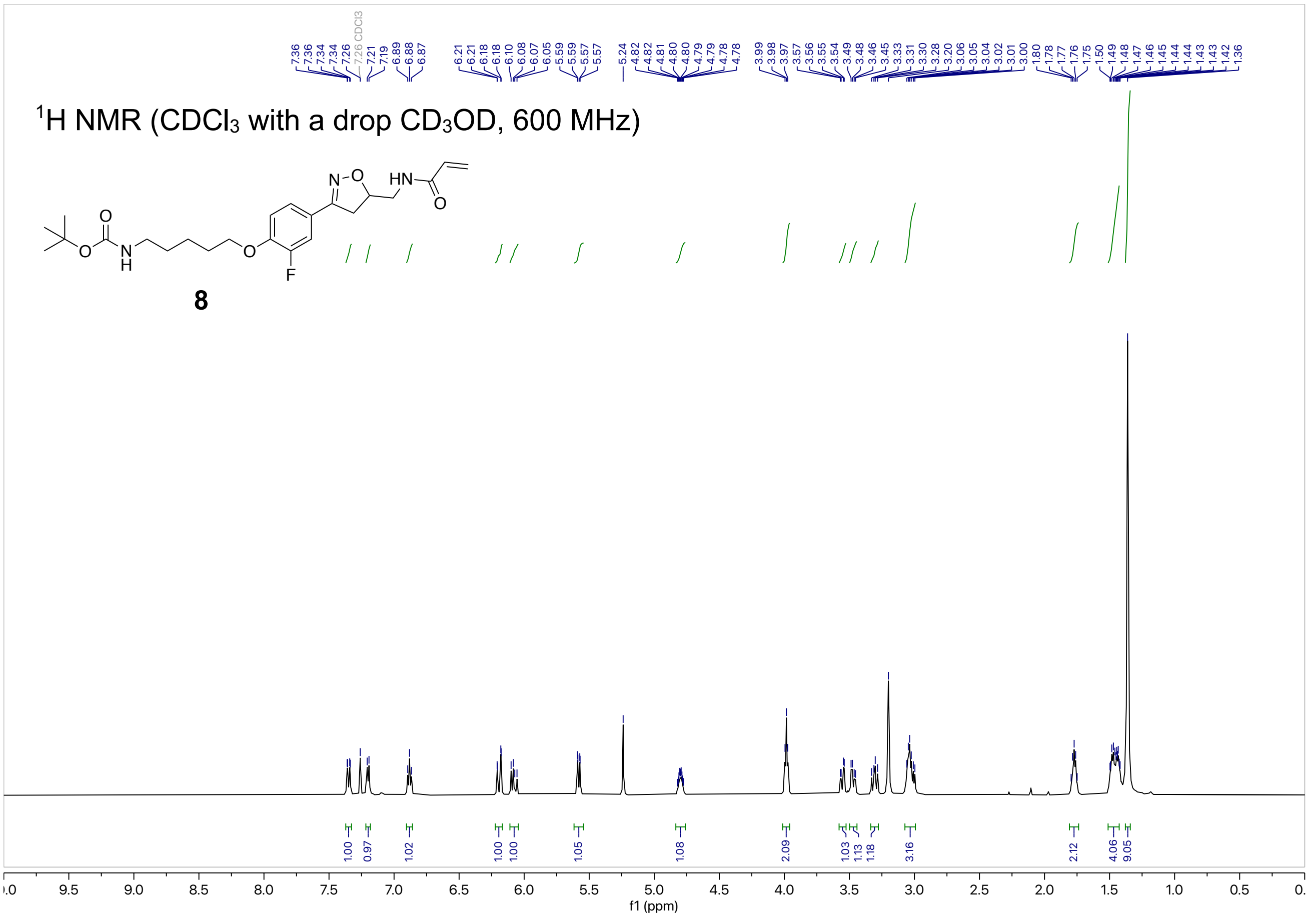

$^1$H NMR (CDCl$_3$ with a drop CD$_3$OD, 600 MHz)
8
f1 (ppm)


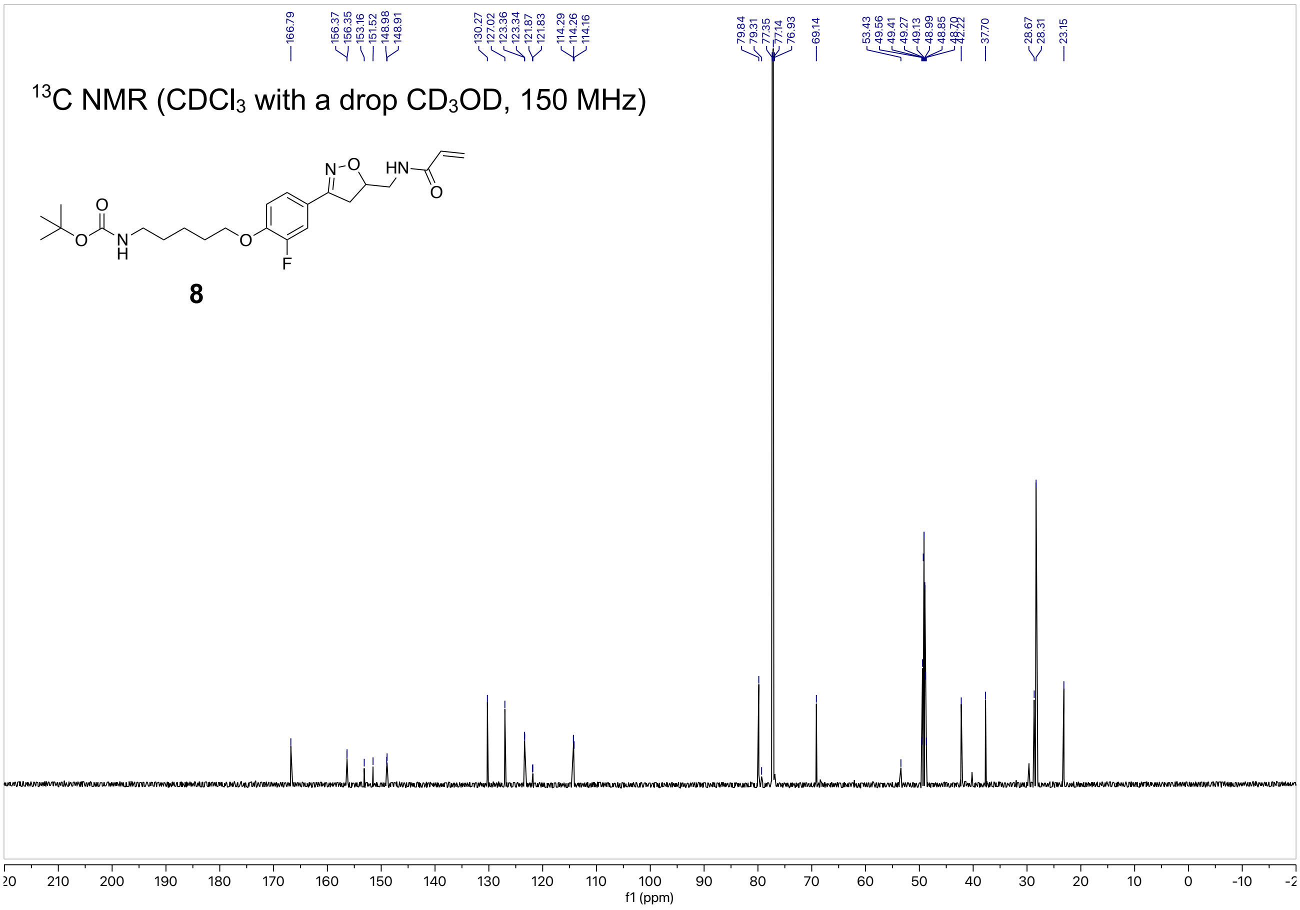

$^{13}$C NMR (CDCl$_3$ with a drop CD$_3$OD, 150 MHz)
8
f1 (ppm)

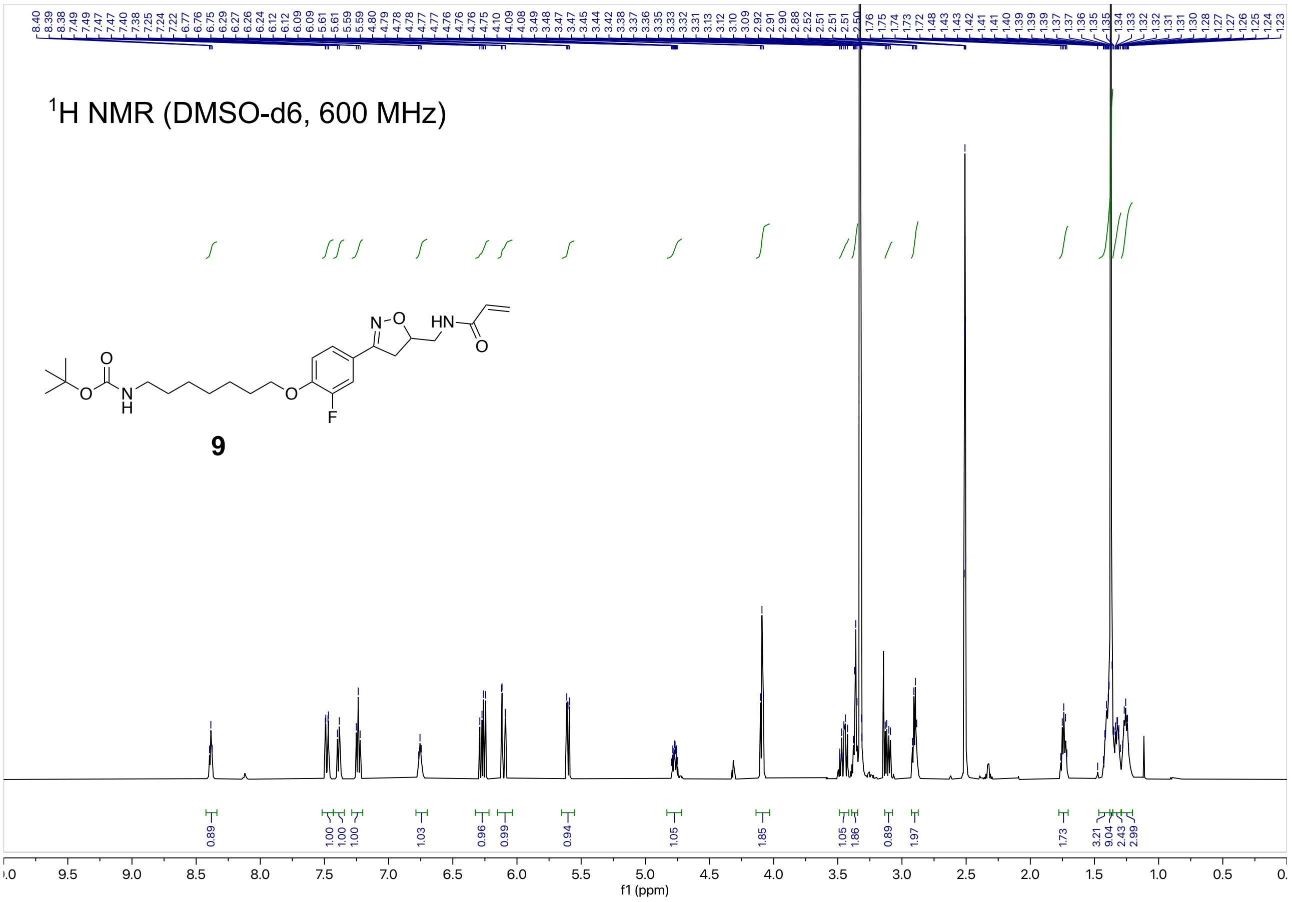

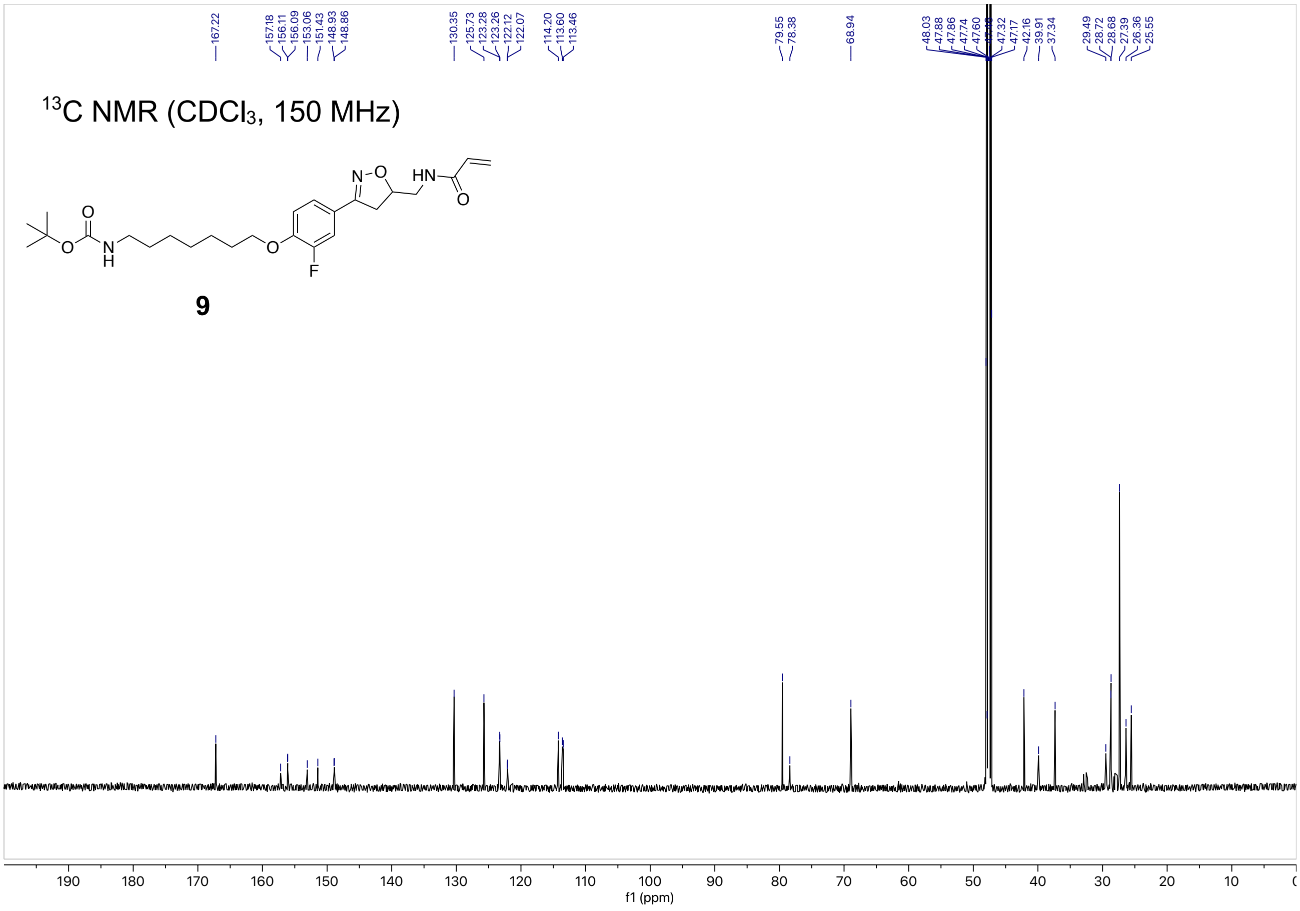

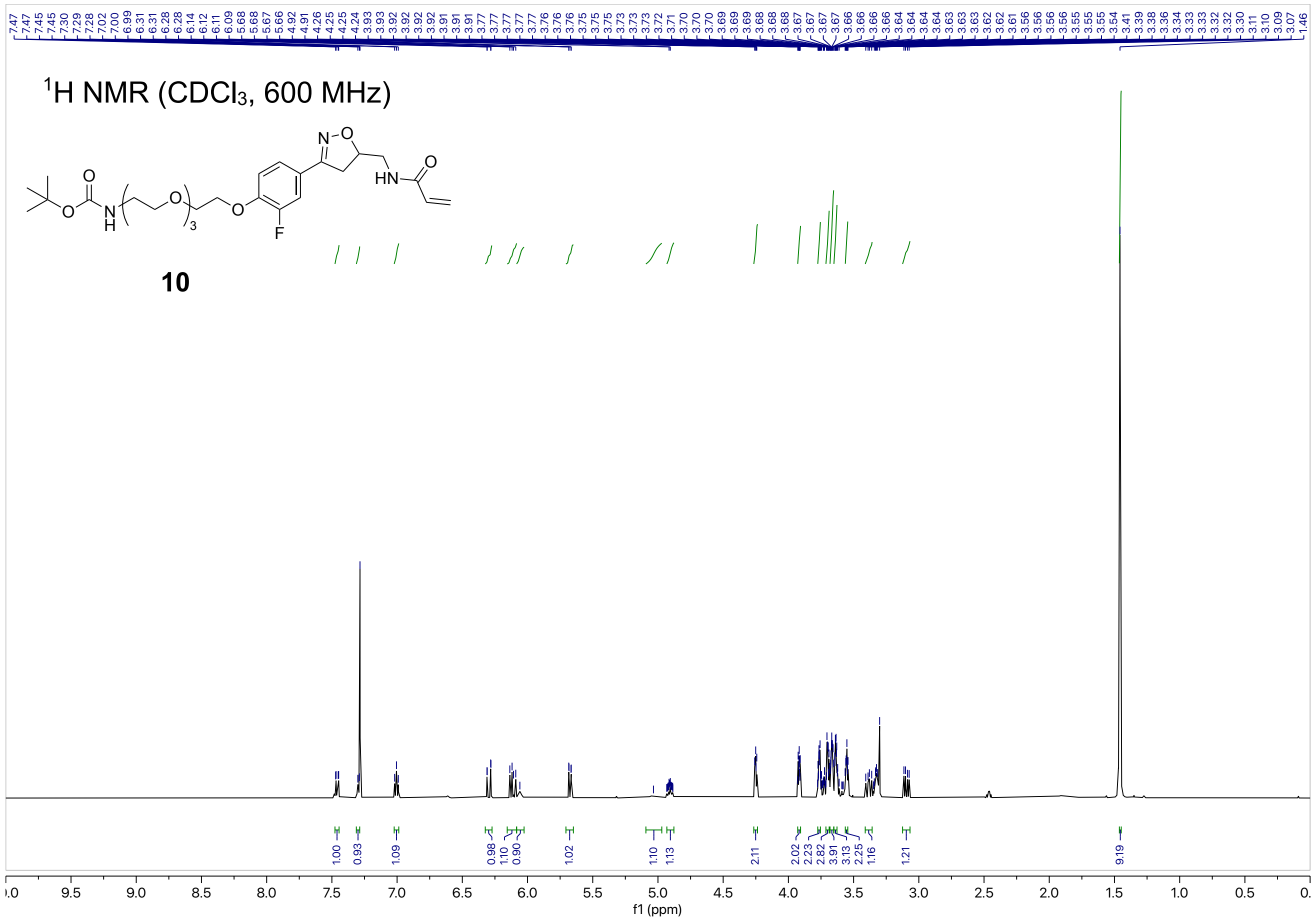

1H NMR (CDCl3, 600 MHz)
10
f1 (ppm)


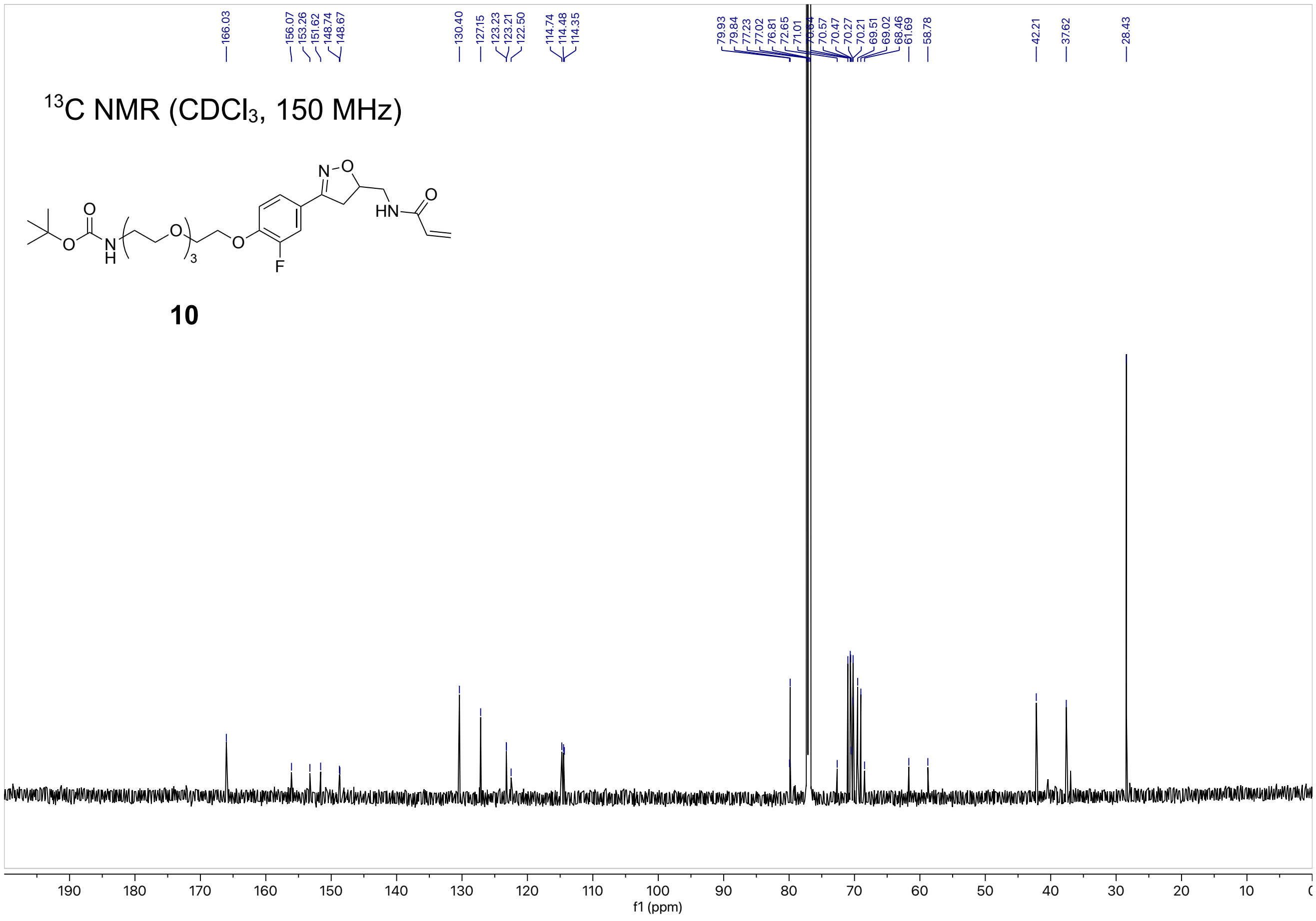

13C NMR (CDCl3, 150 MHz)
10
f1 (ppm)

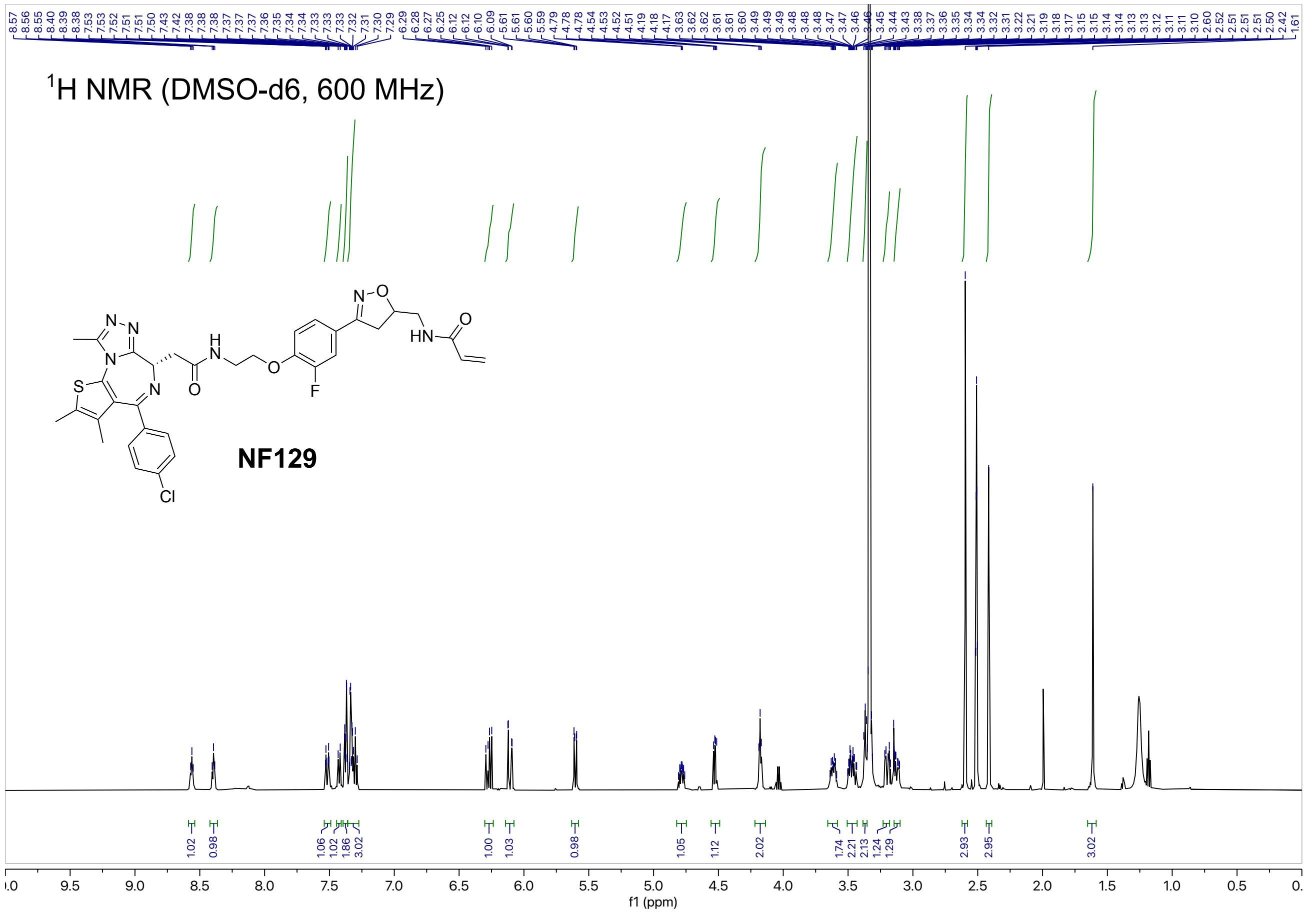
1H NMR (DMSO-d6, 600 MHz)
NF129
f1 (ppm)

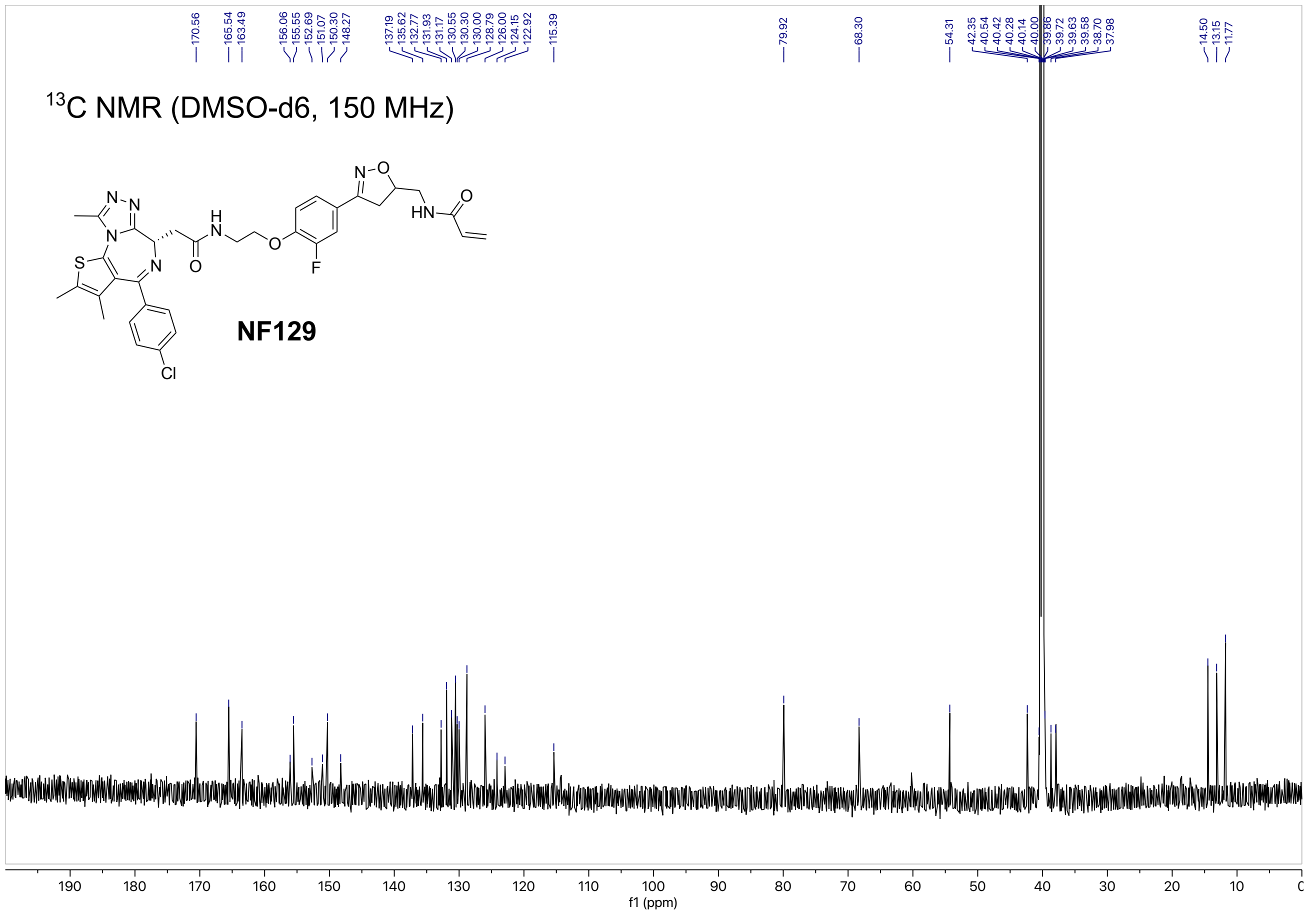
13C NMR (DMSO-d6, 150 MHz)
NF129
f1 (ppm)

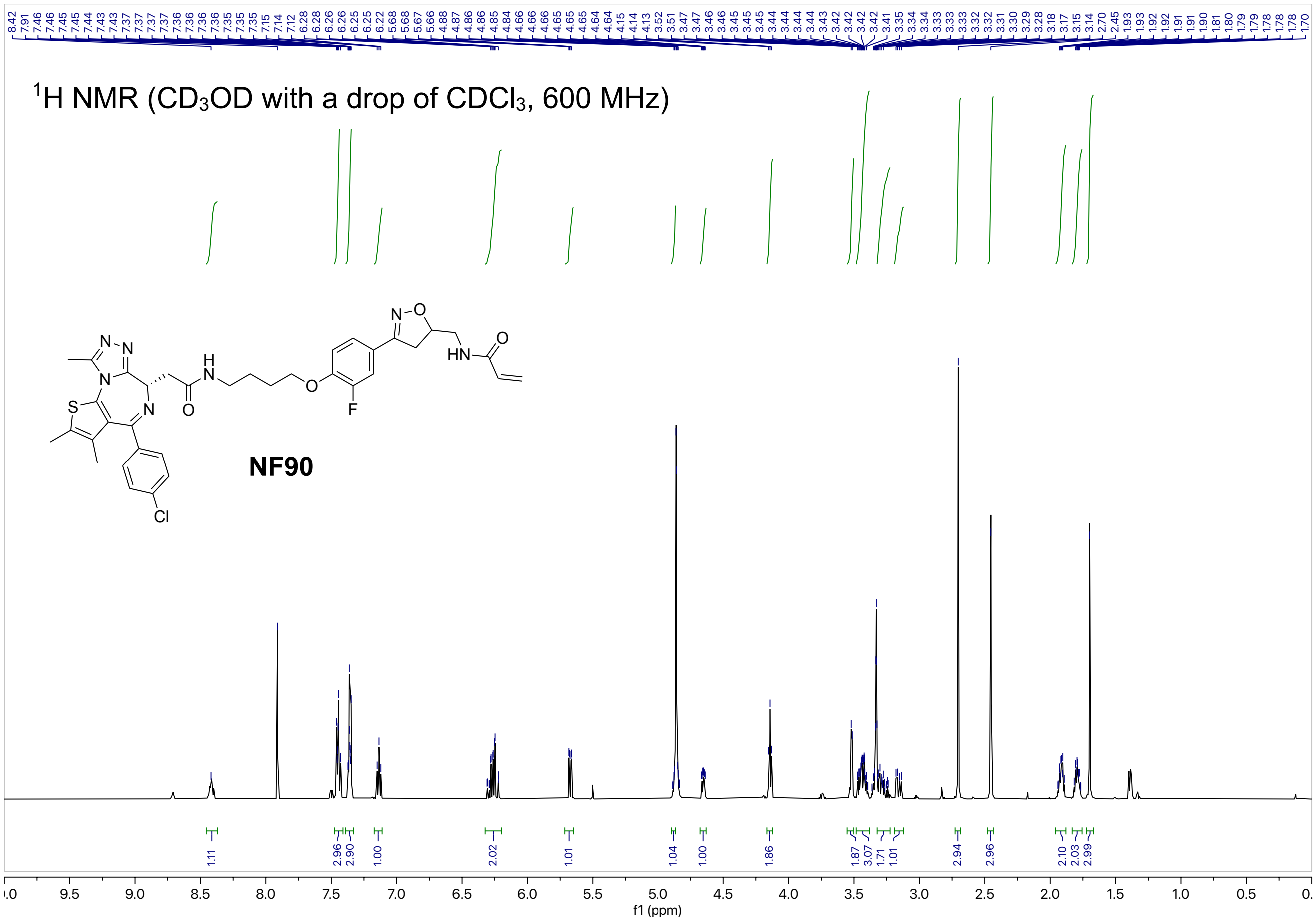
$^1$H NMR ($CD_3OD$ with a drop of $CDCl_3$, 600 MHz)
NF90
f1 (ppm)

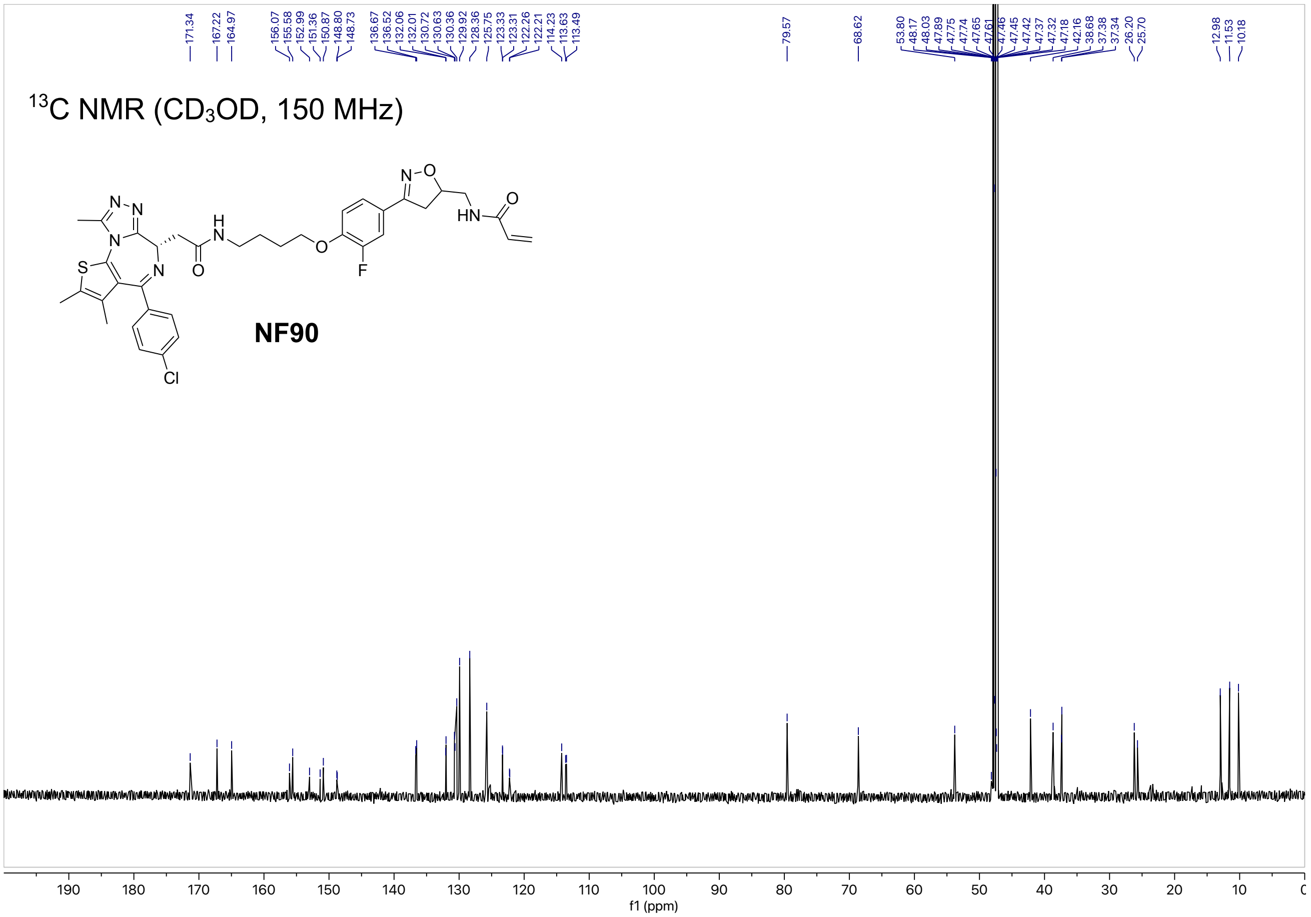
$^{13}$C NMR ($CD_3OD$, 150 MHz)
NF90
f1 (ppm)

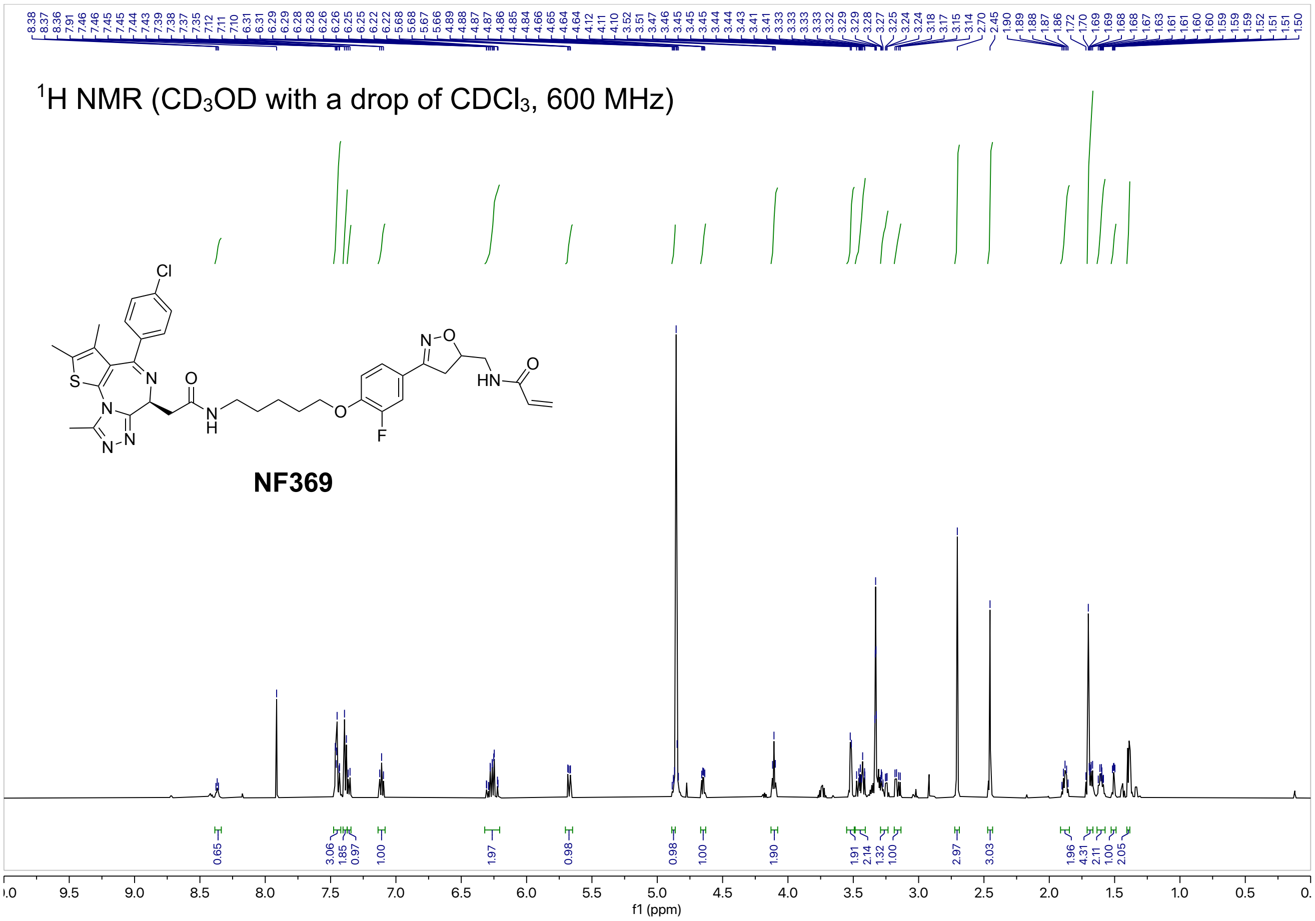

$^1$H NMR (CD$_3$OD with a drop of CDCl$_3$, 600 MHz)
NF369
f1 (ppm)


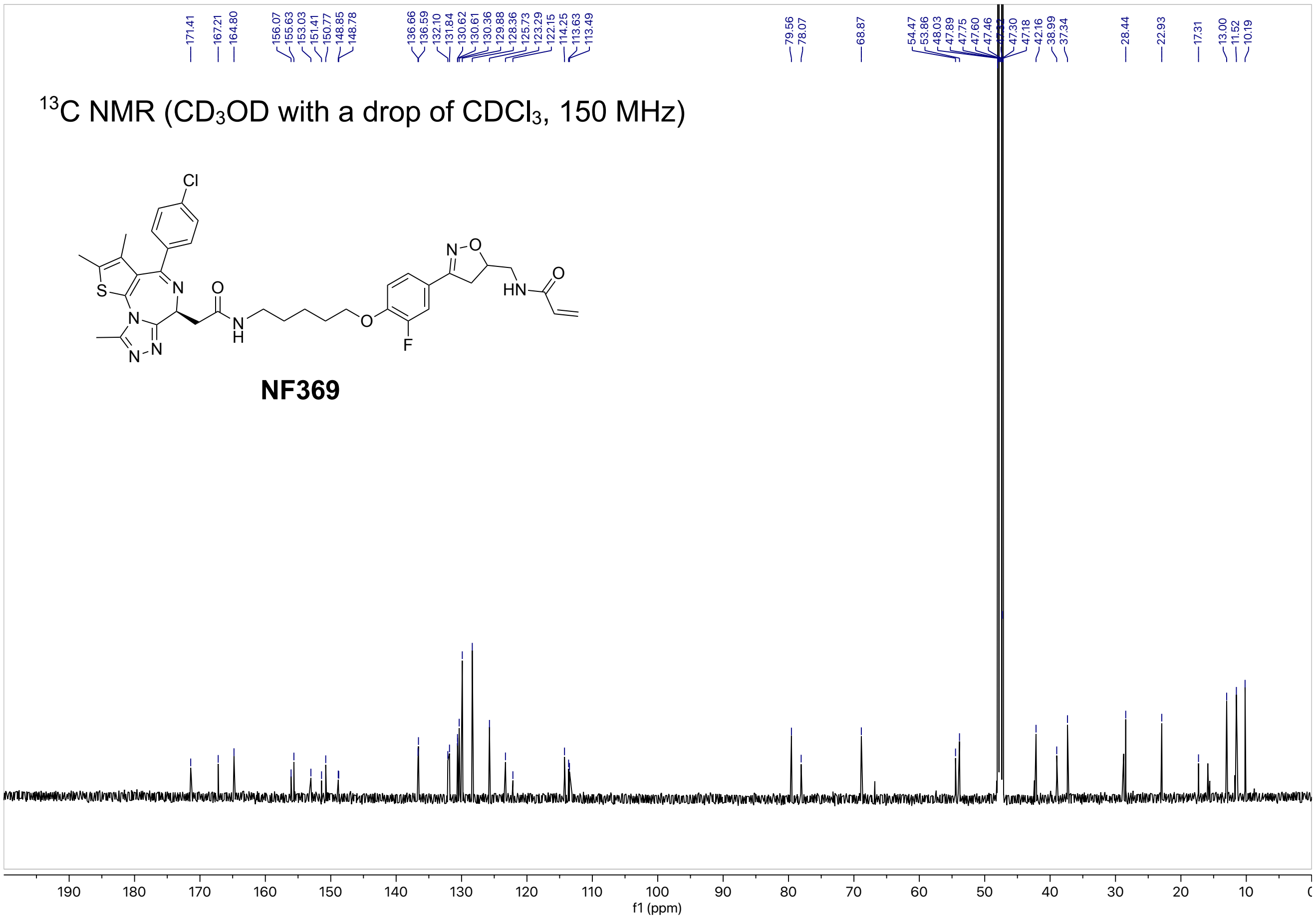

$^{13}$C NMR (CD$_3$OD with a drop of CDCl$_3$, 150 MHz)
NF369
f1 (ppm)

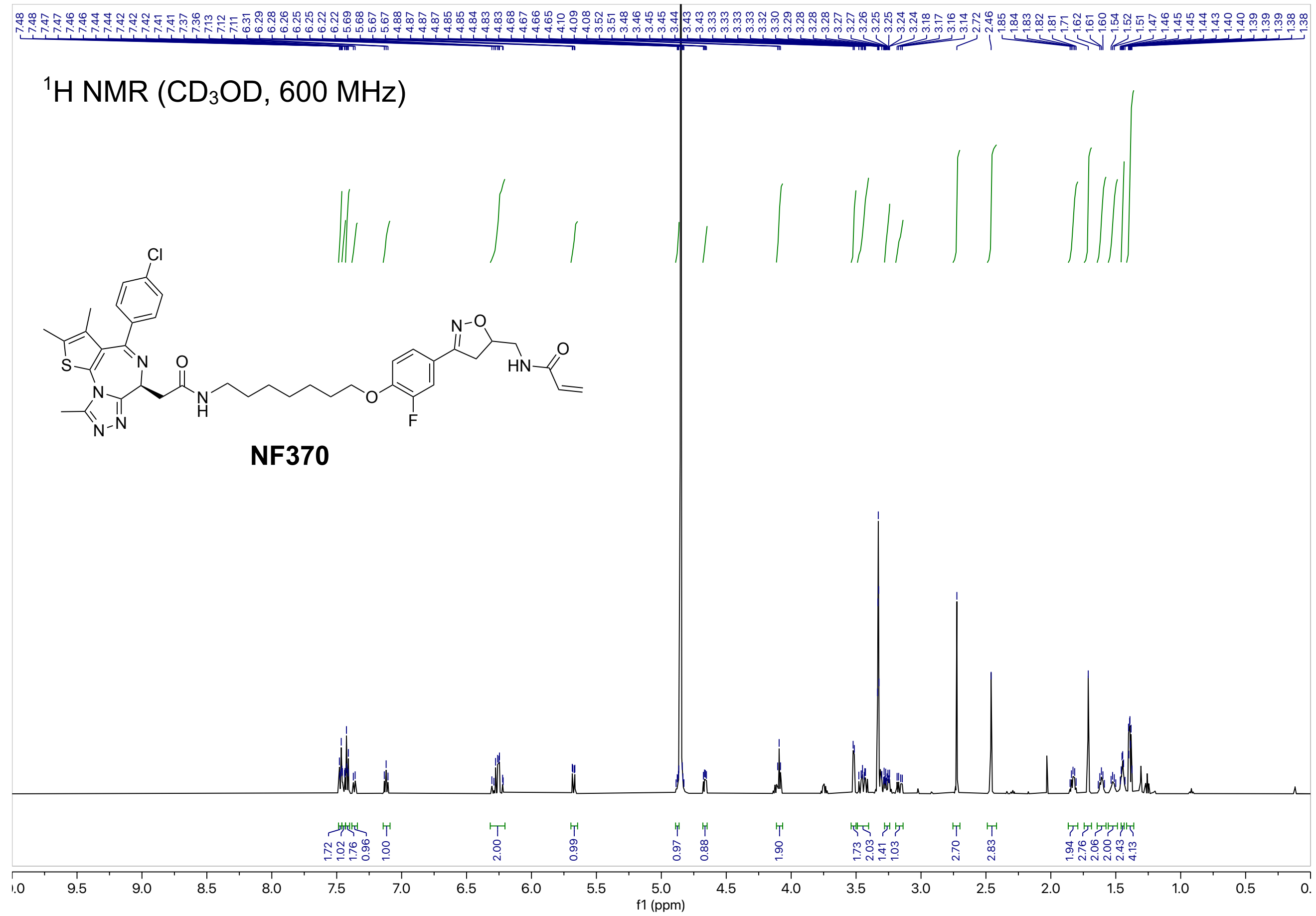
1H NMR (CD3OD, 600 MHz)
NF370
f1 (ppm)

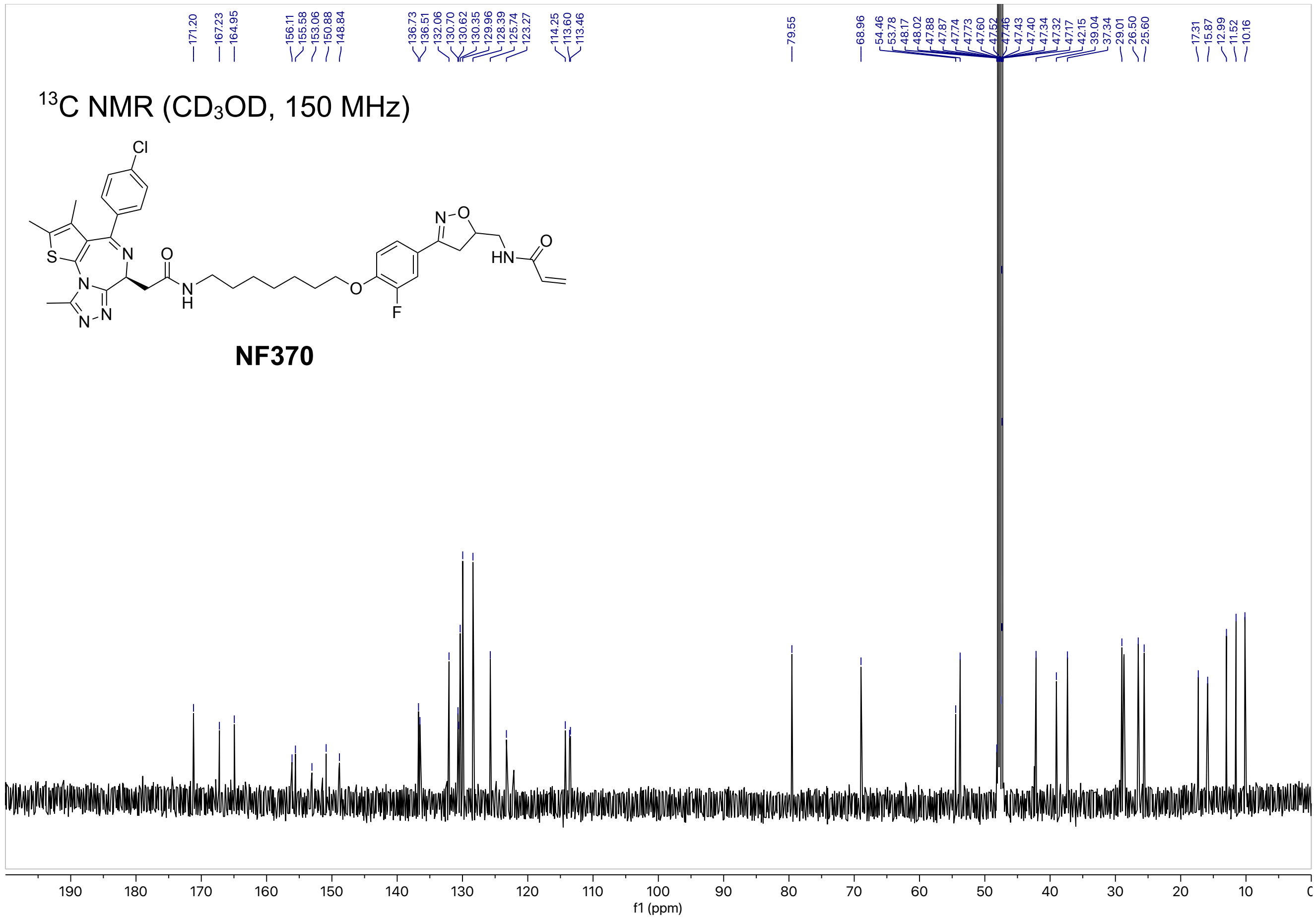
13C NMR (CD3OD, 150 MHz)
NF370
f1 (ppm)

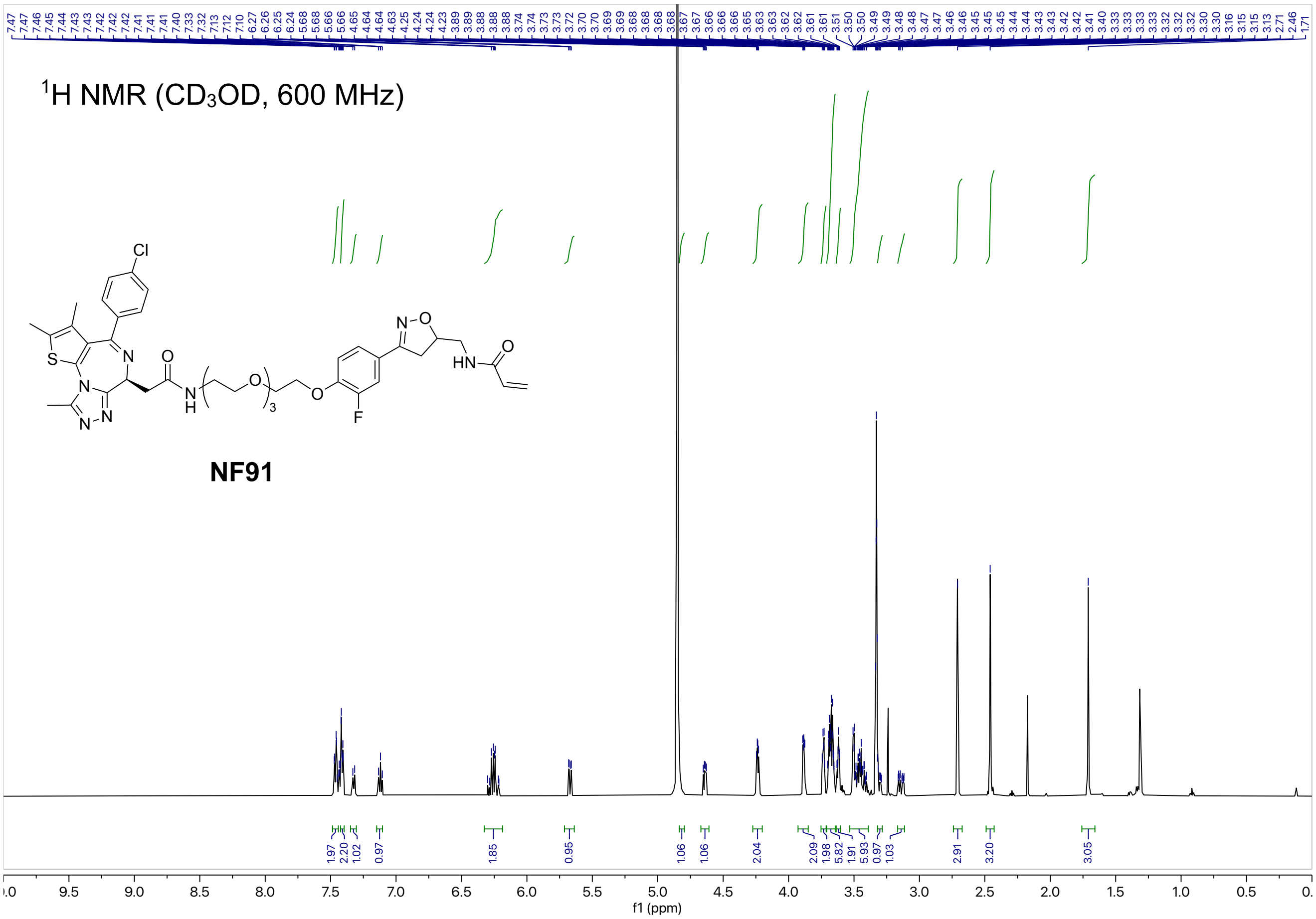

$^1$H NMR ($CD_3OD$, 600 MHz)
NF91
f1 (ppm)


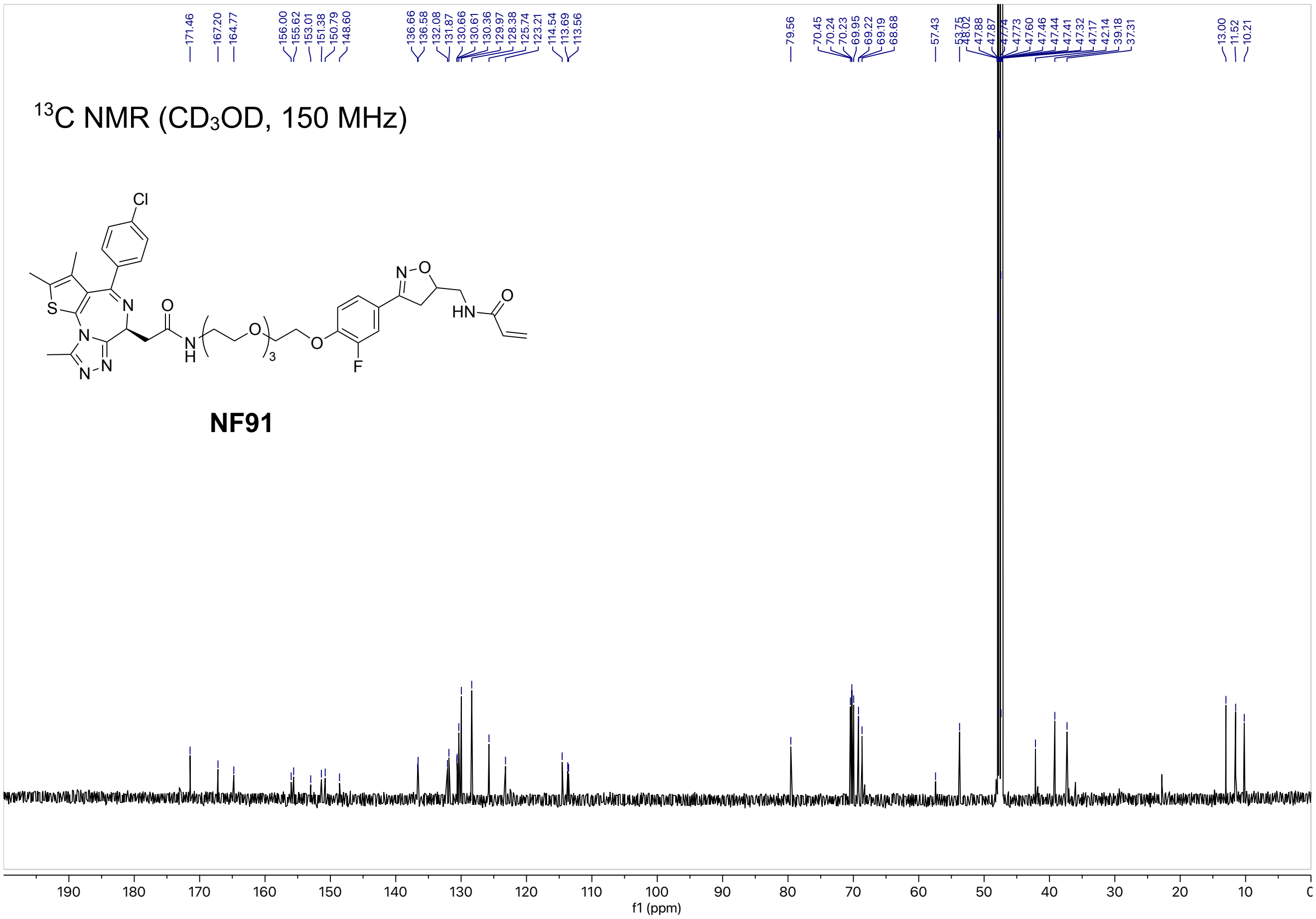

$^{13}$C NMR ($CD_3OD$, 150 MHz)
NF91
f1 (ppm)

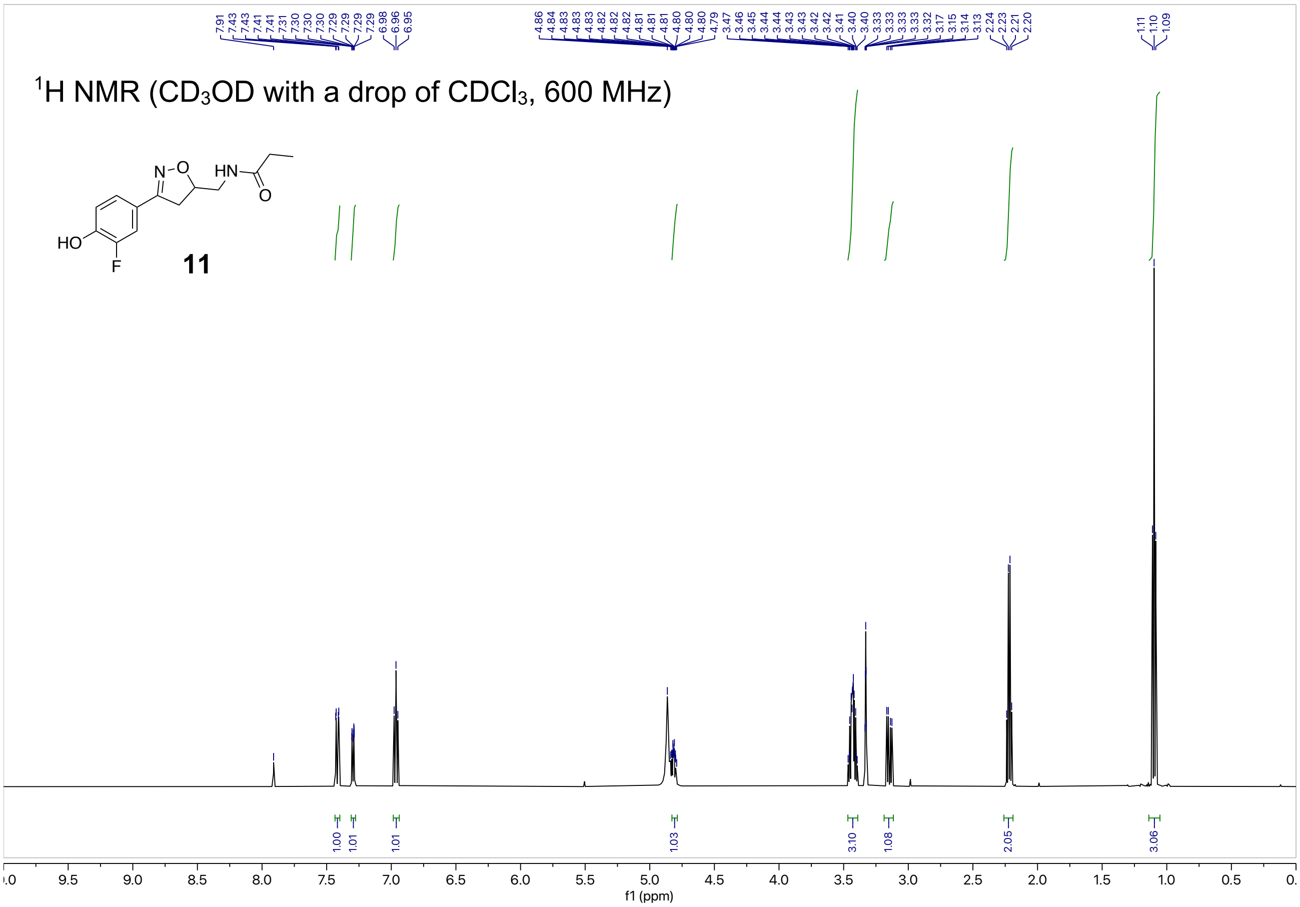

$^1$H NMR (CD$_3$OD with a drop of CDCl$_3$, 600 MHz)
HO
F
HN
O
N-O
11
f1 (ppm)


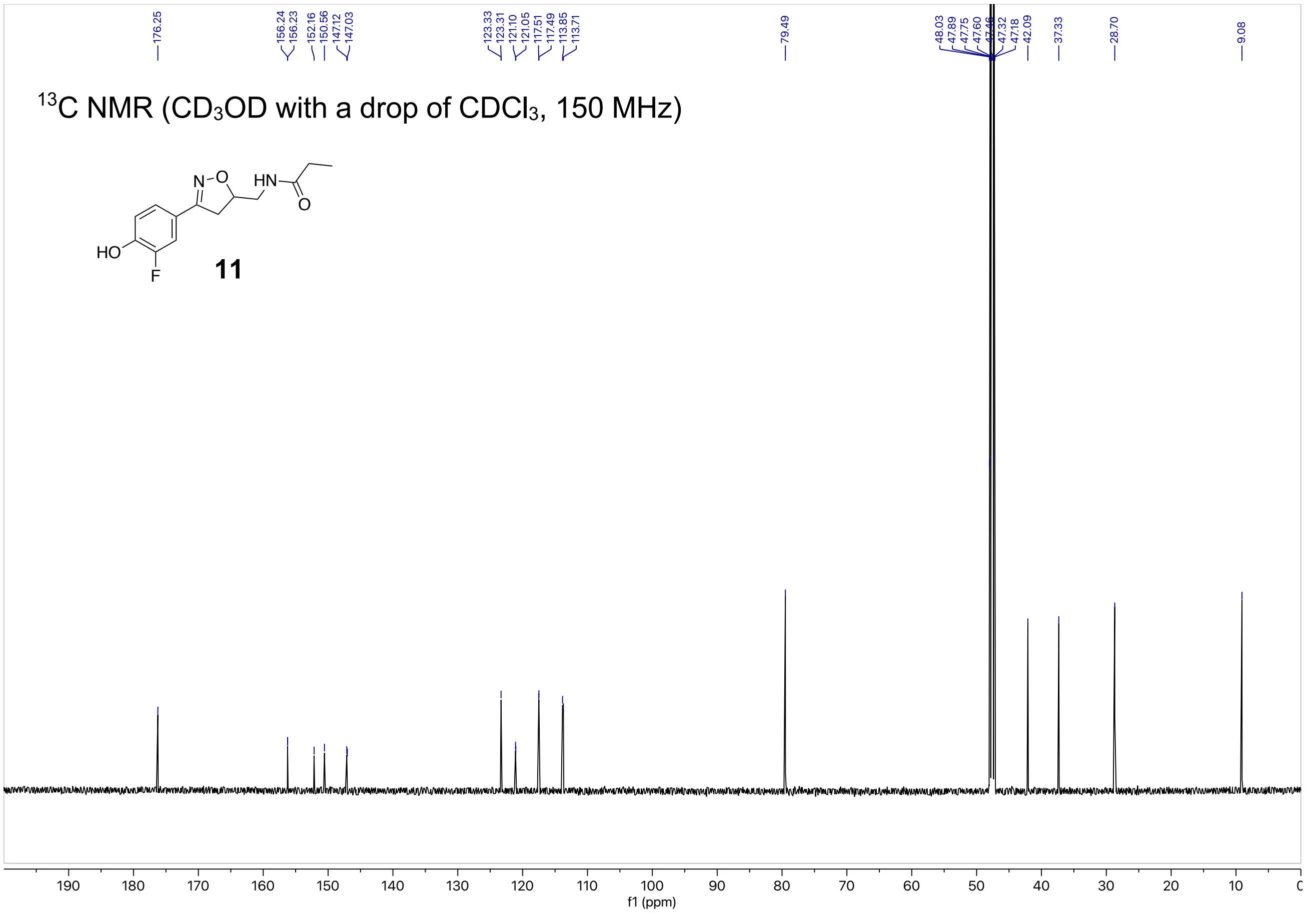

$^{13}$C NMR (CD$_3$OD with a drop of CDCl$_3$, 150 MHz)
HO
F
HN
O
N-O
11
f1 (ppm)

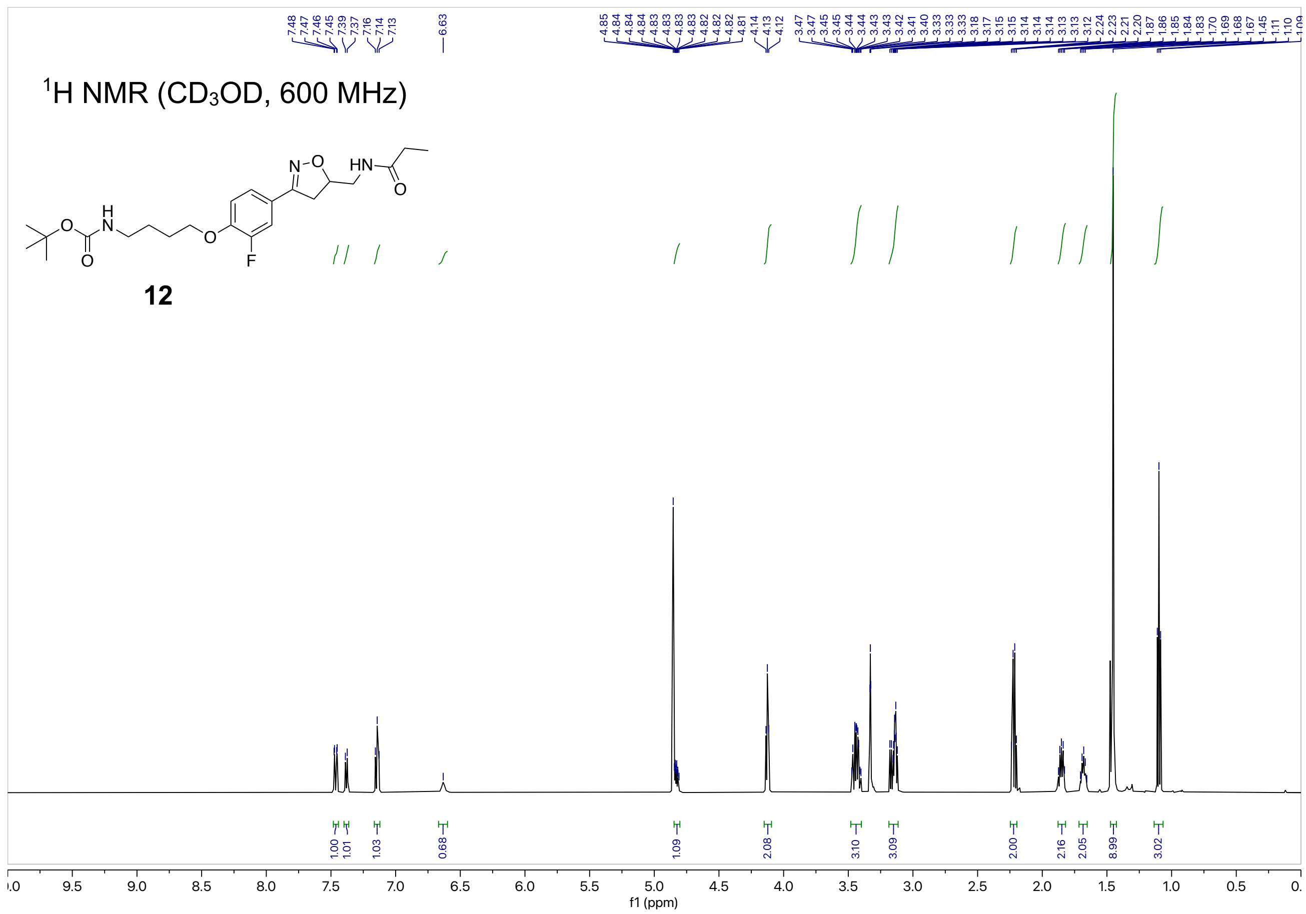

$^1$H NMR ($CD_3OD$, 600 MHz)
12
f1 (ppm)


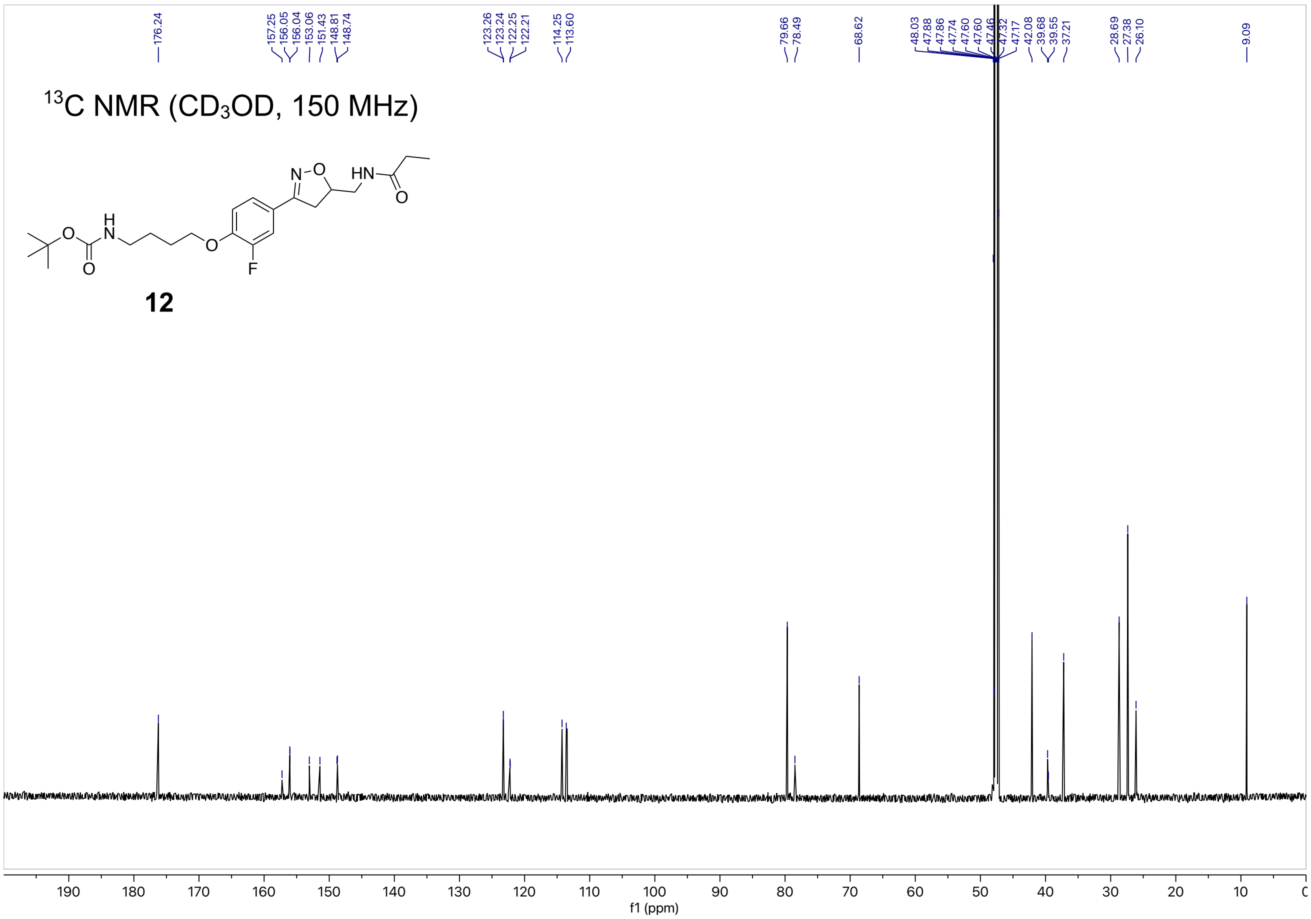

$^{13}$C NMR ($CD_3OD$, 150 MHz)
12
f1 (ppm)

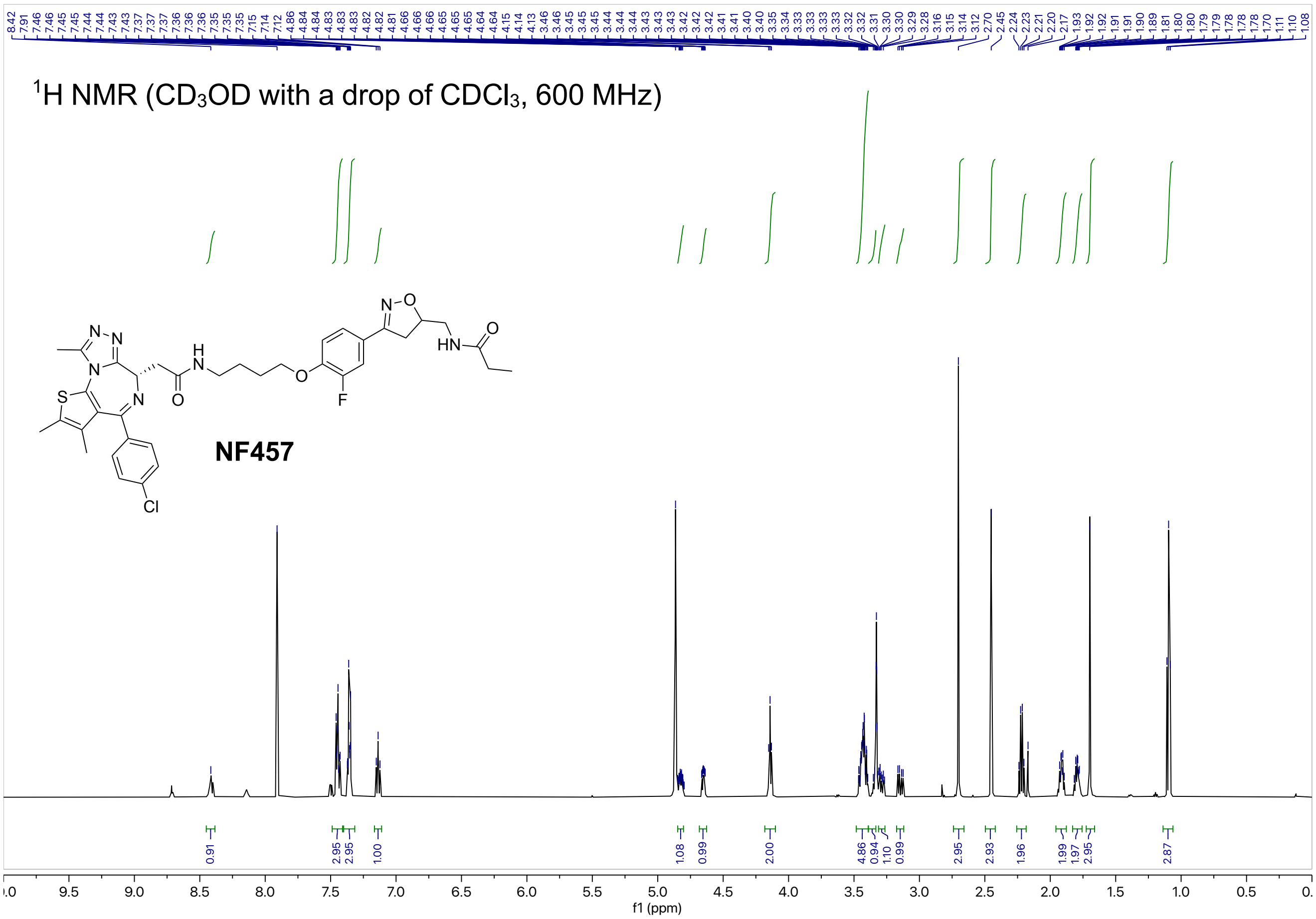

1H NMR (CD3OD with a drop of CDCl3, 600 MHz)
NF457
f1 (ppm)


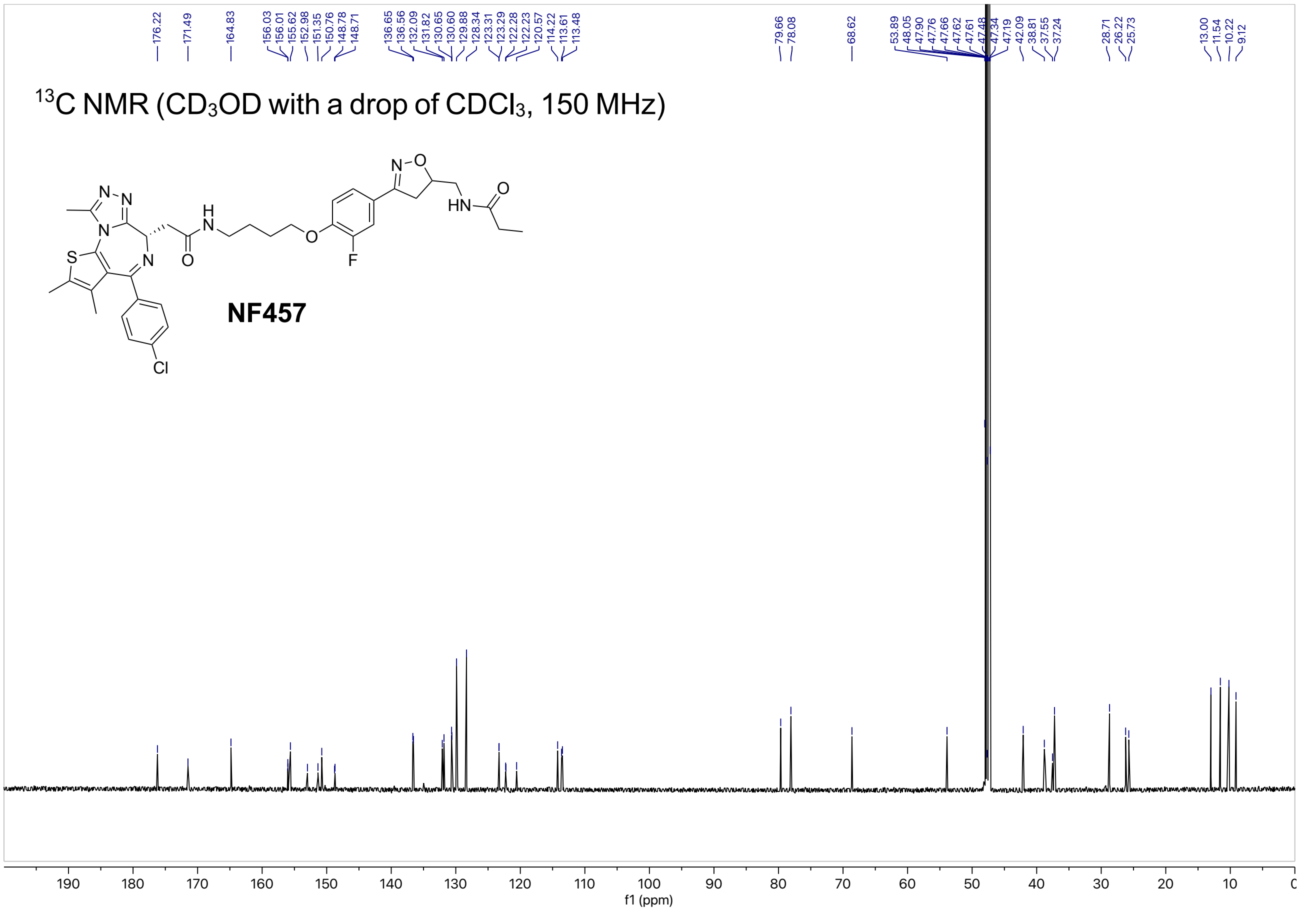

13C NMR (CD3OD with a drop of CDCl3, 150 MHz)
NF457
176.22, 171.49, 164.83, 156.03, 156.01, 155.62, 152.98, 151.35, 150.76, 148.78, 148.71, 136.65, 136.56, 132.09, 131.82, 130.65, 130.60, 129.88, 128.34, 123.31, 123.29, 122.28, 122.23, 120.57, 114.22, 113.61, 113.48, 79.66, 78.08, 68.62, 53.89, 48.05, 47.90, 47.76, 47.62, 47.61, 47.48, 47.34, 47.19, 42.09, 38.81, 37.55, 37.24, 28.71, 26.22, 25.73, 13.00, 11.54, 10.22, 9.12
f1 (ppm)

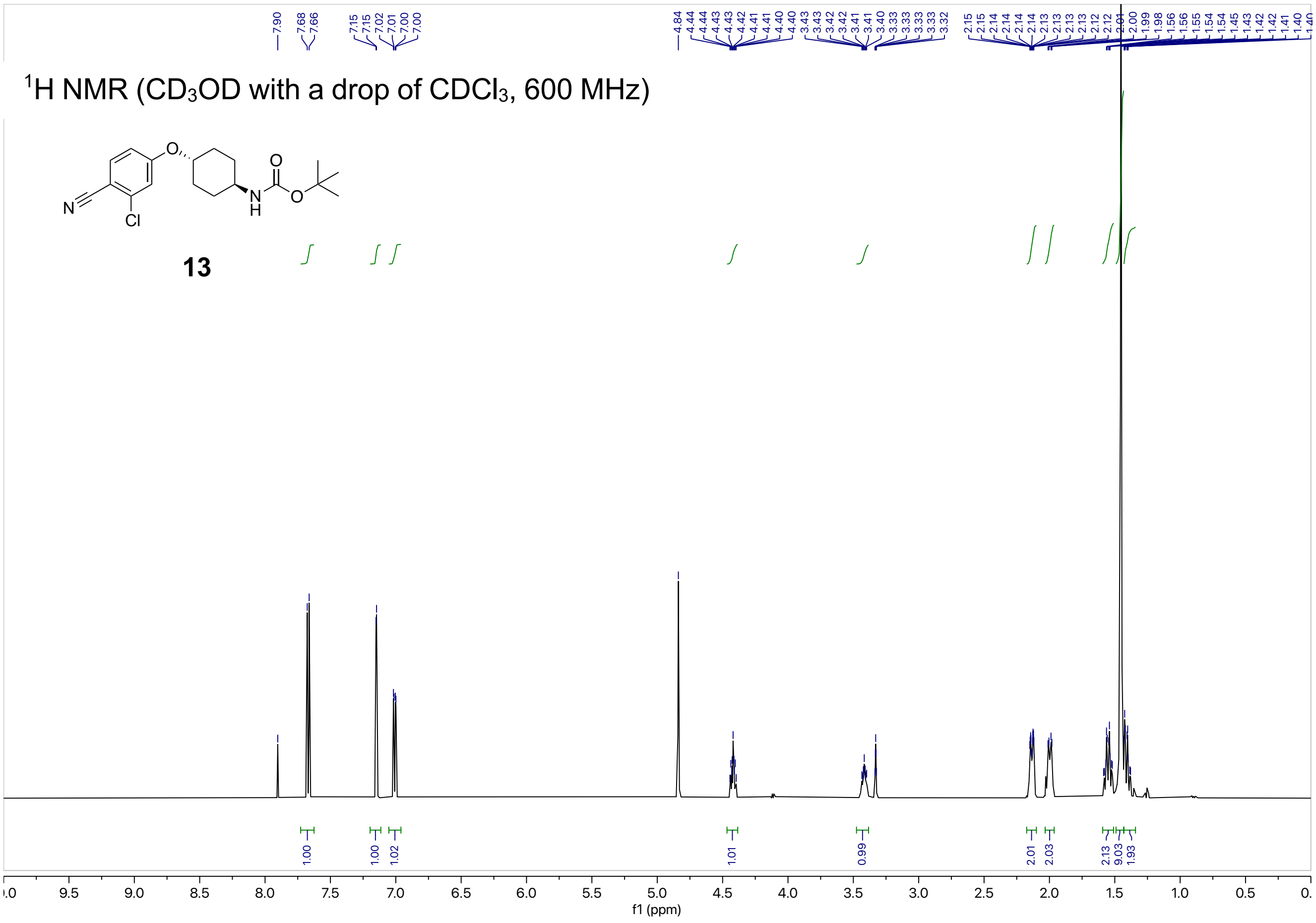

7.90
7.68
7.66
7.15
7.15
7.02
7.01
7.00
7.00
4.84
4.44
4.44
4.43
4.43
4.42
4.41
4.41
4.40
4.40
3.43
3.43
3.42
3.42
3.41
3.41
3.40
3.33
3.33
3.33
3.33
3.32
2.15
2.15
2.14
2.14
2.14
2.14
2.13
2.13
2.13
2.13
2.12
2.12
2.01
2.00
1.99
1.98
1.56
1.56
1.55
1.54
1.54
1.45
1.43
1.42
1.42
1.41
1.40
1.40
$^{1}$H NMR (CD$_3$OD with a drop of CDCl$_3$, 600 MHz)
O
O
N
H
O
N
Cl
13
1.00
1.00
1.02
1.01
0.99
2.01
2.03
2.13
9.03
1.93
9.5
9.0
8.5
8.0
7.5
7.0
6.5
6.0
5.5
5.0
4.5
4.0
3.5
3.0
2.5
2.0
1.5
1.0
0.5
f1 (ppm)


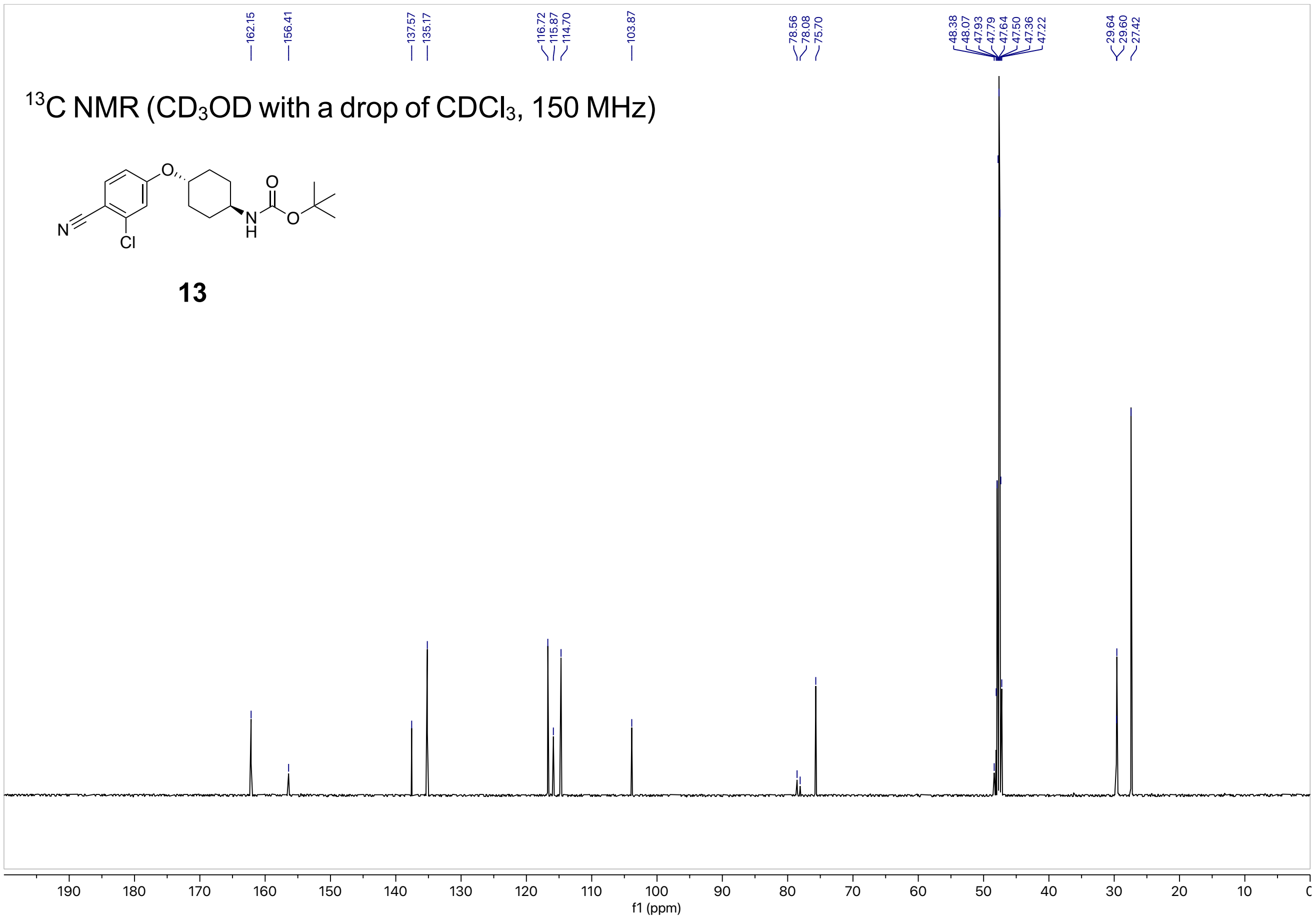

162.15
156.41
137.57
135.17
116.72
115.87
114.70
103.87
78.56
78.08
75.70
48.38
48.07
47.93
47.79
47.64
47.50
47.36
47.22
29.64
29.60
27.42
$^{13}$C NMR (CD$_3$OD with a drop of CDCl$_3$, 150 MHz)
O
O
N
H
O
N
Cl
13
190
180
170
160
150
140
130
120
110
100
90
80
70
60
50
40
30
20
10
f1 (ppm)

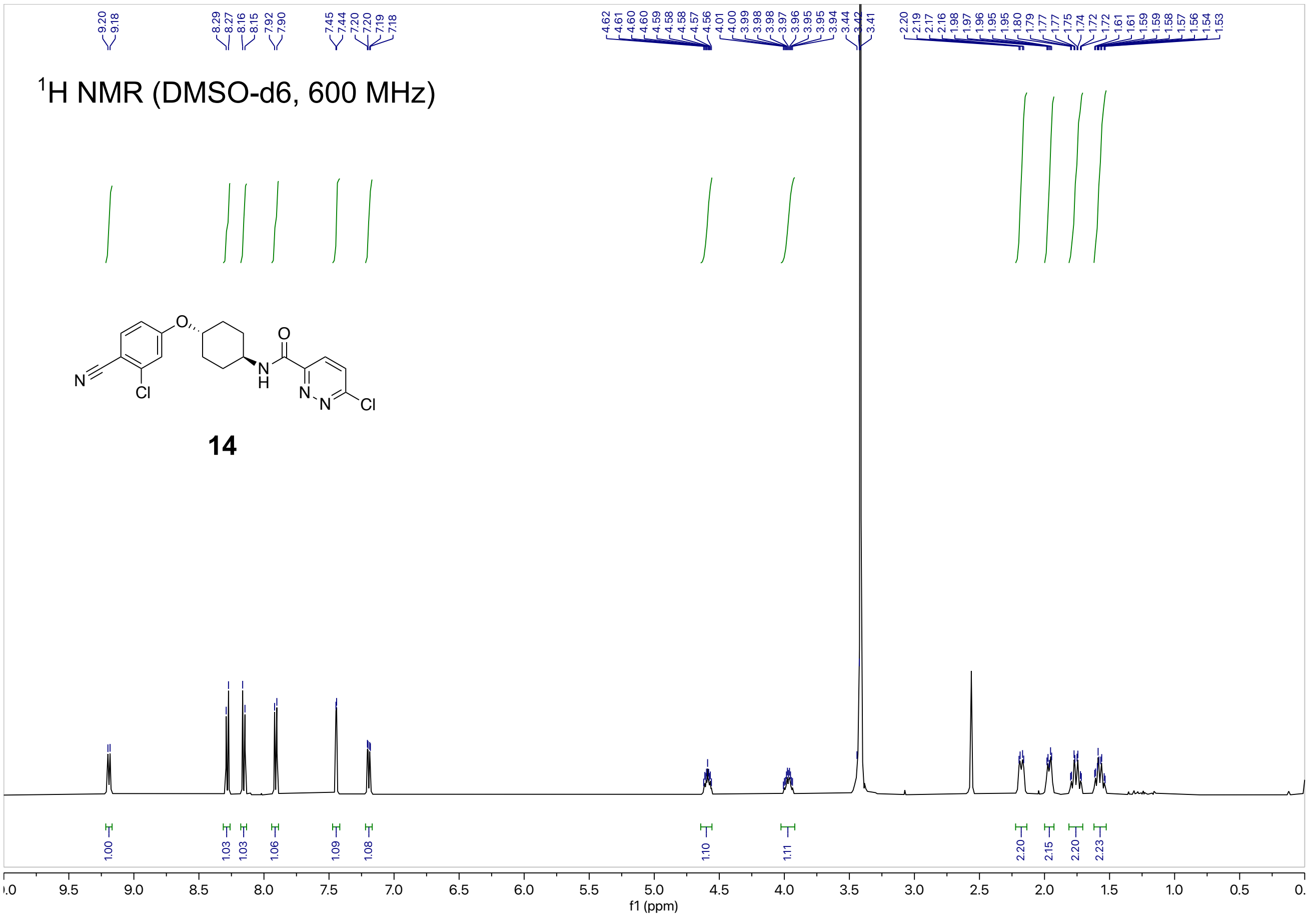
1H NMR (DMSO-d6, 600 MHz)
9.20 9.18 8.29 8.27 8.16 8.15 7.92 7.90 7.45 7.44 7.20 7.20 7.19 7.18
4.62 4.61 4.60 4.60 4.59 4.58 4.58 4.57 4.56 4.01 4.00 3.99 3.98 3.98 3.97 3.96 3.95 3.95 3.94 3.44 3.42 3.41
2.20 2.19 2.17 2.16 1.98 1.97 1.96 1.95 1.95 1.80 1.79 1.77 1.77 1.75 1.74 1.72 1.72 1.61 1.61 1.59 1.58 1.57 1.56 1.54 1.53
N
O
Cl
N
H
O
N
N
Cl
14
1.00 1.03 1.03 1.06 1.09 1.08 1.10 1.11 2.20 2.15 2.20 2.23
f1 (ppm)

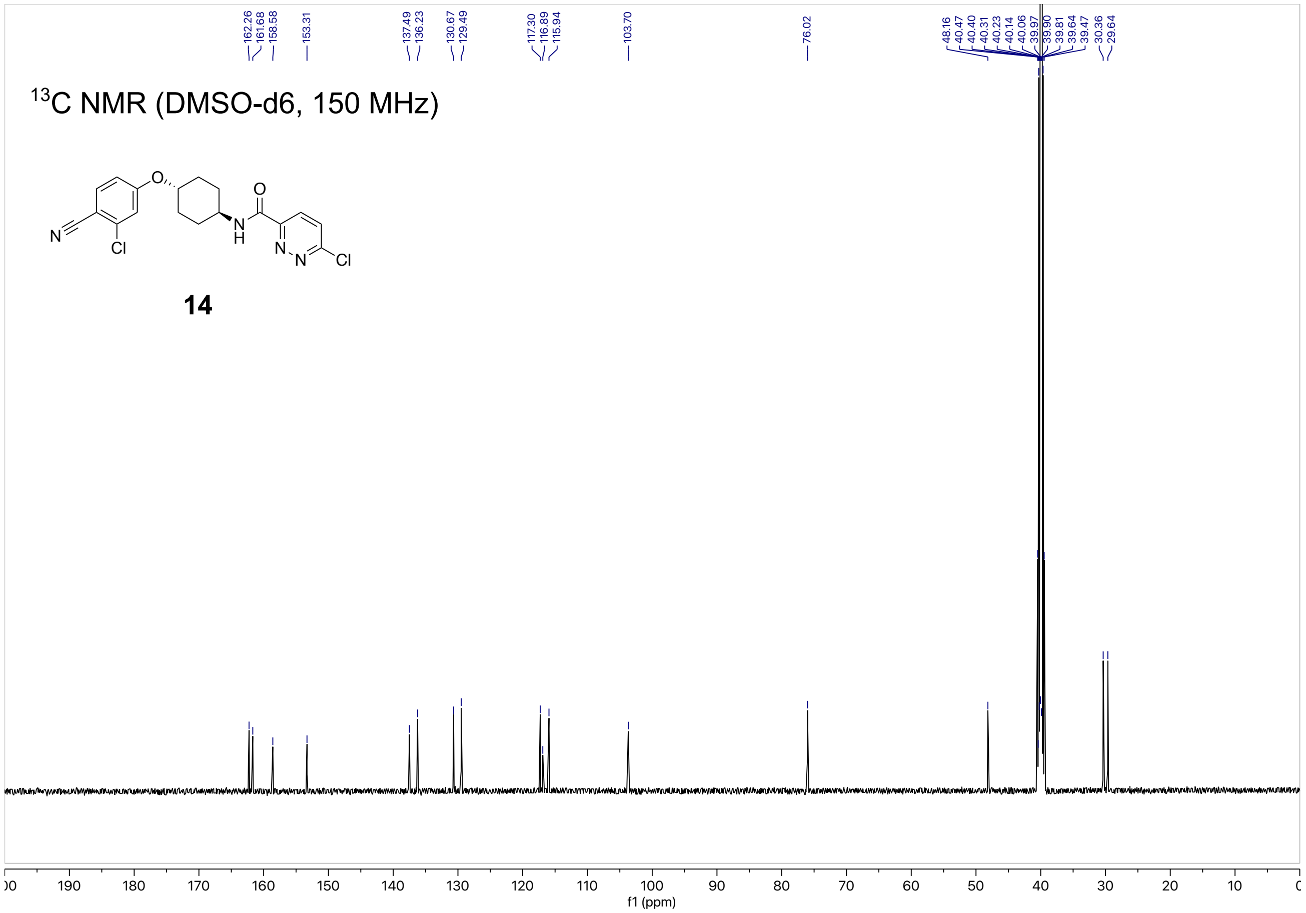
13C NMR (DMSO-d6, 150 MHz)
162.26 161.68 158.58 153.31 137.49 136.23 130.67 129.49 117.30 116.89 115.94 103.70 76.02
48.16 40.47 40.40 40.31 40.23 40.14 40.06 39.97 39.90 39.81 39.64 39.47 30.36 29.64
N
O
Cl
N
H
O
N
N
Cl
14
f1 (ppm)

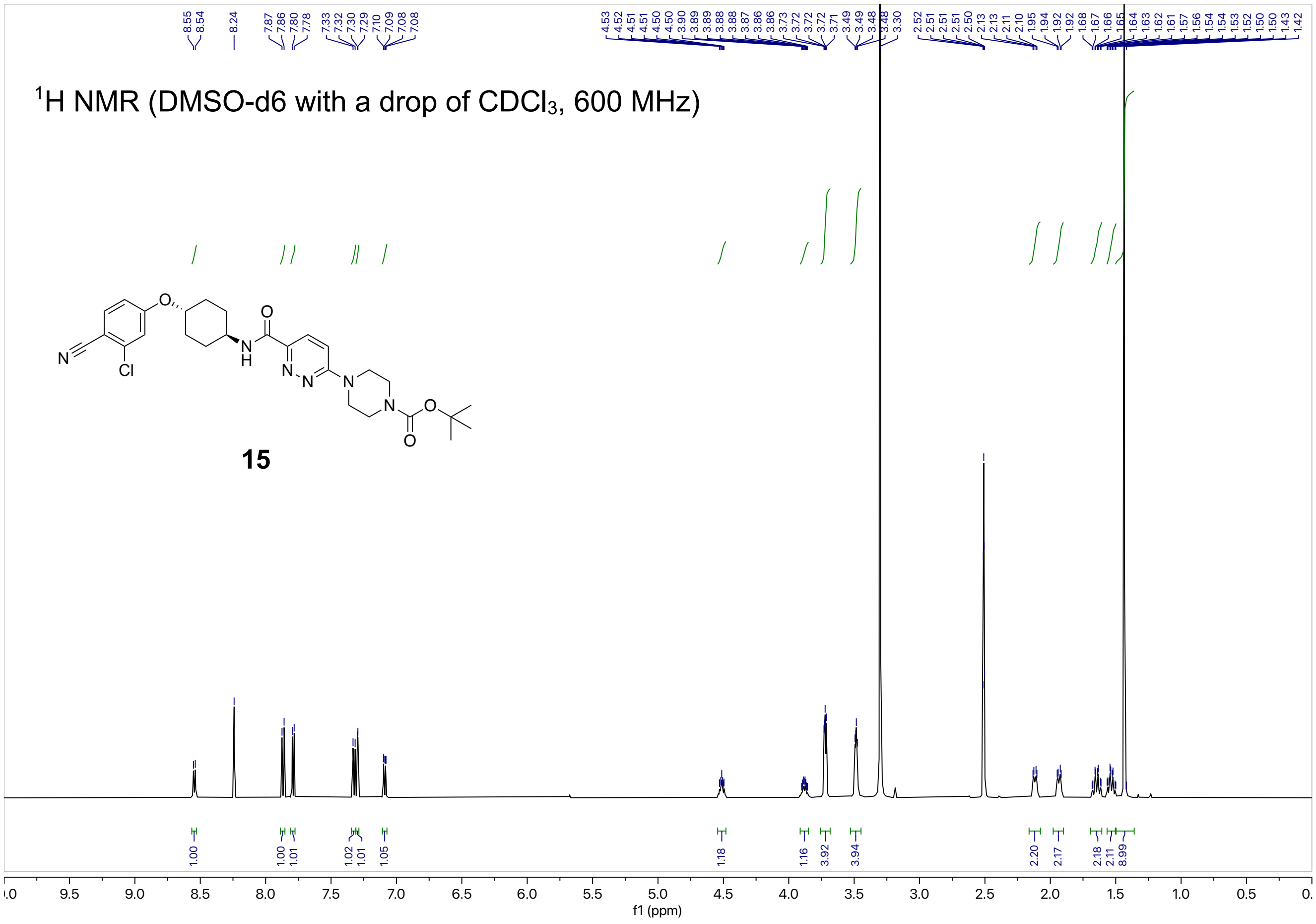

$^1$H NMR (DMSO-d6 with a drop of CDCl3, 600 MHz)
15
f1 (ppm)


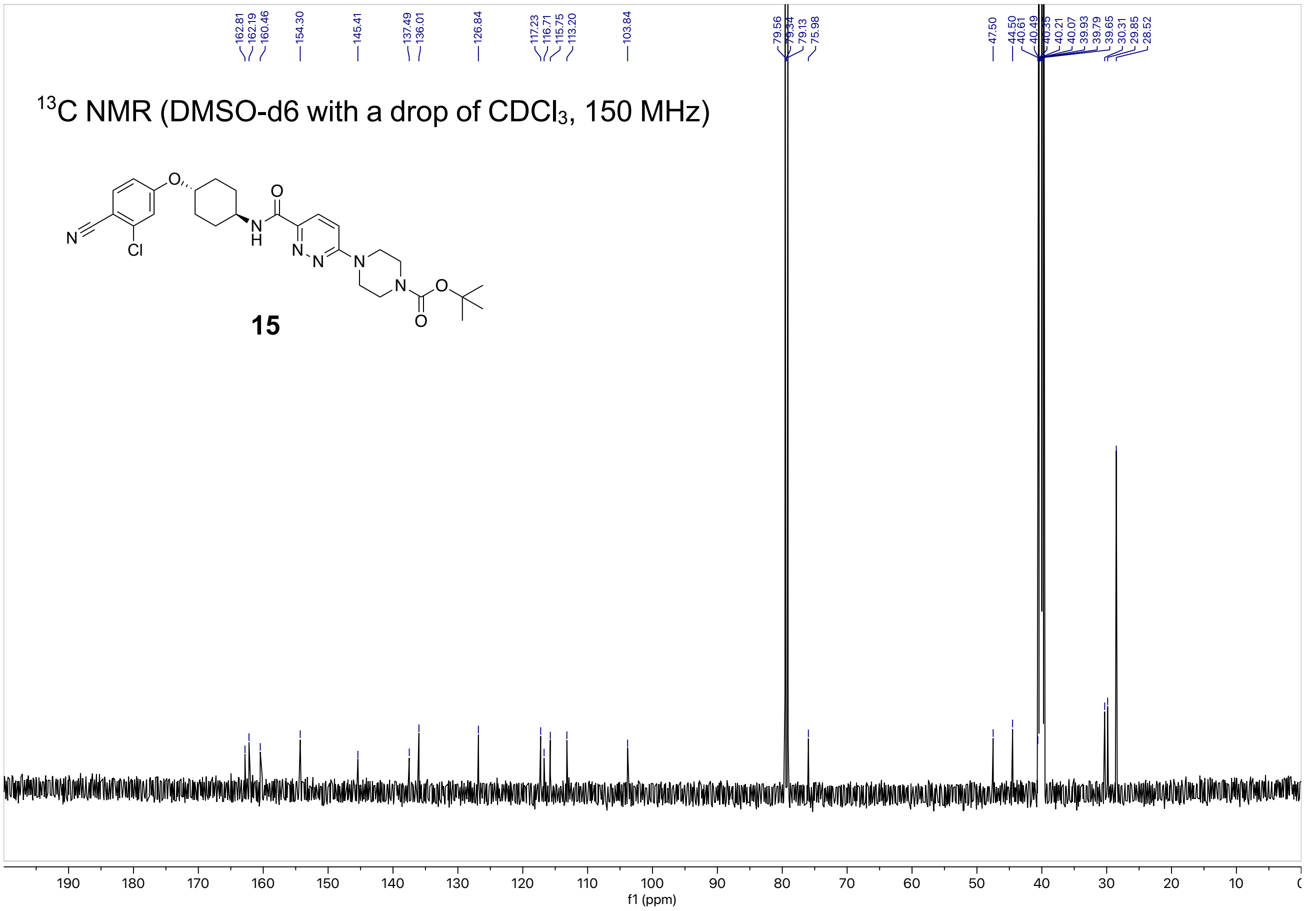

$^{13}$C NMR (DMSO-d6 with a drop of CDCl3, 150 MHz)
15
f1 (ppm)

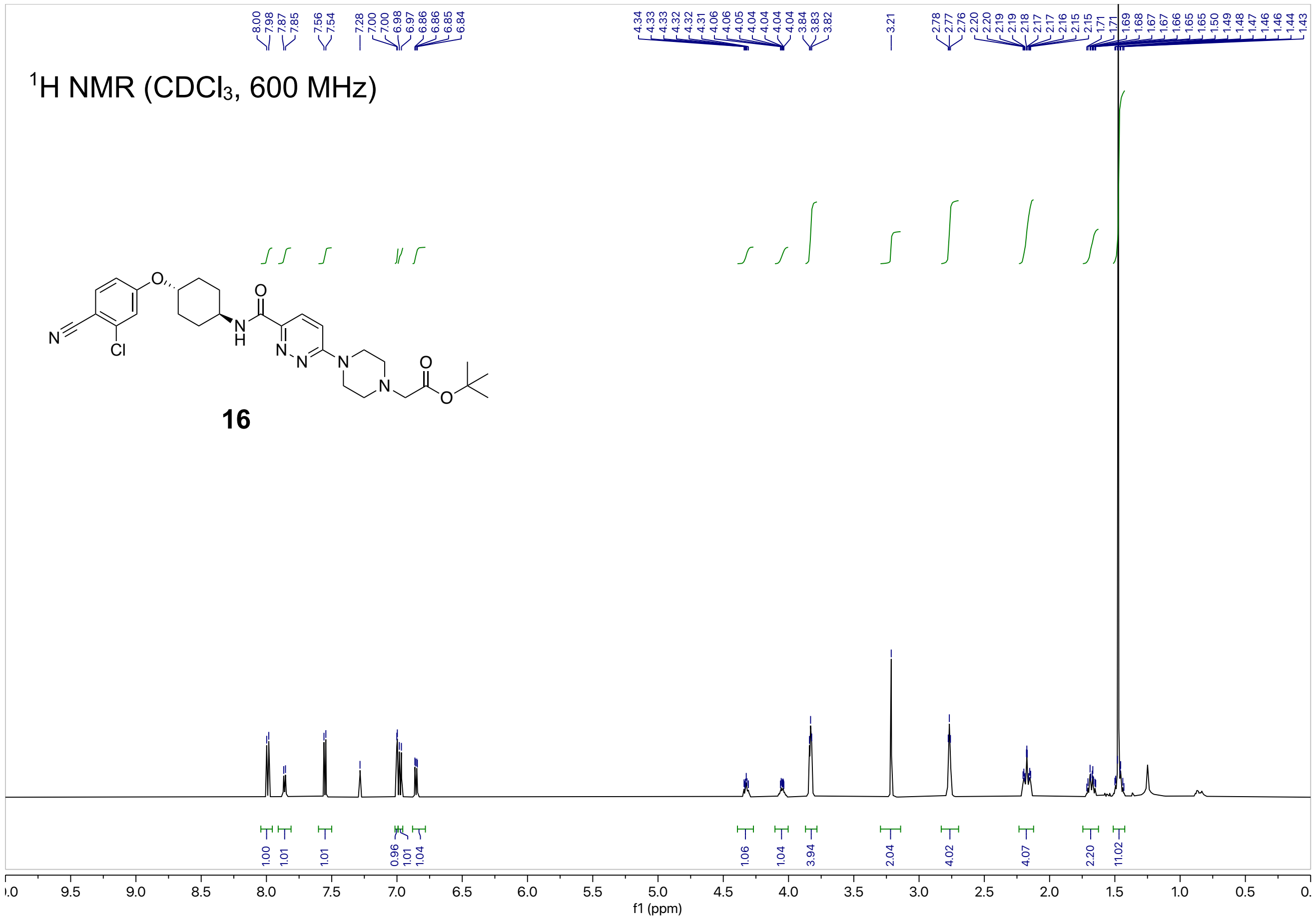

$^1$H NMR (CDCl$_3$, 600 MHz)
16
f1 (ppm)


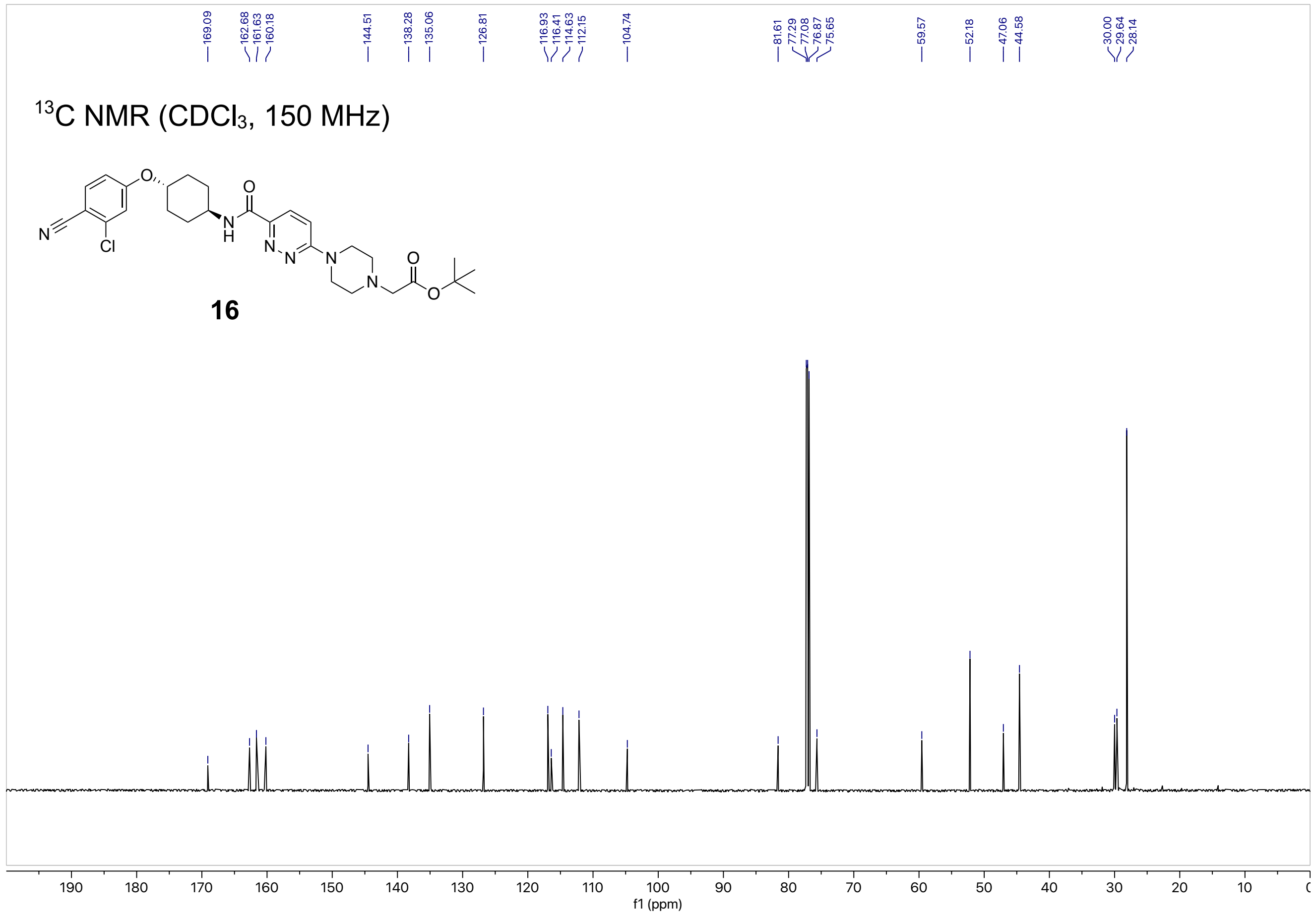

$^{13}$C NMR (CDCl$_3$, 150 MHz)
16
f1 (ppm)

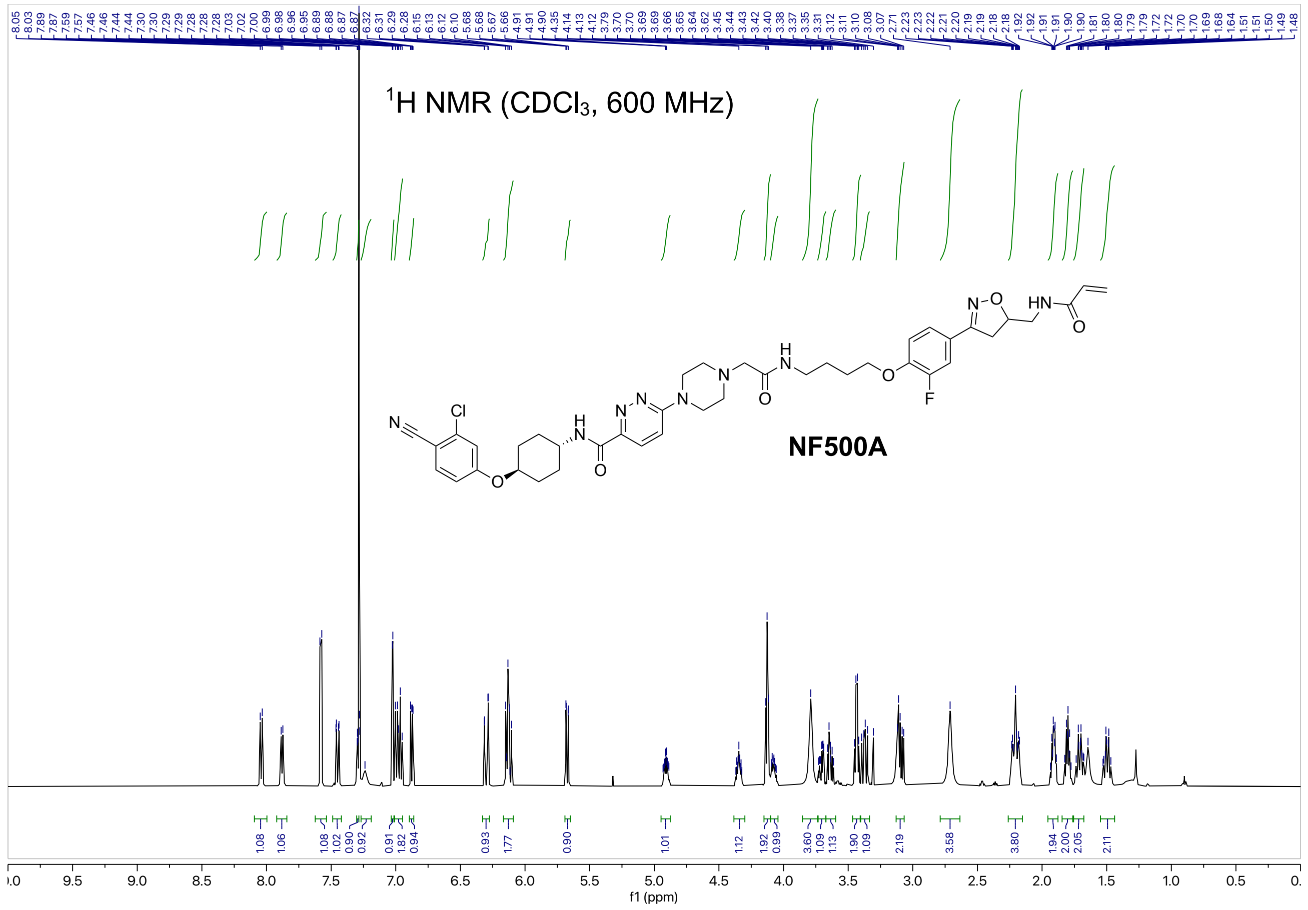

1H NMR (CDCl3, 600 MHz)
NF500A
f1 (ppm)


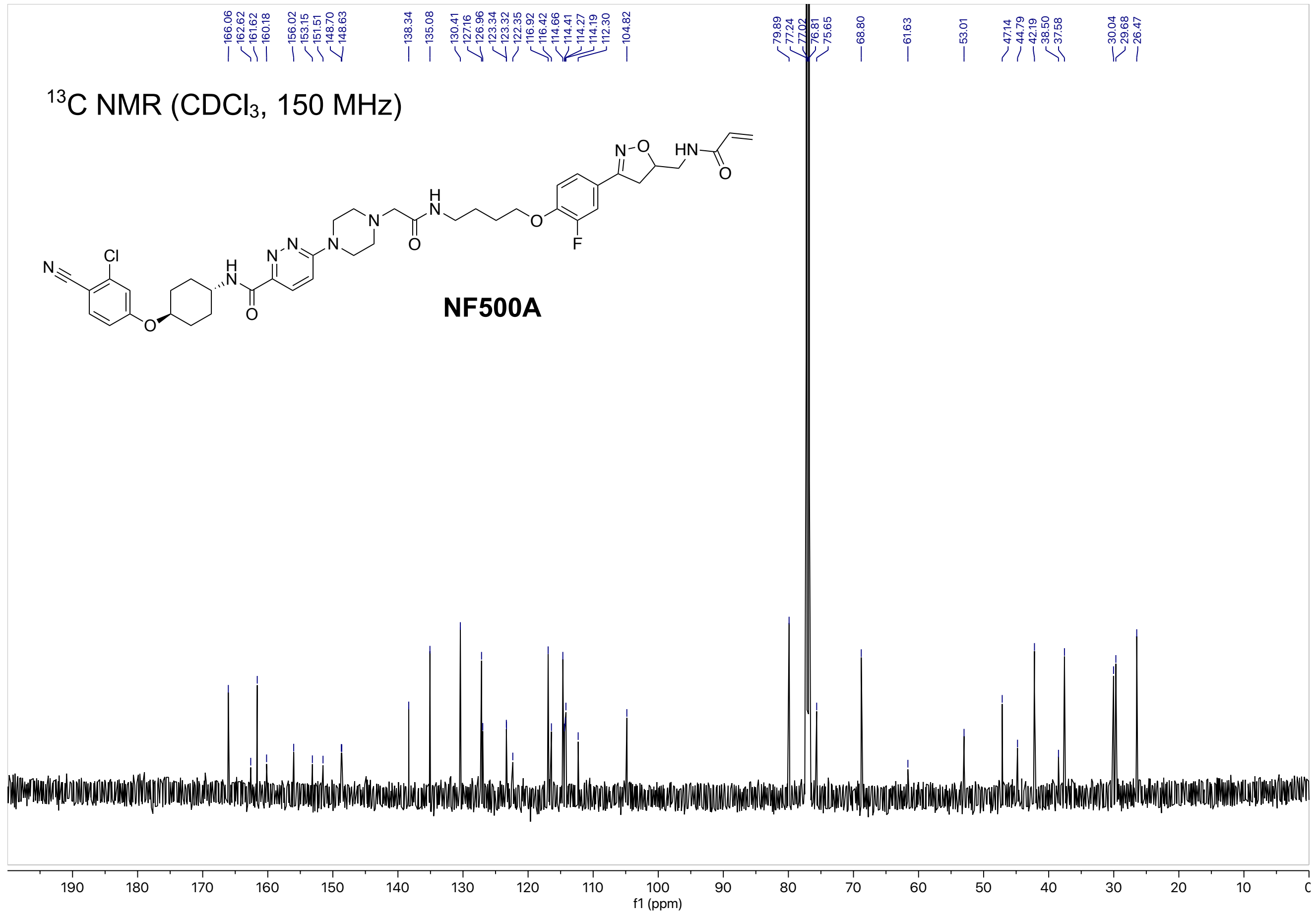

13C NMR (CDCl3, 150 MHz)
NF500A
f1 (ppm)

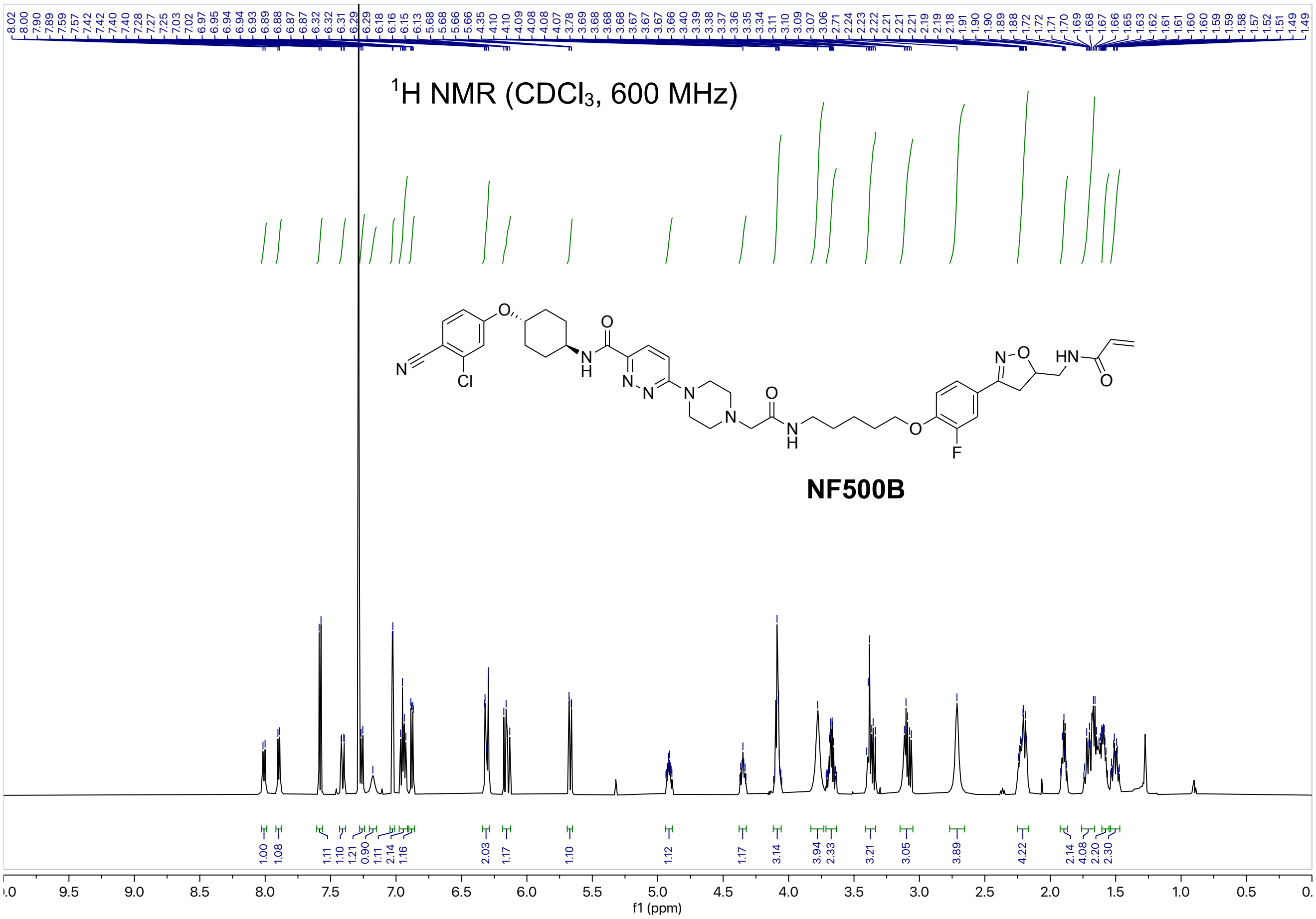

¹H NMR (CDCl₃, 600 MHz)
NF500B
f1 (ppm)


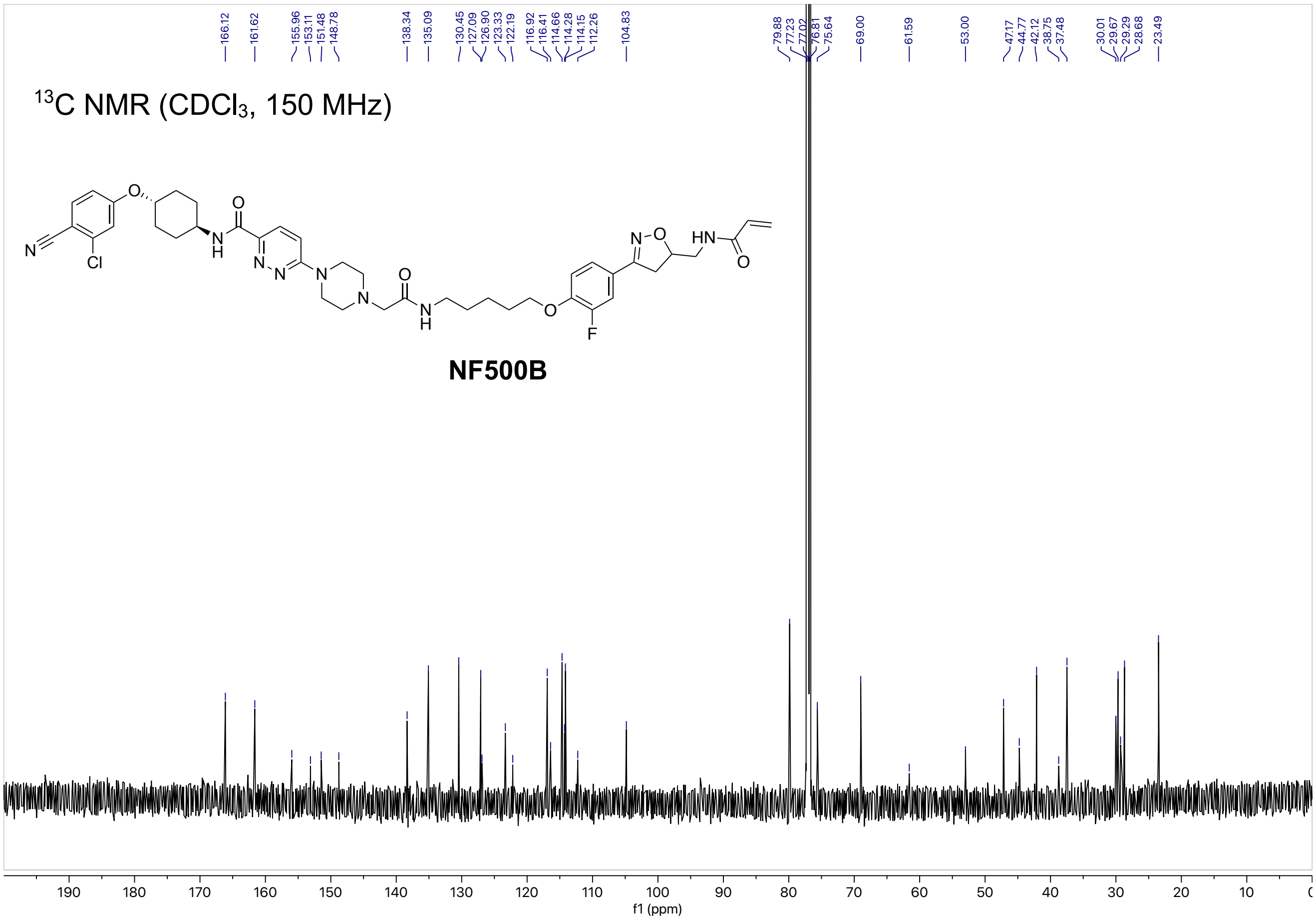

166.12
161.62
155.96
153.11
151.48
148.78
138.34
135.09
130.45
127.09
126.90
123.33
122.19
116.92
116.41
114.66
114.28
114.15
112.26
104.83
79.88
77.23
77.02
76.81
75.64
69.00
61.59
53.00
47.17
44.77
42.12
38.75
37.48
30.01
29.67
29.29
28.68
23.49
¹³C NMR (CDCl₃, 150 MHz)
NF500B
f1 (ppm)

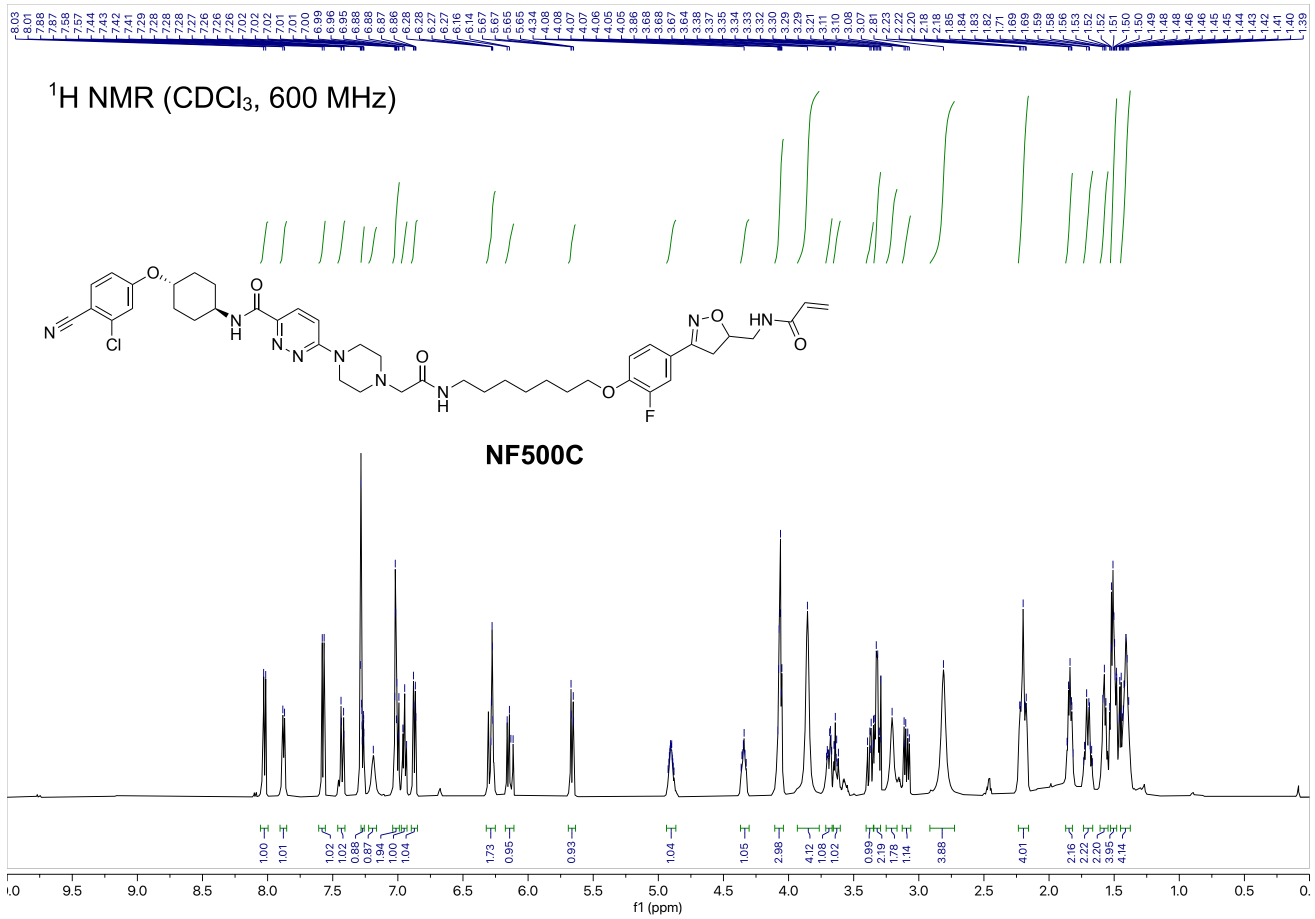
$^1$H NMR ($CDCl_3$, 600 MHz)
NF500C
f1 (ppm)

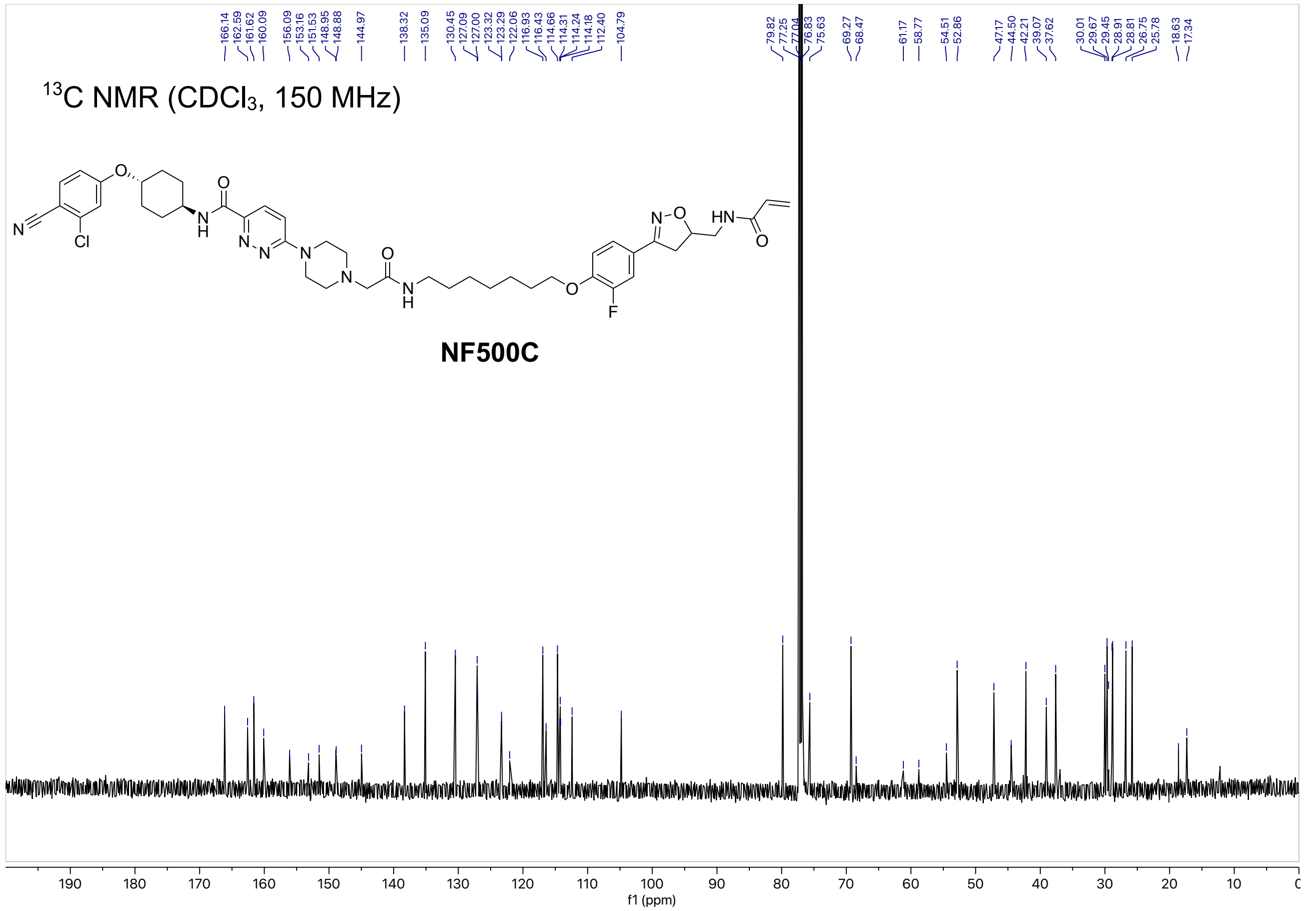
$^{13}$C NMR ($CDCl_3$, 150 MHz)
NF500C
f1 (ppm)

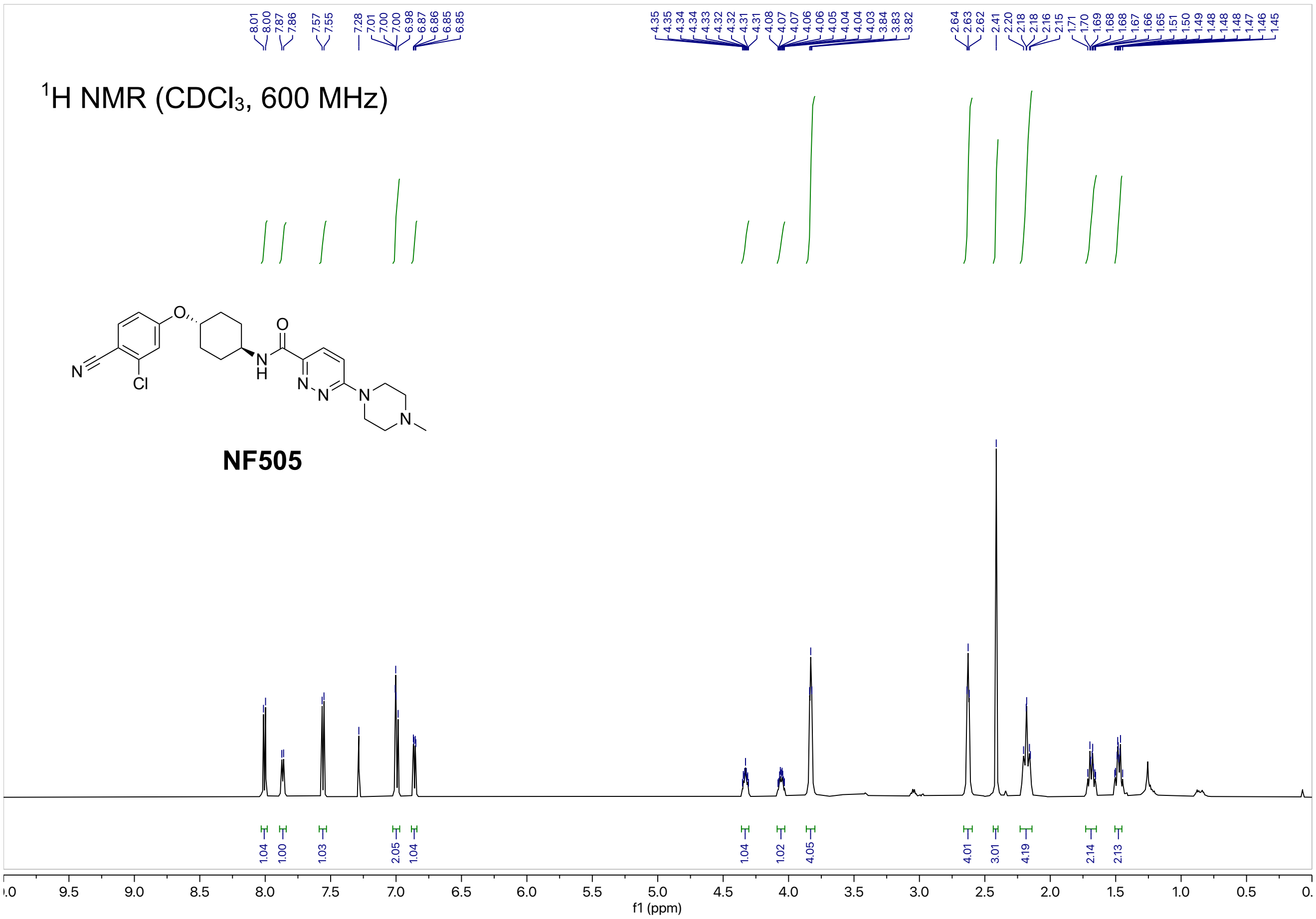

$^{1}$H NMR (CDCl$_3$, 600 MHz)
NF505
f1 (ppm)


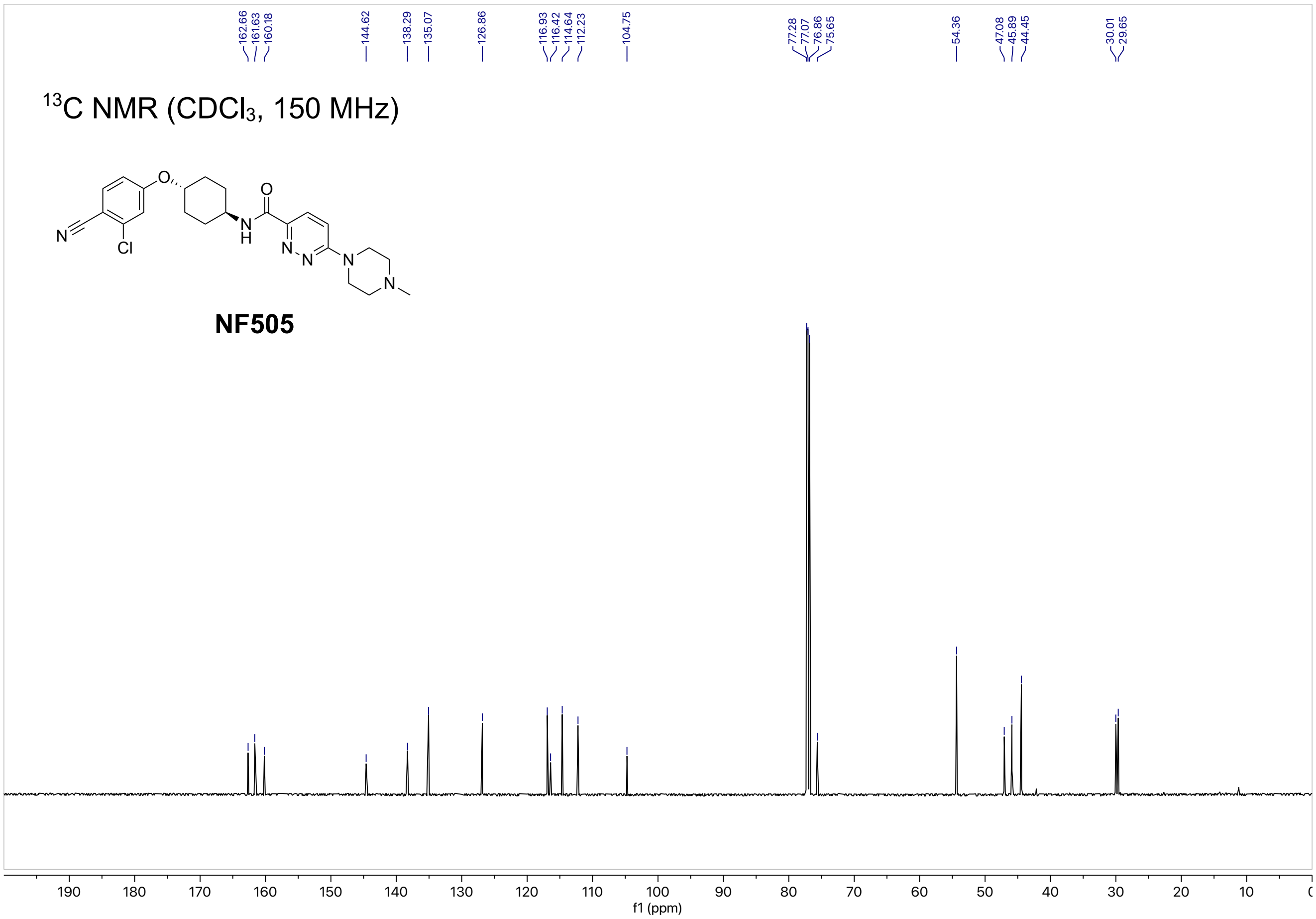

$^{13}$C NMR (CDCl$_3$, 150 MHz)
NF505
f1 (ppm)

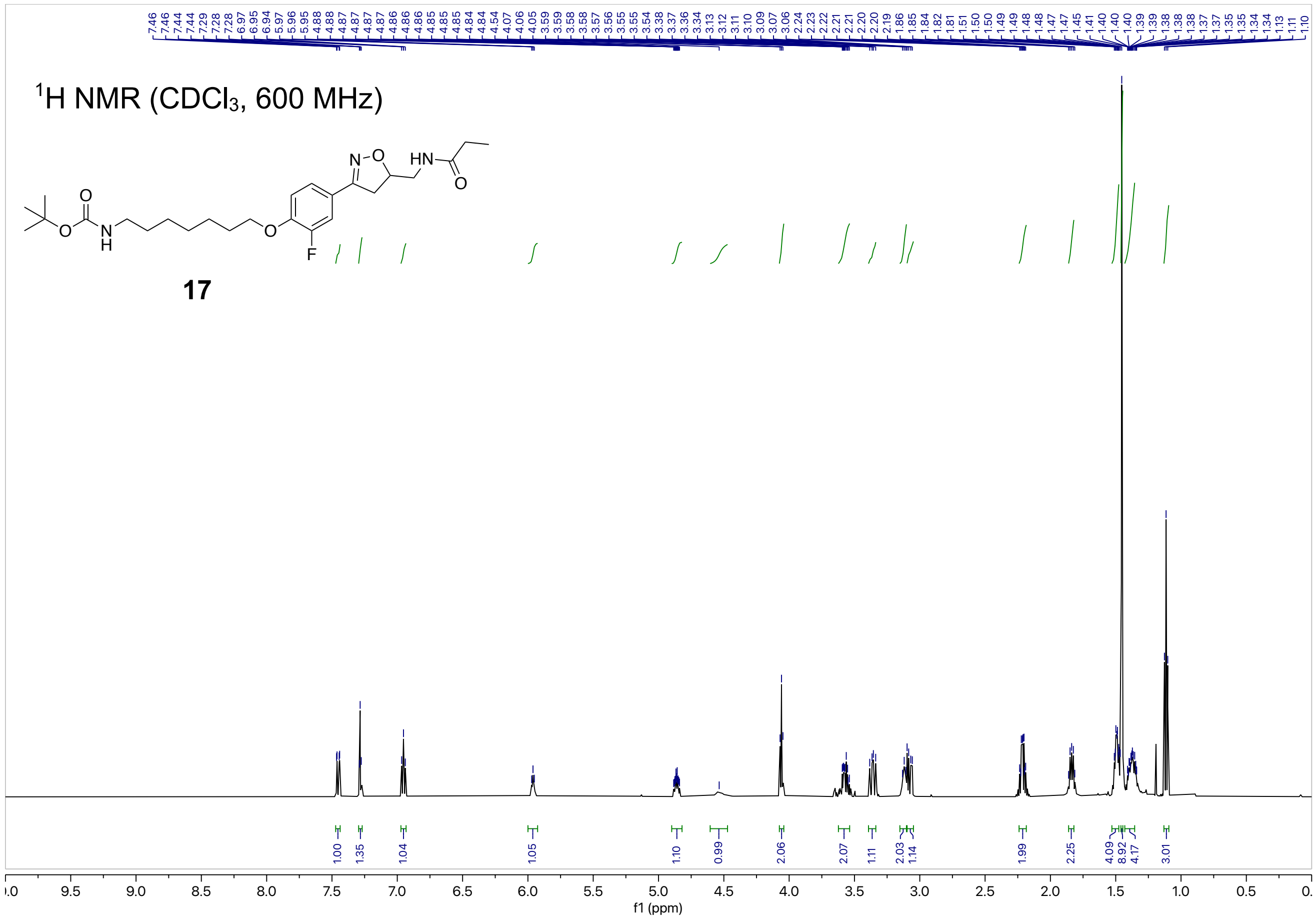
$^1$H NMR (CDCl$_3$, 600 MHz)
17
f1 (ppm)

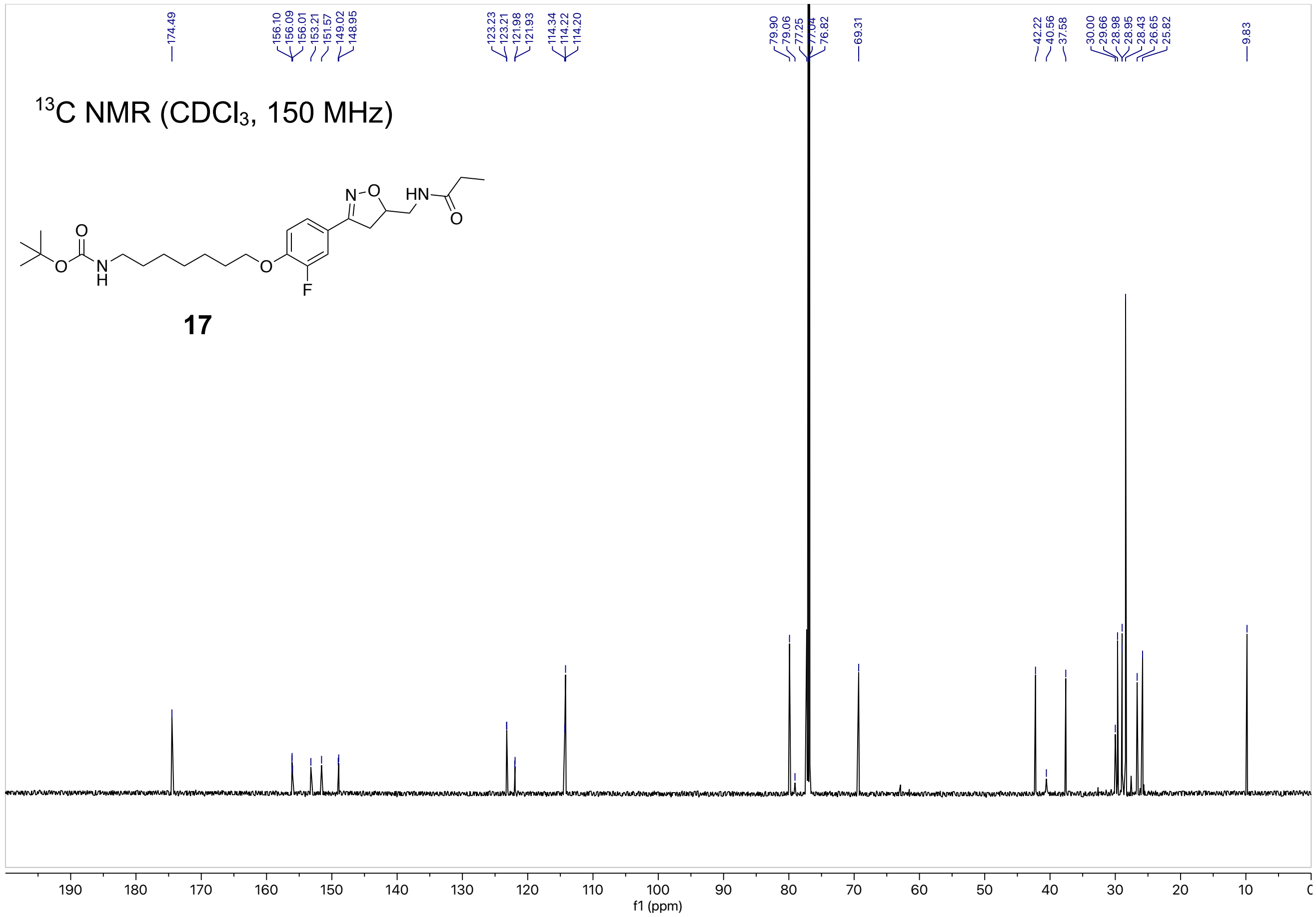
$^{13}$C NMR (CDCl$_3$, 150 MHz)
17
f1 (ppm)

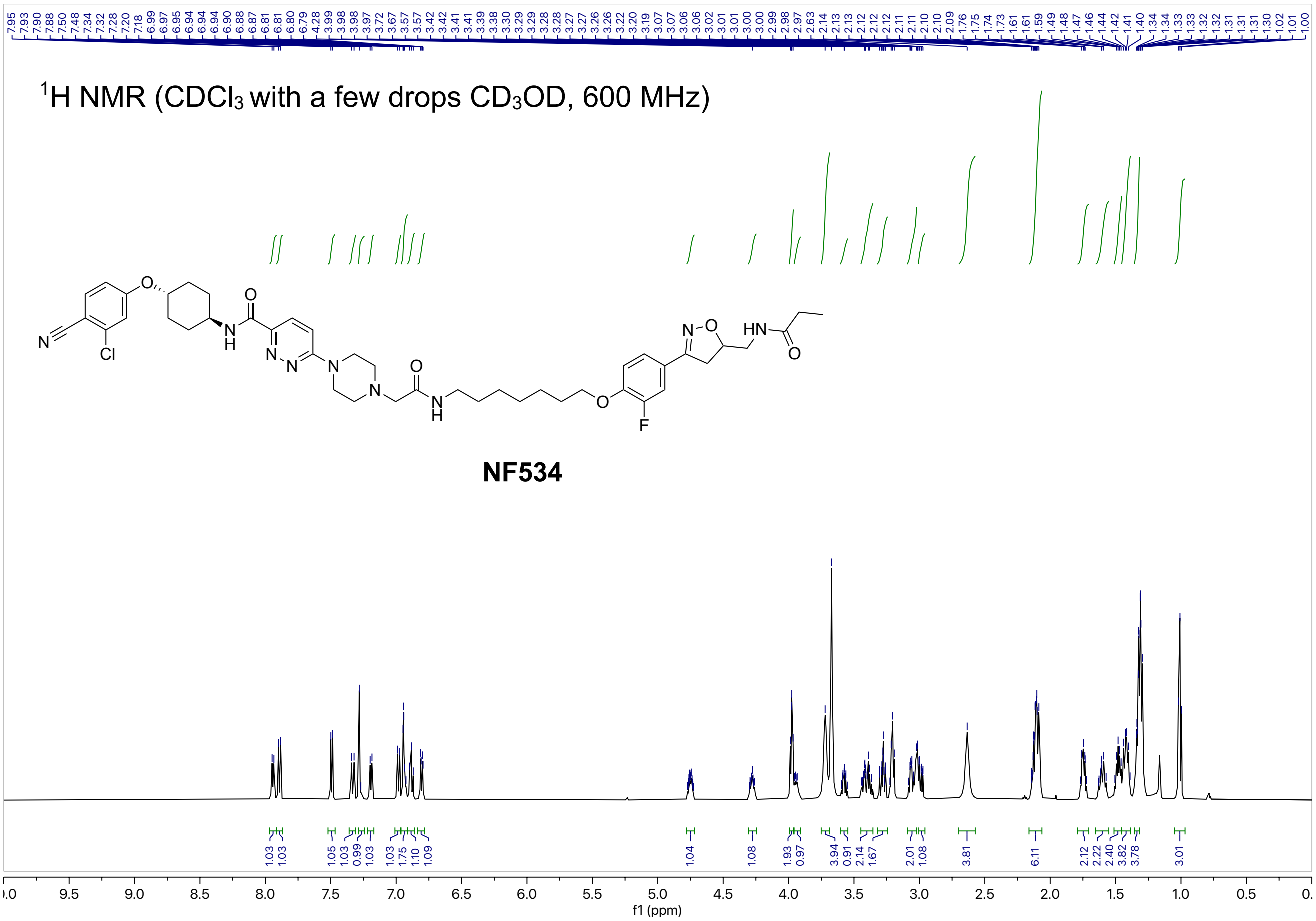
1H NMR (CDCl3 with a few drops CD3OD, 600 MHz)
NF534
f1 (ppm)

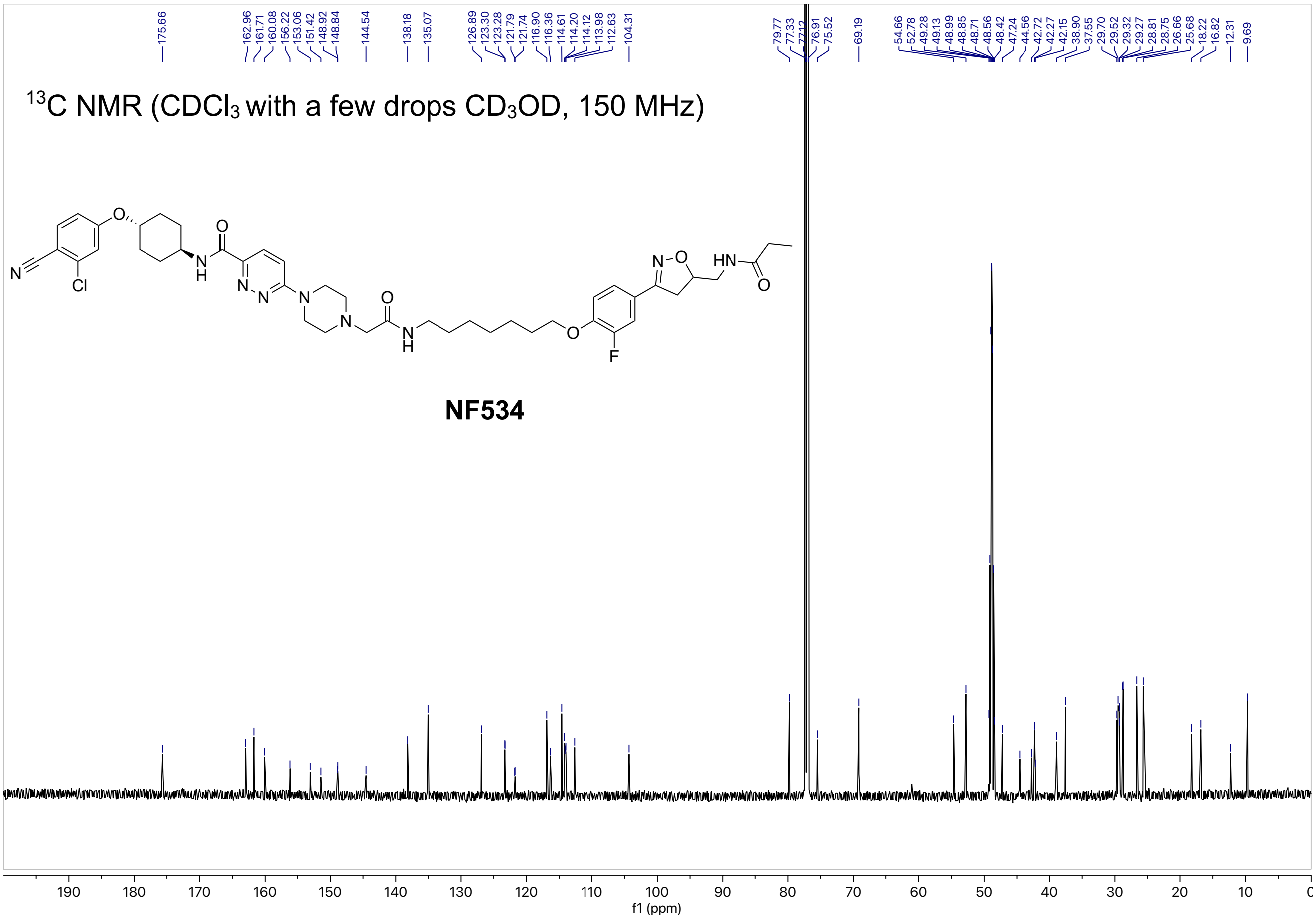
13C NMR (CDCl3 with a few drops CD3OD, 150 MHz)
NF534
f1 (ppm)
